# PhysioFit: a software to quantify cell growth parameters and extracellular fluxes

**DOI:** 10.1101/2023.10.12.561695

**Authors:** Loïc Le Grégam, Yann Guitton, Floriant Bellvert, Stéphanie Heux, Fabien Jourdan, Jean-Charles Portais, Pierre Millard

## Abstract

**Summary:** Quantification of growth parameters and extracellular uptake and production fluxes is central in systems and synthetic biology. Fluxes can be estimated using various mathematical models by fitting time-course measurements of the concentration of cells and extracellular substrates and products. A single tool is available to non-computational biologists to calculate extracellular fluxes, but it is hardly interoperable and is limited to a single hard-coded growth model. We present our open-source flux calculation software, PhysioFit, which can be used with any growth model and is interoperable by design. PhysioFit includes some of the most common growth models, and advanced users can implement additional models to calculate extracellular fluxes and other growth parameters for metabolic systems or experimental setups that follow alternative kinetics. PhysioFit can be used as a Python library and offers a graphical user interface for intuitive use by end-users and a command-line interface to streamline integration into existing pipelines.

**Availability and Implementation:** PhysioFit is implemented in Python 3 and was tested on Windows, Unix and MacOS platforms. PhysioFit is also freely available online at https://workflow4metabolomics.org. The source code, the data and the documentation are freely distributed under GPL3 license at https://github.com/MetaSys-LISBP/PhysioFit/ and https://physiofit.readthedocs.io/.

## Introduction

Quantification of growth rates and extracellular uptake and production fluxes is an essential task for addressing fundamental and applied questions in the fields of systems and synthetic biology, biotechnology, and health (Millard *et al*., 2023; Peiro *et al*., 2019; Hui *et al*., 2020). Estimating extracellular fluxes involves the use of mathematical models to fit time-course measurements of cell concentrations and extracellular substrates and products (Peiro *et al*., 2019; Murphy and Young, 2013). Several models have been proposed to relate these dynamic measurements to extracellular fluxes and other growth parameters (such as growth rates or lag times) (Baranyi and Roberts, 1994; Nielsen and Villadsen, 1992; Poccia *et al*., 2014). However, in most cases, model-based flux quantification is performed using custom-made scripts, which can be error-prone and lack the robustness and reproducibility necessary for comprehensive flux studies. Currently, there is only one software tool available to biologists for calculating extracellular fluxes, namely, Extracellular Time-Course Analysis (ETA) (Murphy and Young, 2013). While ETA represents a significant step toward more accessible metabolic studies, it is limited by a single hard-coded growth model, a reliance on a proprietary programming language, and a lack of interoperability with other upstream and downstream computational tools. The open-source Python package pyFOOMB (Hemmerich *et al*., 2021) has been developed to overcome some of these limitations, allowing users to build and analyze their own flux models using efficient numerical methods. However, pyFOOMB can only be used programmatically and lacks certain features that would make extracellular flux studies more accessible and reproducible for biologists, such as statistical methods to evaluate goodness-of-fit and identify the most appropriate flux models, a graphical user interface, or a file logger to save calculation parameters and process information along with the results. This deficiency in user-friendly, versatile software significantly constrains the scope, throughput, and reproducibility of metabolic flux studies.

Here, we present PhysioFit, a Python tool designed to be interoperable and user-friendly, allowing biologists with no prior computational experience to use it effectively. PhysioFit comes with common growth models and can be extended with additional models to calculate extracellular fluxes and other growth parameters for metabolic systems or experimental setups that follow alternative kinetics.

### Methods and implementation

The overall process implemented in PhysioFit is shown in Figure 1. After loading the required input data containing the time-course concentrations of biomass and extracellular metabolites (provided as a *tsv* file), the user can select an appropriate growth model and processing parameters (number of Monte-Carlo iterations, initial values and bounds on model parameters, and standard deviations on measurements). Extracellular fluxes and growth parameters included in the model are then estimated by fitting the measured concentrations. All parameters *p* are estimated by minimizing the objective function *c* defined as the weighted sum of squared errors:

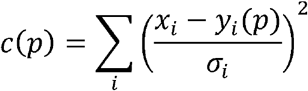

where *x*_*i*_ is the experimental value of data point *i*, with an experimental standard deviation *σ*_*i*_, and *y*_*i*_(*p*) is the corresponding value simulated by the model. The objective function *c* is minimized with the differential evolution optimization algorithm (Storn and Price, 1997), and the best solution is refined using the L-BFGS-B method (Byrd *et al*., 1995) to ensure convergence to a local optimum. Confidence intervals on fluxes and growth parameters are estimated using a Monte-Carlo analysis, and plots are generated for visual inspection of the fitted profiles. The goodness-of-fit is evaluated based on a χ^2^ statistical test, and the Akaike information criteria (both classical and corrected, AIC and AICc) (Symonds and Moussalli, 2011) allow users to compare and rank different models, as explained in the next section. Finally, PhysioFit generates *i*) a *tsv* file containing the estimated fluxes and growth parameters, *ii*) a *txt* file containing the statistical results (χ^2^ test and AIC), *iii*) a yaml configuration file containing the run parameters, *iv*) *pdf* and *svg* plots of simulation and measurements, and v) a *log* file containing detailed information on the calculation process. These files allow users to quickly assess the quality of the calculated fluxes and contain all necessary information to share repeatable extracellular flux analyses, thereby enhancing the quality and reproducibility of flux studies.

**Figure 1.**
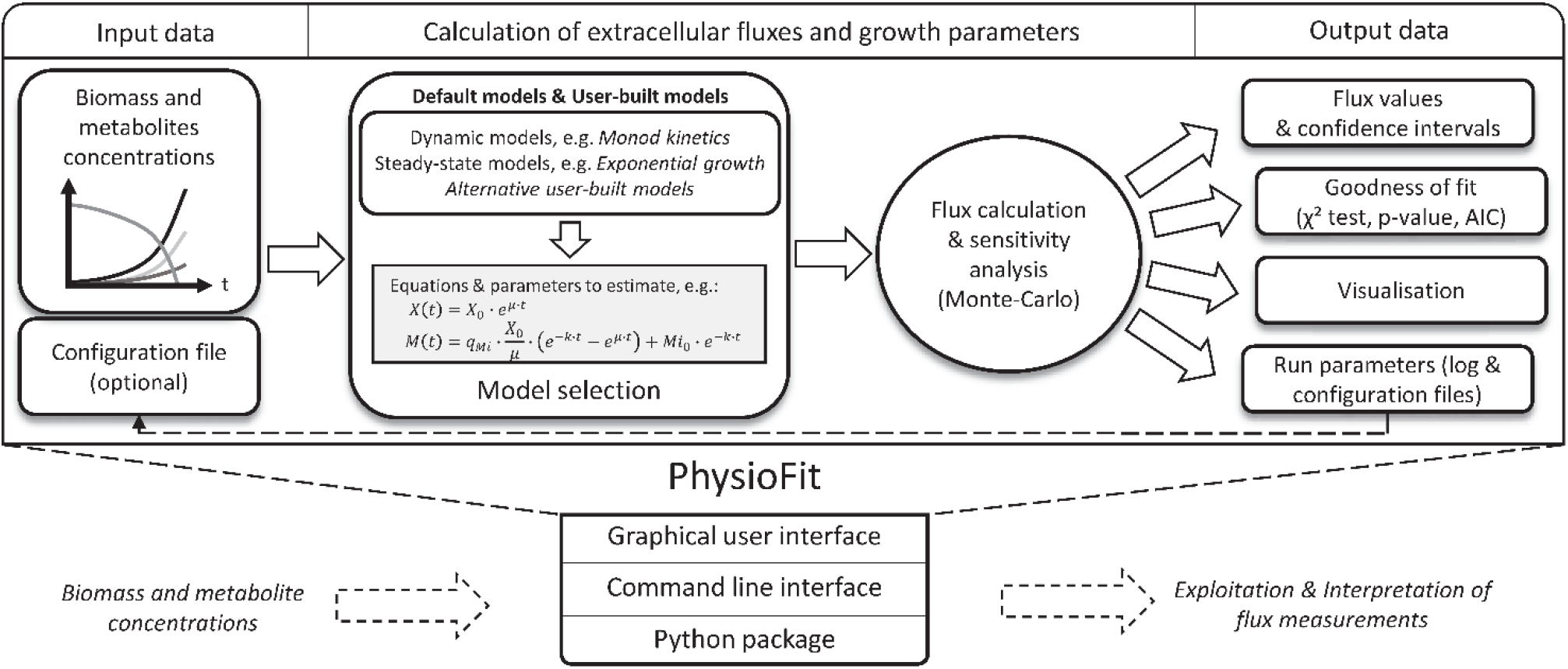
Overview of PhysioFit. Input data (time-course concentrations of biomass and extracellular metabolites) are fitted using a model (selected by the user), and flux calculation results are returned with associated statistics and plots. Different interfaces are available to enable utilization of PhysioFit in computational workflows.

## Flux models

A flux model must contain equations that describe the dynamics of biomass and metabolite concentrations (used for simulations) and the list of all parameters with their default values and bounds (used for optimization). These equations are typically expressed as:

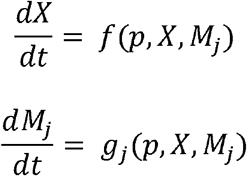

where *X* is the biomass concentration, *M*_*j*_ is the concentration of the metabolite *j*, and *p* is the vector of parameters. For instance, during exponential growth of *Escherichia coli* on dihydroxyacetone (DHA), a carbon source that is subject to non-enzymatic degradation (Peiro *et al*., 2019), the functions *f* and *g* correspond to:

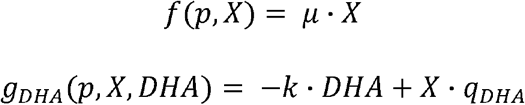

with parameters *μ* (growth rate), *k* (degradation constant of DHA) and *q*_*DHA*_ (DHA uptake flux). Note that PhysioFit can solve ODEs-based models using classical numerical methods or, when known, using analytical solutions, the latter of which can substantially reduce calculation times.

As detailed in the Supplementary information and in the documentation, PhysioFit comes with common flux models, including a dynamic Monod model and four steady-state models of exponential growth. These models cover typical growth experiments conducted in metabolic studies. Advanced users can create additional models, with a model template provided with PhysioFit and a detailed tutorial on model construction available in the documentation.

Since different models can be used to fit a dataset, we provide guidelines for model comparison to help users select the most appropriate model. Briefly, candidate models that differ in terms of structure or complexity can be used to fit the data and then compared based on statistical results (Symonds and Moussalli, 2011). The model with the lowest AIC value is considered the best-fitting model among the candidates. However, it is crucial to consider the differences in AIC values between models, as models with low ΔAIC values (typically < 2) are considered to have similar support from the data. Users can follow the guidelines detailed in the documentation to ensure that the selected model optimally balances goodness-of-fit and model complexity for reliable extracellular flux calculations.

### Validation in the context of high-throughput flux studies

We have implemented unit tests to validate the main features of PhysioFit, including i) data loading, ii) model initialization, iii) simulation and iv) parameter estimation. In particular, each model is tested by i) comparing a synthetic dataset (simulated from known parameters using analytical solutions or via the pyFOOMB package) to the dataset simulated by PhysioFit from the same parameters and ii) comparing the parameters estimated by PhysioFit from the synthetic dataset to the parameters used for simulations. For all the models, relative differences between the expected and calculated dynamics and parameters remain below 1 %.

We have also conducted external validations by calculating extracellular fluxes and growth rates from datasets taken from the literature. The complete validation dataset, which covers the different models included with PhysioFit, contains a total of 223 growth experiments carried out on wild-type and mutant *E. coli* and *S. cerevisiae* strains – 197 experiments from (Bergès *et al*., 2021), 25 experiments from (Peiro *et al*., 2019), 1 experiment from (Zentou *et al*., 2019). All calculations were performed in a few minutes, confirming the applicability of PhysioFit for high-throughput extracellular flux studies. The fluxes and other parameters estimated by PhysioFit are in good agreement with the expected flux values (r^2^ > 0.99).

Detailed validation results can be found in the Supplementary information. Overall, these tests on synthetic and real-world datasets validate both the software and the models implemented.

### Interfaces and interoperability

PhysioFit can be used by biologists with no prior computational experience via a Graphical User Interface. Moreover, since extracellular flux analyses are often part of larger computational workflows (e.g. for intracellular flux calculations by Flux Balance Analysis or by ^13^C-fluxomics), PhysioFit has been designed to maximize its interoperability with upstream and downstream tools. PhysioFit can be used directly as a Python module or via a Command-Line Interface and uses standard input and output formats that facilitate data exchange with other tools. As a step towards the development of automated, high-throughput fluxomics workflows, we have implemented PhysioFit on Workflow4Metabolomics (Giacomoni *et al*., 2015) (https://workflow4metabolomics.usegalaxy.fr), a collaborative portal for the metabolomics and fluxomics community. Here, it can be used as a standalone tool and easily integrated into user-made workflows.

## Conclusion

Here we present PhysioFit, a highly versatile tool that enables the calculation of extracellular fluxes using any growth model with a user-friendly graphical interface. A command line interface and a Python package also enable integrating PhysioFit into metabolomics and fluxomics workflows. PhysioFit’s modularity makes it an ideal tool for conducting quantitative studies of virtually any biological system, from microbial to higher cells. PhysioFit will benefit a wide range of fields, including systems biology, synthetic biology, and biotechnology.

## Acknowledgements

We thank Lindsay Peyriga and Cécilia Bergès (MetaToul-MetaboHUB, Toulouse, France) for testing the software, and Sophie Colombié and Sylvain Prigent (INRAE, Bordeaux, France) for insightful discussions.

