## Supplementary material for "PhysioFit: a software to quantify cell growth parameters and extracellular fluxes": Supplementary_information.pdf

**-**

To validate PhysioFit, we implemented five commonly used flux models to quantify extracellular fluxes. This document describes the models included by default with PhysioFit, the testing strategy employed, and the validation results obtained.

### 1. Models

PhysioFit includes flux models that cover classical situations encountered in metabolic studies. Advanced users can build additional models by following the guidelines provided at PhysioFit Documentation. We also offer support at our GitHub repository. If you have developed a new model, it might be useful and valuable to the fluxomics community! Please keep in touch with us to discuss the possibility of including your model among the built-in models shipped with PhysioFit.

#### a. Steady-state models

PhysioFit includes different steady-state models of exponential growth, where fluxes remain constant over time, as typically observed in batch experiments. A first model, which can account for i) non-enzymatic degradation of certain metabolites and ii) growth lag, is described by the following system of ordinary differential equations:

$$\frac{dX}{dt} = \begin{cases} 0 \ if \ t < t_{lag} \\ \mu \cdot X \ else \end{cases} \qquad (eq. 1)$$

$$\frac{dM_i}{dt} = \begin{cases} -k \cdot M_i \ if \ t < t_{lag} \\ -k \cdot M_i + X \cdot q_{M_i} \ else \end{cases} \qquad (eq. 2)$$

where $t$ is time, μ is growth rate, $X$ is the biomass concentration, $t_{lag}$ is the lag time, $k$ is the first-order degradation constant of the metabolite $M_i$ and $q_{M_i}$ is its exchange (uptake or production) flux. Note that since $q_{M_i}$ is positive (negative) when $M_i$ is produced (consumed), the sign of $q_{M_i}$ can be used to automatically identify products and substrates.

Integrating equations 1-2 provides the following analytical functions:

$$X(t) = \begin{cases} X_0 \ if \ t < t_{lag} \\ X_0 \cdot e^{\mu \cdot (t - t_{lag})} \ else \end{cases} \qquad (eq. 3)$$

$$M_i(t) = \begin{cases} M_i^0 \cdot e^{-k \cdot t} \ if \ t < t_{lag} \\ q_{M_i} \cdot \frac{X_0}{\mu + k} \cdot \left(e^{\mu \cdot (t - t_{lag})} - e^{-k \cdot (t - t_{lag})}\right) + M_i^0 \cdot e^{-k \cdot t} \ else \end{cases} \qquad (eq. 4)$$

Three additional models are derived from this general model (without degradation, without lag phase, and without degradation nor lag phase).

In the absence of a lag phase (i.e. $t_{lag} = 0$), equations 3-4 simplifies to:

$$X(t) = X_0 \cdot e^{\mu \cdot t} \qquad (eq. 5)$$

$$M_i(t) = q_{M_i} \cdot \frac{X_0}{\mu + k} \cdot \left(e^{\mu \cdot t} - e^{-k \cdot t}\right) + M_i^0 \cdot e^{-k \cdot t} \qquad (eq. 6)$$

In the absence of degradation (i.e. $k = 0$), equation 4 simplifies to:

$$M_i(t) = \begin{cases} M_i^0 \ if \ t < t_{lag} \\ q_{M_i} \cdot \frac{X_0}{\mu} \cdot \left(e^{\mu \cdot (t - t_{lag})} - 1\right) + M_i^0 \ else \end{cases} \qquad (eq. 7)$$

In the absence of both degradation and lag (i.e. $t_{lag} = 0$ and $k = 0$), equations 3-4 simplifies to:

$$X(t) = X_0 \cdot e^{\mu \cdot t} \qquad (eq. 8)$$

$$M_i(t) = q_{M_i} \cdot \frac{X_0}{\mu} \cdot (e^{\mu \cdot t} - 1) + M_i^0 \qquad (eq. 9)$$

For simulations, these models are solved using directly the analytical solutions.

#### a. Dynamic model

We have also implemented a dynamic model where fluxes and growth rate are represented by Monod kinetics, for one substrate and one product. In this model, the time course concentrations of biomass (X), substrate (S) and product (P) are described by the following system of ordinary differential equations:

$$\frac{dS}{dt} = -X \cdot q_S \qquad (eq. 10)$$

$$\frac{dP}{dt} = q_S \cdot product\_yield \qquad (eq. 11)$$

$$\frac{dX}{dt} = q_S \cdot biomass\_yield \qquad (eq. 12)$$

where $q_S$ is the substrate uptake flux. The dependence of $q_S$ on the substrate concentration is expressed by the Monod rate law:

$$q_S = qS_{max} \cdot \frac{S}{K_M + S} \qquad (eq. 13)$$

where $qS_{max}$ is the maximal substrate uptake rate and $K_M$ is the "half-velocity constant" (the value of at which $q_S = 2 \cdot qS_{max}$).

For simulations, this model is solved by numerical methods, using the *lsoda* solver of Scipy (https://docs.scipy.org/doc/scipy/reference/generated/scipy.integrate.LSODA.html).

### 2. Test strategy

We used two complementary approaches to validate PhysioFit:

- **Unit Tests**: These were used to validate the main functions of PhysioFit, including i) data loading, ii) model initialization, iii) simulation, iv) parameter estimation, and v) sensitivity analysis. Specifically, we validated the core features of PhysioFit and the default models using synthetic datasets simulated from known parameters.
- **Flux Calculation from Real-World Datasets**: This approach involved comparing the results obtained with PhysioFit to published extracellular fluxes and growth rates to ensure accuracy and reliability.

### 3. Validation using synthetic data

For each model, we have generated a synthetic dataset from known parameters using analytical solutions (see, for example, equations 8-9) or via the pyFOOMB package. Using these parameters and the corresponding dataset, we have designed two unit tests per model:

- Comparison of the synthetic dataset to the dataset simulated by PhysioFit from the parameters used for simulations.
- Comparison of the parameters estimated by PhysioFit from the synthetic dataset to the parameters used for simulations.

The set of parameters and the corresponding biomass and metabolite dynamics can be found in the test folder of PhysioFit repository here. For all the models, relative differences between the expected and calculated dynamics and parameters remain below 1 %, thereby validating the models and the simulation and parameter estimation routines of PhysioFit.

### 4. Validation using experimental data

We have also validated PhysioFit using data taken from previous extracellular flux studies. The complete validation dataset contains a total of 223 growth experiments of wild-type and mutant *E. coli* and *S. cerevisiae* strains (197 experiments from Bergès et al., 25 experiments from Peiro et al., 1 experiment from Zentou et al.) that cover different models provided with PhysioFit. The three validation datasets are available from the original publications and can be accessed here, where we also provide all the calculation results. As detailed below, fluxes and other parameters estimated by PhysioFit are in good agreement with the published values ($r^2 > 0.99$).

#### a. Data from Bergès et al. (2021)

We implemented the steady-state model described by equations 8-9 to quantify the glucose uptake, acetate production and growth rates of 180 wild-type and mutant *Escherichia coli* strains grown on glucose in M9 medium, as detailed in (Berges, et al., 2021). The time-course concentrations of glucose (Glc), acetate (Ace) and biomass (X) were measured in 197 growth experiments.

Experimental data and simulation results for the best fit of a replicate of the wild-type strain are shown below as example. For all the strains, simulations are in good agreement with the experimental data.

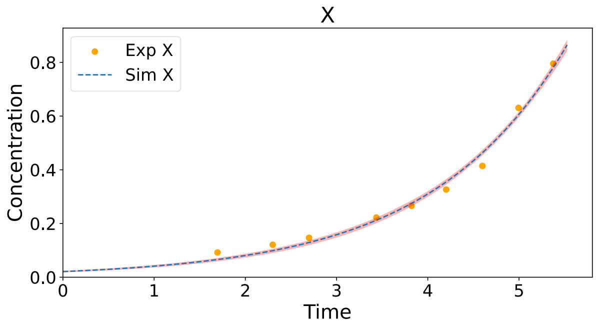


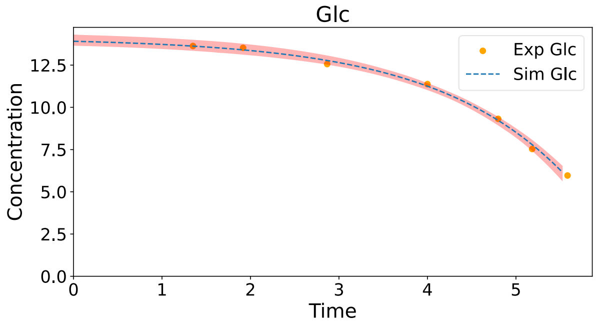


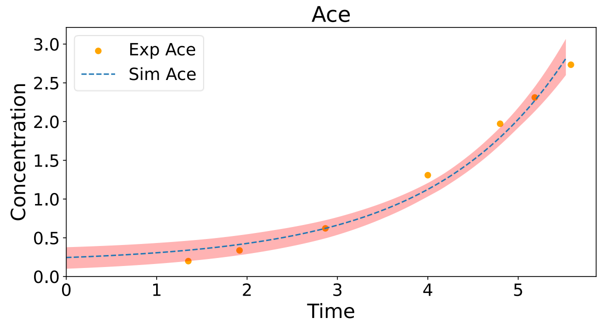


As shown below, the published and estimated parameters - initial concentration of each species (*X_0*, *Glc_0*, *Ace_0*) and fluxes (growth rate; glucose uptake rate, *Glc_q*; and acetate production rate, *Ace_q*) - are in excellent agreement, with $r^2 > 0.99$ for the 197 experiments.

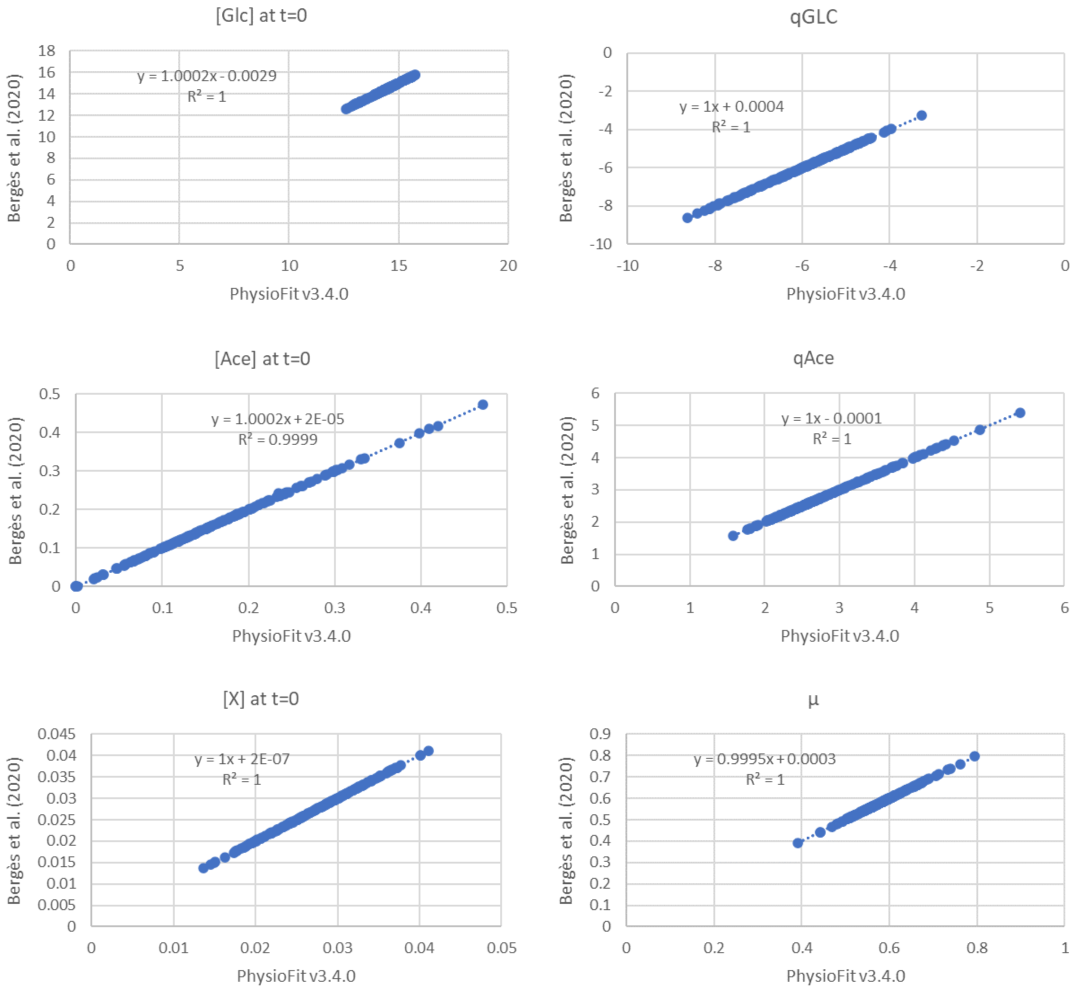


### b. Data from Peiro et al. (2019)

As a second example, we implemented and tested the model with lag and (non-enzymatic) degradation of carbon sources such as DHA or glutamine, as described by equations 1-2. The experimental data taken from (Peiro, et al., 2019) contains a total of 25 growth experiments from 6 (wild type and mutant) *E. coli* strains grown on DHA. The degradation constant for DHA was set to $8.64 \cdot 10^{-3}$, as determined experimentally in the publication. For all strains, we determined the growth and DHA uptake rates. For strains that produce glycerol, the glycerol production rate was also quantified. Simulation results for the best fit obtained for the Δ*gldA* strain (see below) are in good agreement with the experimental data.

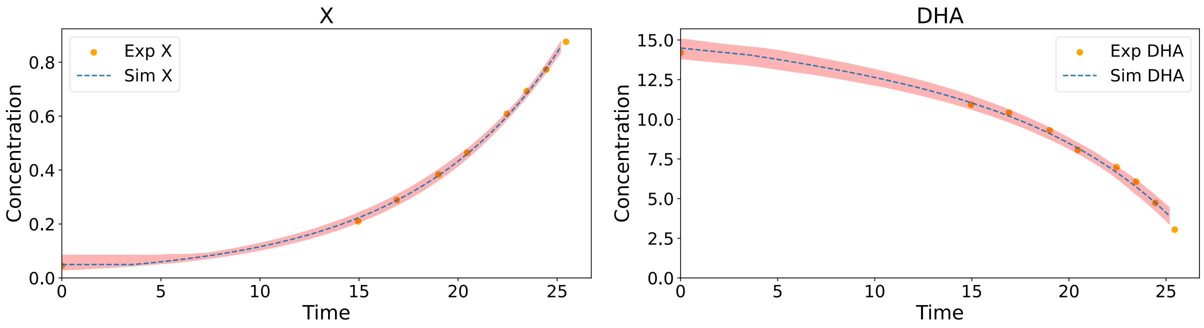


As shown below, the published and estimated parameters - initial concentration of each species (*X_0*, *DHA_0*, *GLY_0*), fluxes (growth rate; DHA uptake rate, *DHA_q*; glycerol production rate, *GLY_q*) and lag time - are in good agreement, with $r^2 > 0.99$ for the 25 experiments.

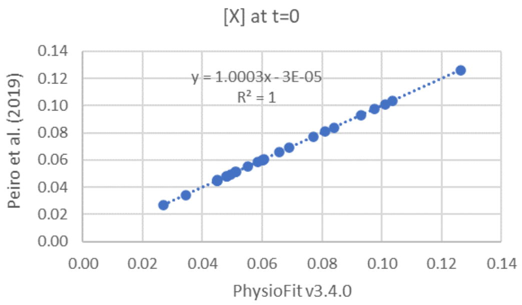


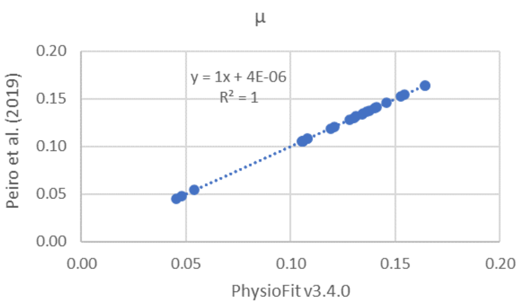


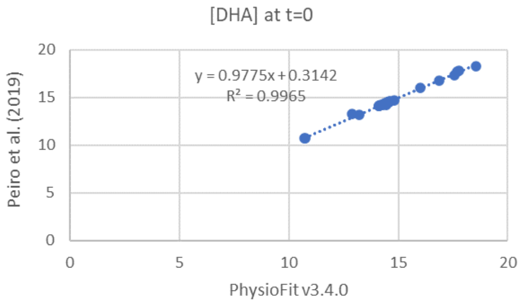


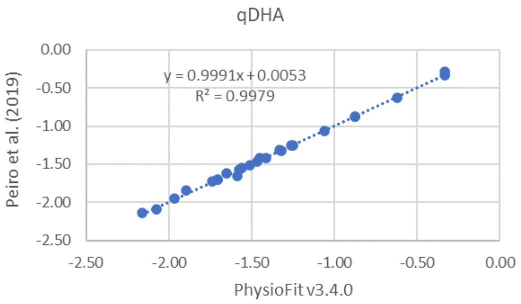


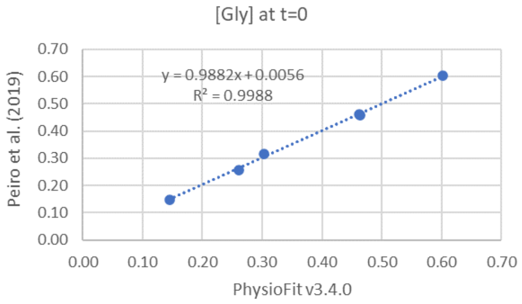


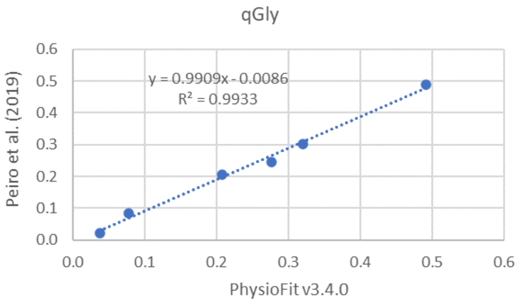


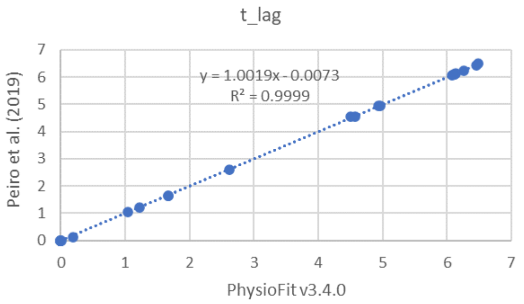


#### c. Dynamic (Monod) model

We also implemented a dynamic model where fluxes and growth are represented by Monod kinetics, as detailed in (Zentou, et al., 2019) and described by equations 10-13. This model was used to estimate the biomass and ethanol yields of the yeast *Saccharomyces cerevisiae* during a batch fermentation of molasses.

Simulation results for the best fit (see below) are in good agreement with the experimental data.

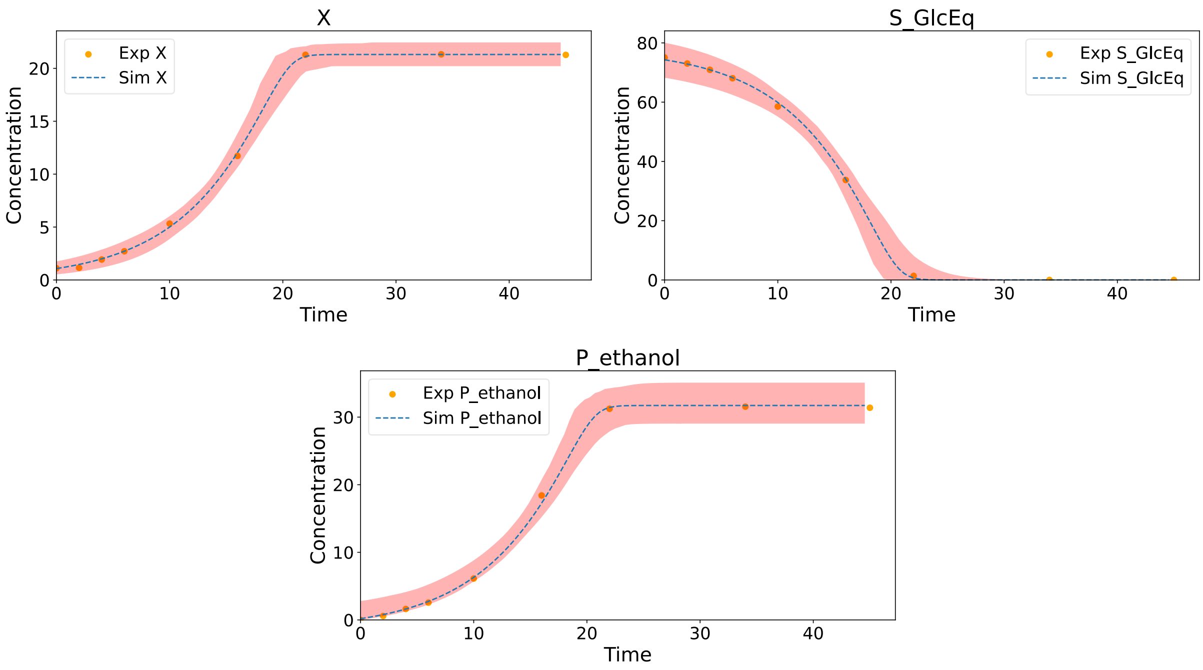


The initial concentration of each species (*X_0*, *S_GlcEq_s_0*, *P_ethanol_p_0*) and the growth parameters (biomass yield, *y_BM*; ethanol yield, *P_ethanol_y_P*; affinity for the substrate, *S_GlcEq_km*; maximal substrate uptake rate, *S_GlcEq_qsmax*) are provided below with their associated confidence intervals.

| | **PhysioFit v3.4.0** | | | | | | **Zentou et al., 2019** | |
|---|---|---|---|---|---|---|---|---|
| **parameter** | **optimal** | **mean** | **sd** | **median** | **CI_2.5** | **CI_97.5** | **val** | **sd** |
| X_0 | 1.076 | 1.094 | 0.325 | 1.069 | 0.527 | 1.752 | *n.a.* | *n.a.* |
| y_BM | 0.272 | 0.274 | 0.016 | 0.273 | 0.247 | 0.308 | 0.286 | *n.a.* |
| S_GlcEq_km | 8.253 | 11.781 | 20.229 | 8.224 | 1.000 | 43.689 | *n.a.* | *n.a.* |
| S_GlcEq_qsmax | 0.632 | 0.673 | 0.227 | 0.622 | 0.479 | 1.088 | *n.a.* | *n.a.* |
| S_GlcEq_s_0 | 74.297 | 74.057 | 3.183 | 74.006 | 68.229 | 80.061 | 75 | *n.a.* |
| P_ethanol_y_P | 0.424 | 0.417 | 0.033 | 0.417 | 0.354 | 0.478 | 0.431 | *n.a.* |
| P_ethanol_p_0 | 0.182 | 0.887 | 0.912 | 0.712 | 0.001 | 2.783 | *n.a.* | *n.a.* |

*n.a.*: not available in the original publication

Parameters obtained with PhysioFit are consistent with the available published values.
