## Supplementary figures and images for "PhysioFit: a software to quantify cell growth parameters and extracellular fluxes"

### KEIO_ROBOT1_3.pdf

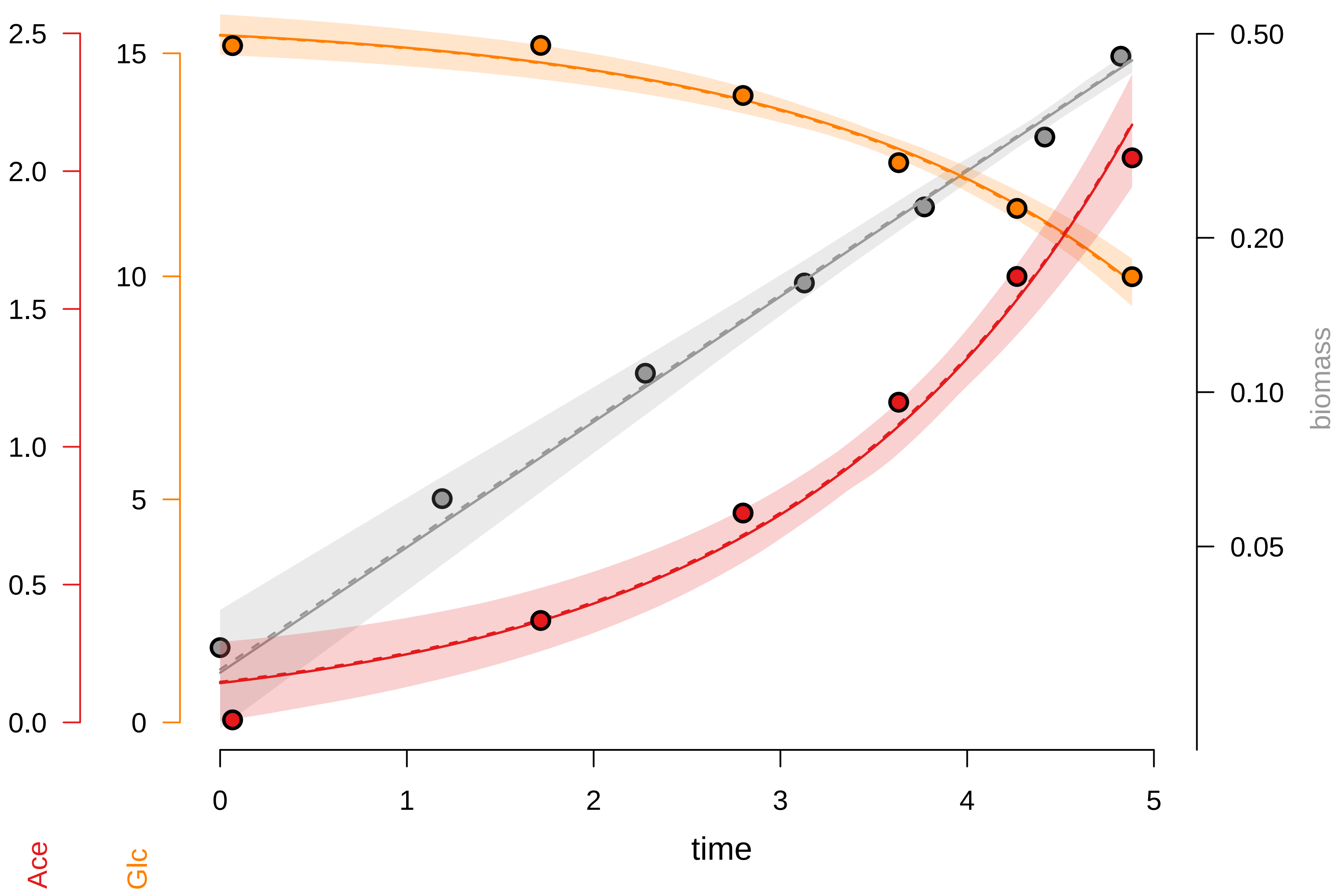

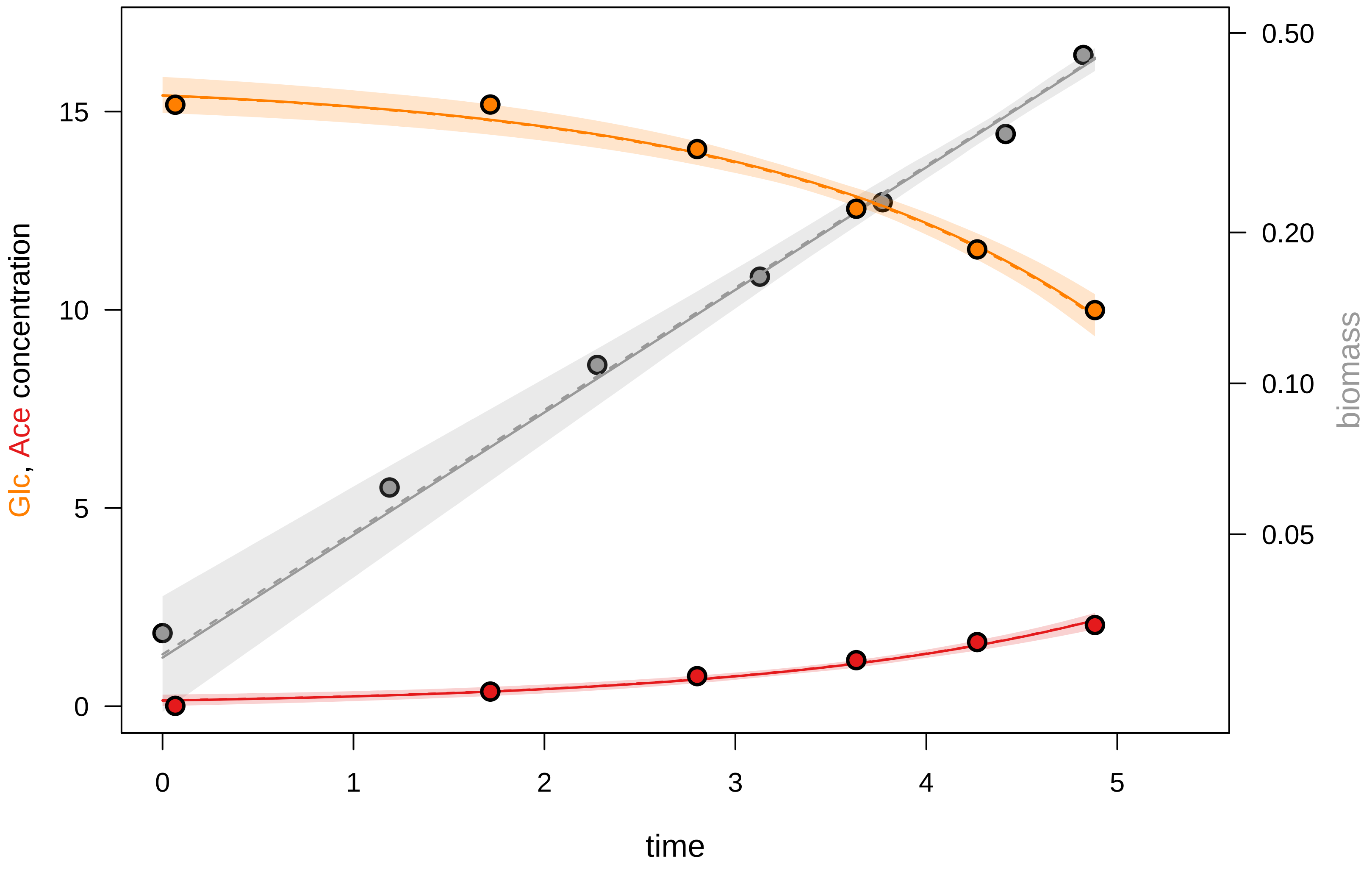

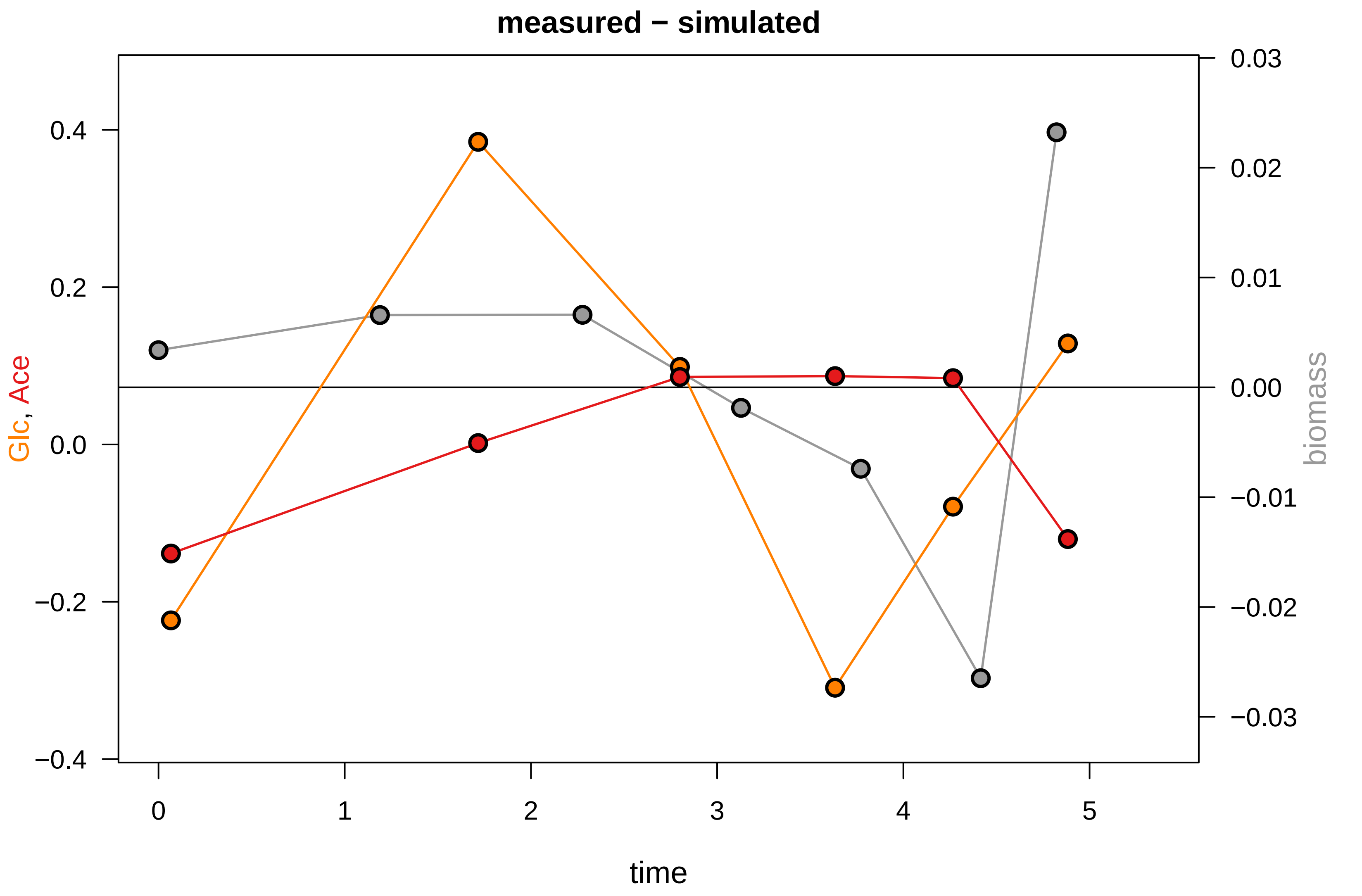

### KEIO_ROBOT1_27.pdf

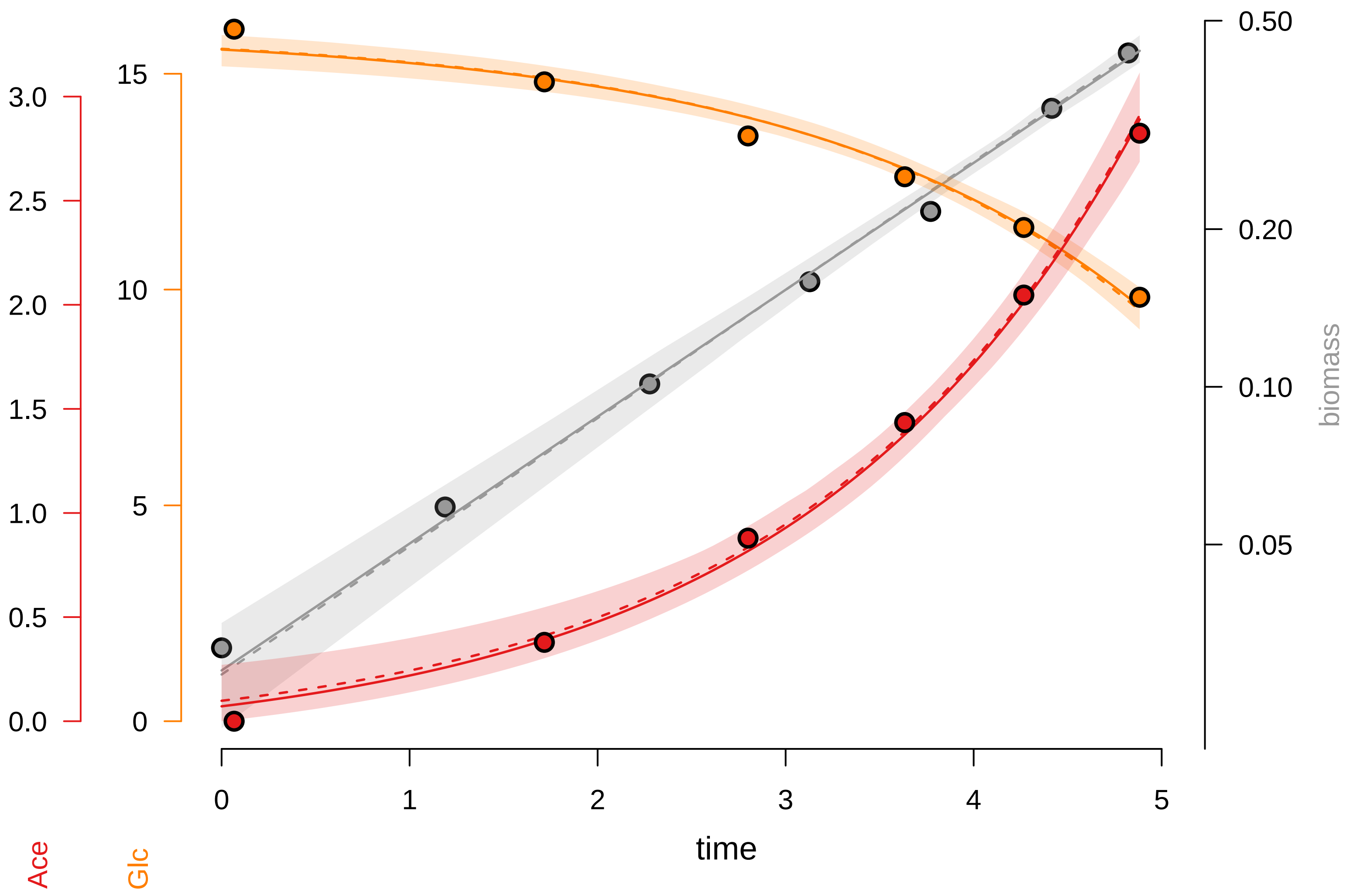

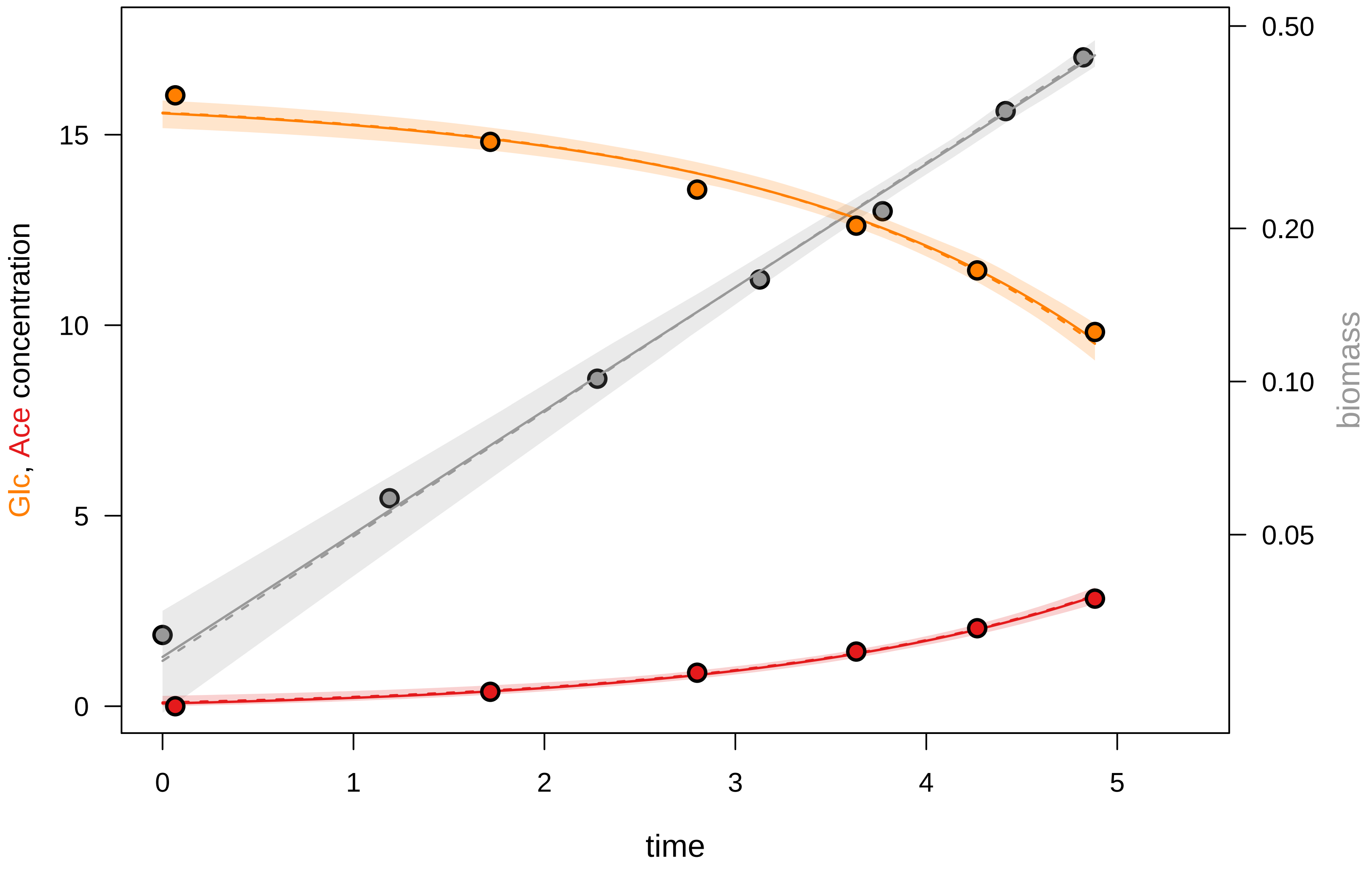

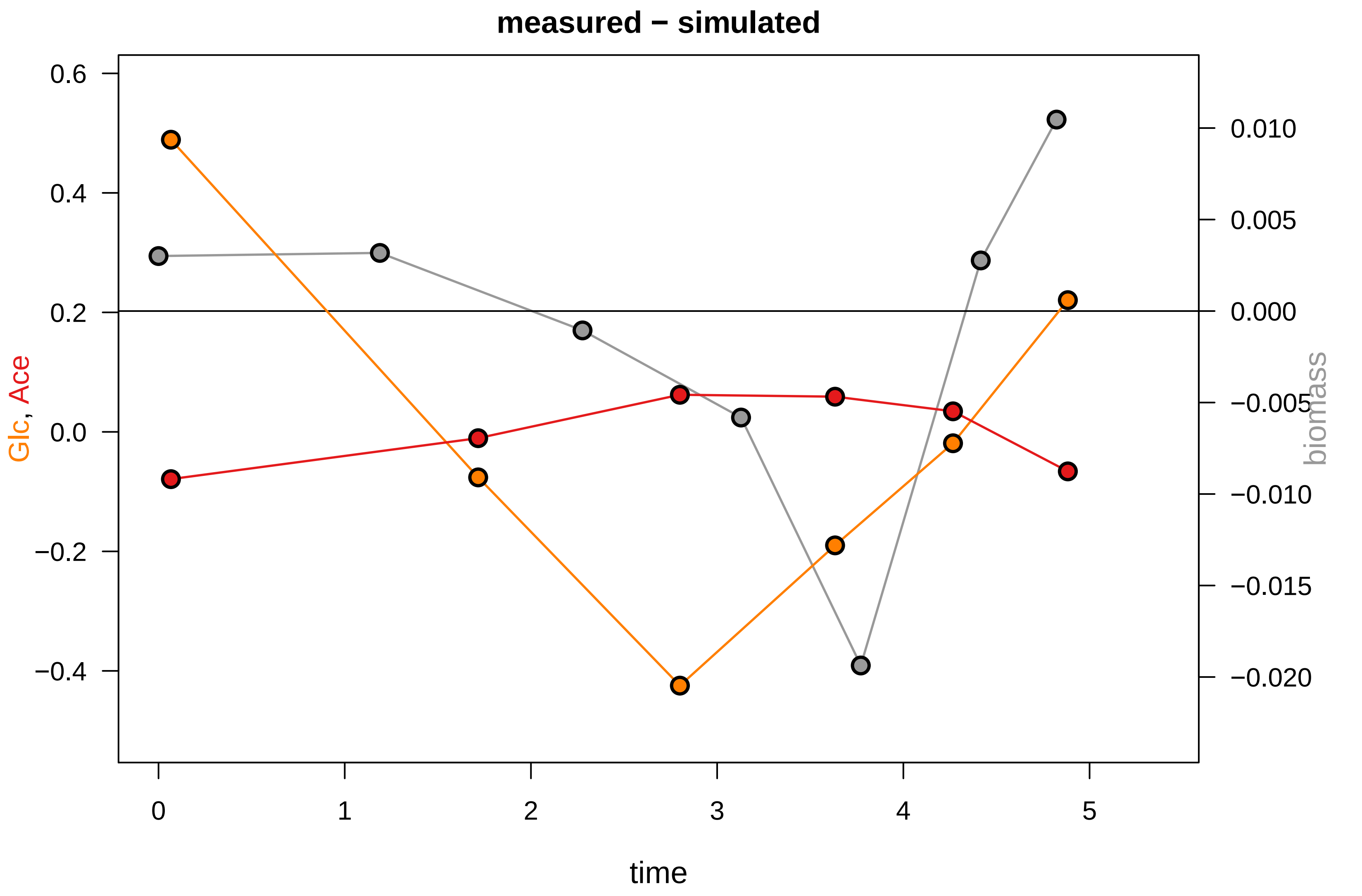

### KEIO_ROBOT1_34.pdf

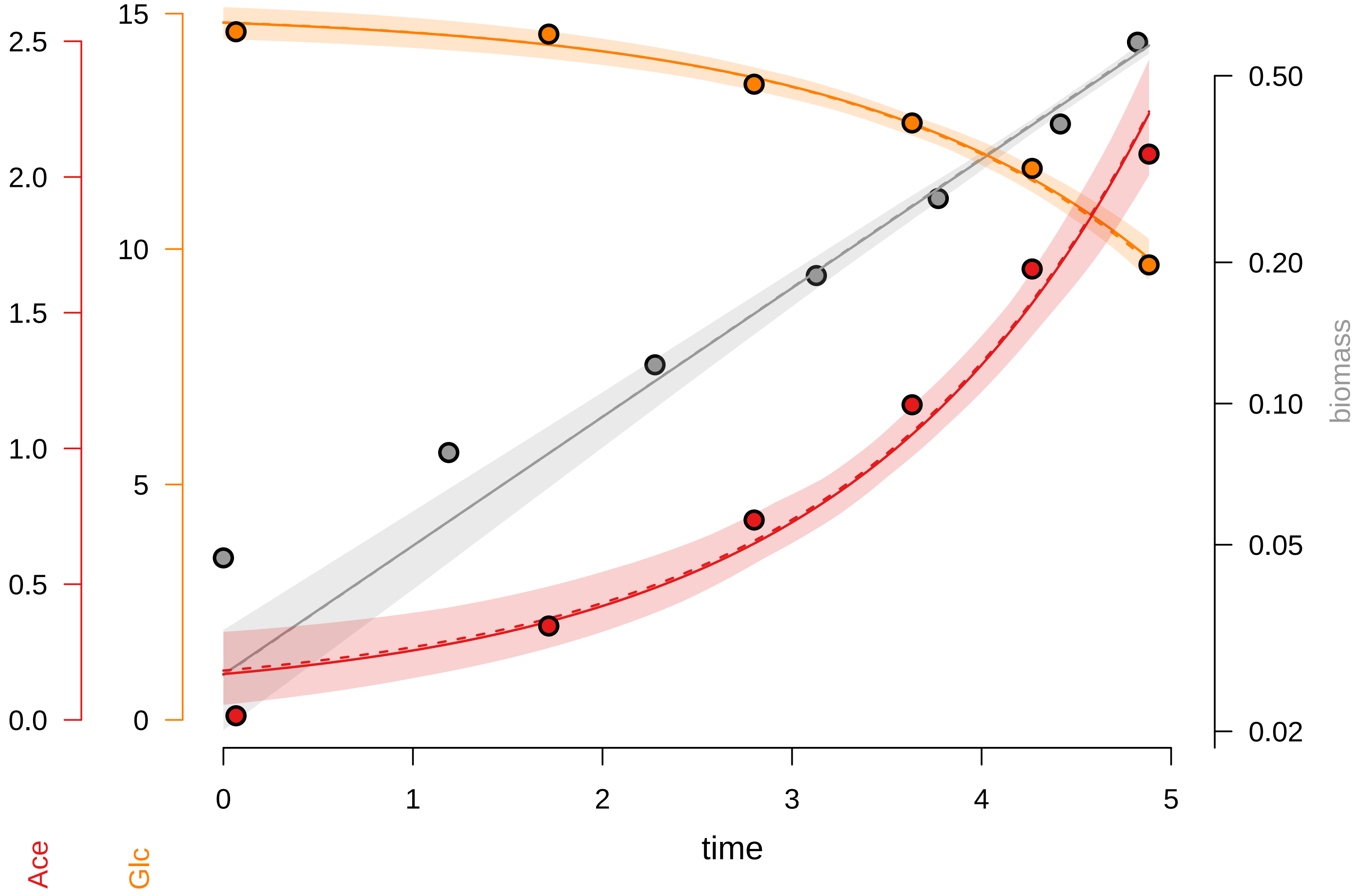

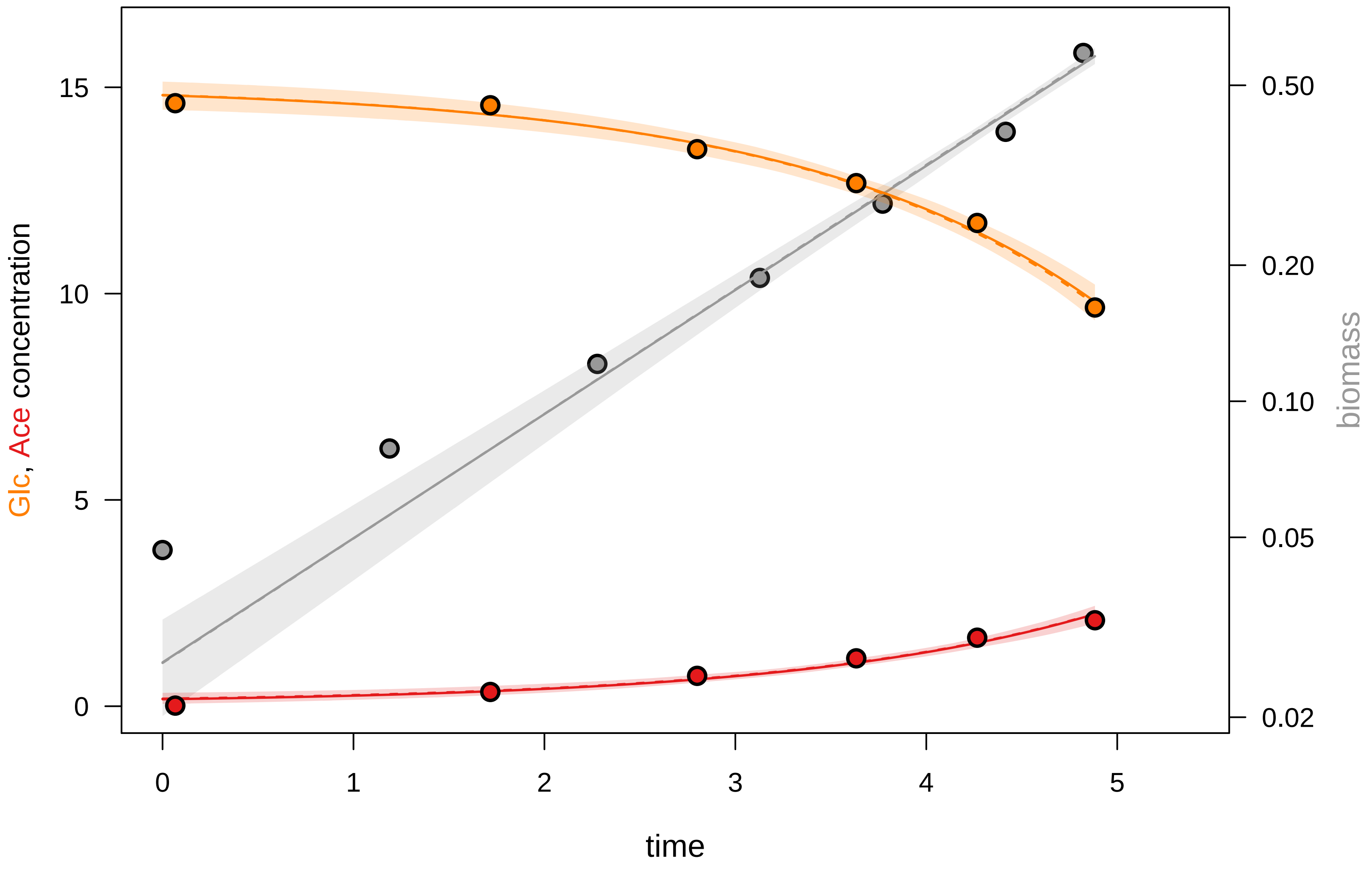

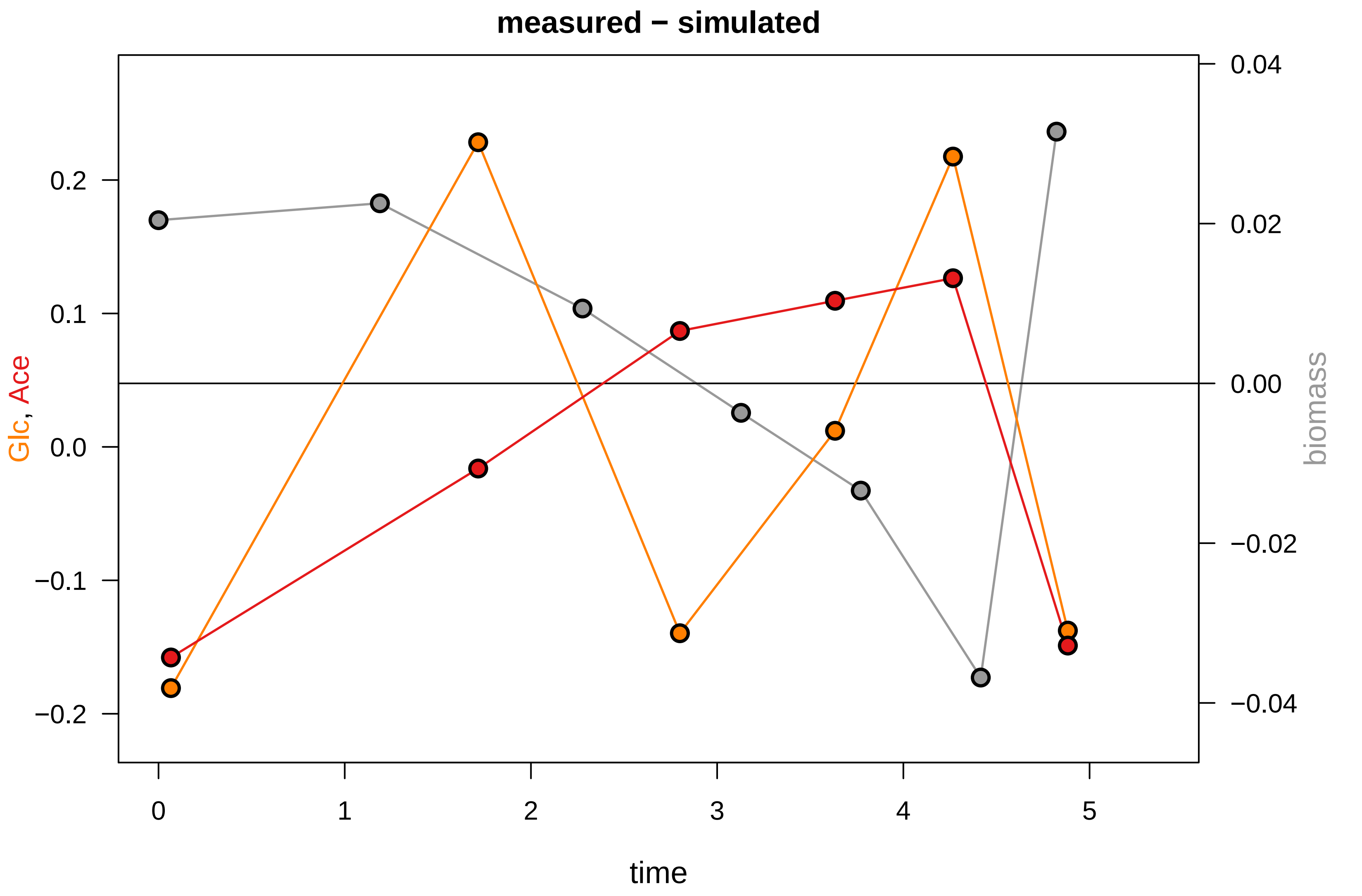

### KEIO_ROBOT1_36.pdf

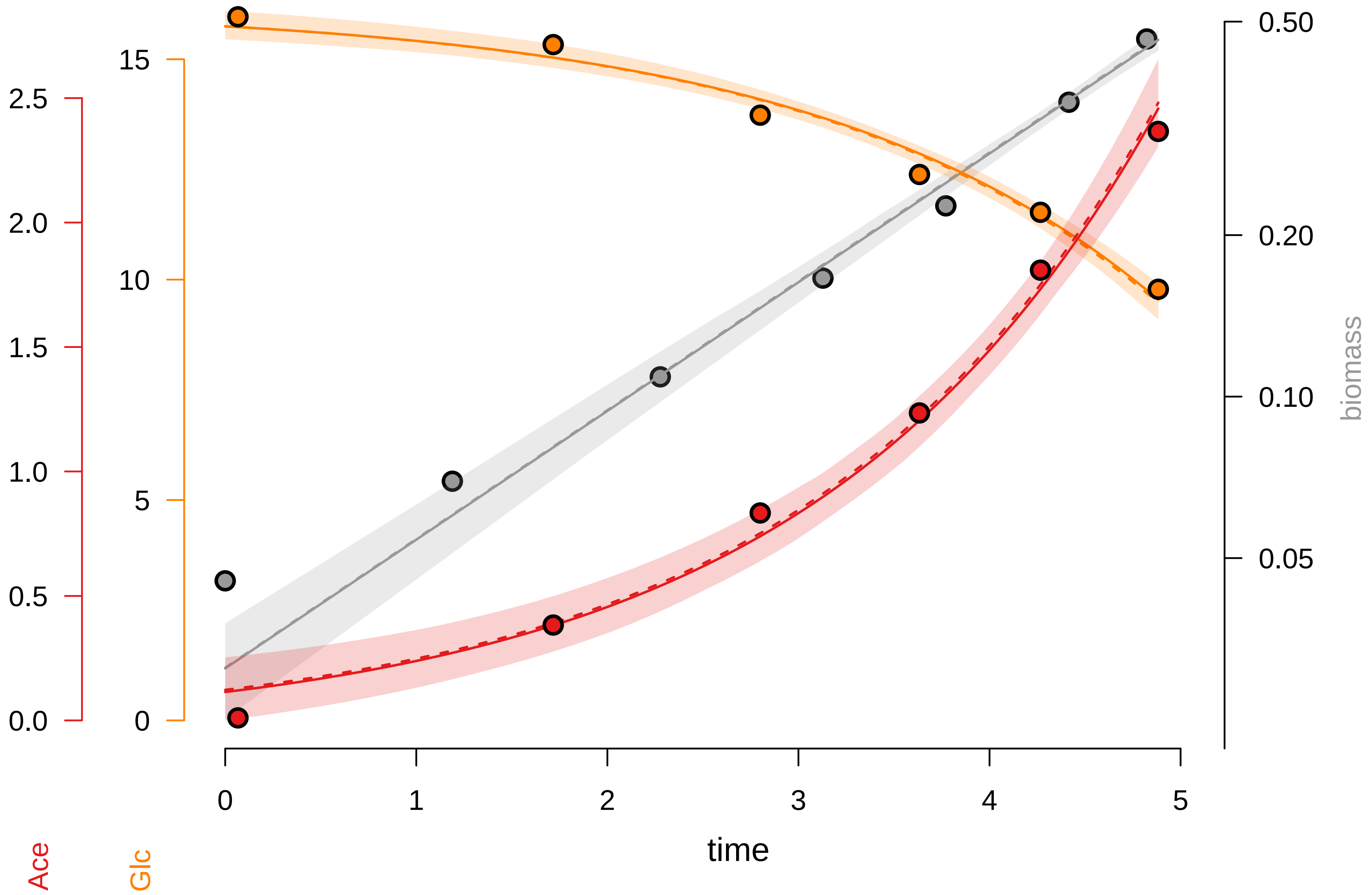

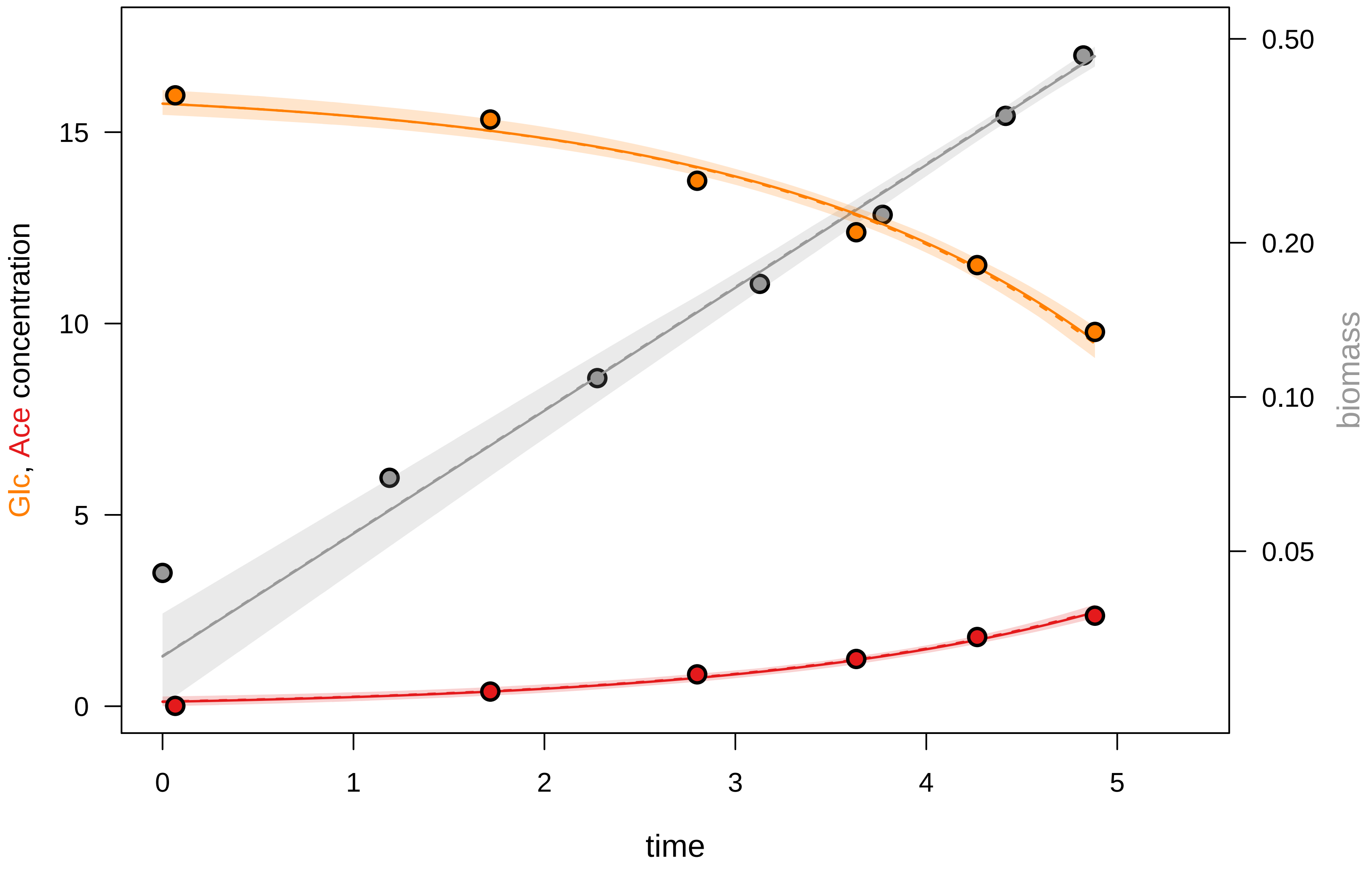

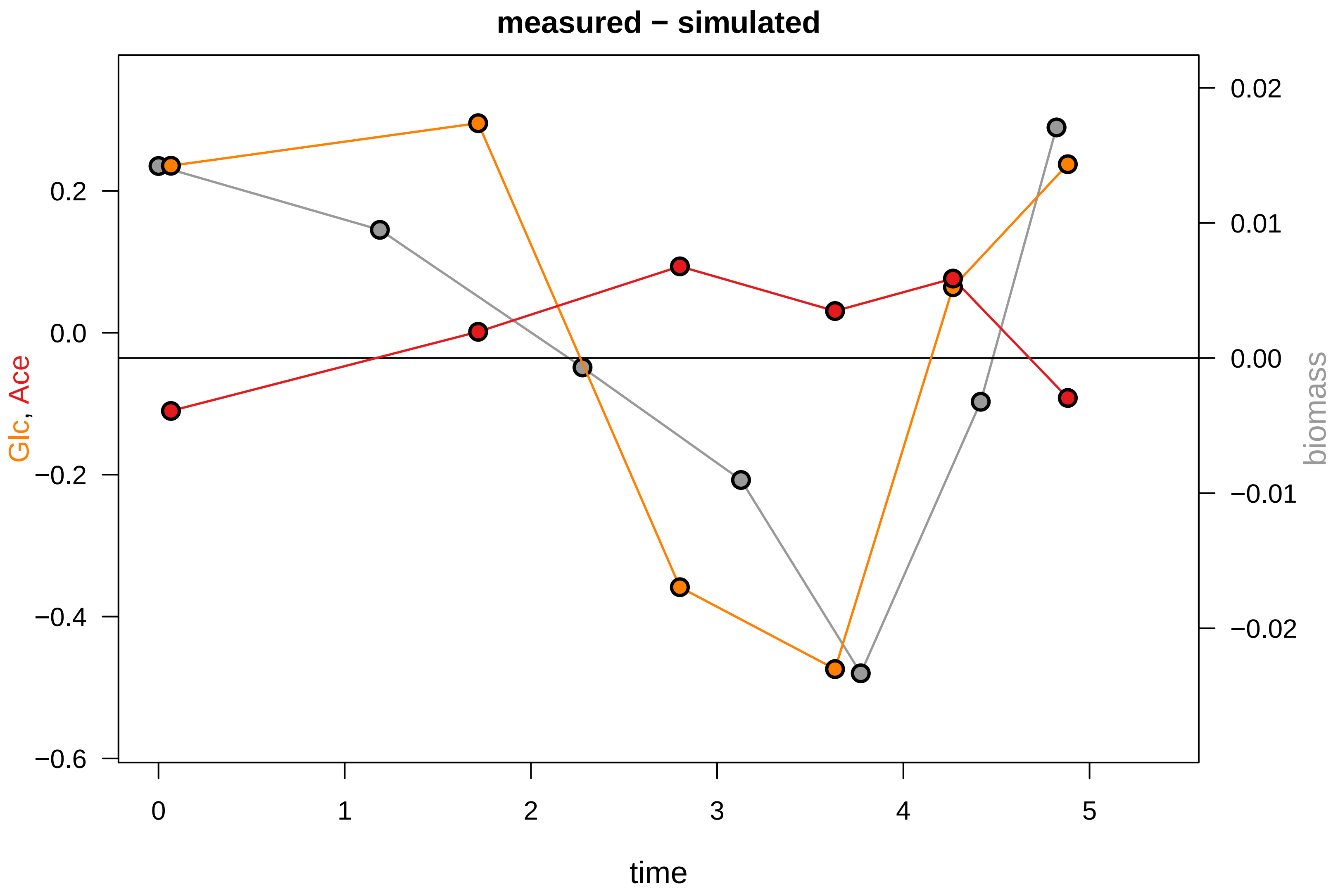

### KEIO_ROBOT1_37.pdf

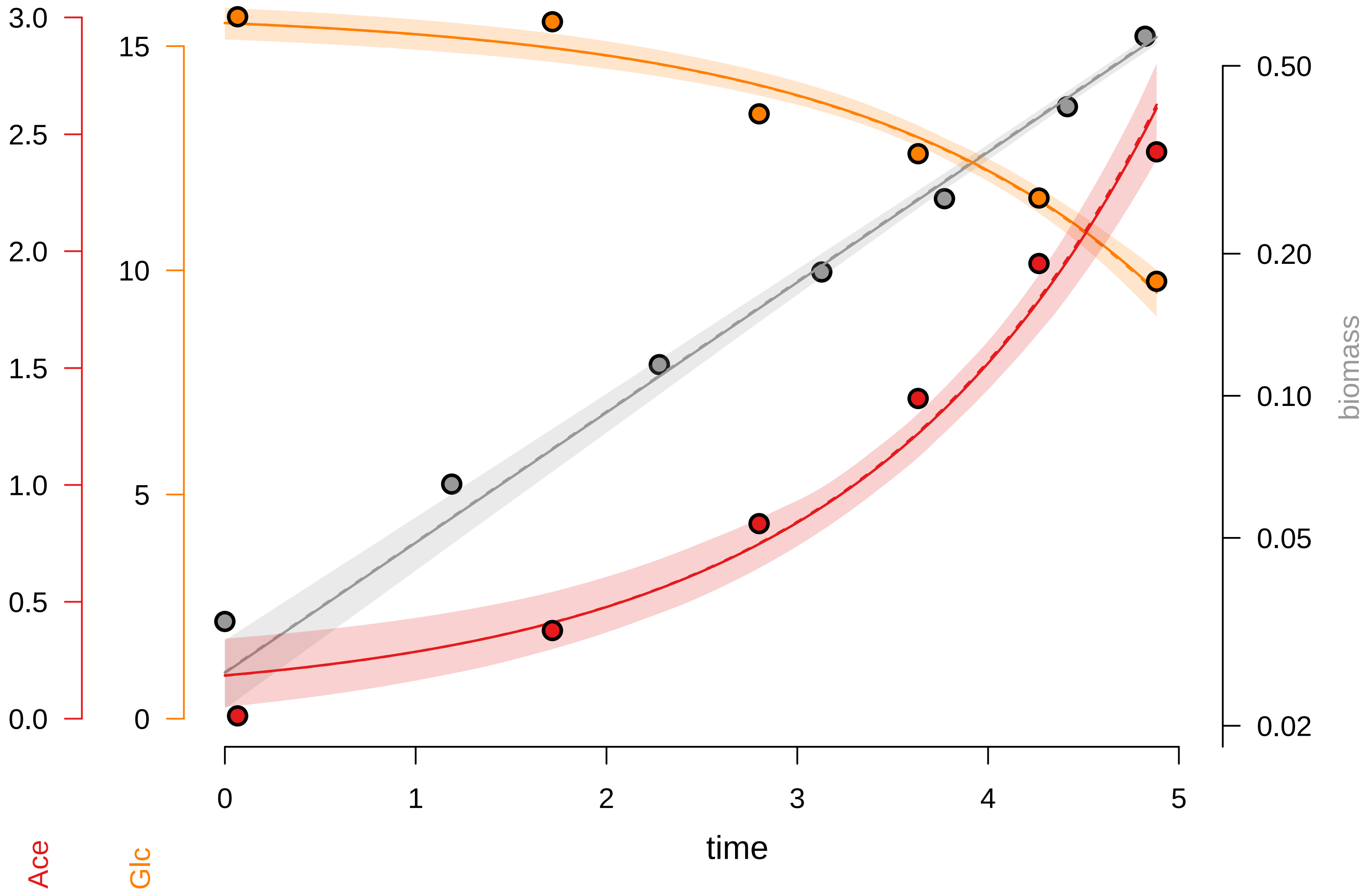

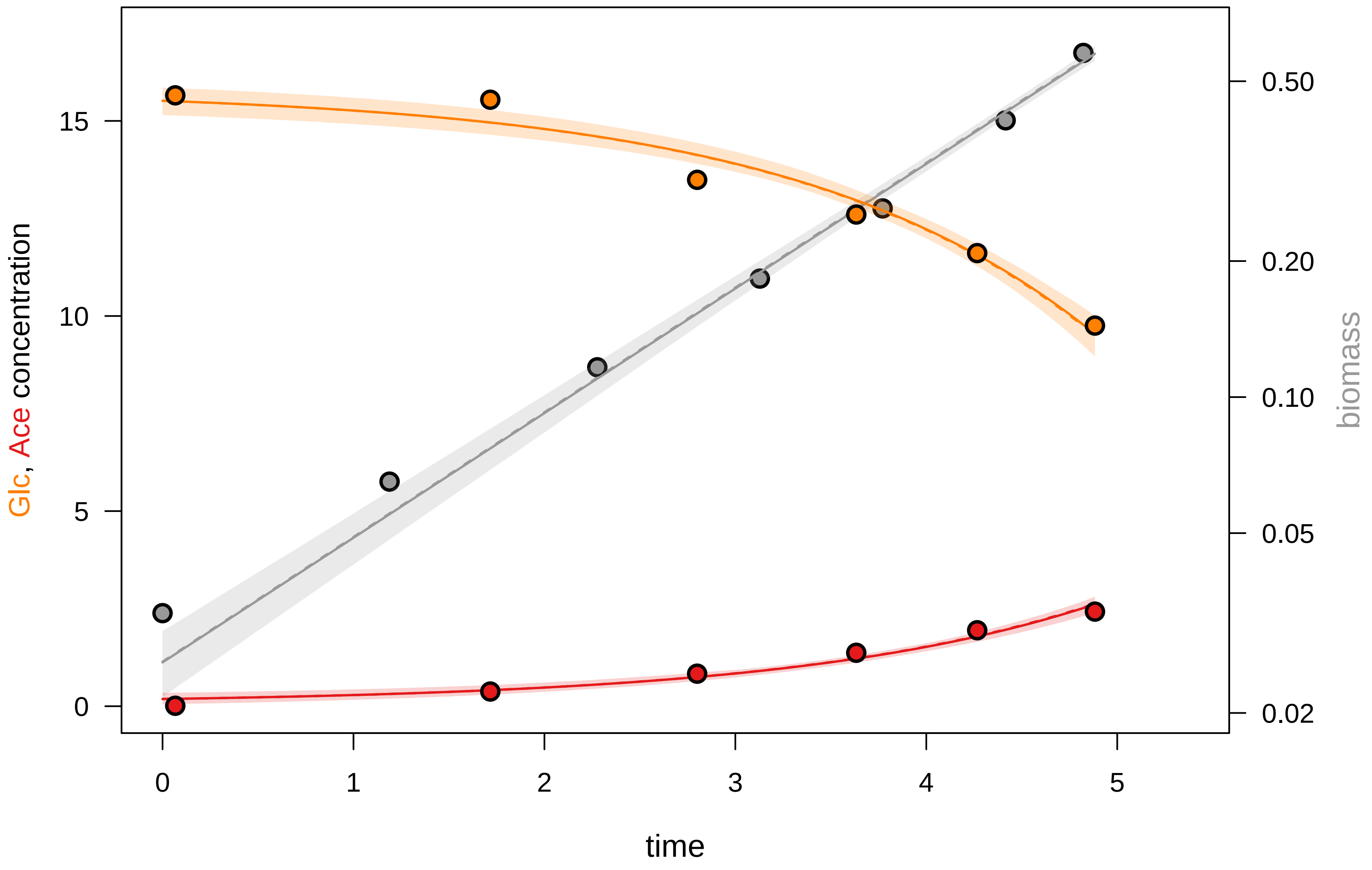

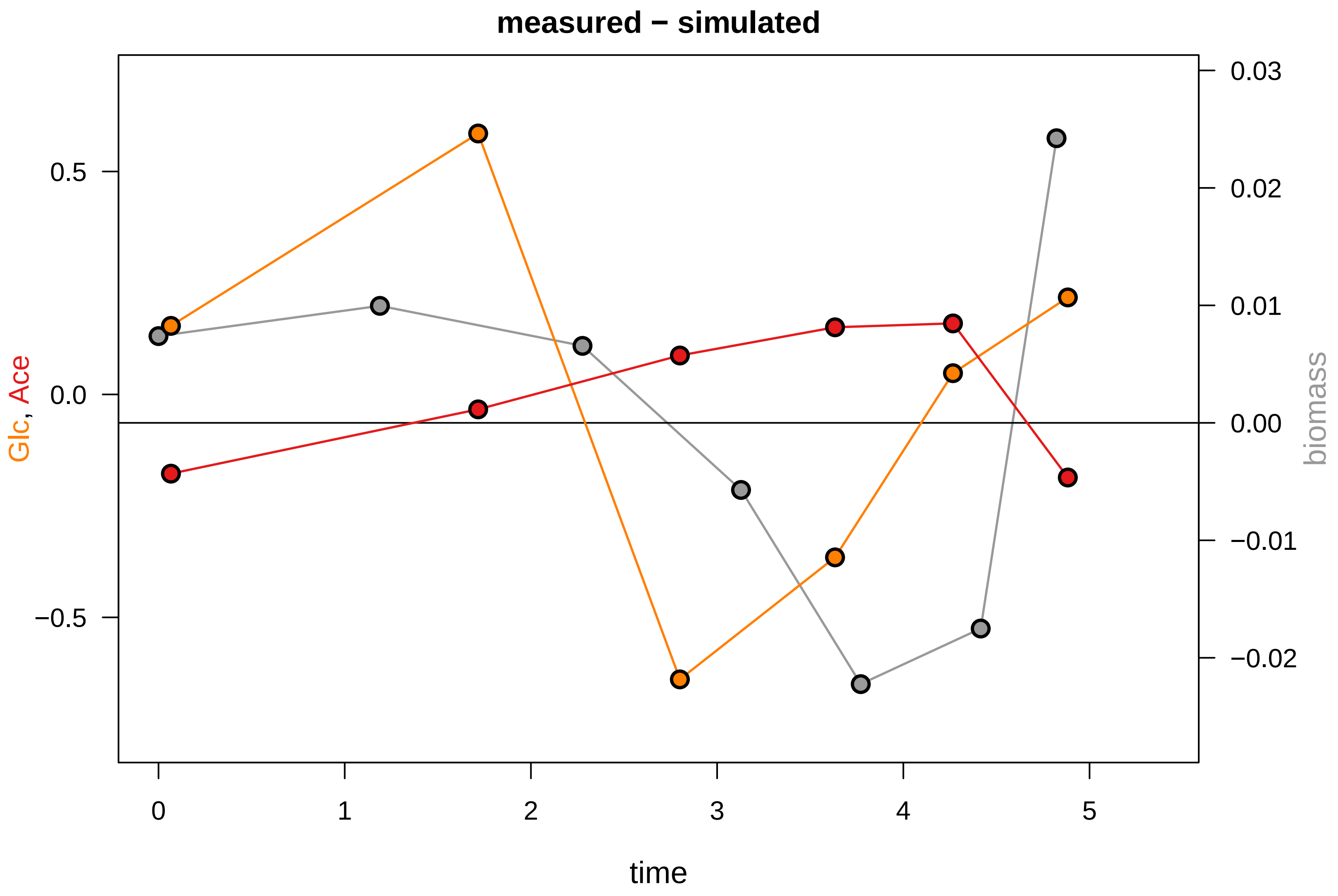

### KEIO_ROBOT1_38.pdf

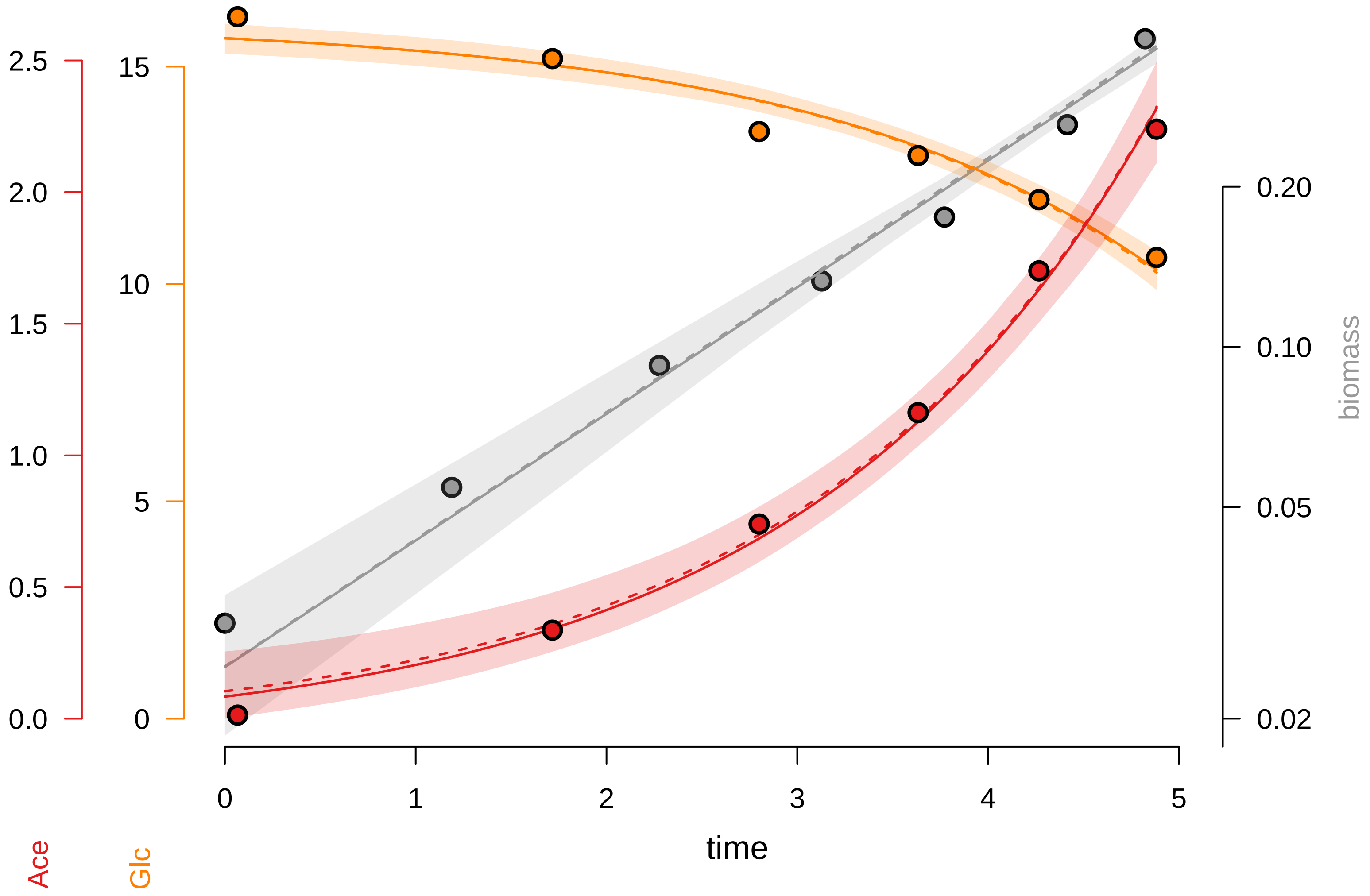

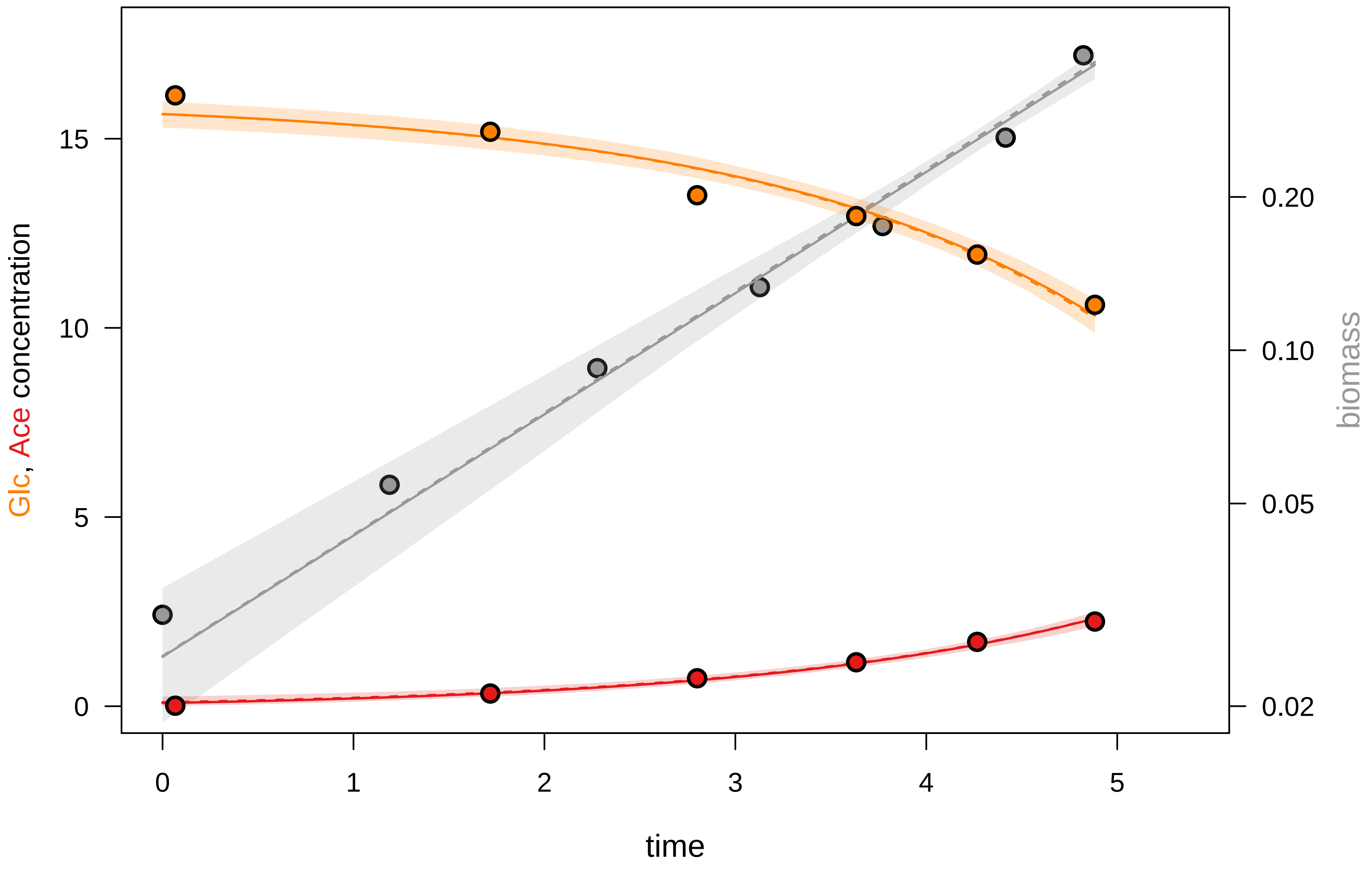

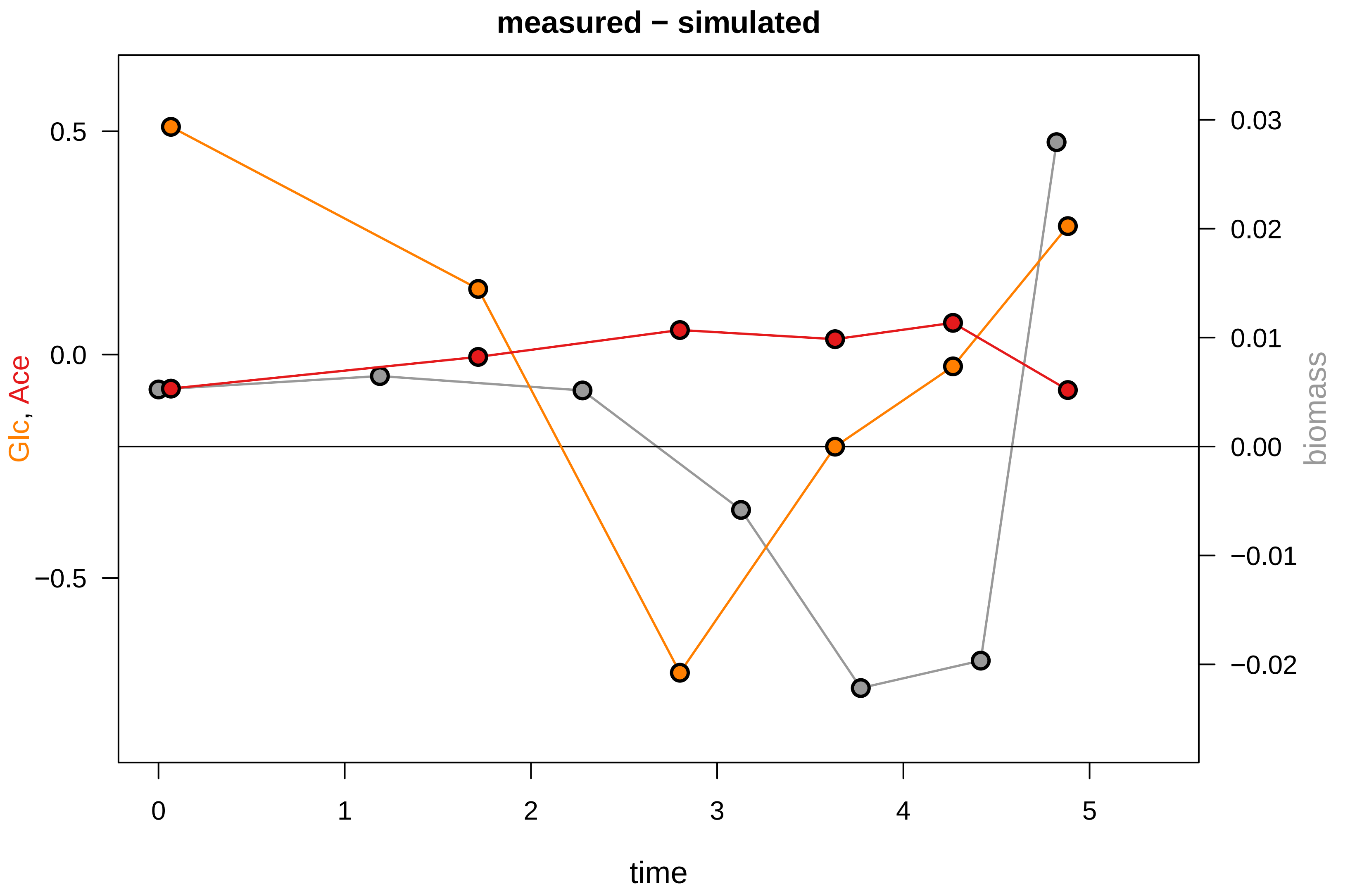

### KEIO_ROBOT1_40.pdf

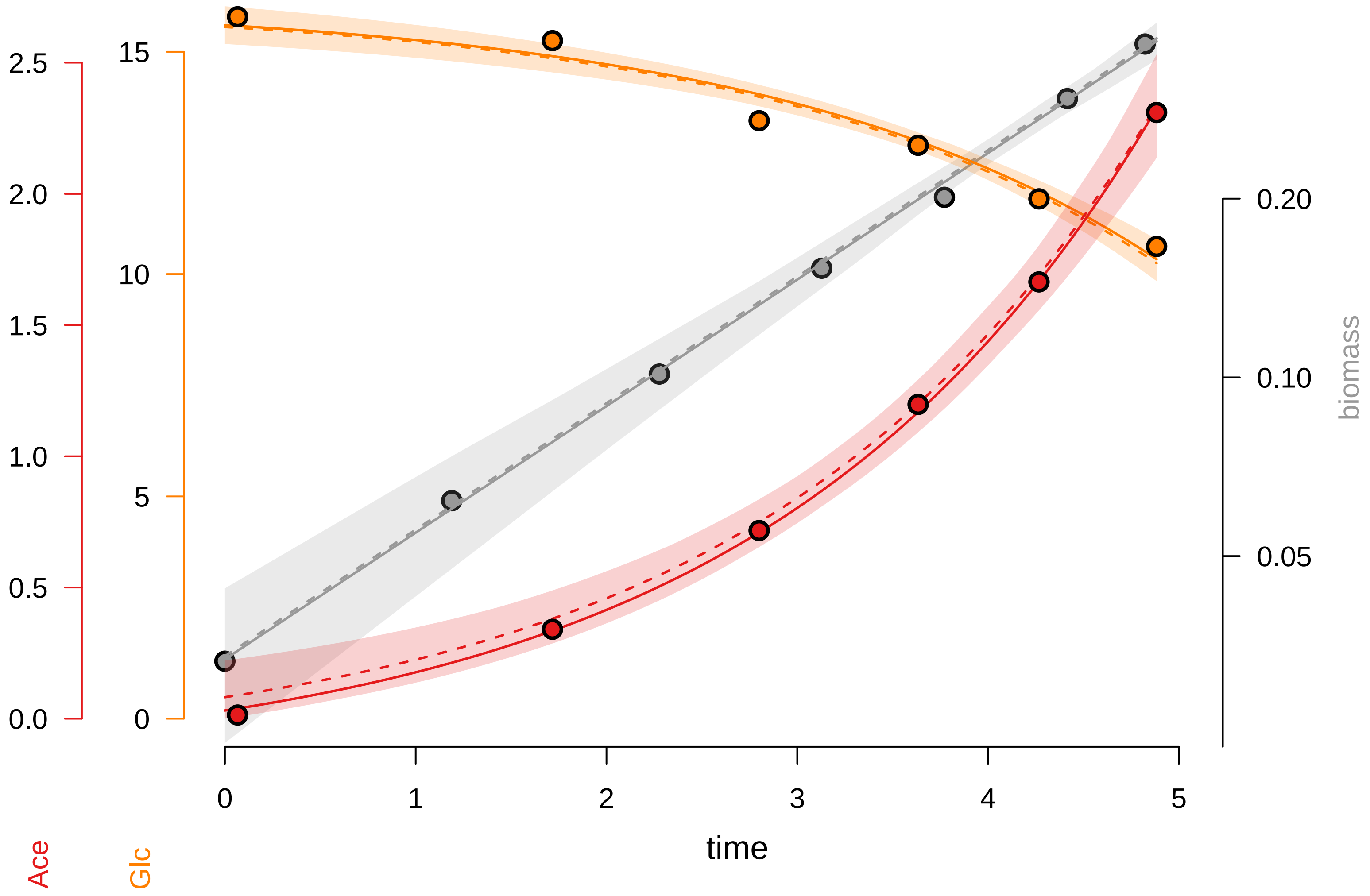

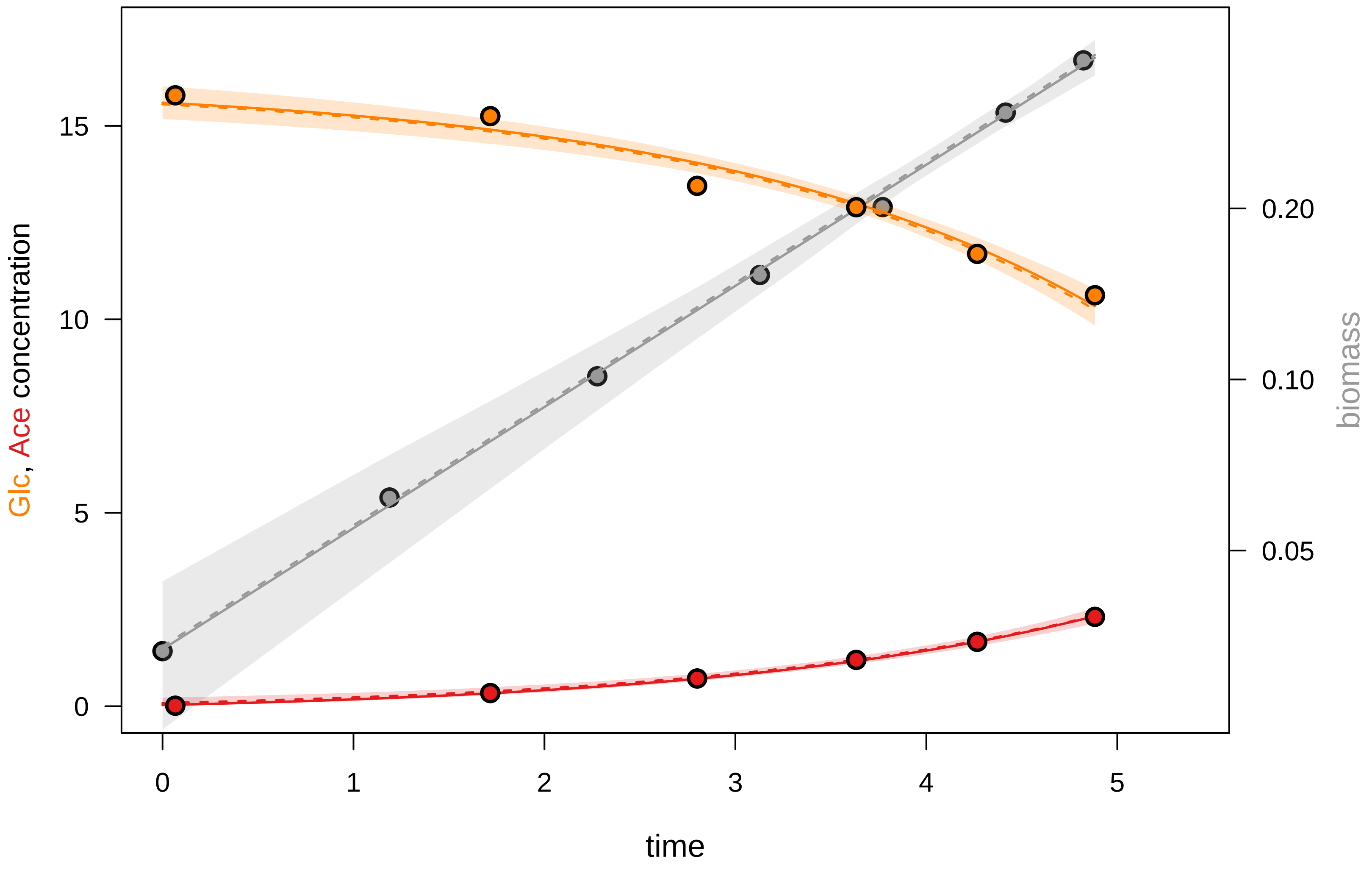

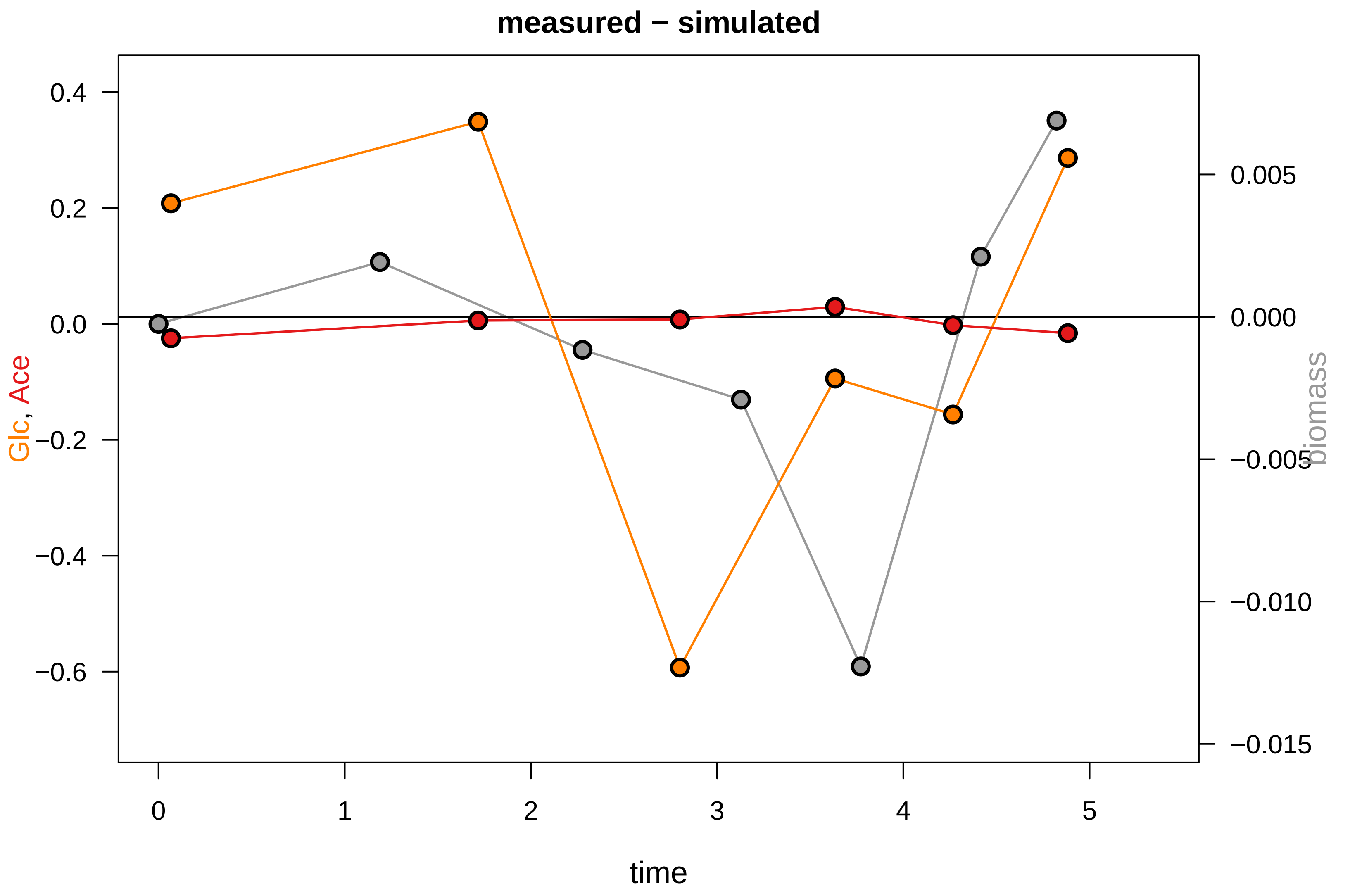

### KEIO_ROBOT1_42.pdf

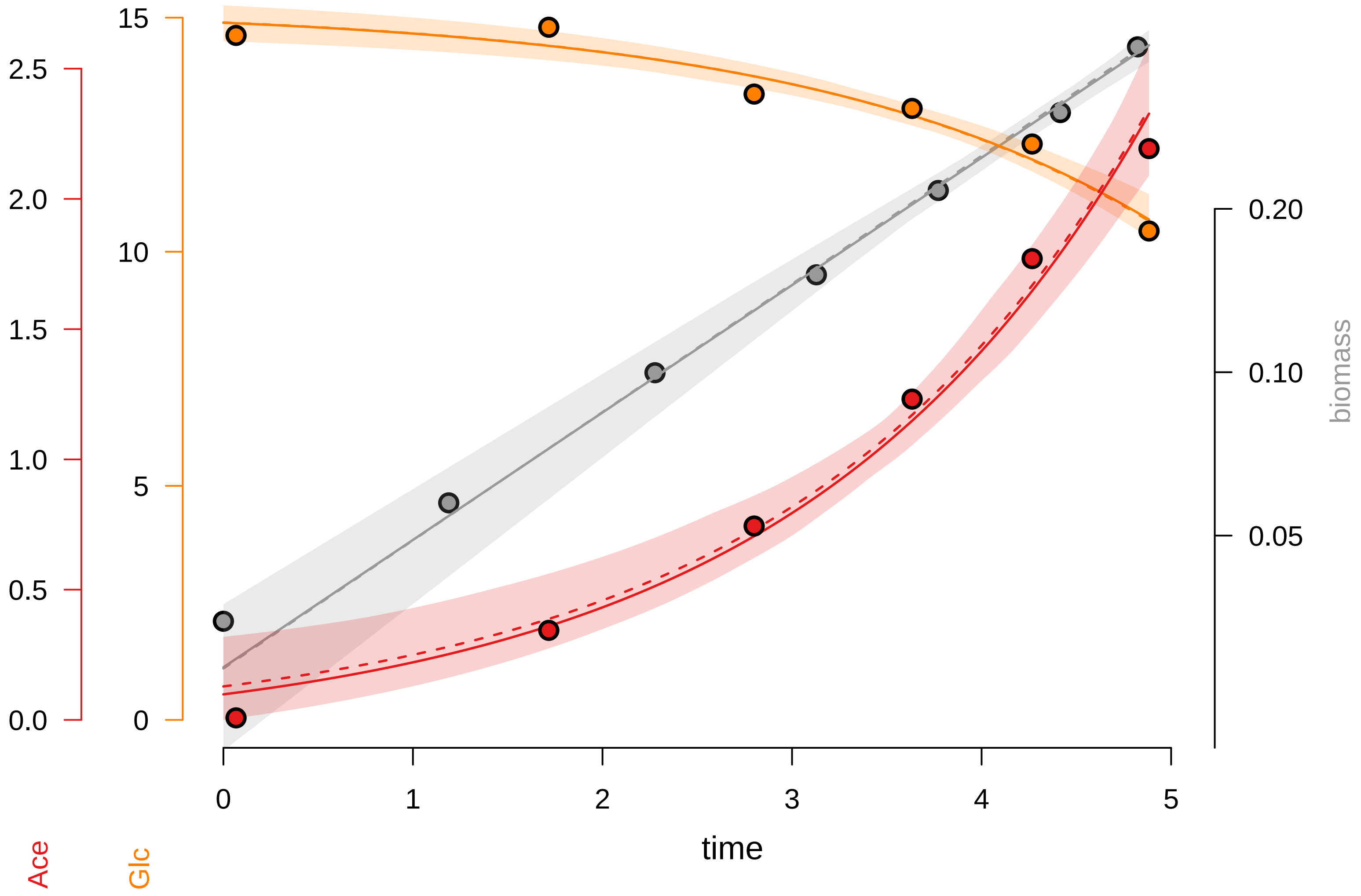

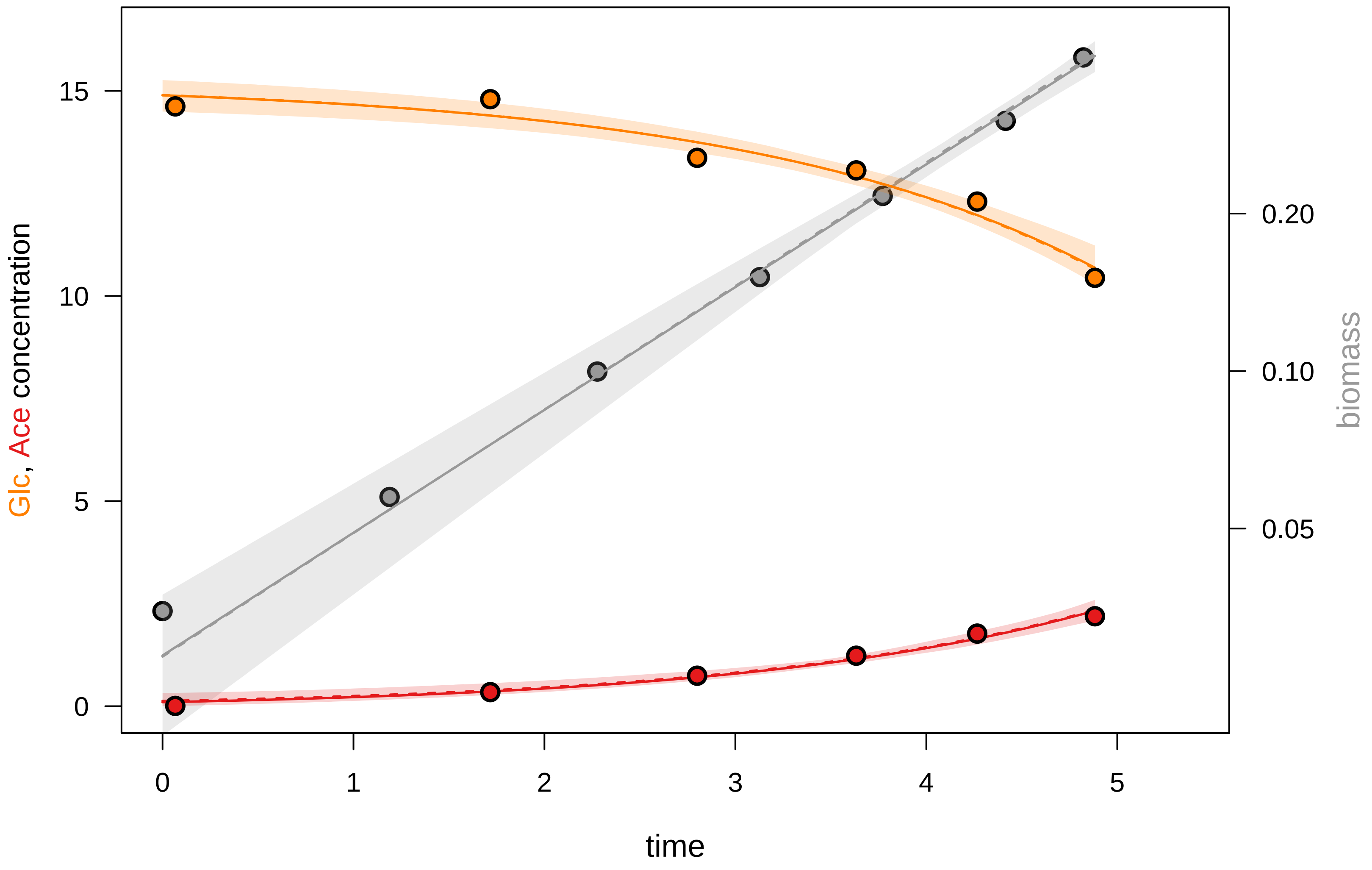

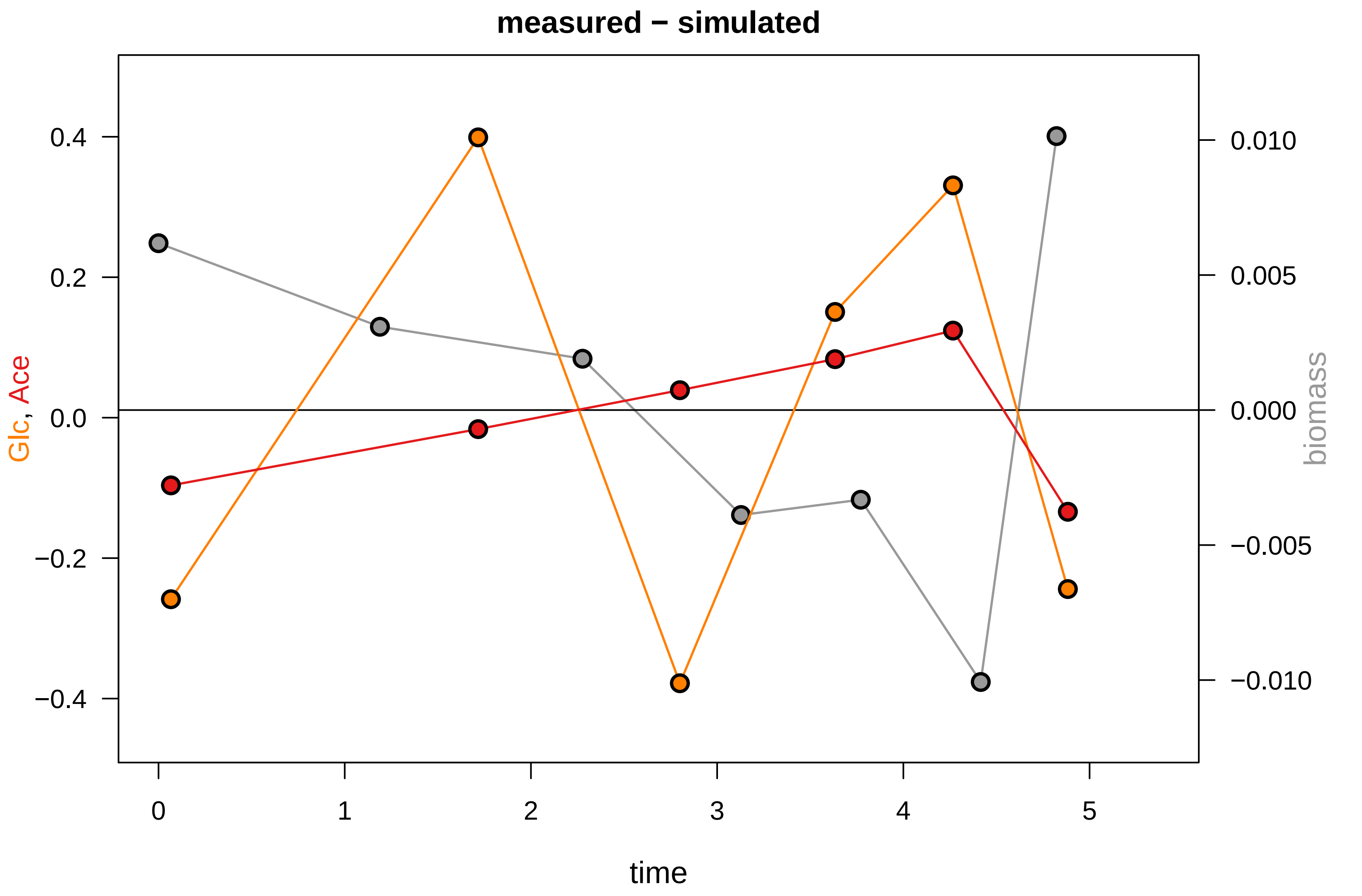

### KEIO_ROBOT1_43.pdf

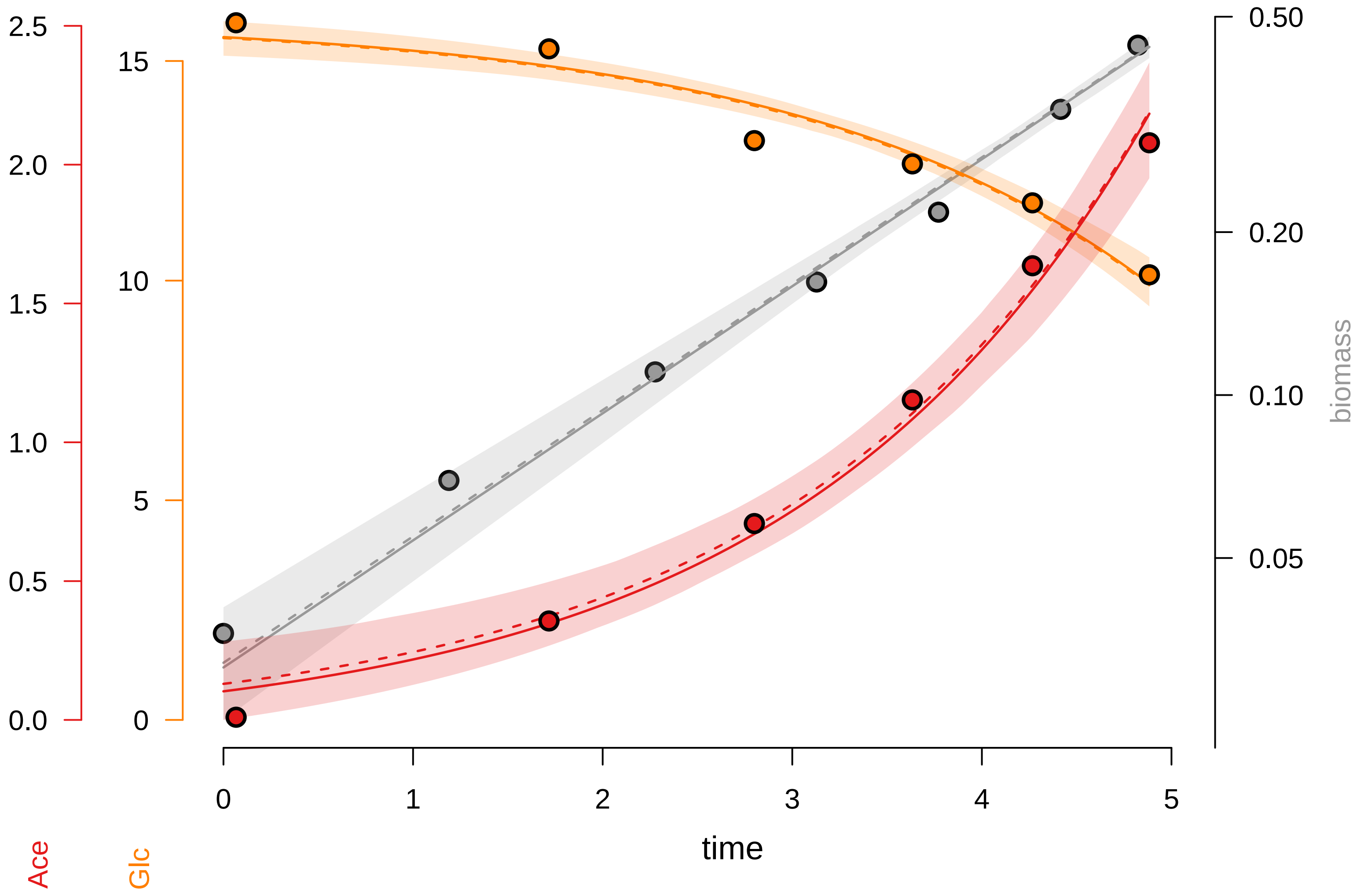

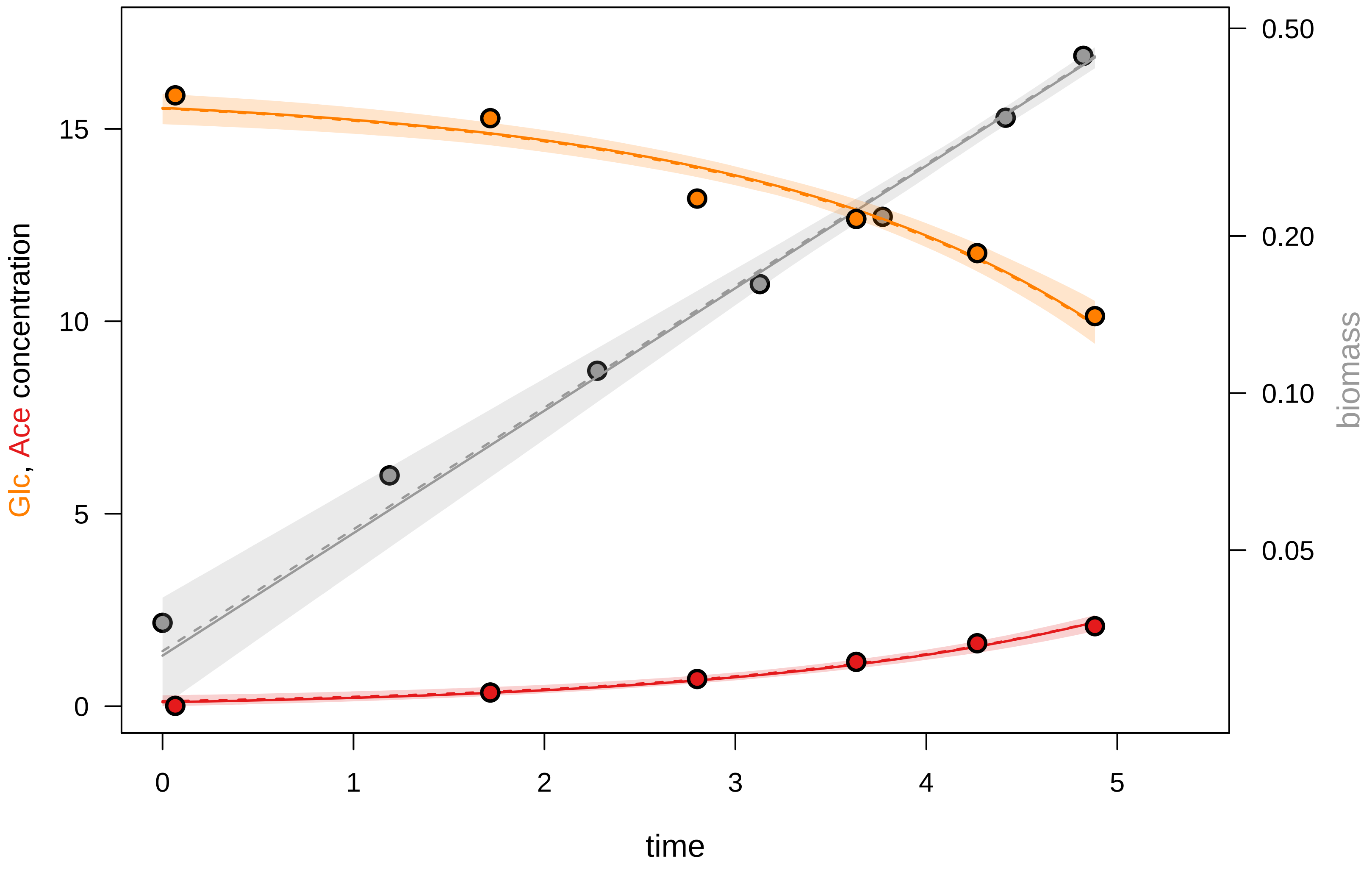

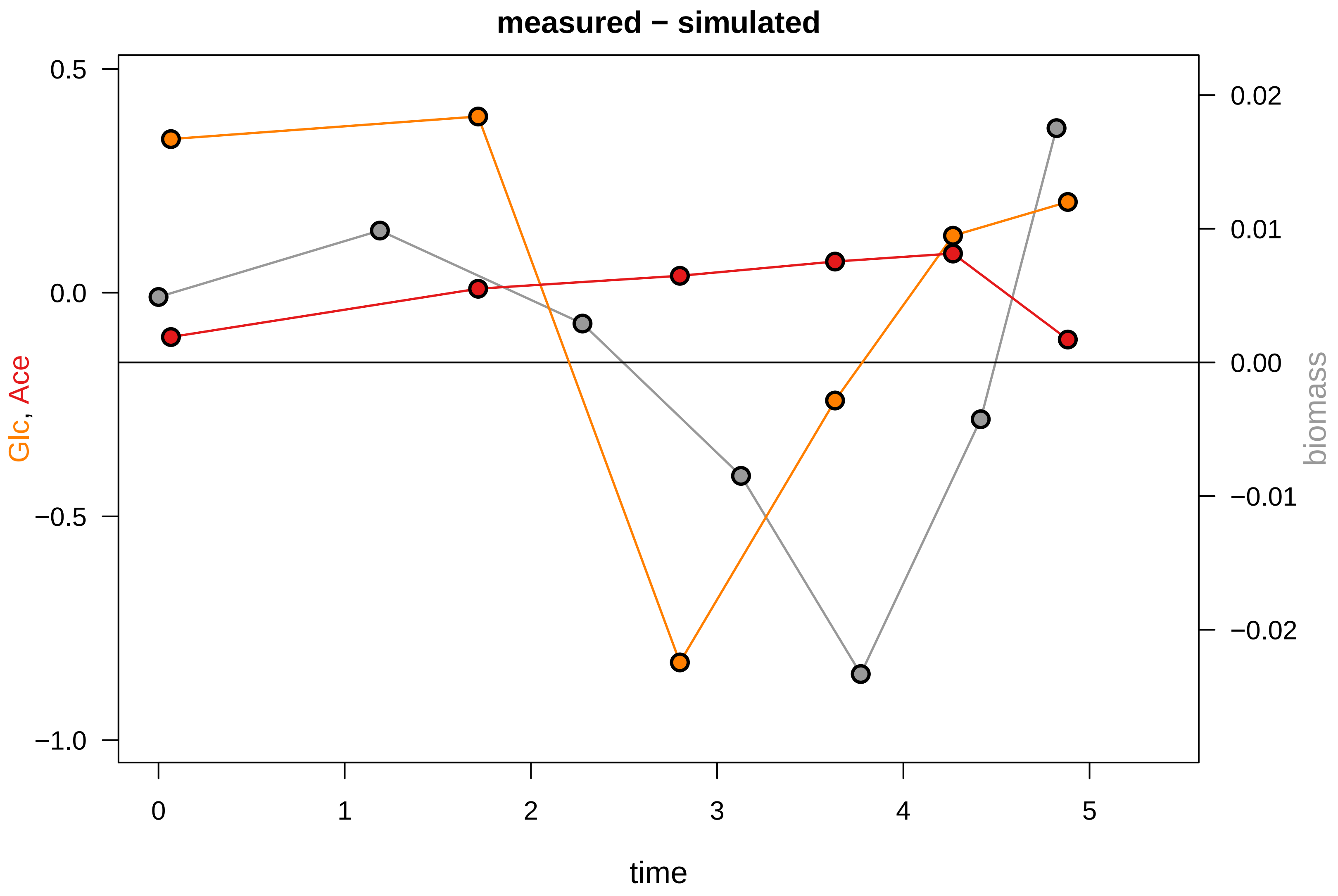

### KEIO_ROBOT1_46.pdf

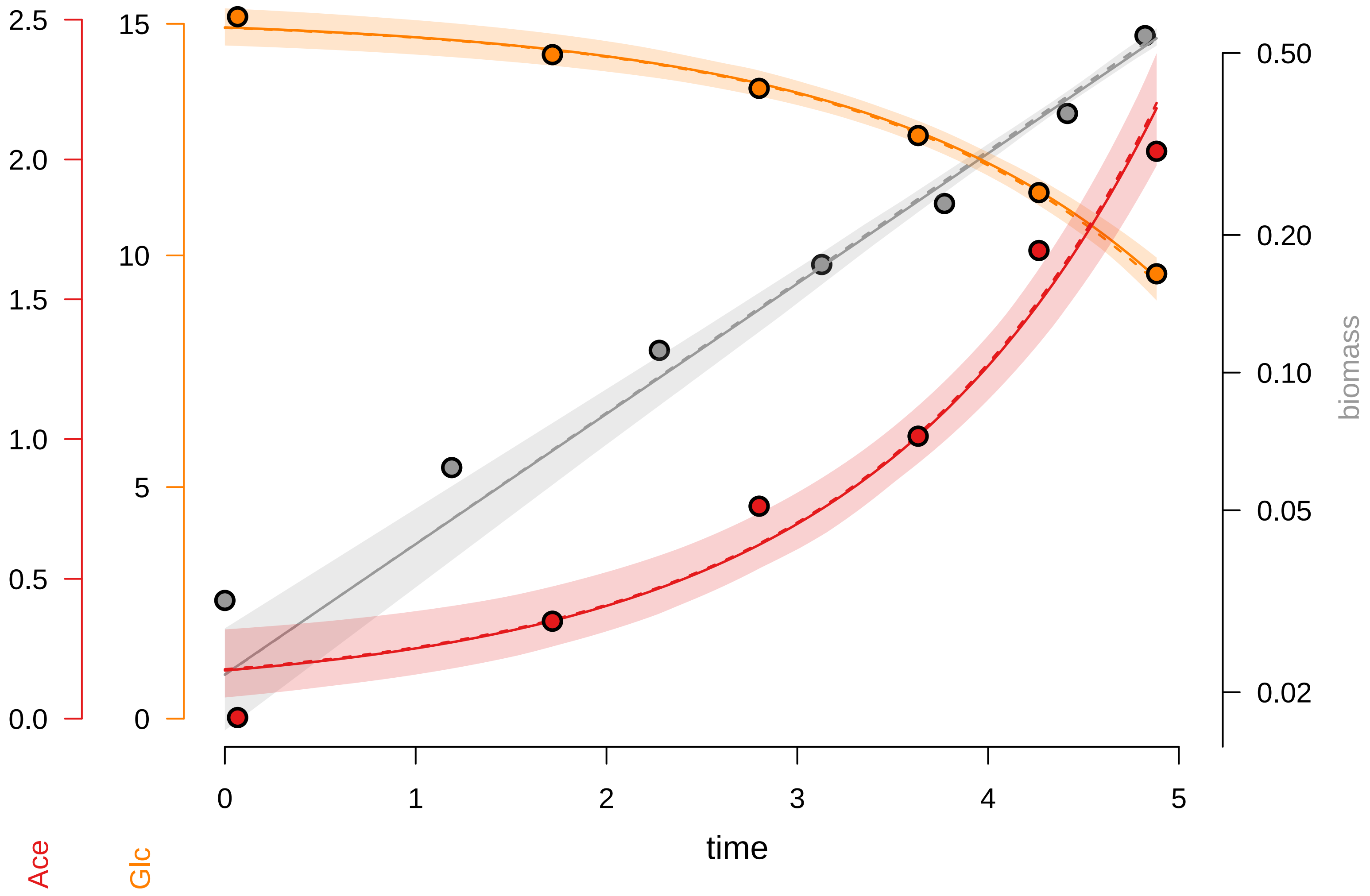

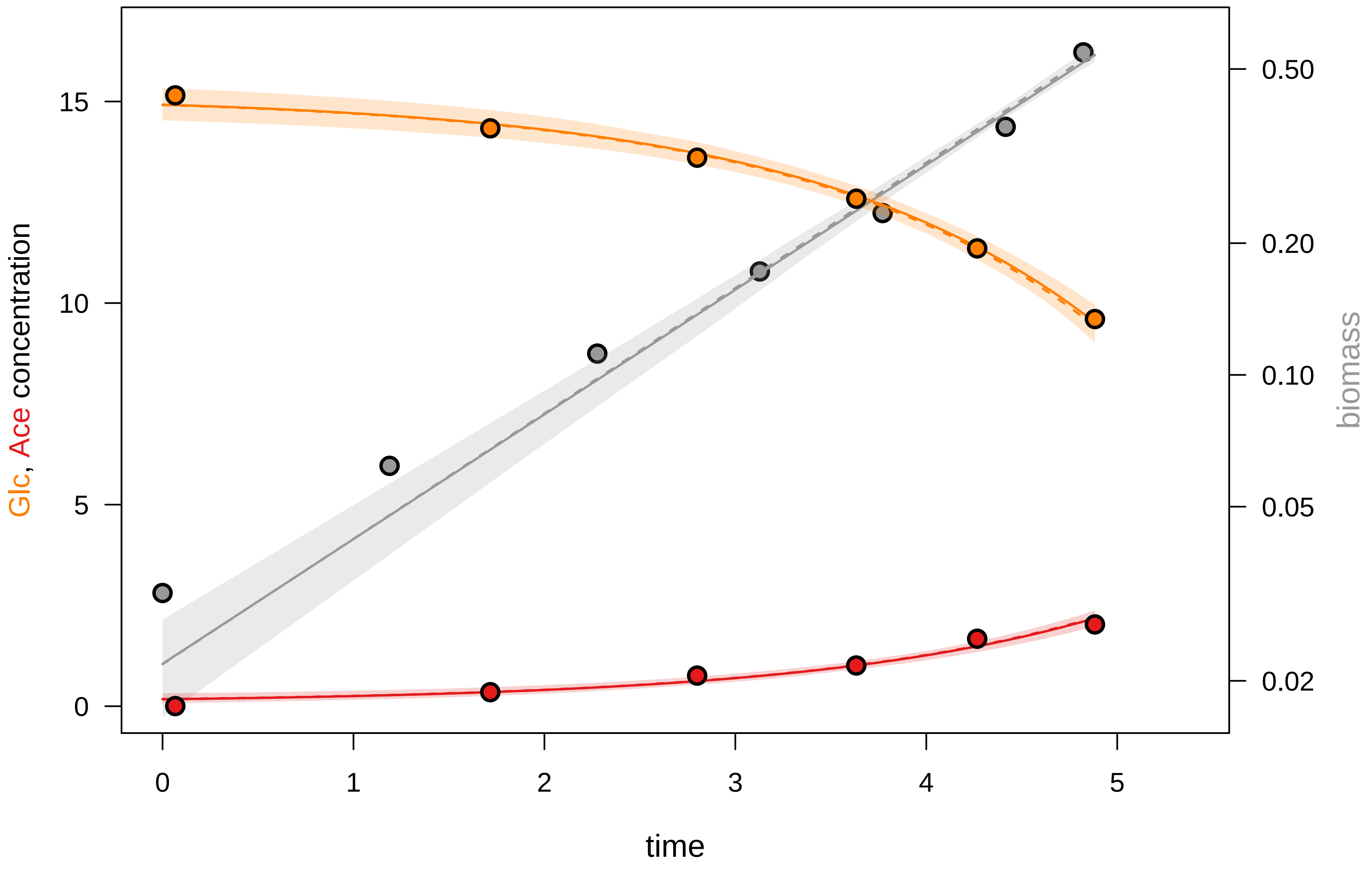

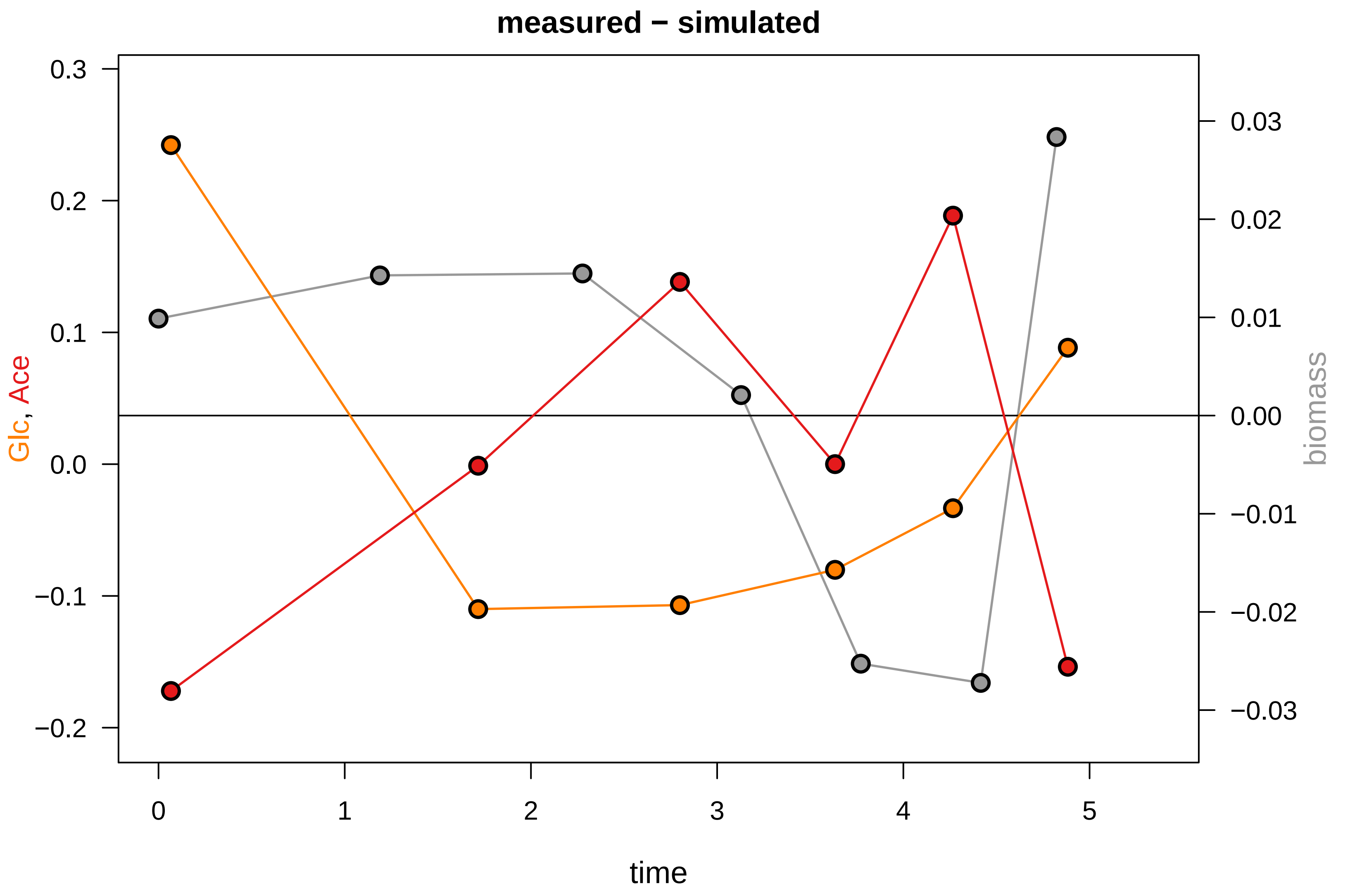

### KEIO_ROBOT2_5.pdf

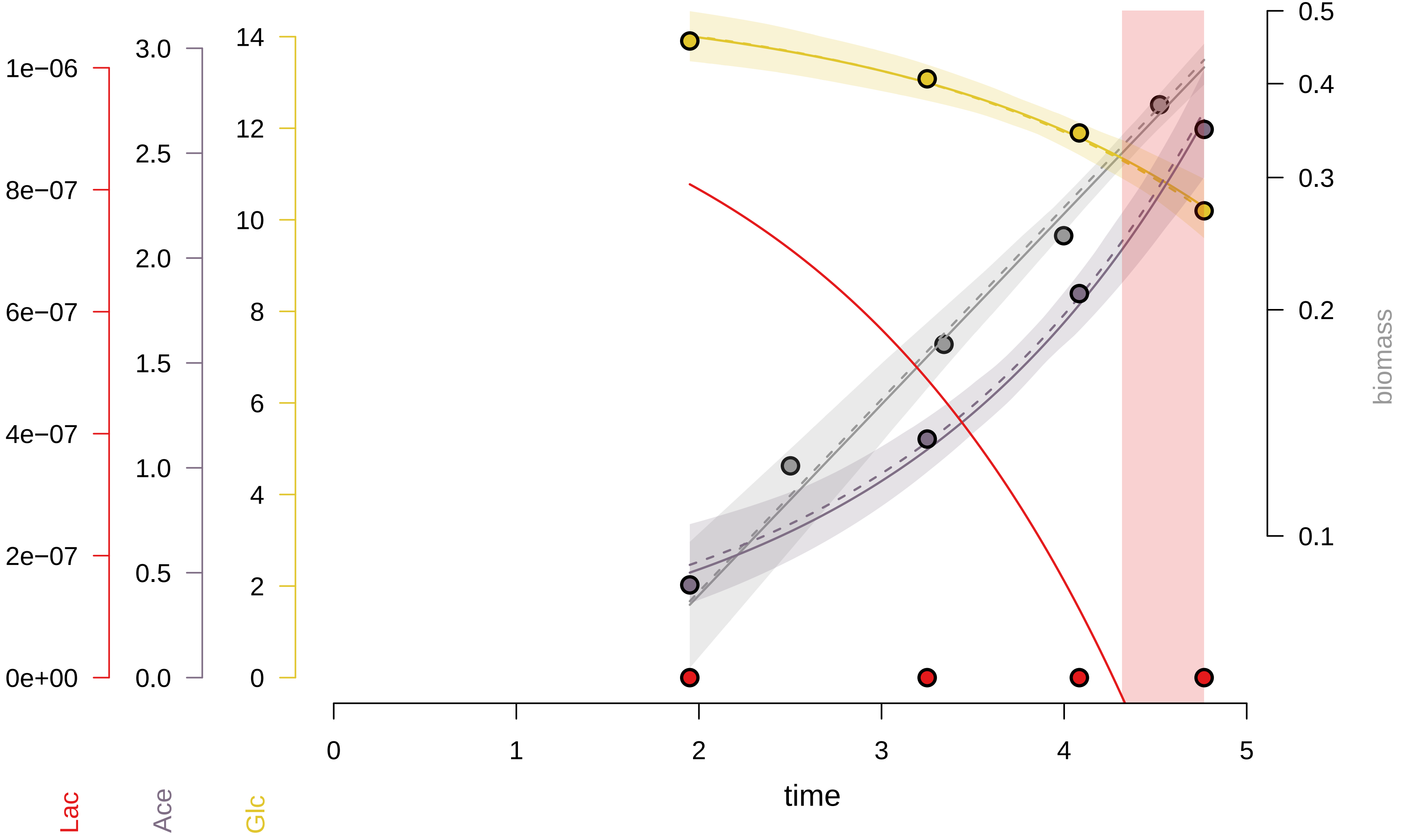

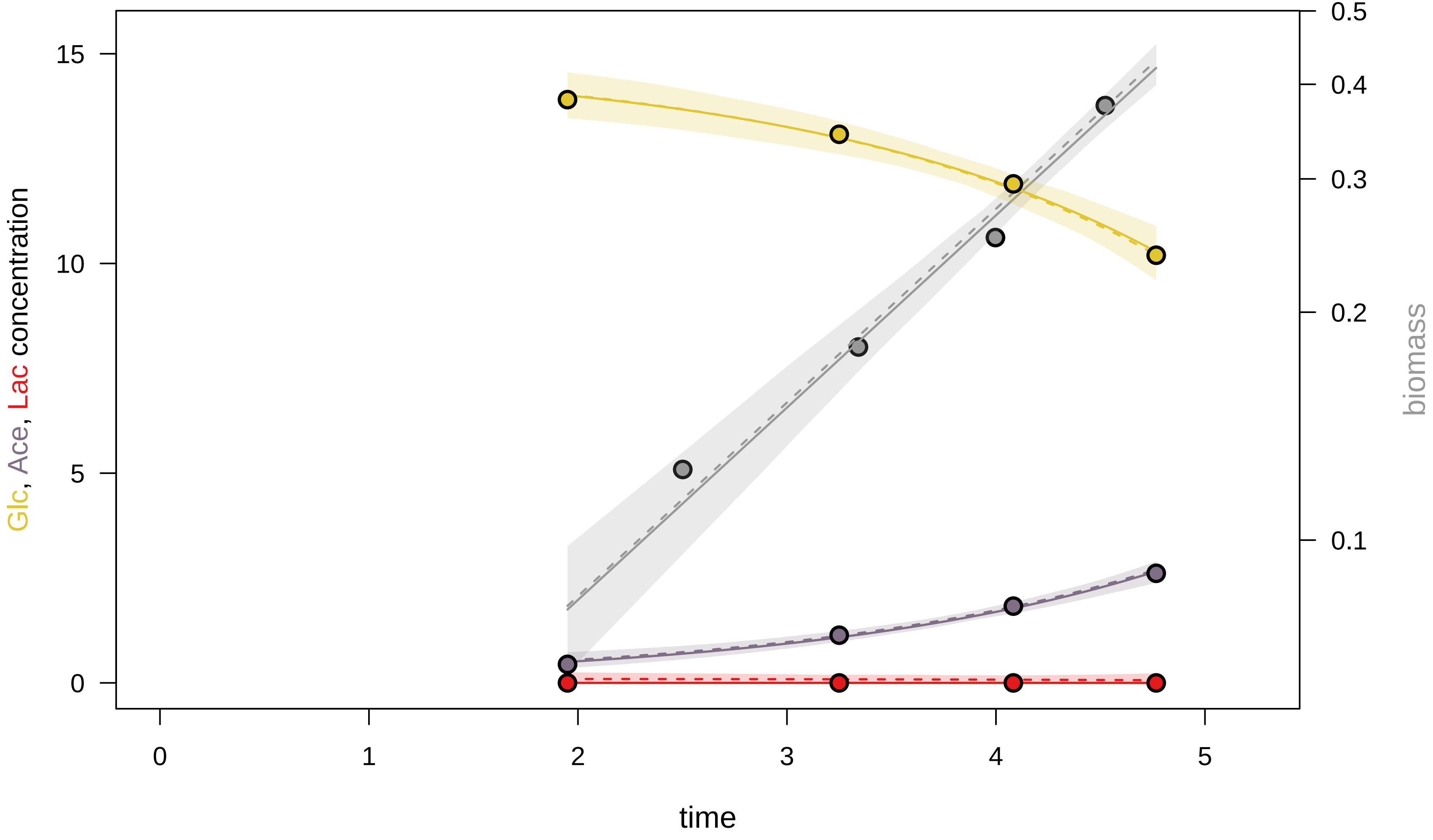

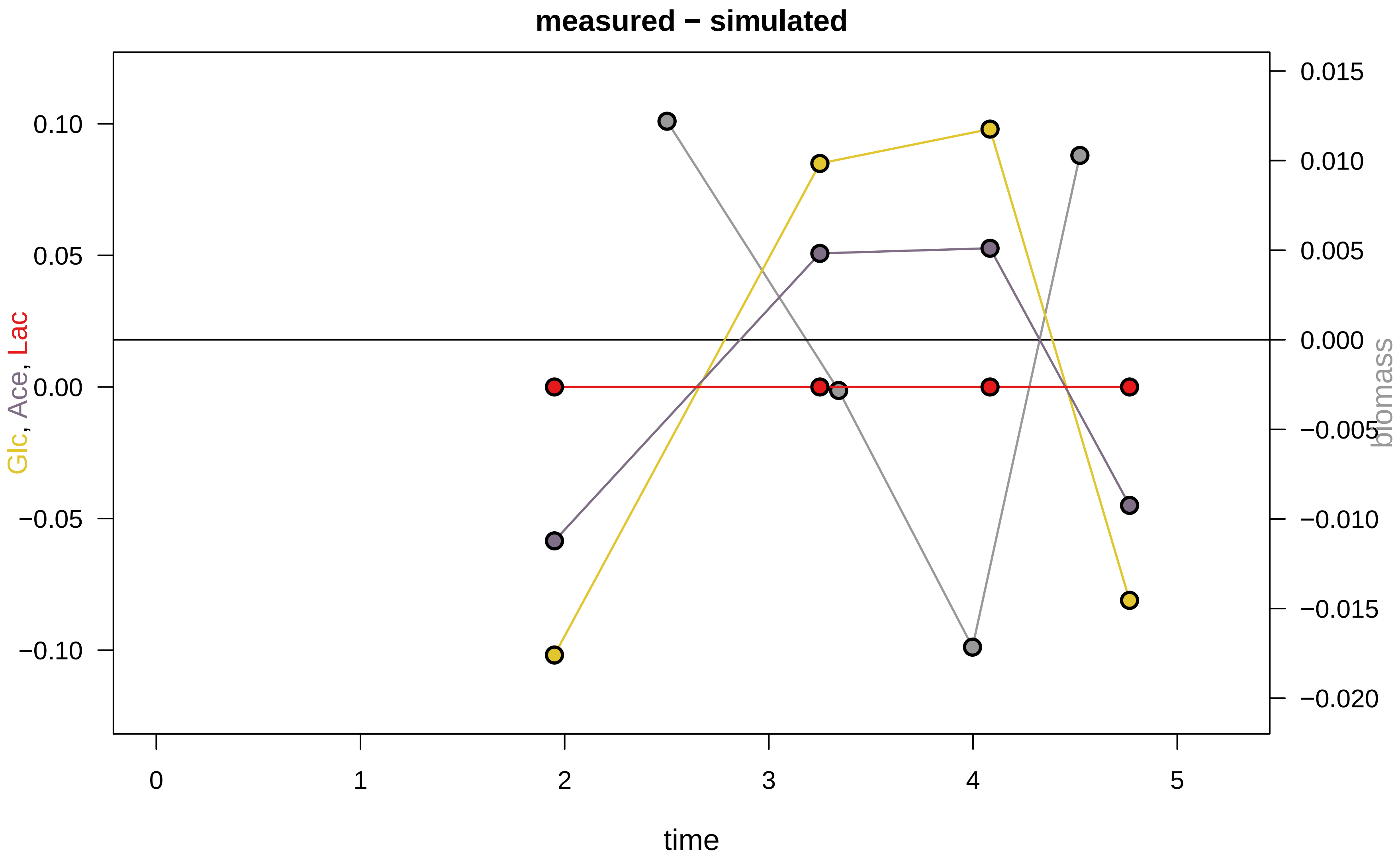

### KEIO_ROBOT2_6.pdf

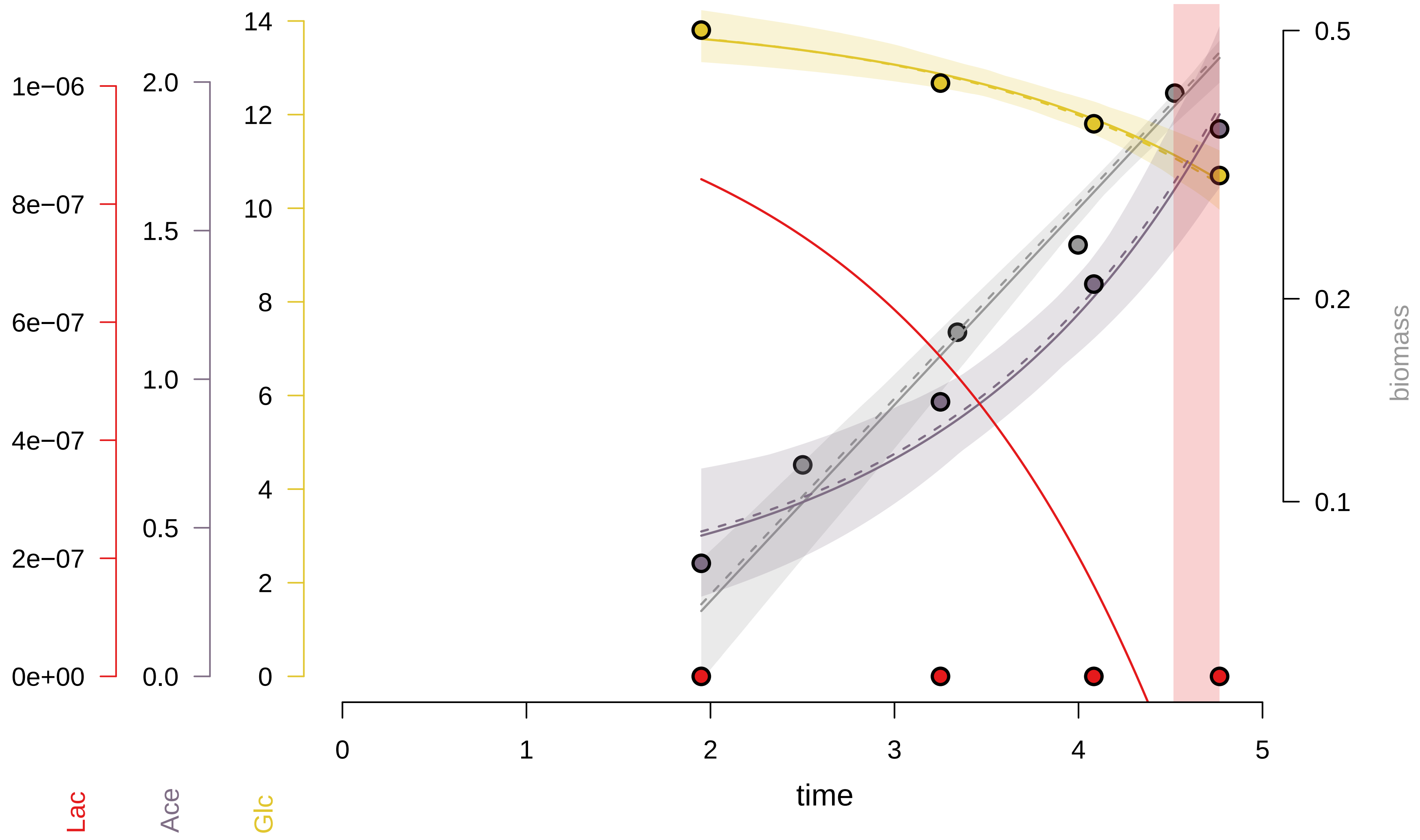

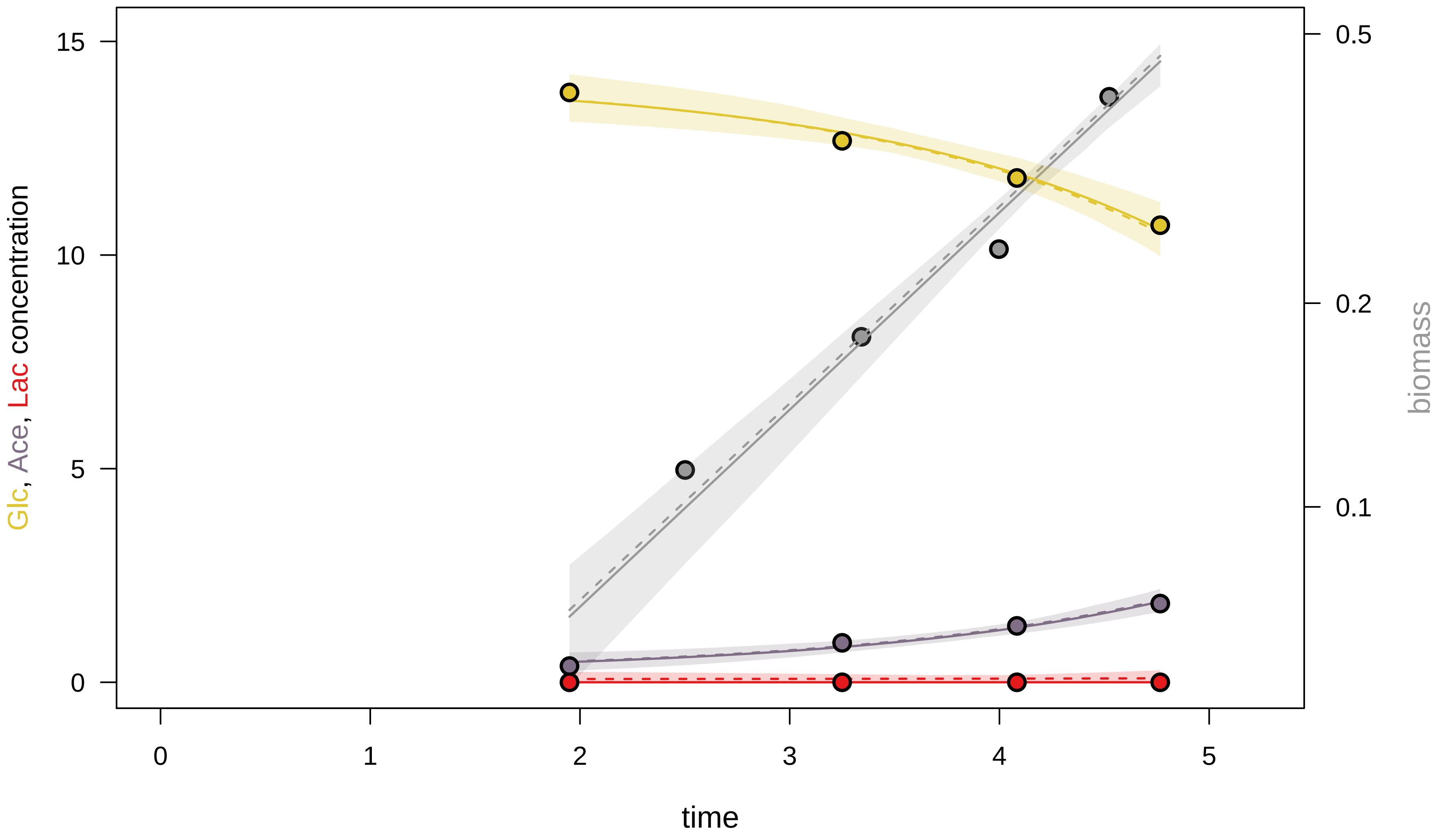

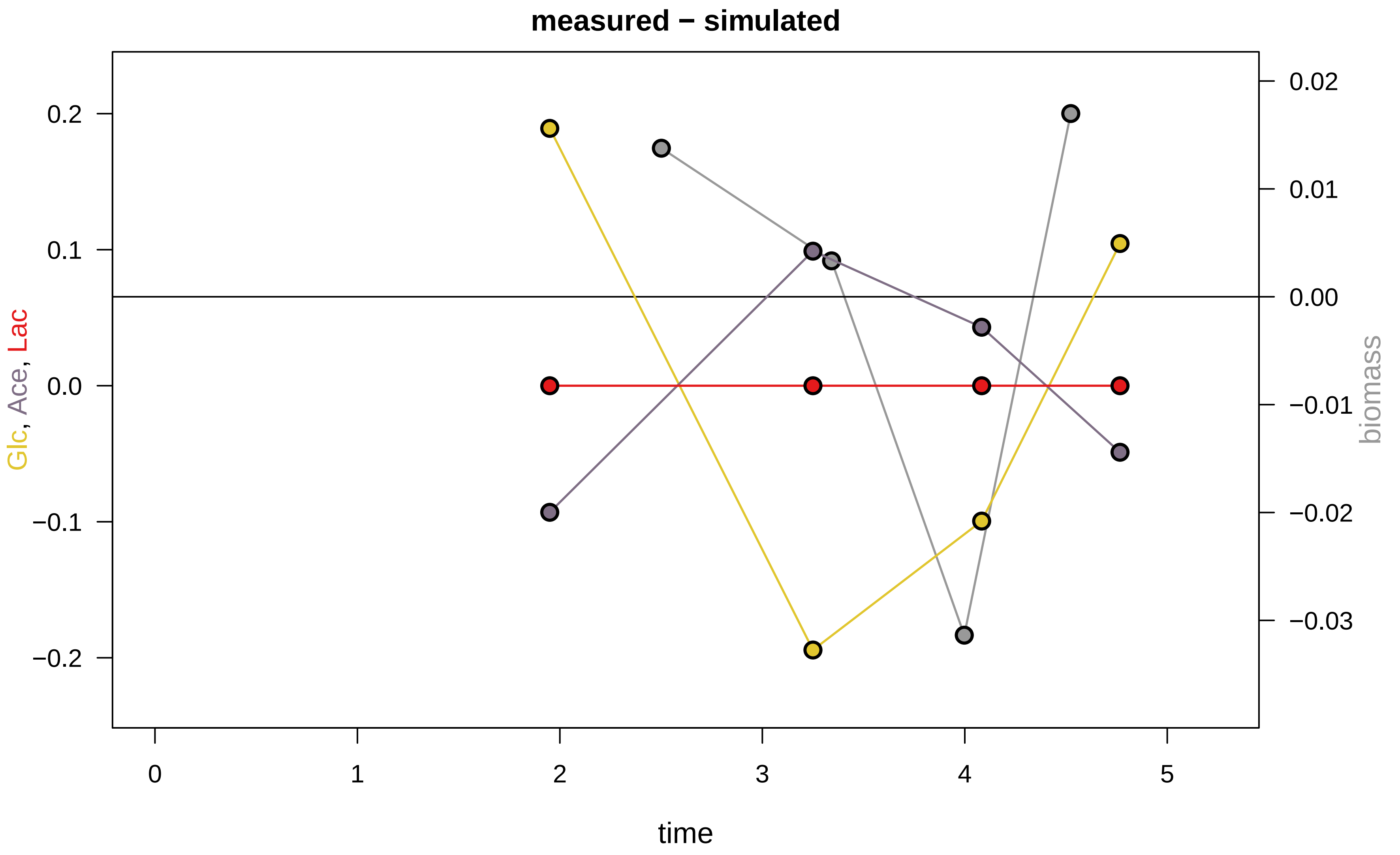

### KEIO_ROBOT2_8.pdf

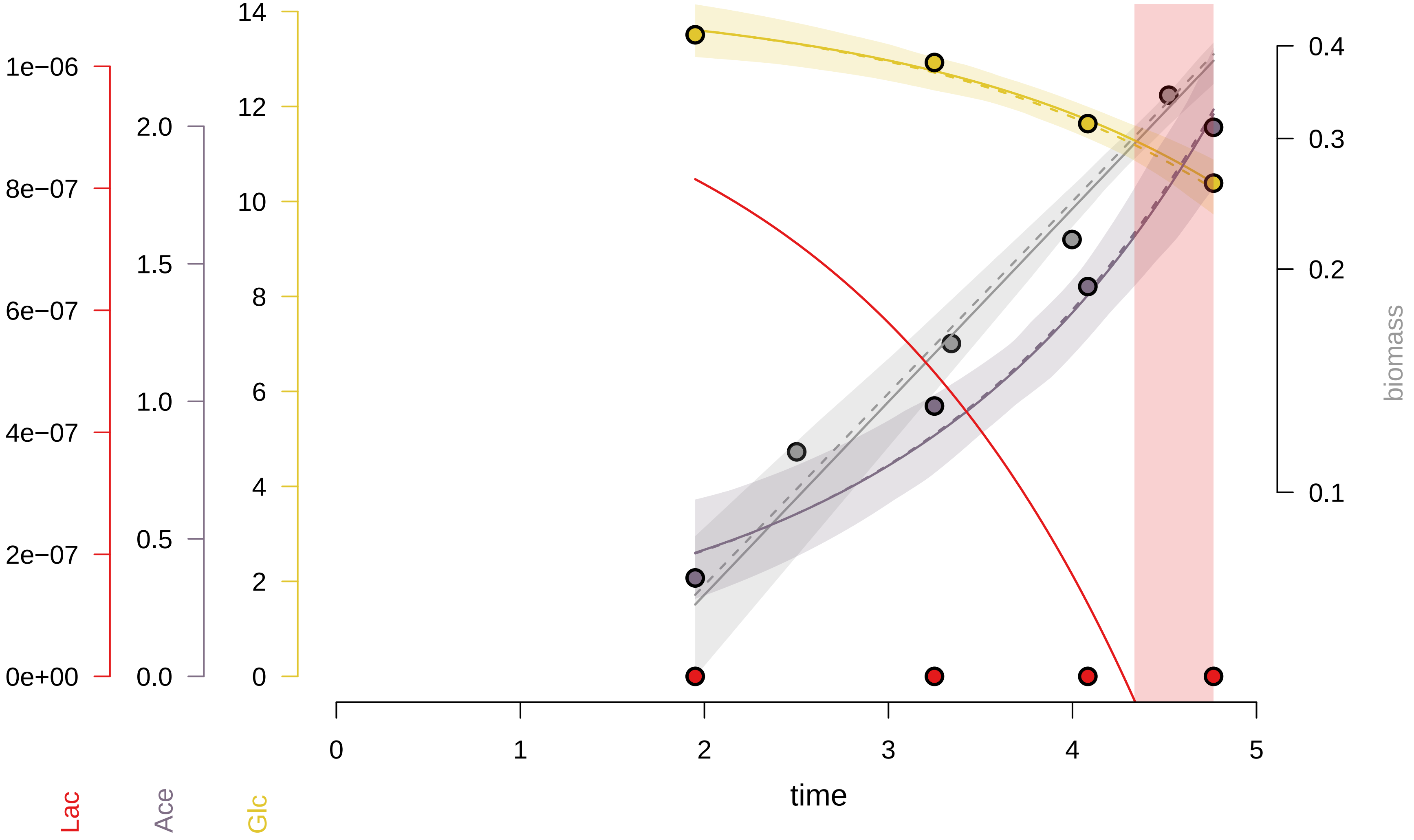

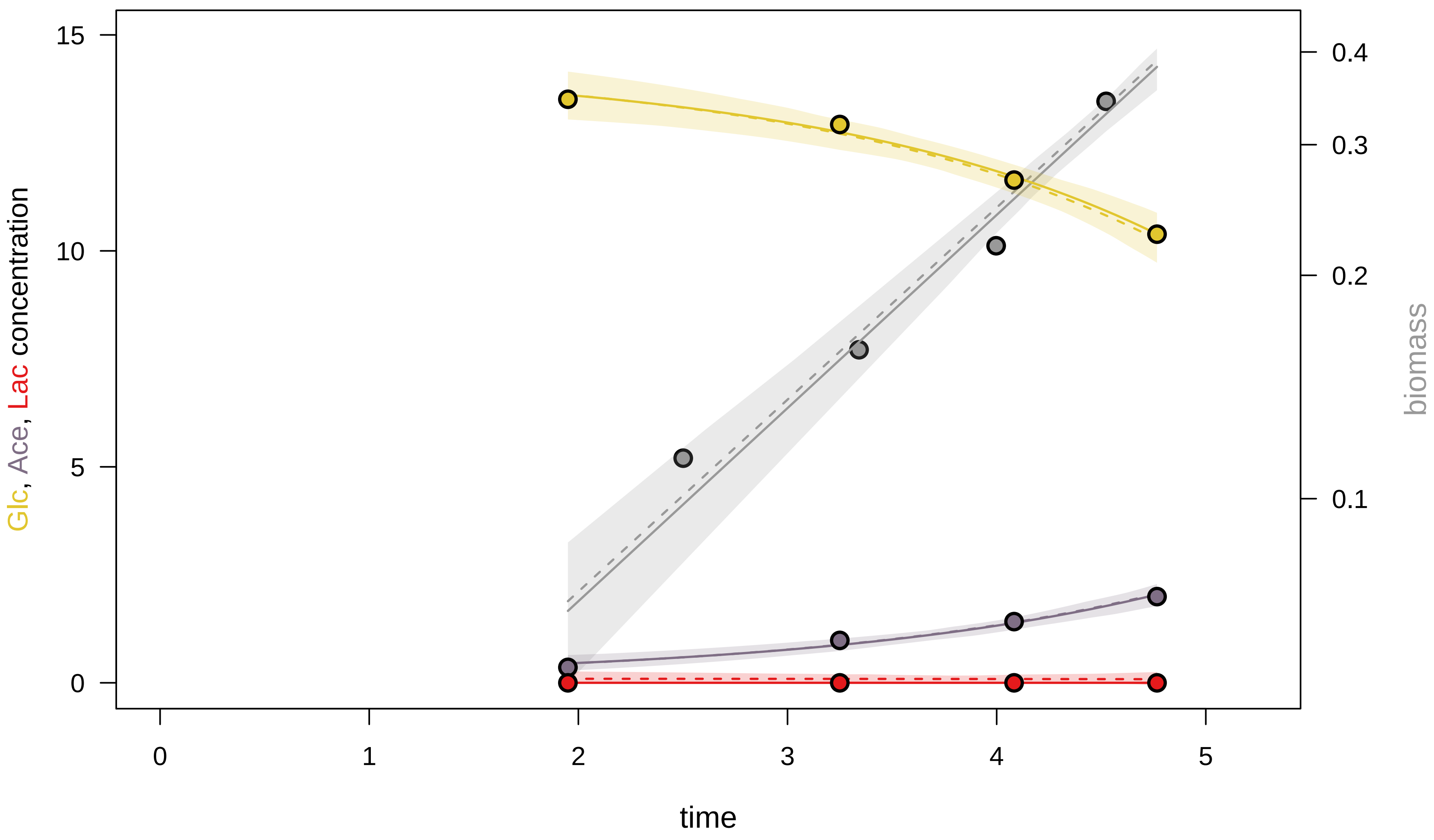

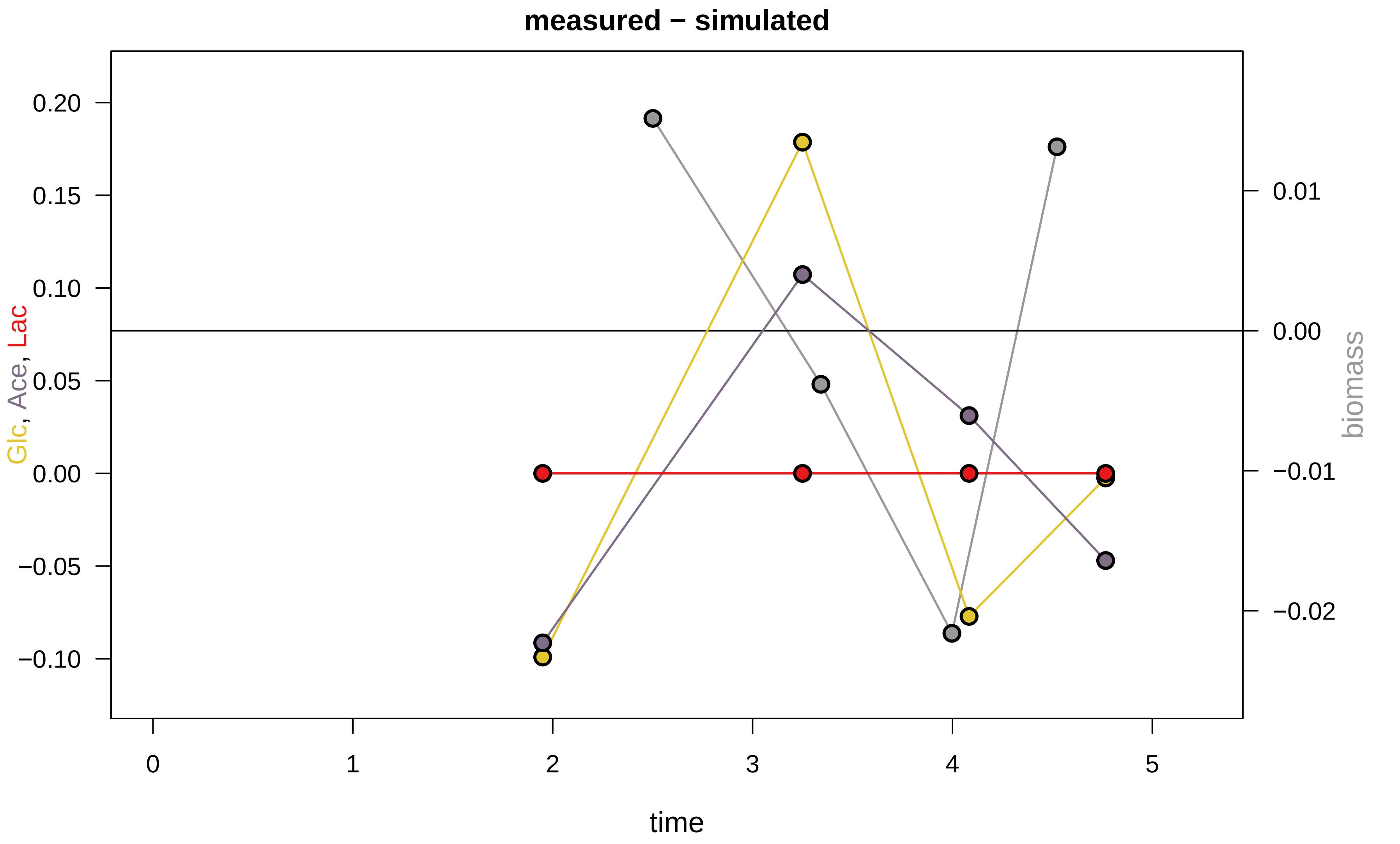

### KEIO_ROBOT2_15.pdf

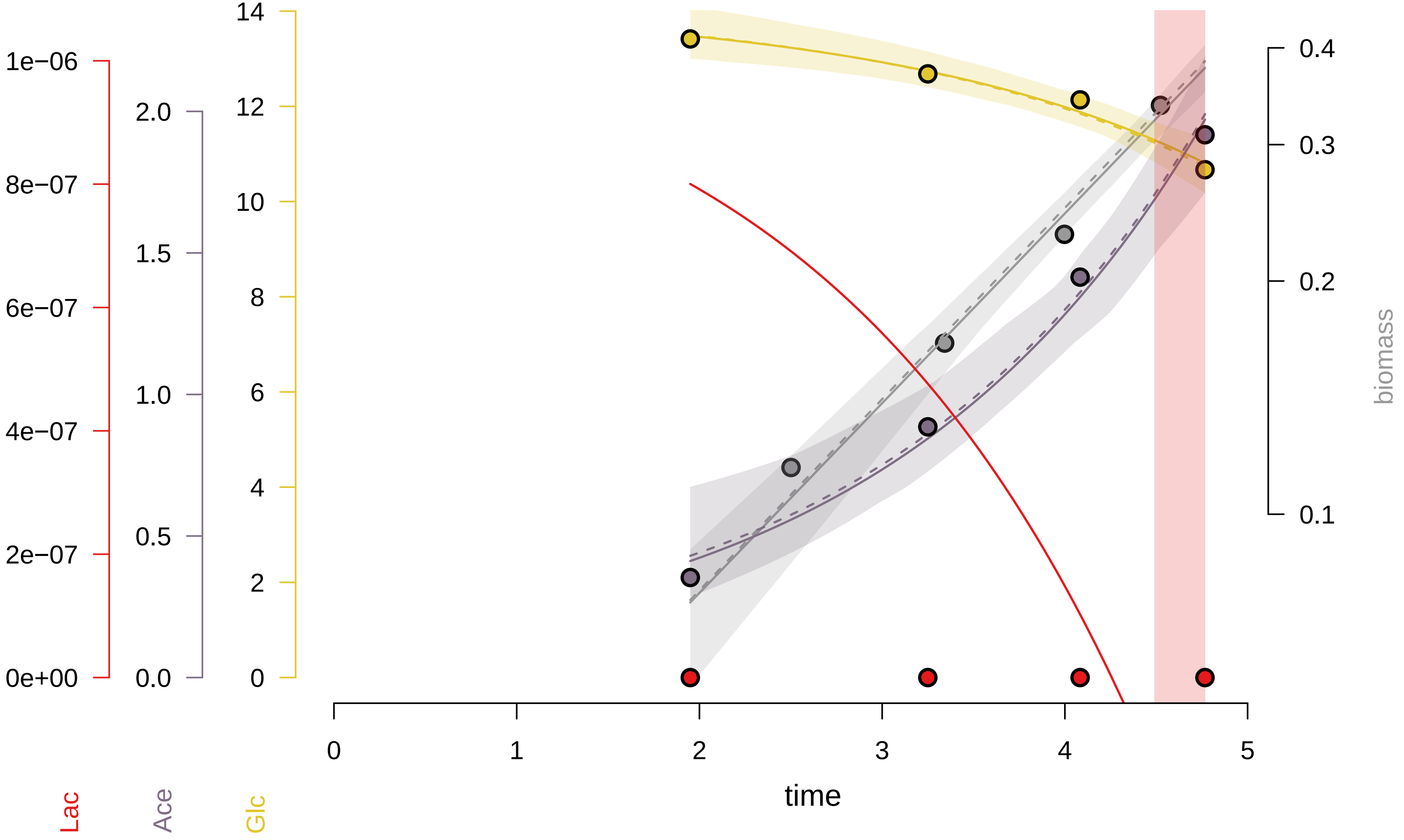

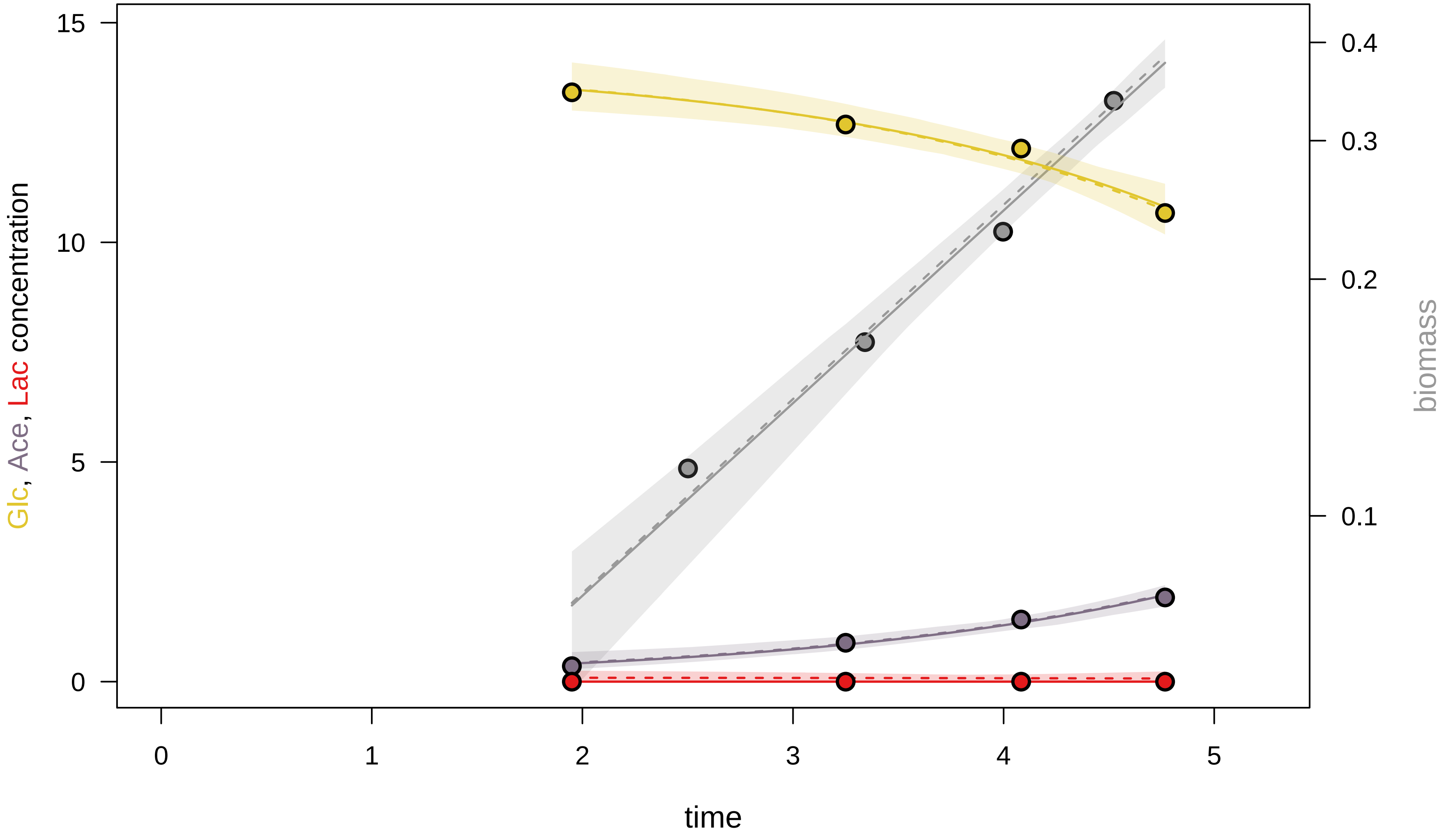

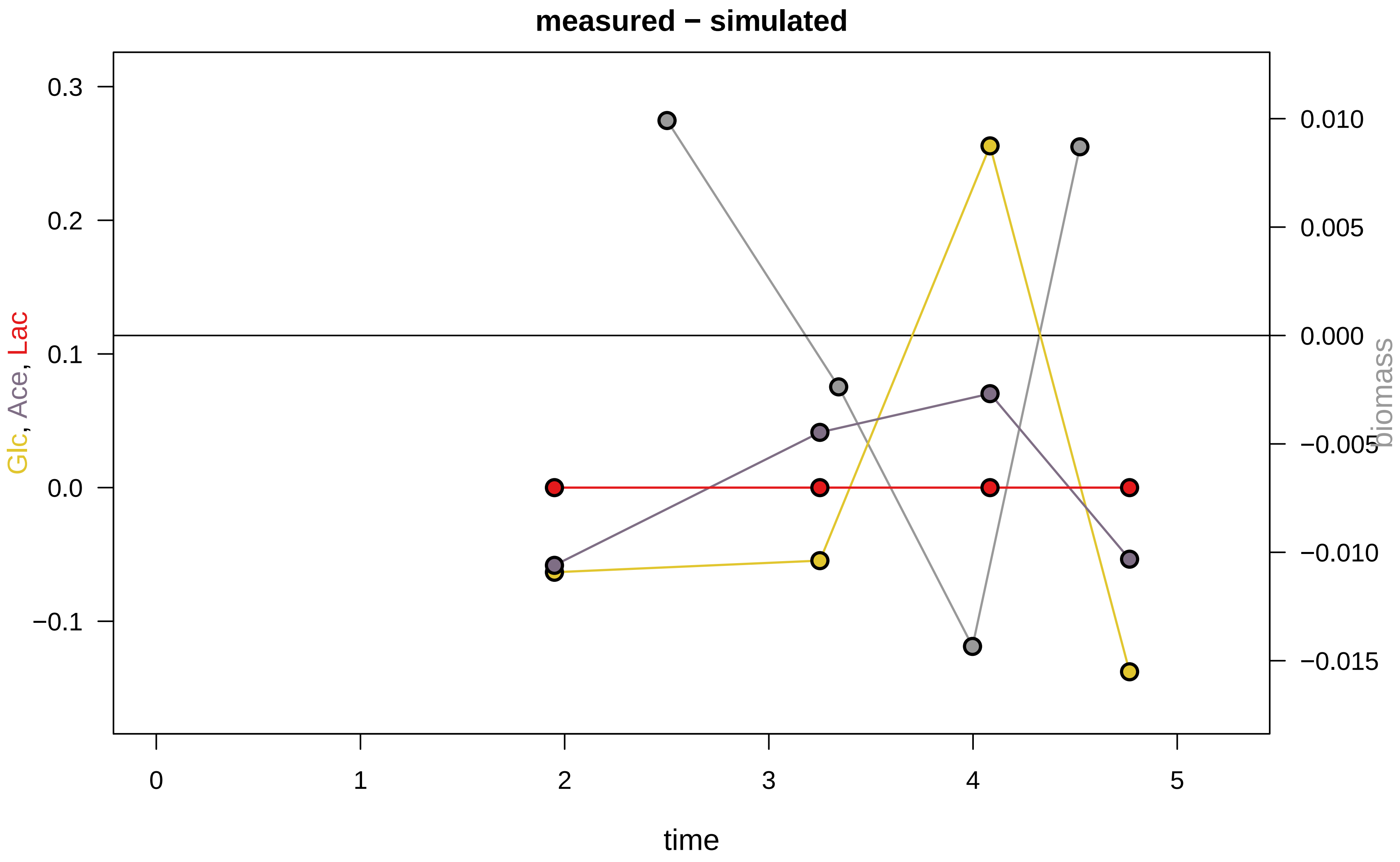

### KEIO_ROBOT2_16.pdf

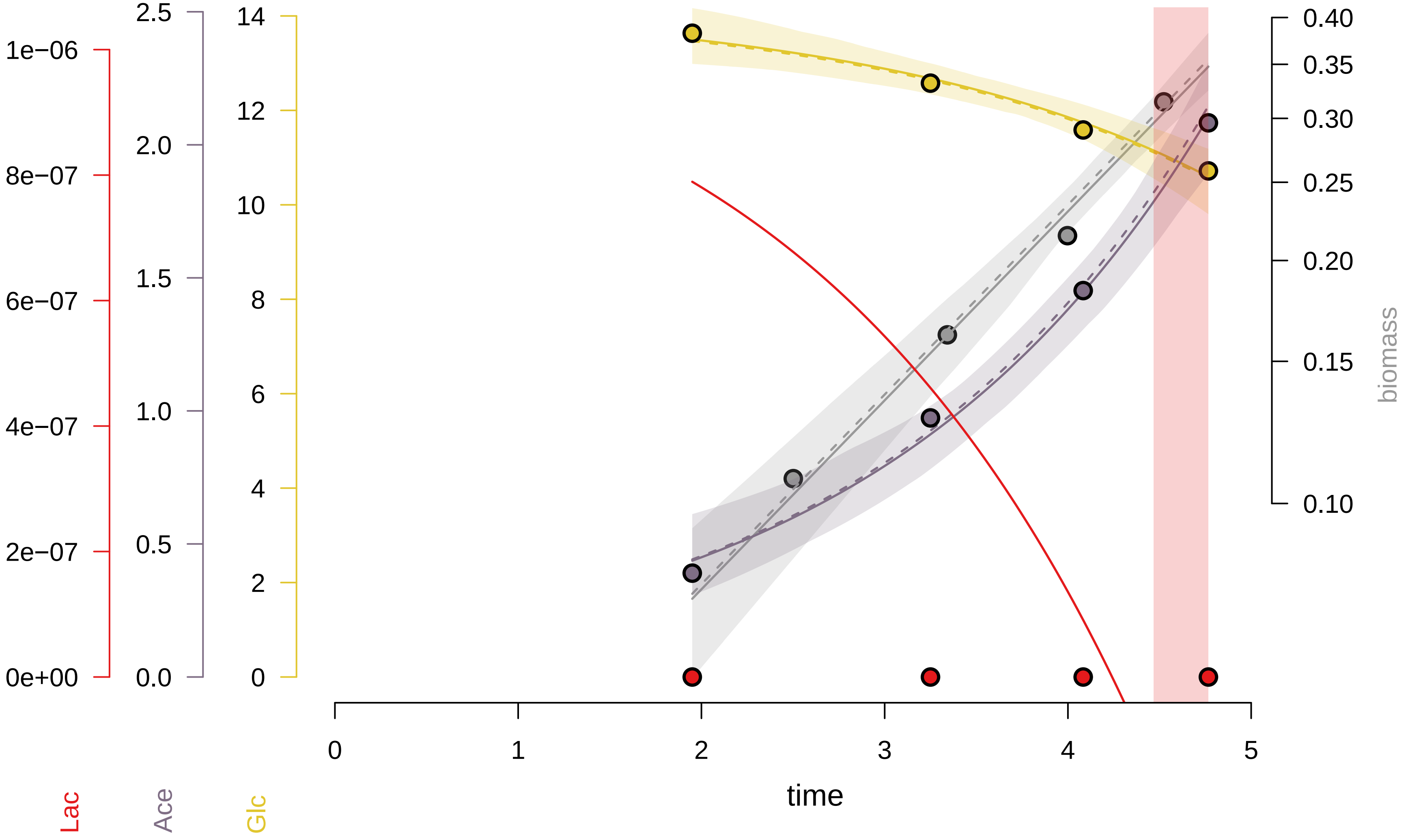

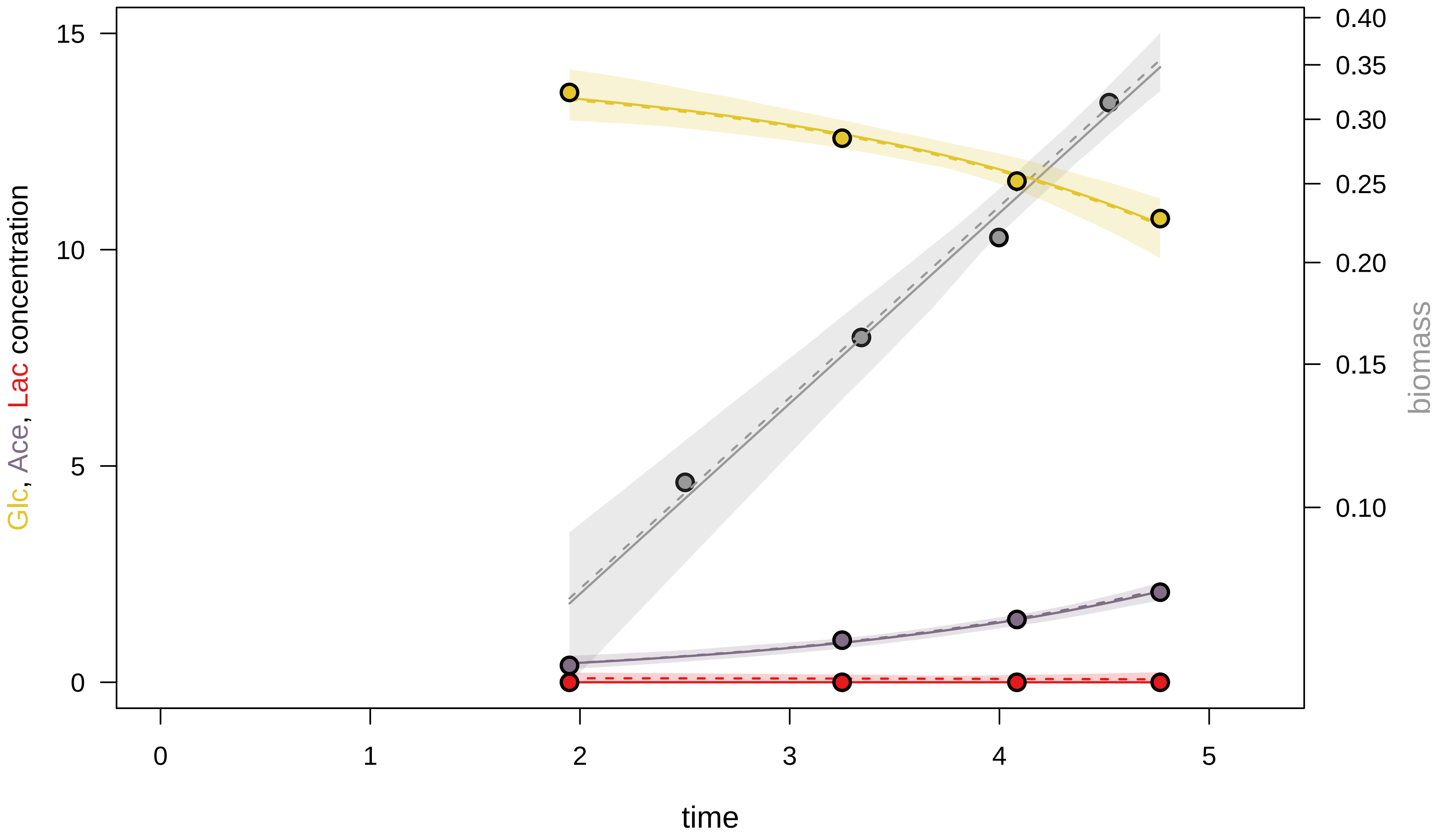

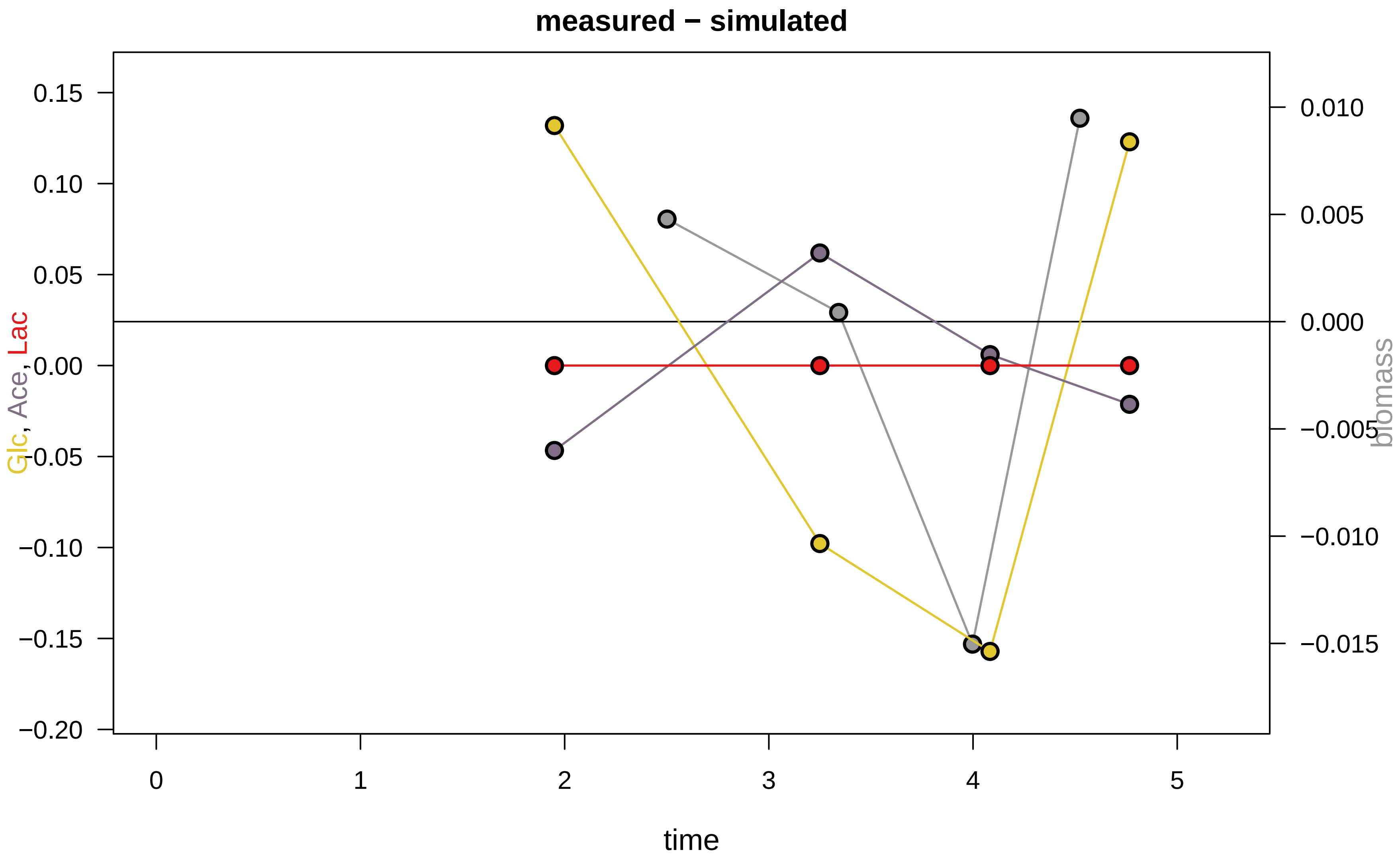

### KEIO_ROBOT2_17.pdf

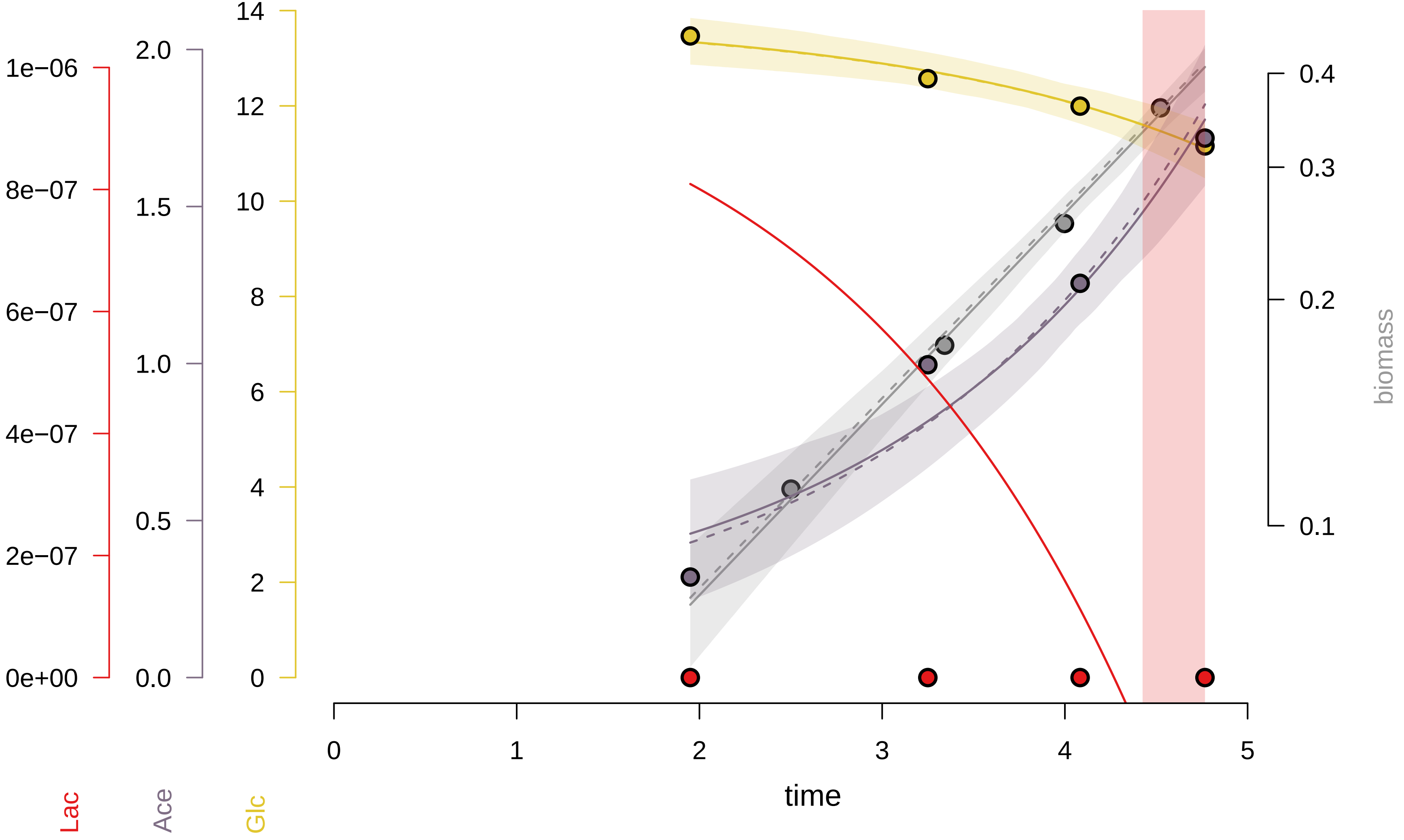

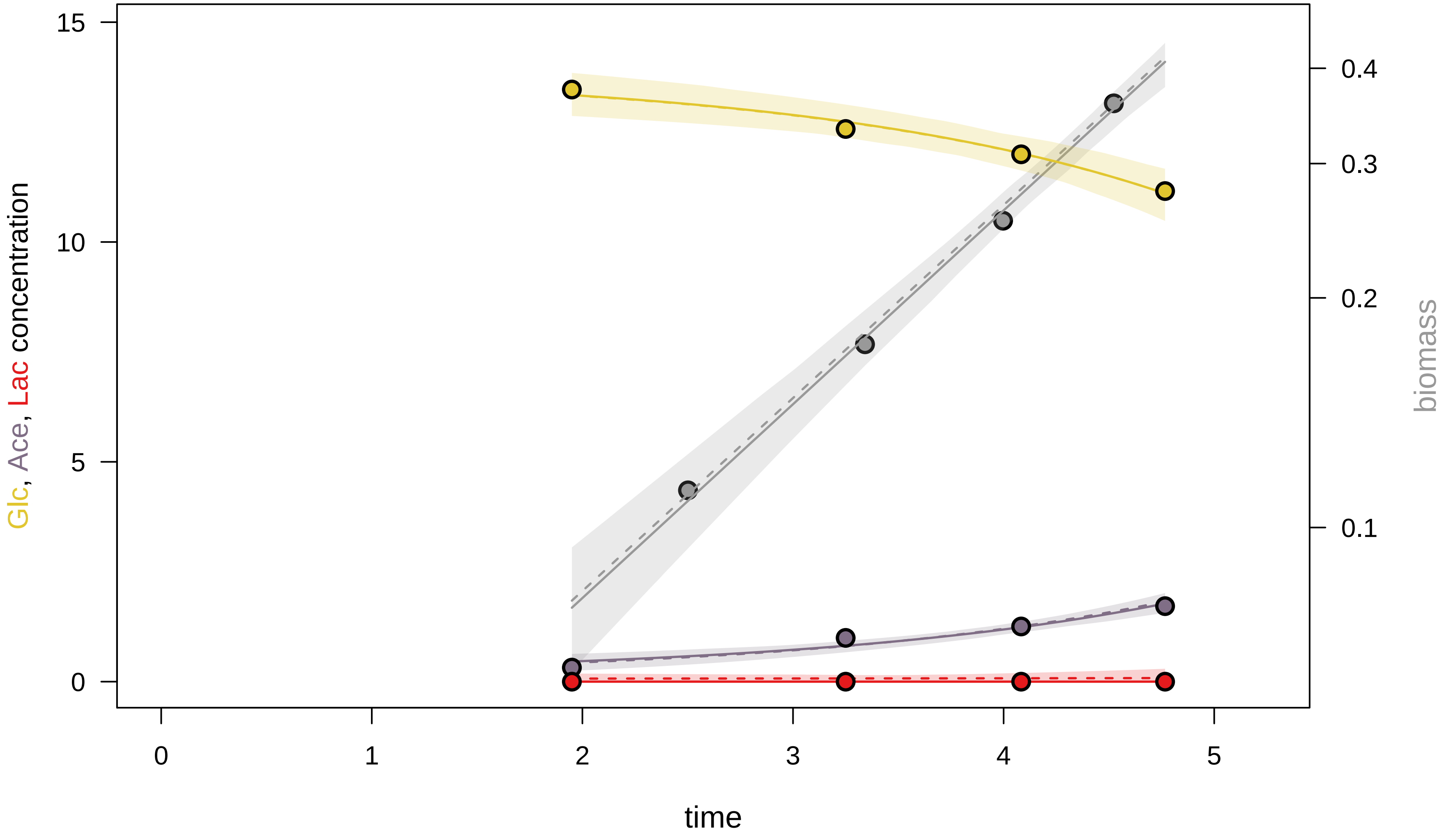

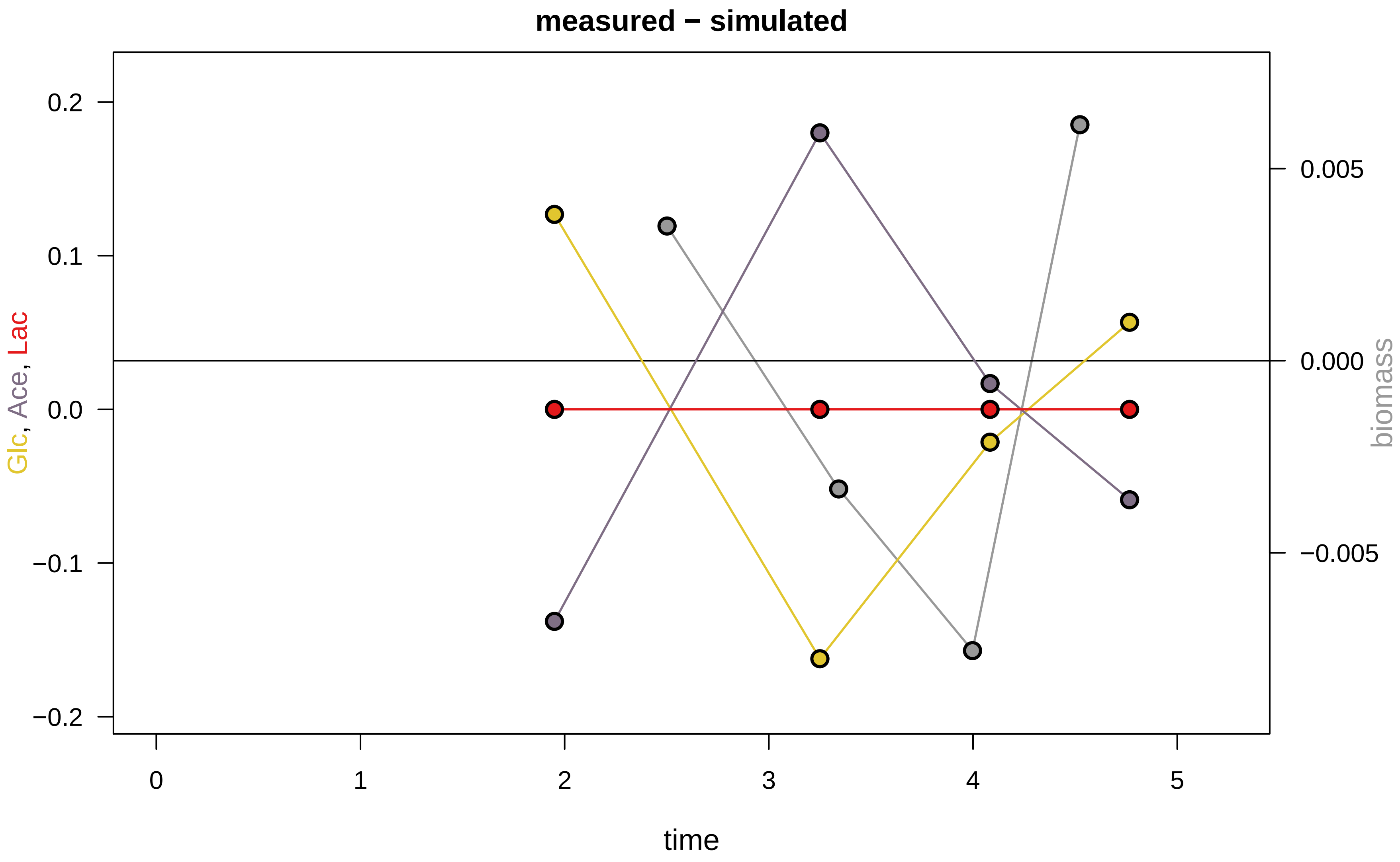

### KEIO_ROBOT2_19.pdf

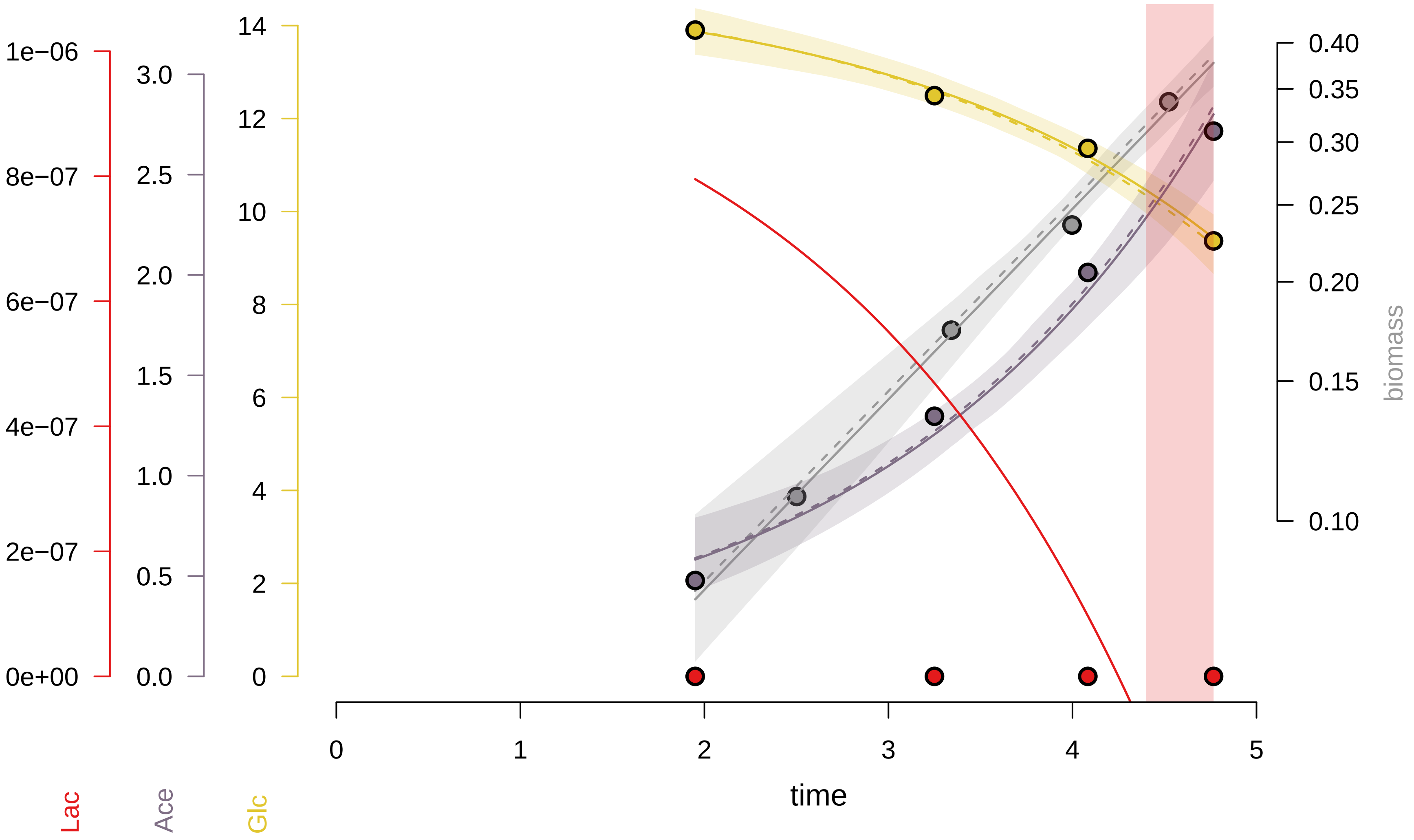

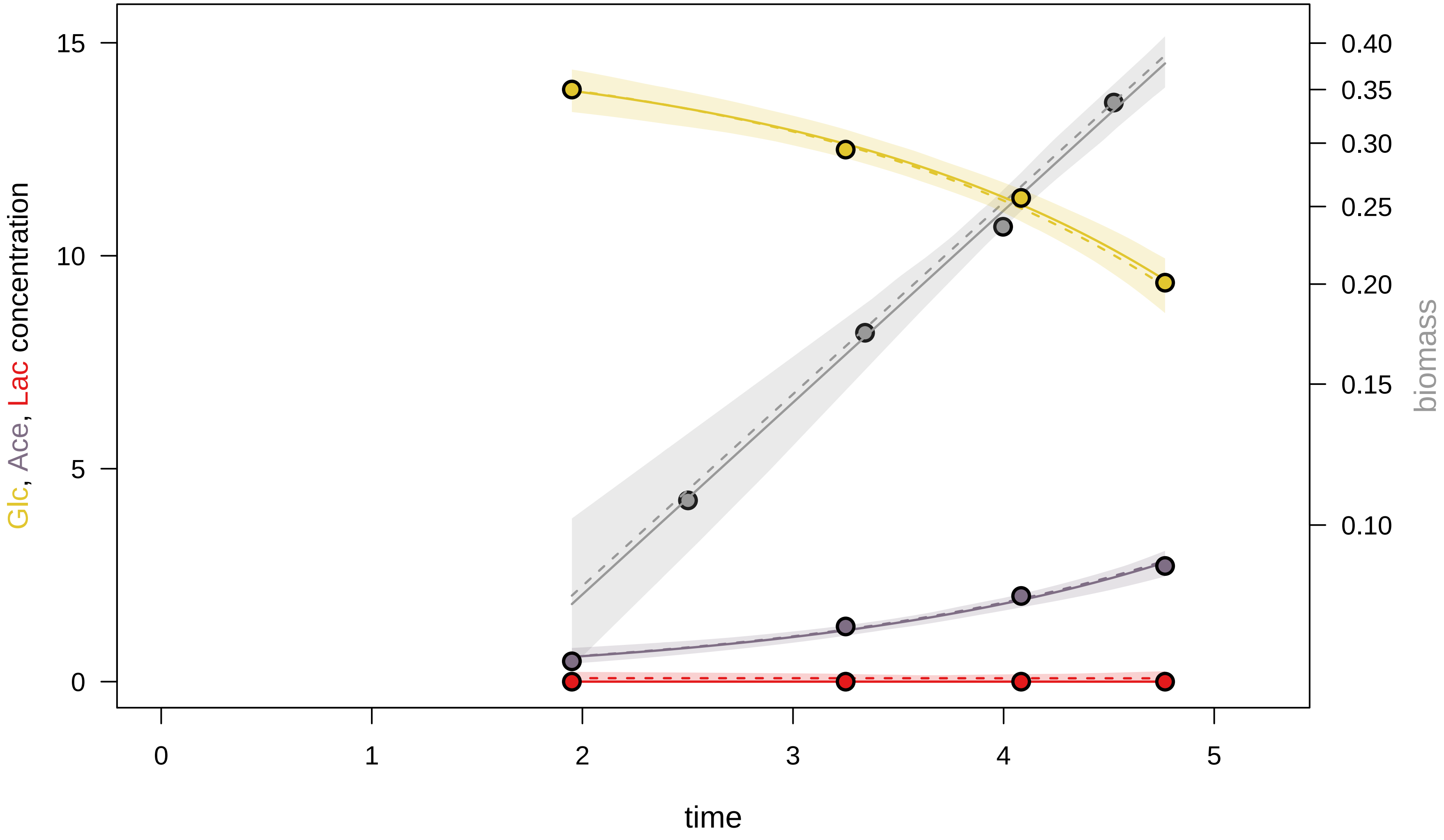

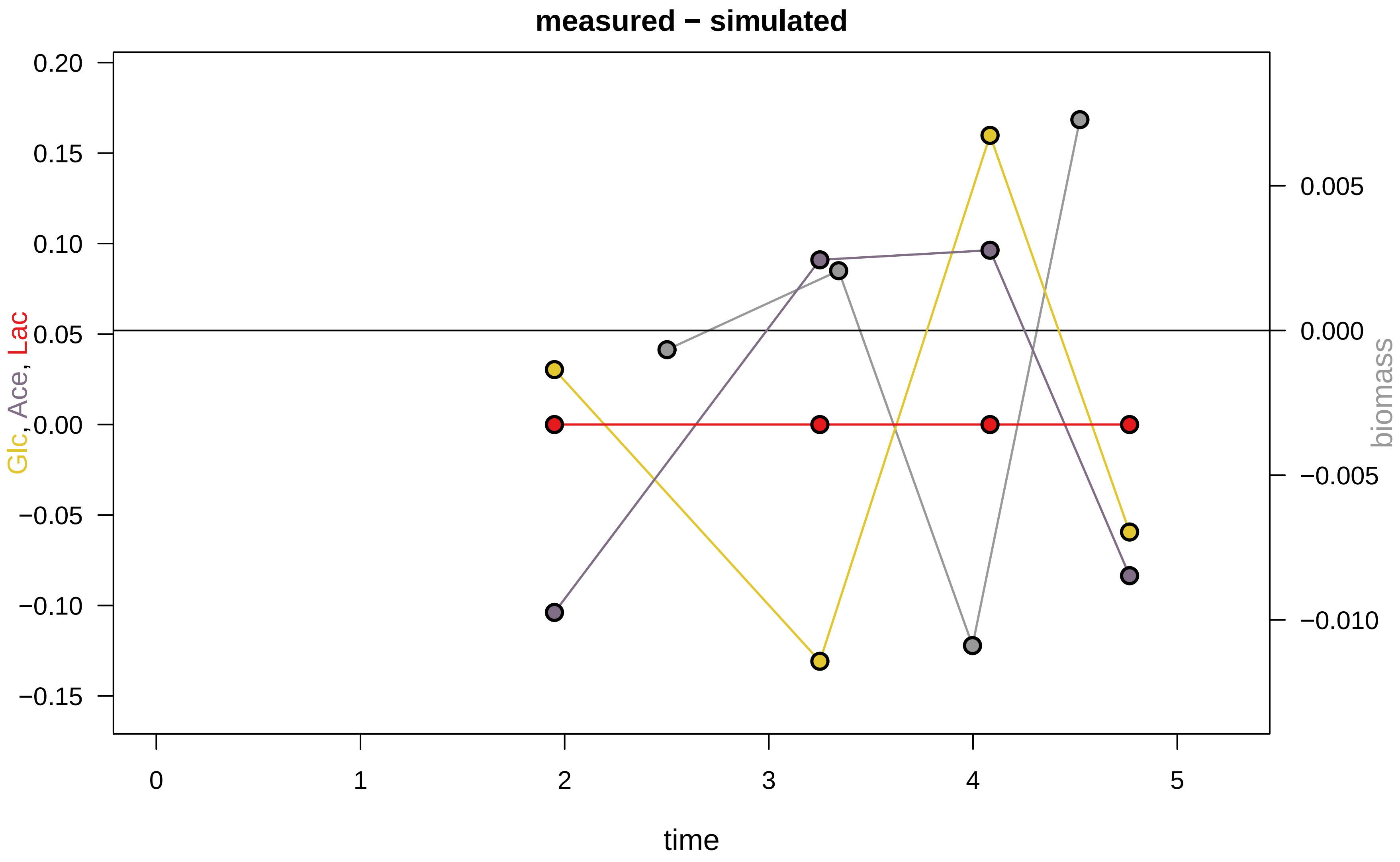

### KEIO_ROBOT2_22.pdf

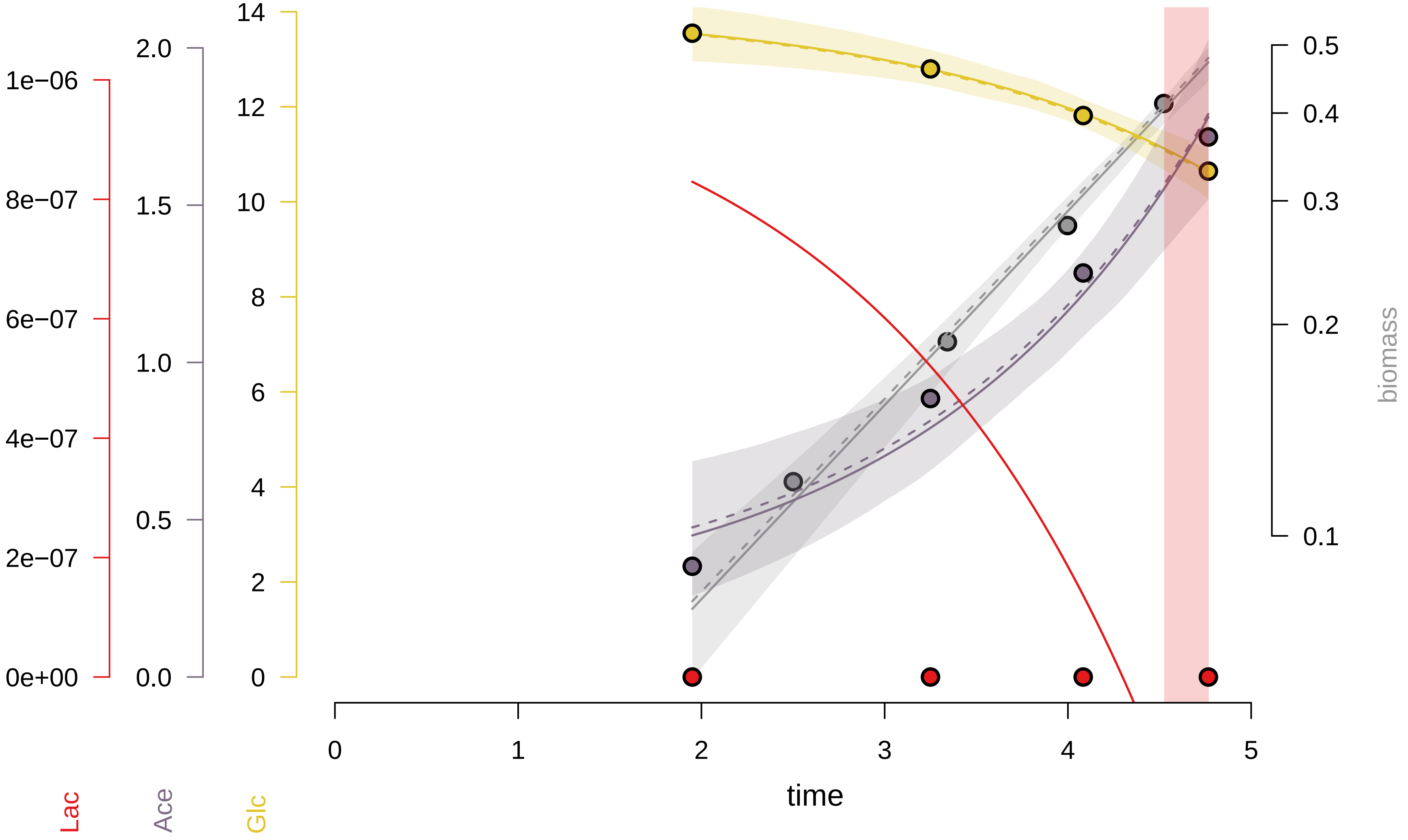

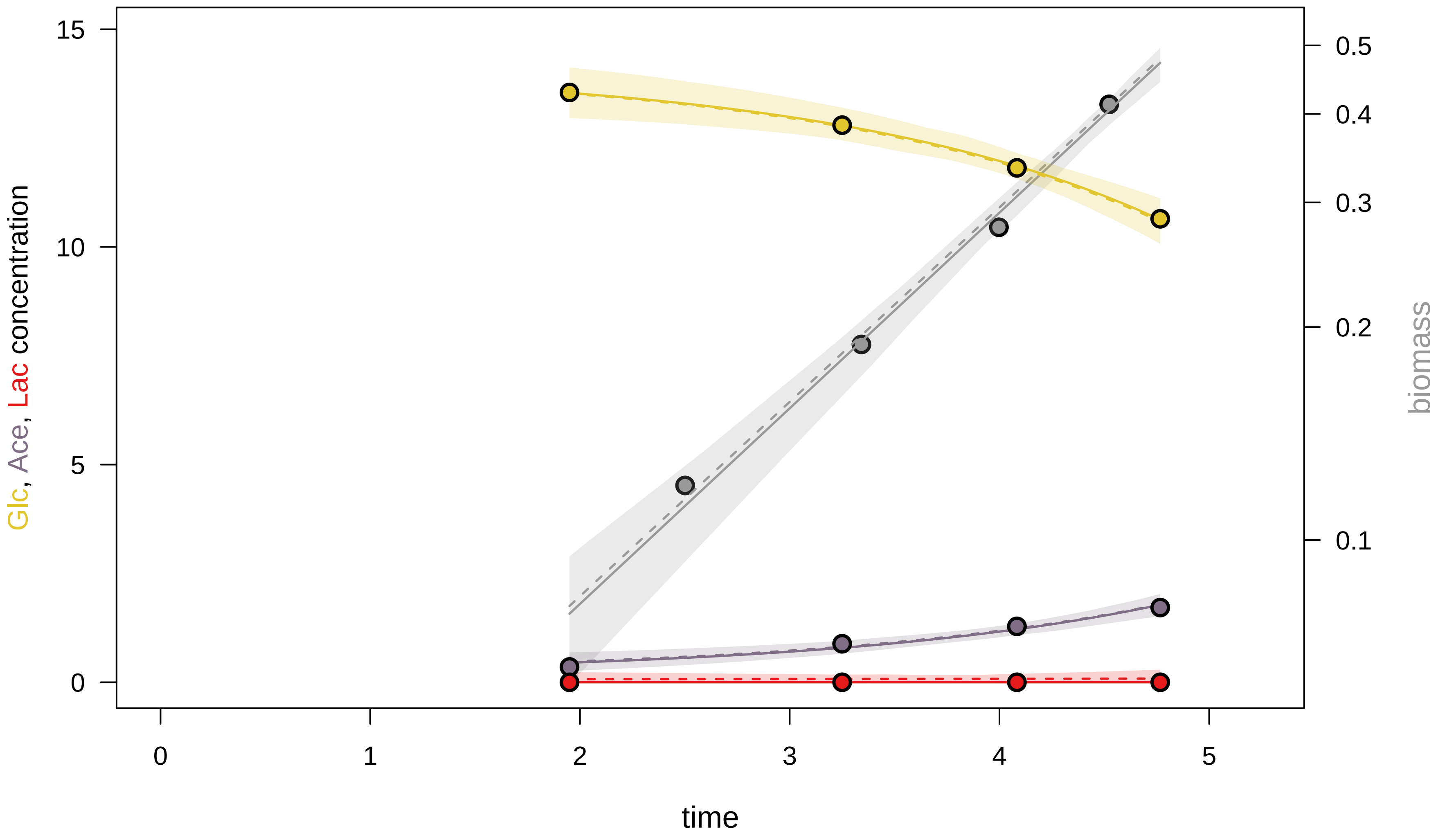

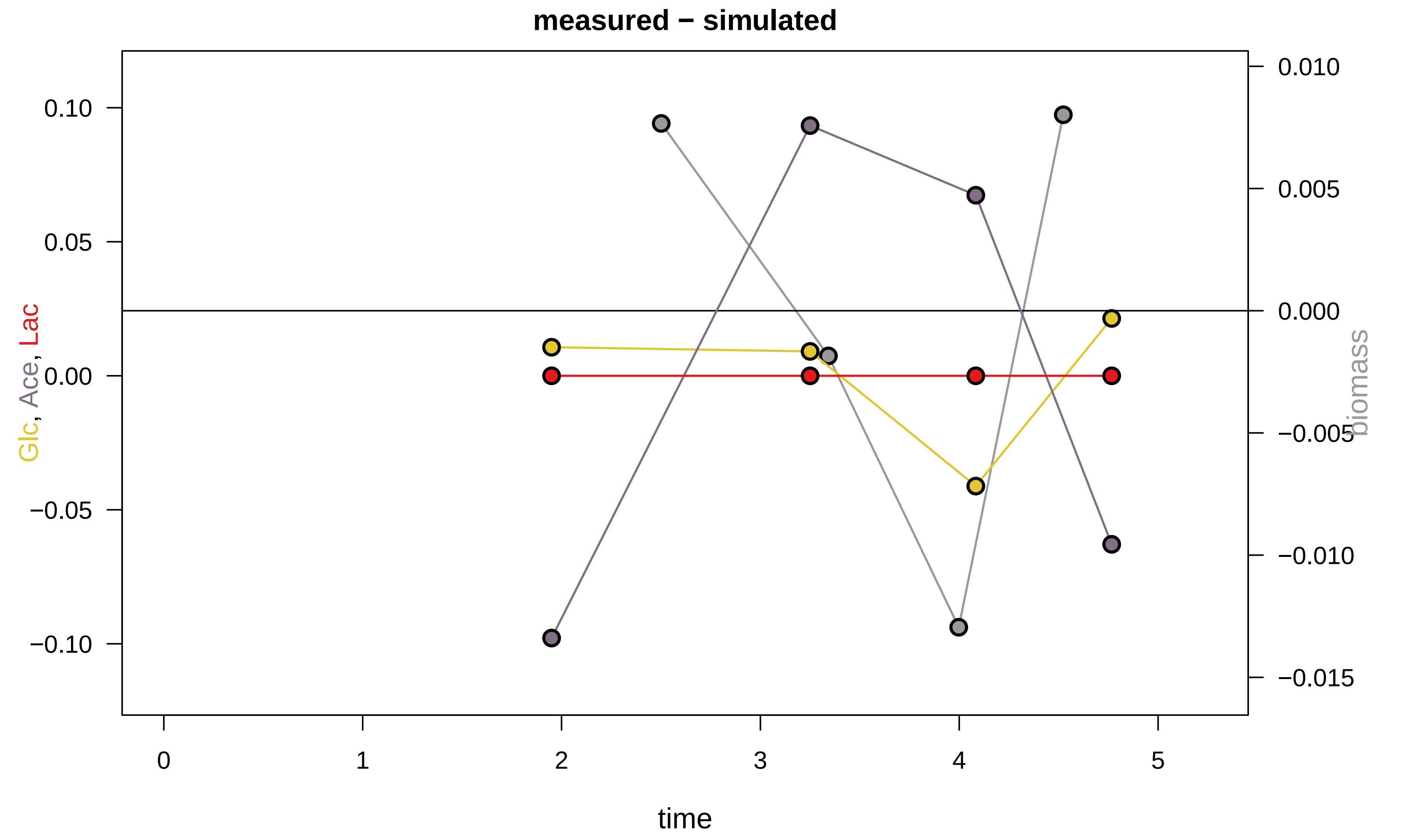

### KEIO_ROBOT2_24.pdf

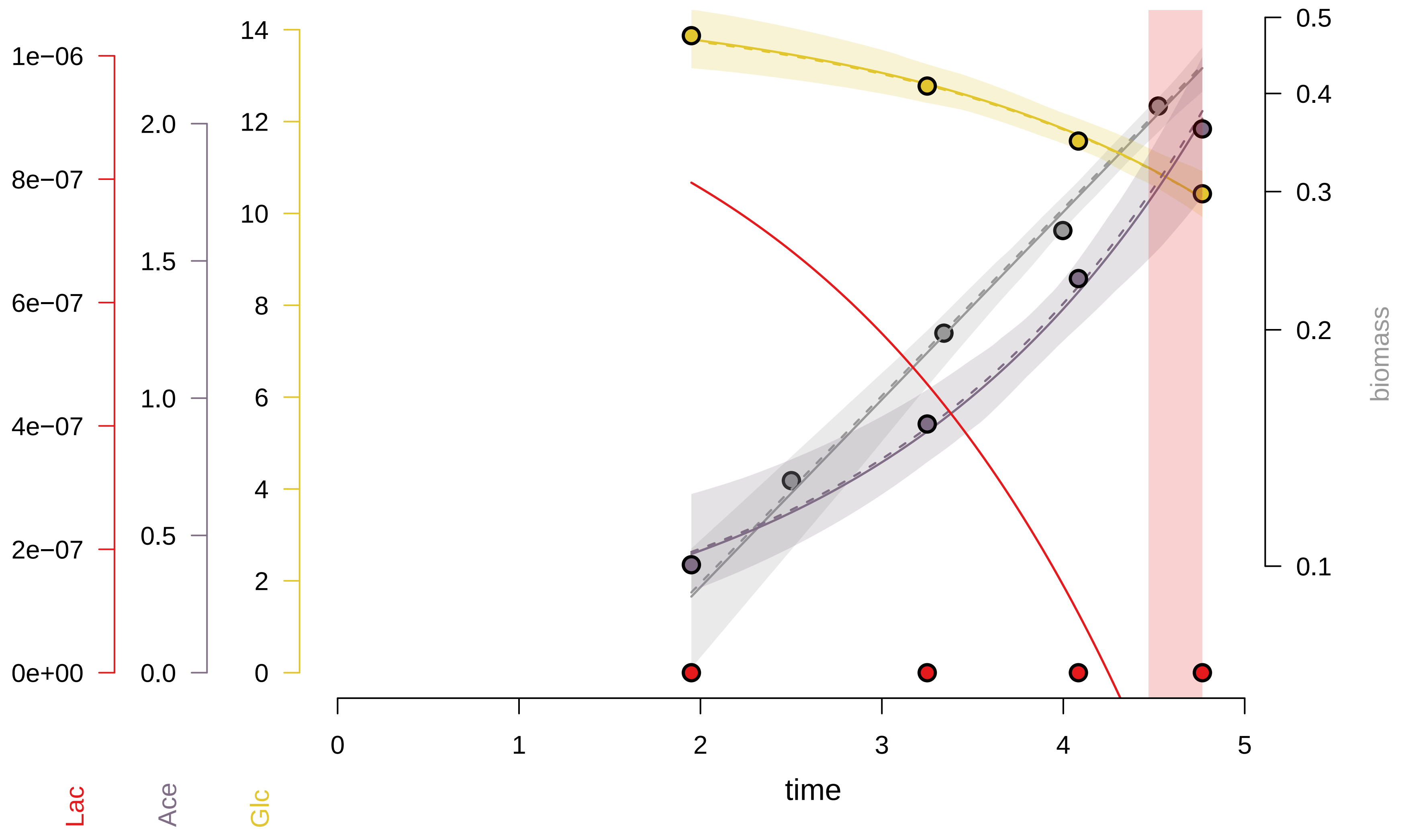

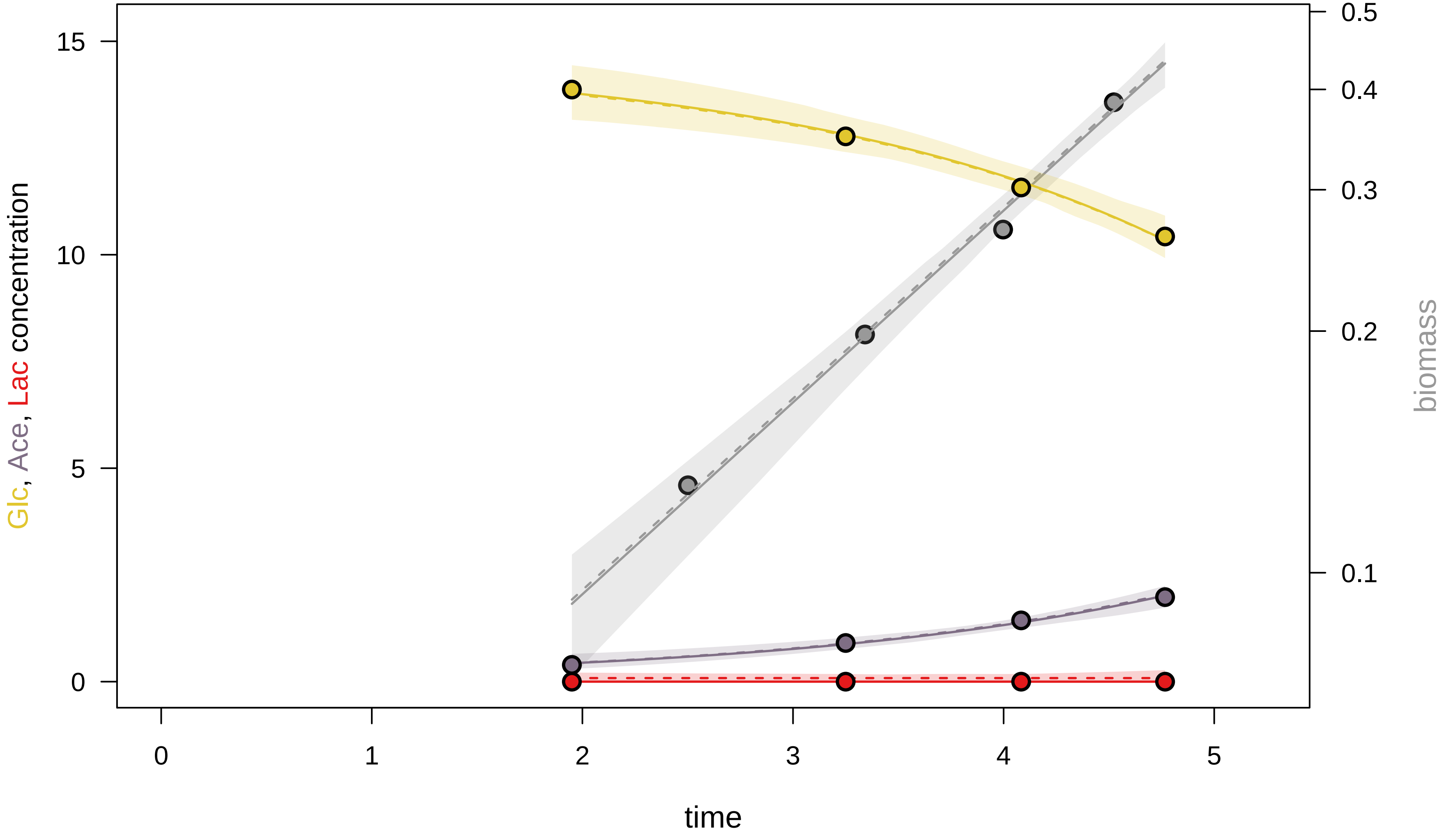

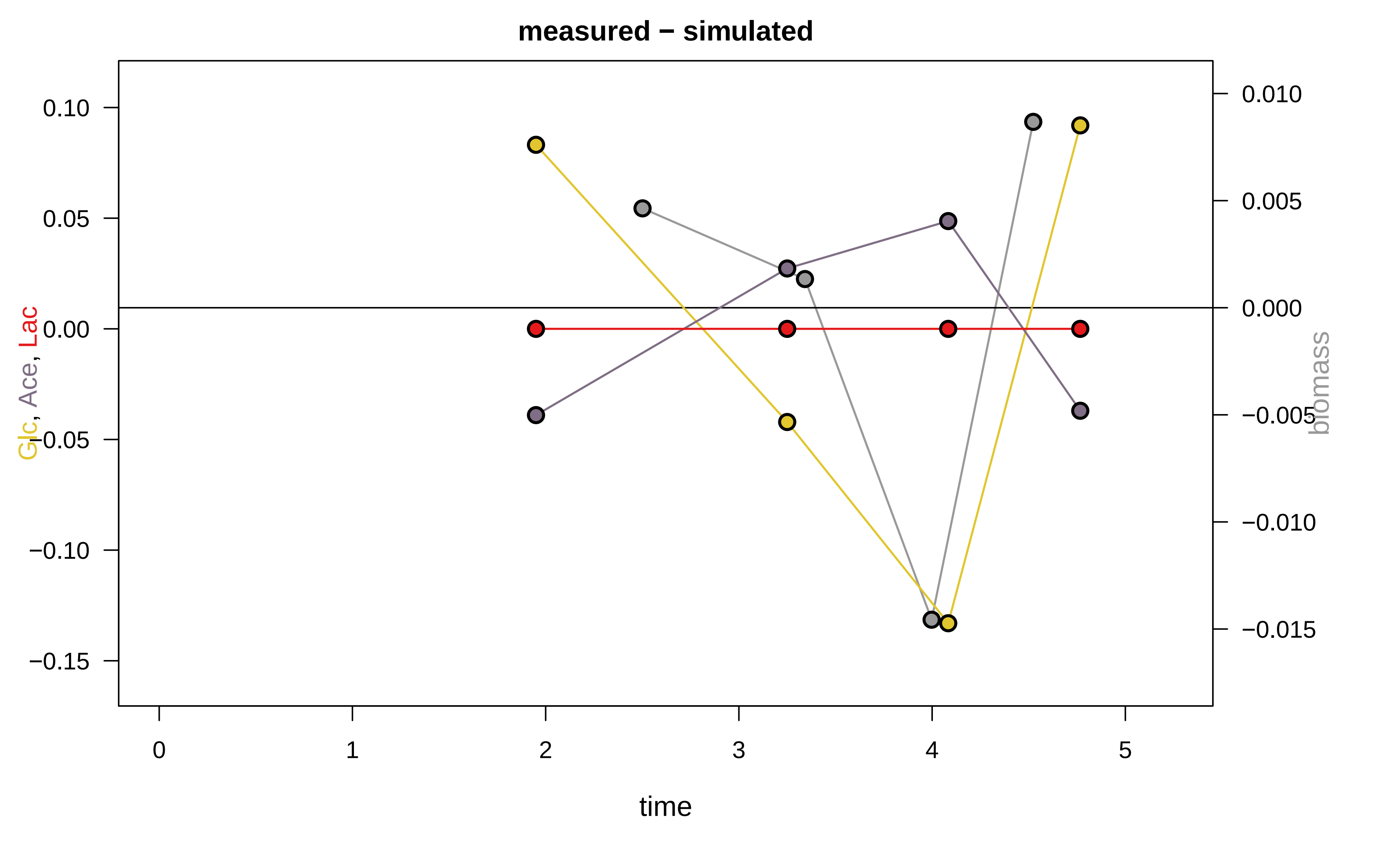

### KEIO_ROBOT2_28.pdf

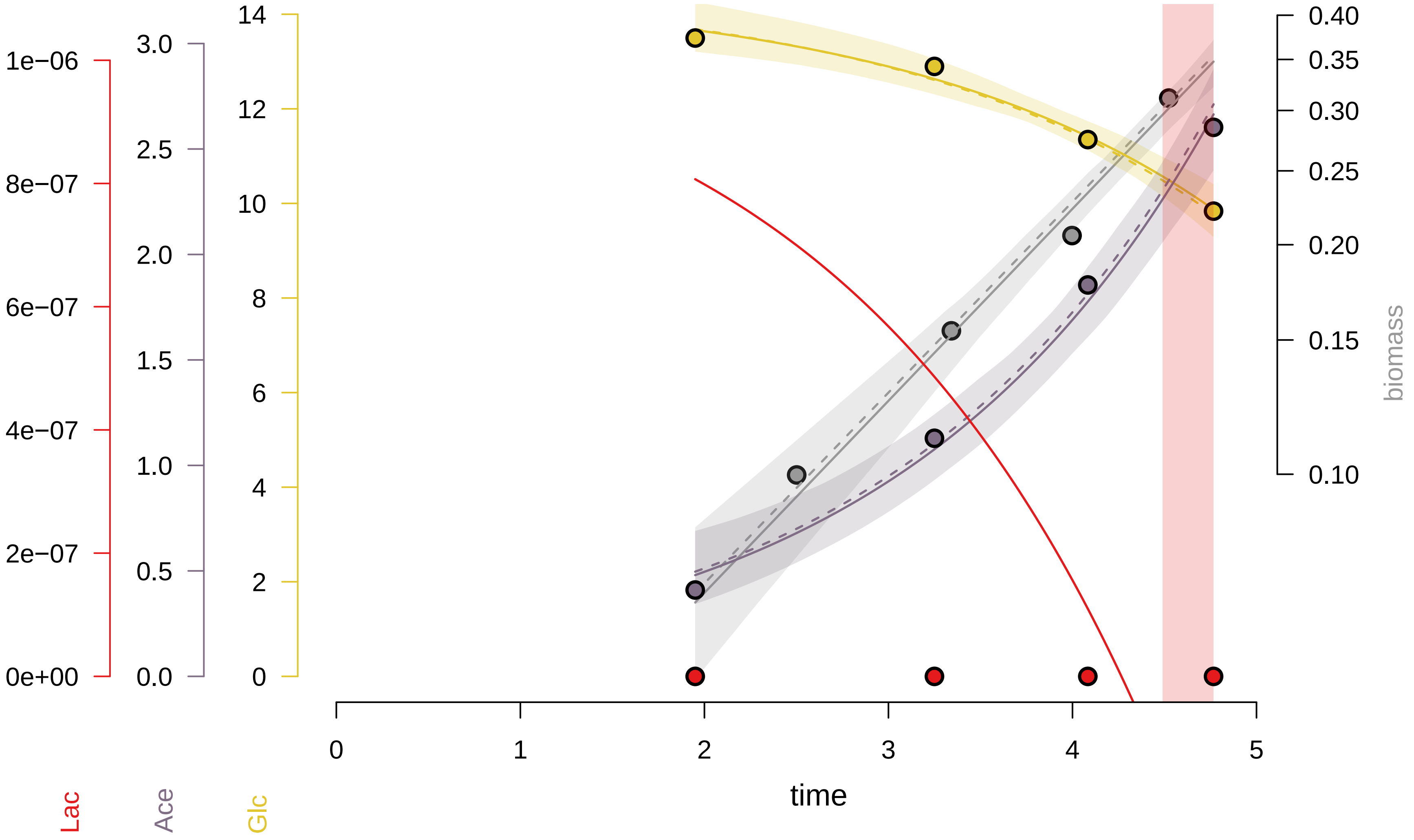

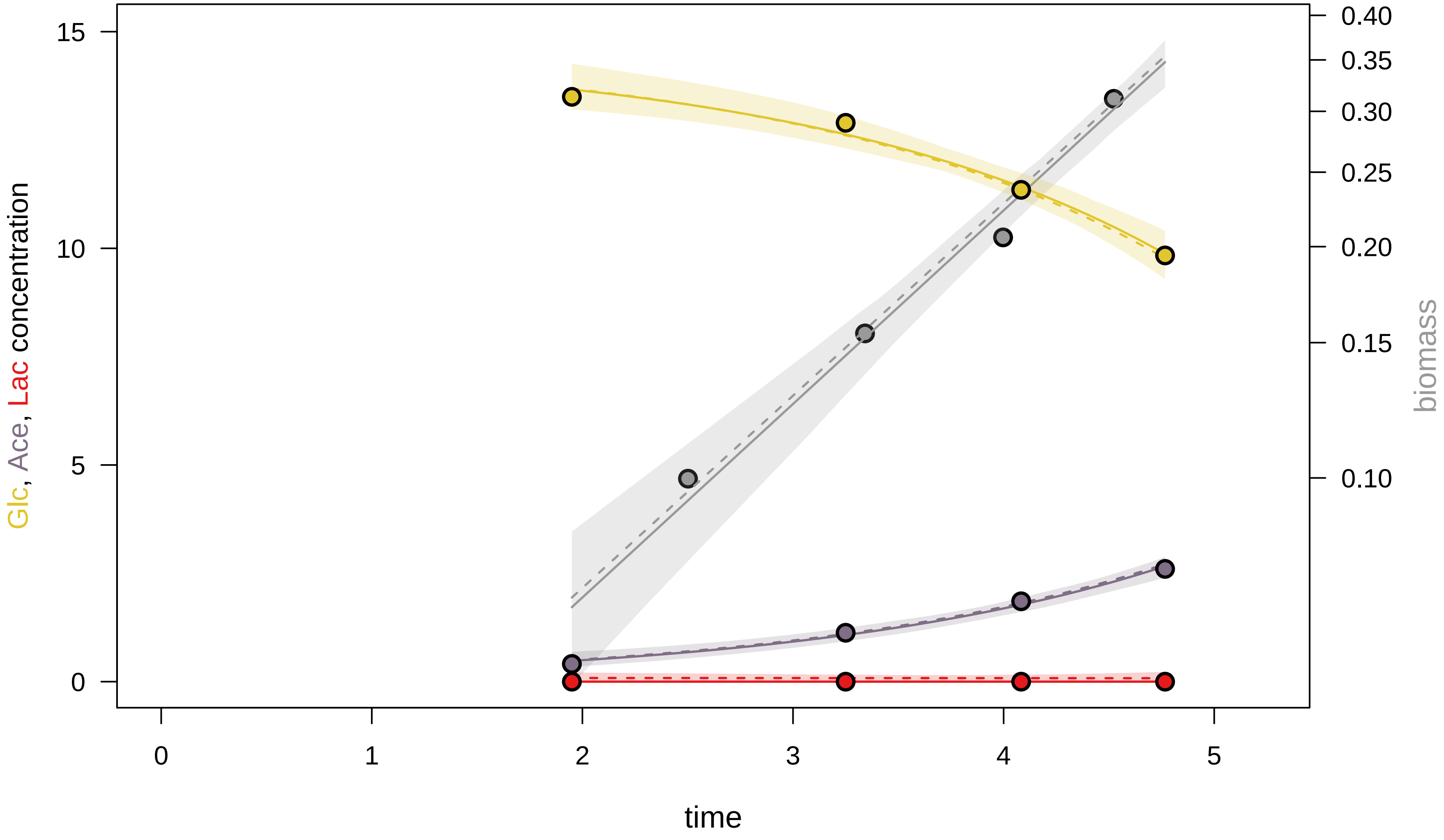

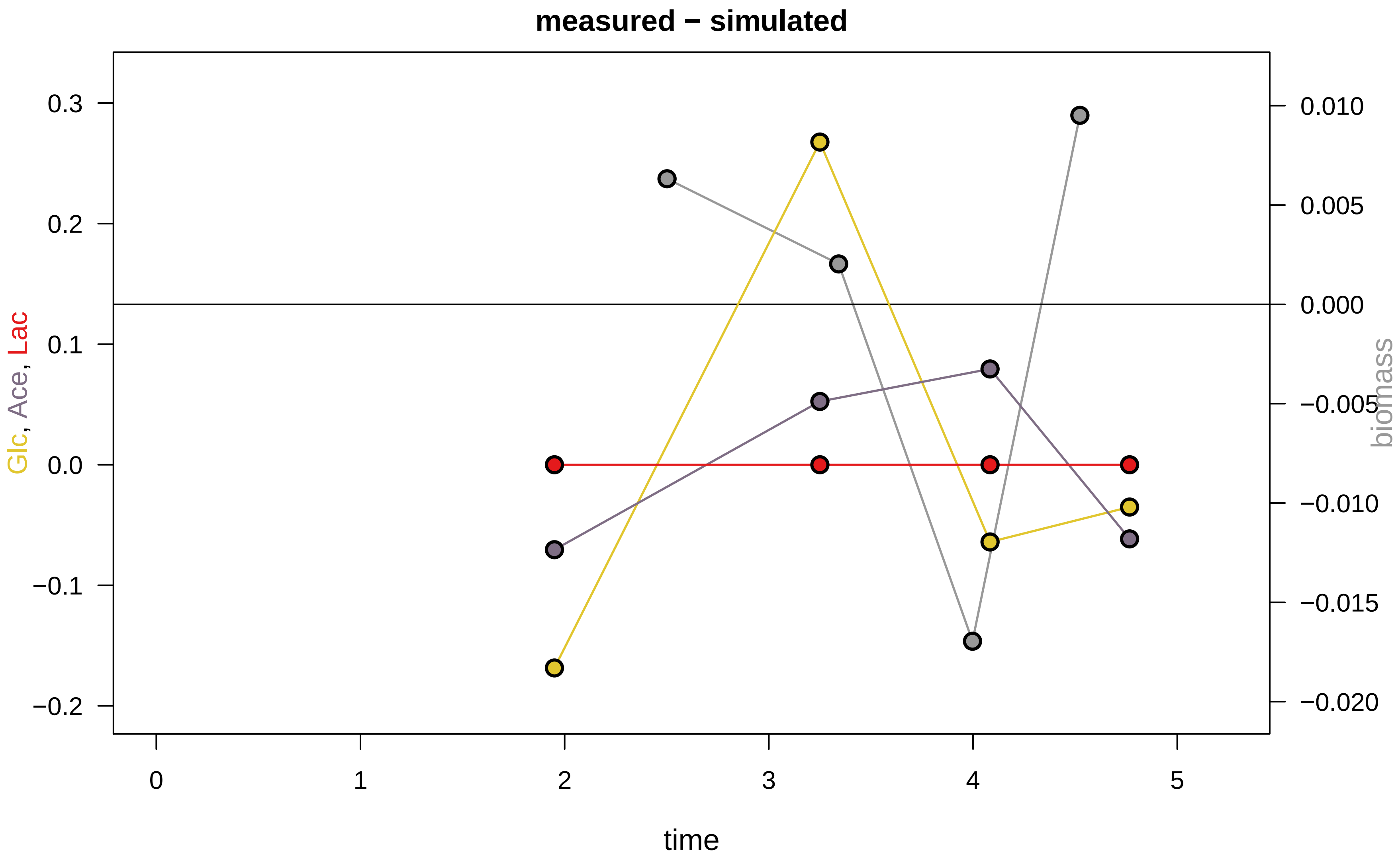

### KEIO_ROBOT2_32.pdf

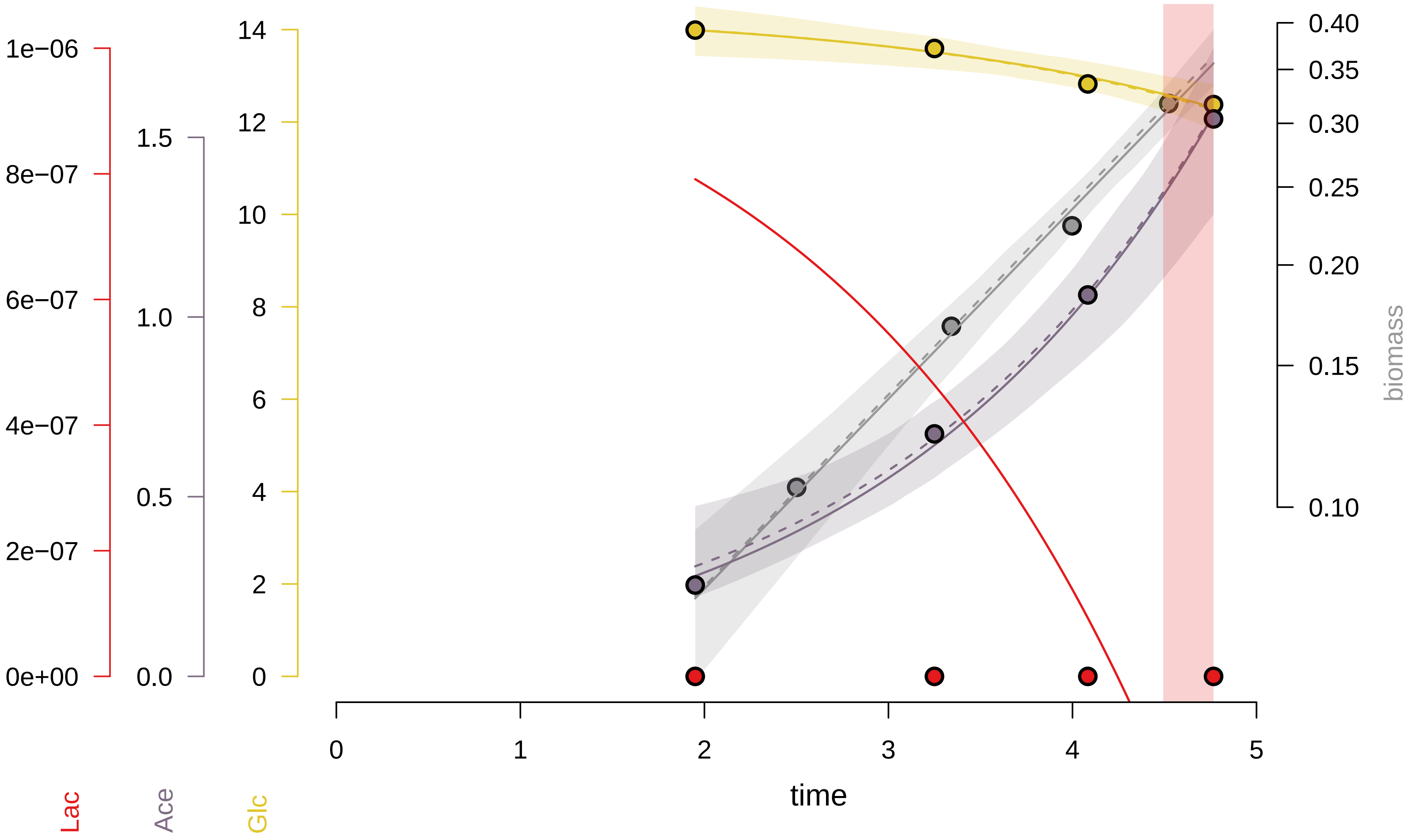

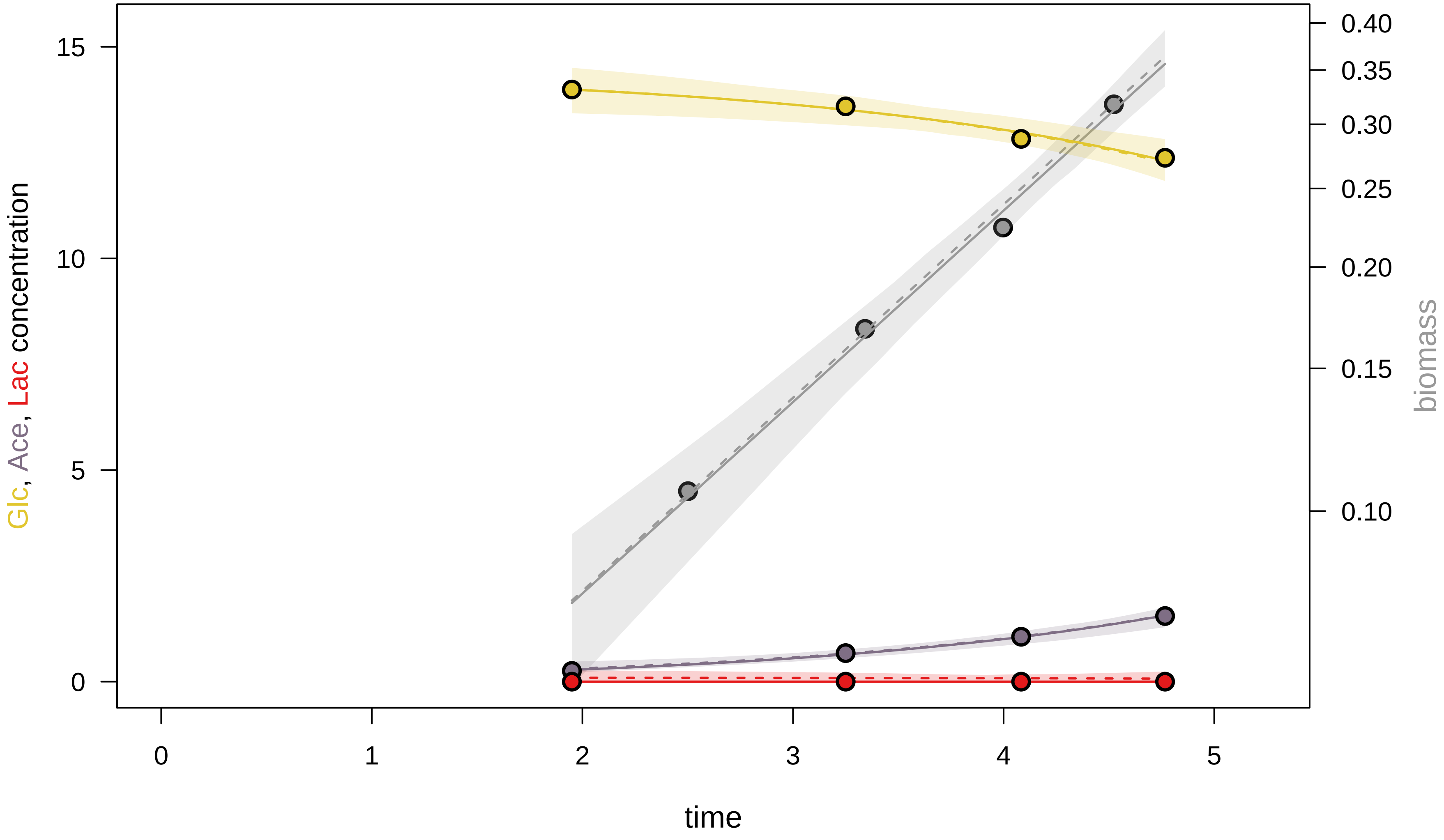

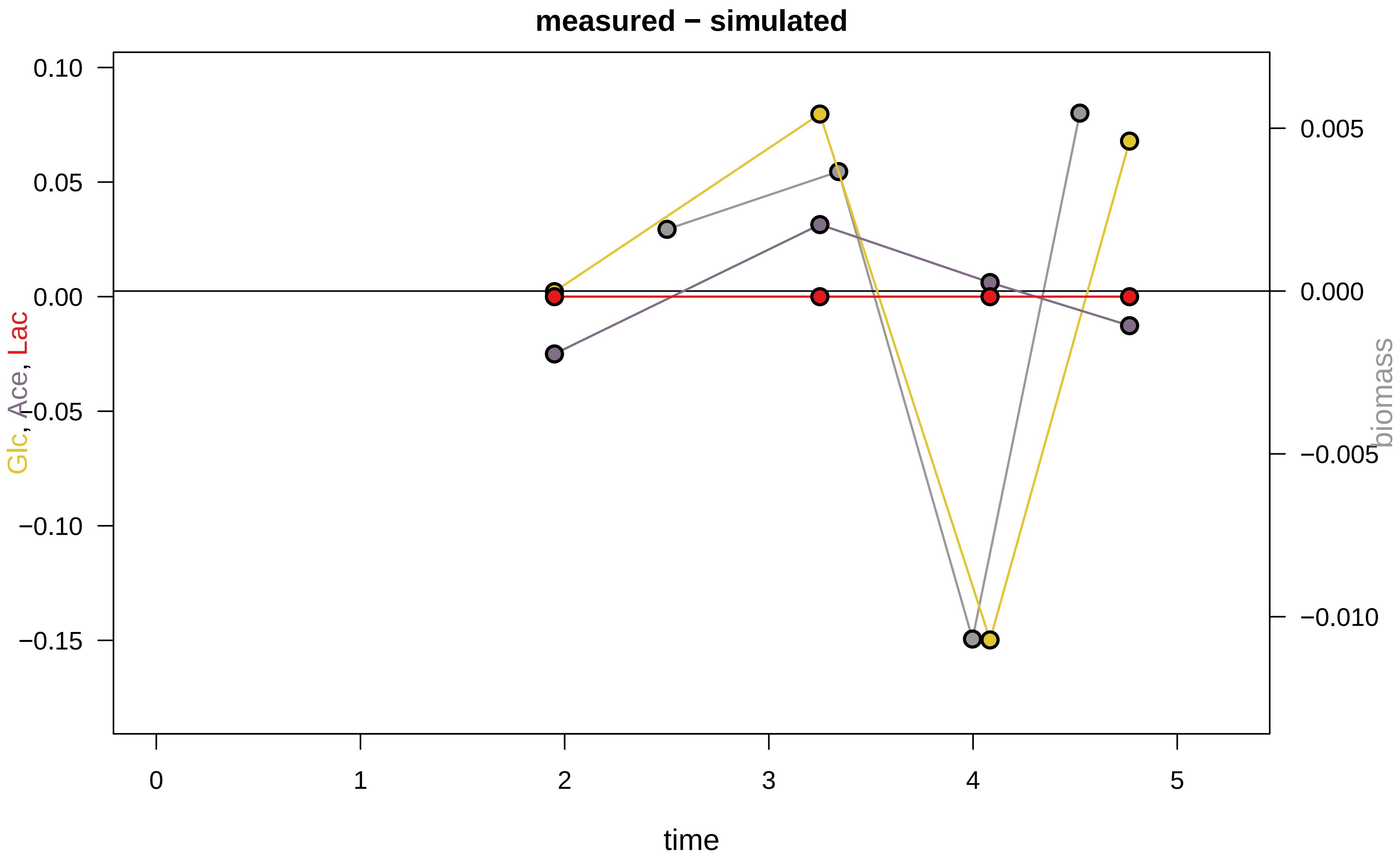

### KEIO_ROBOT2_42.pdf

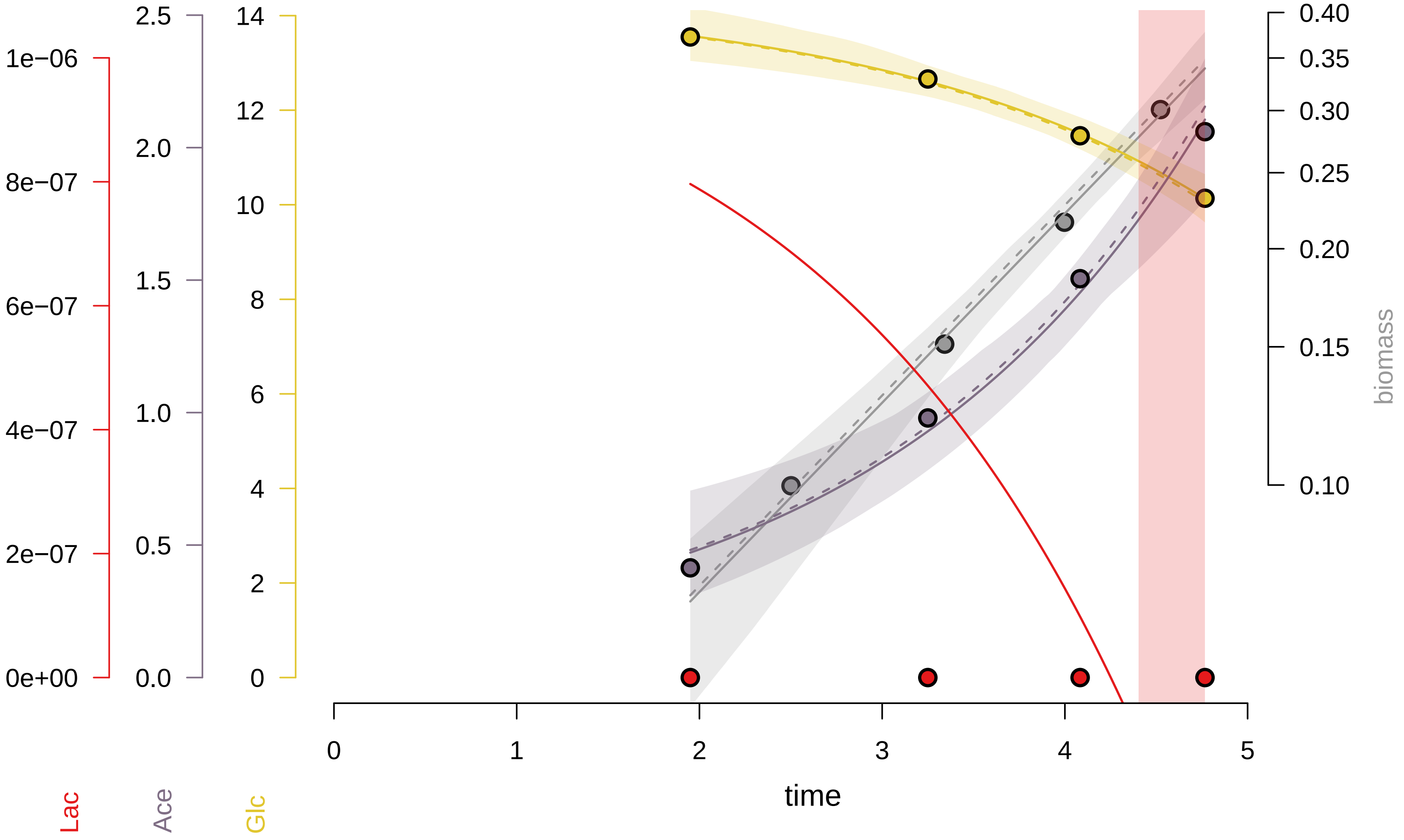

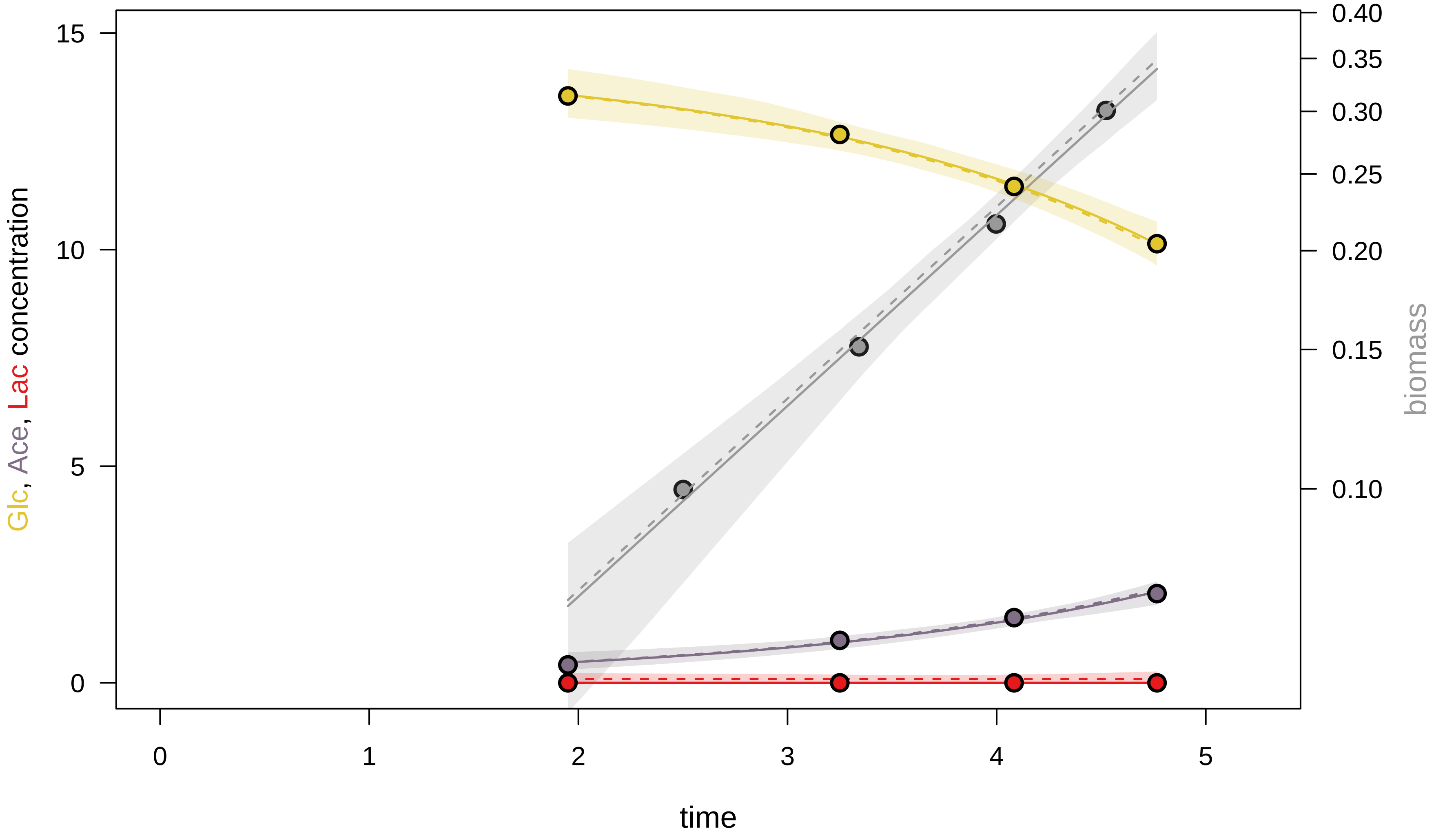

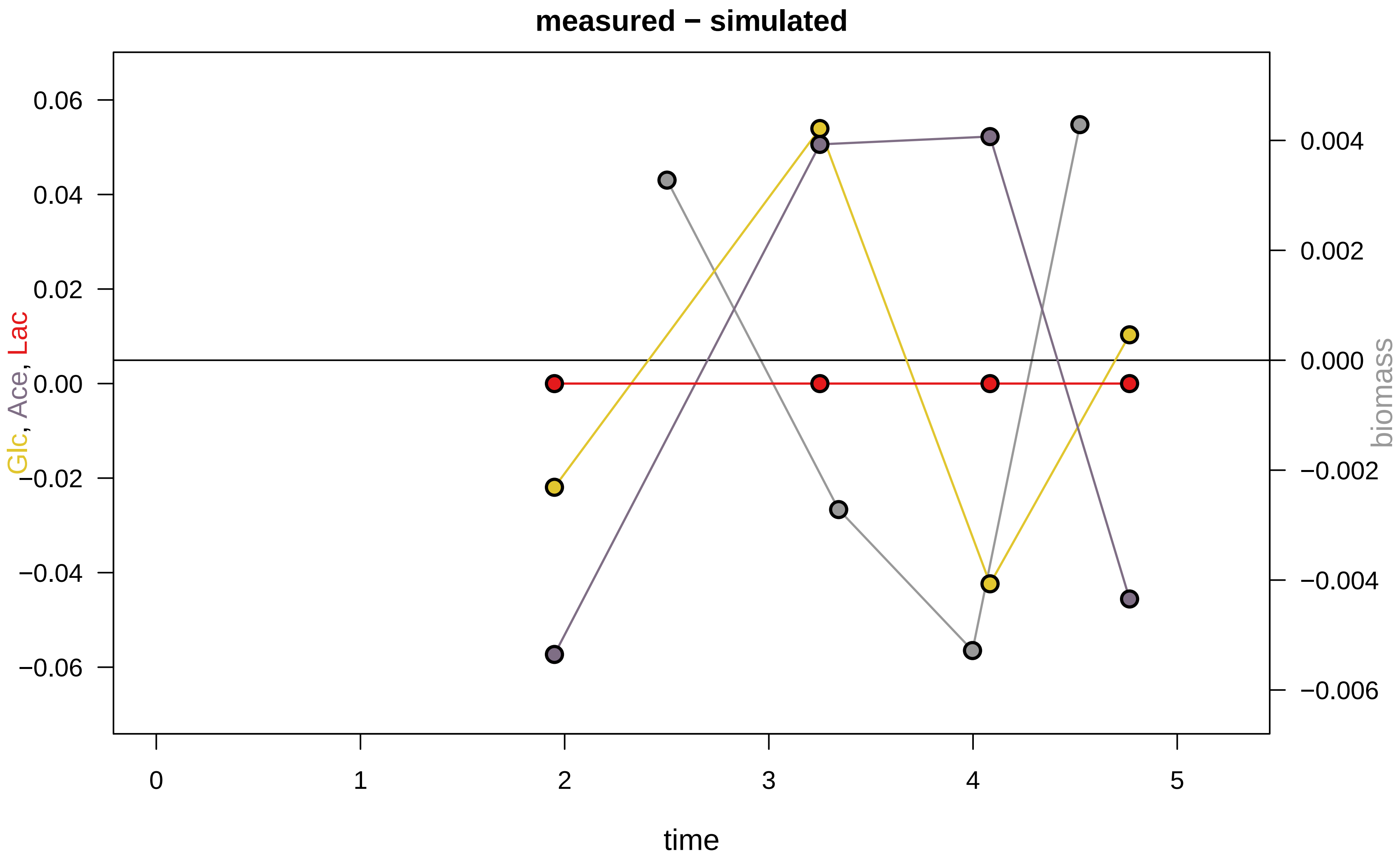

### KEIO_ROBOT2_46.pdf

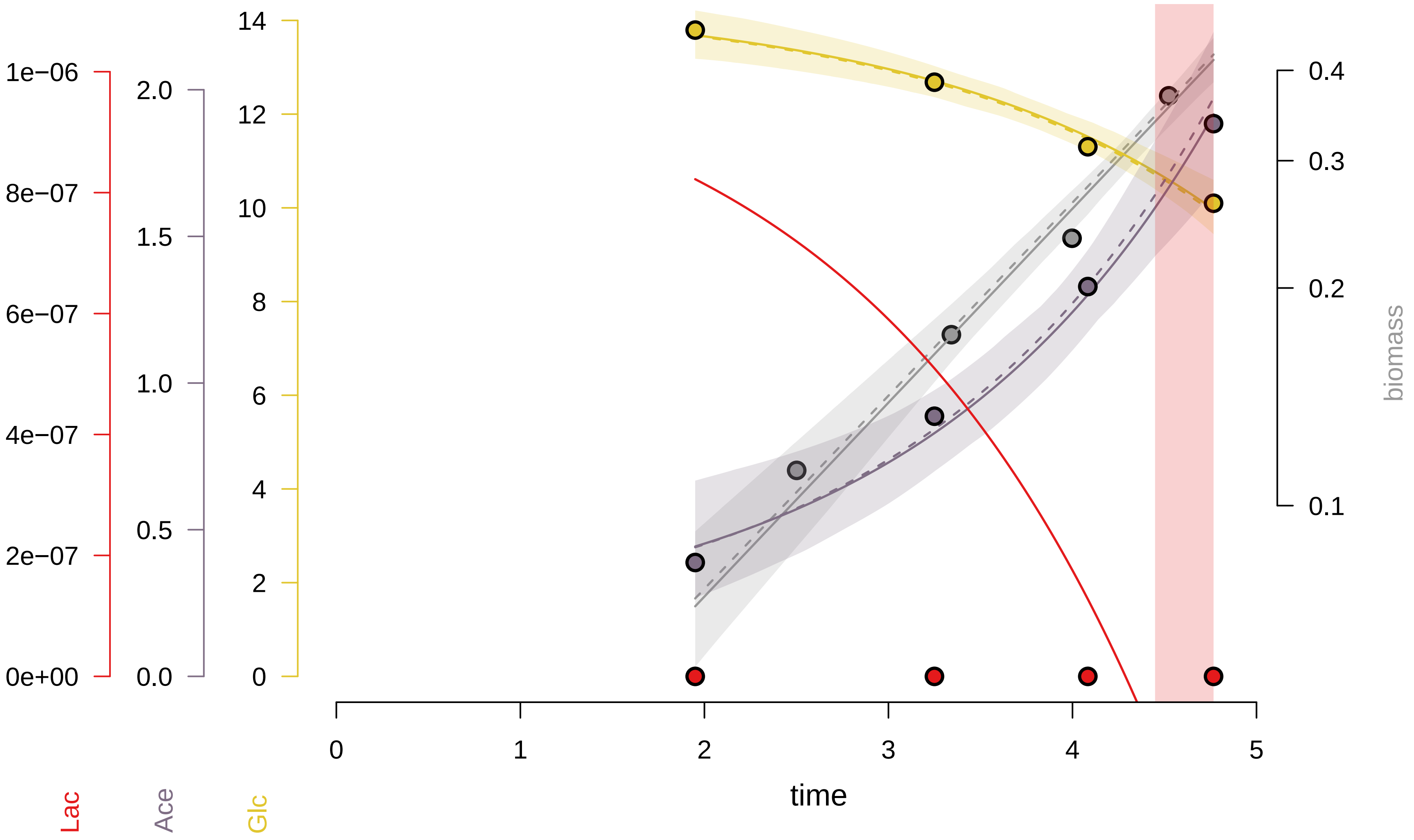

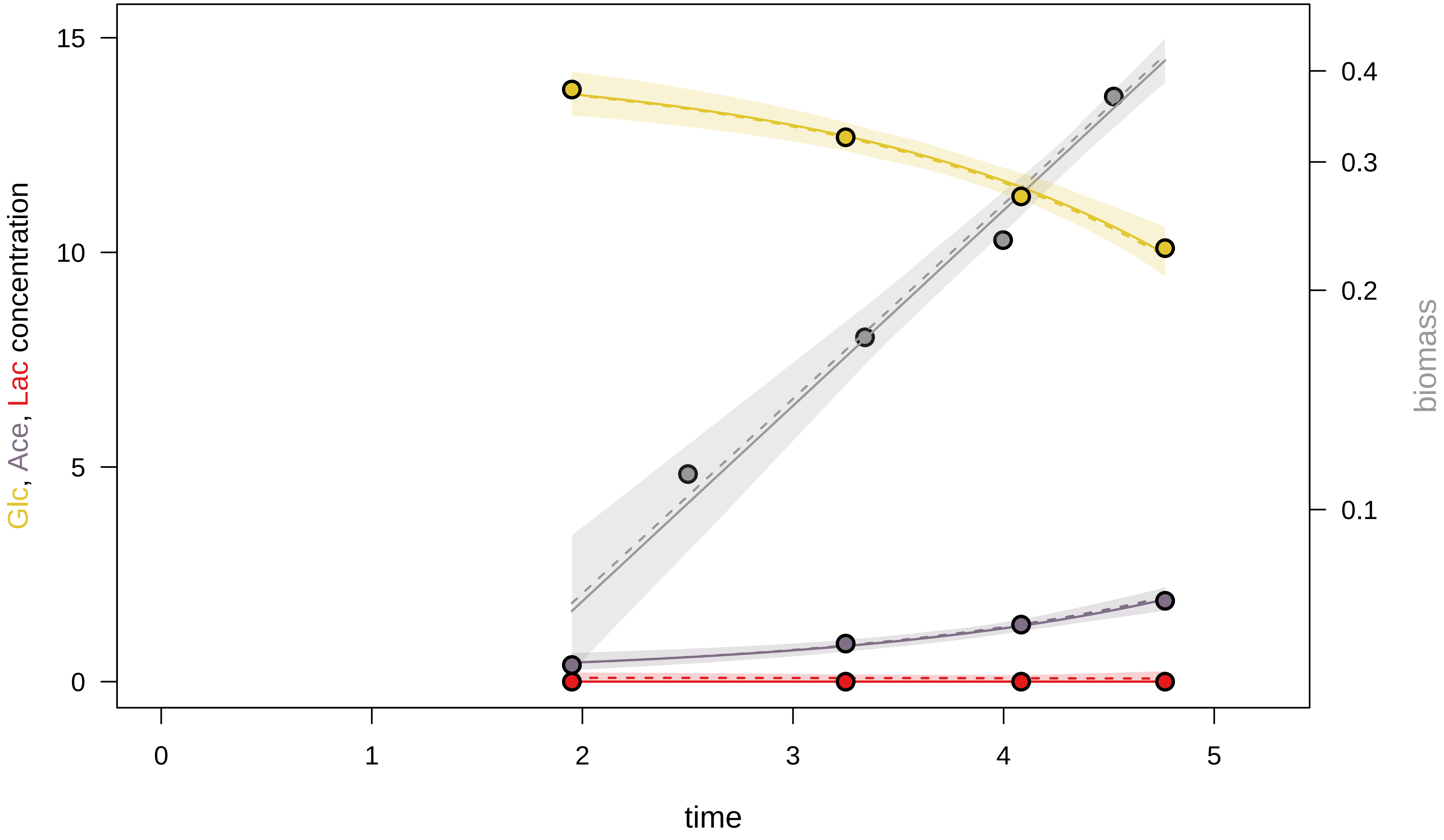

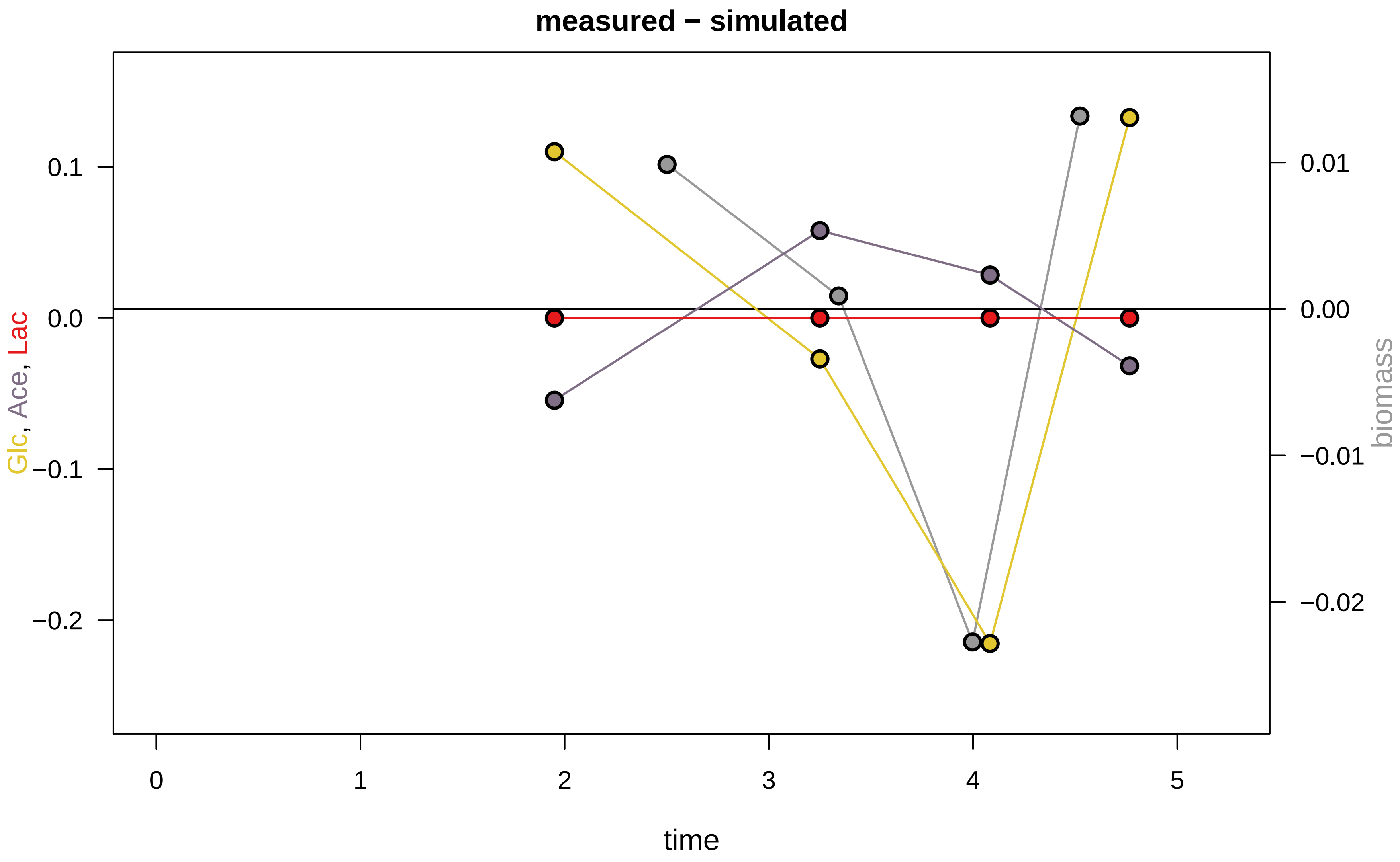

### KEIO_ROBOT3_15.pdf

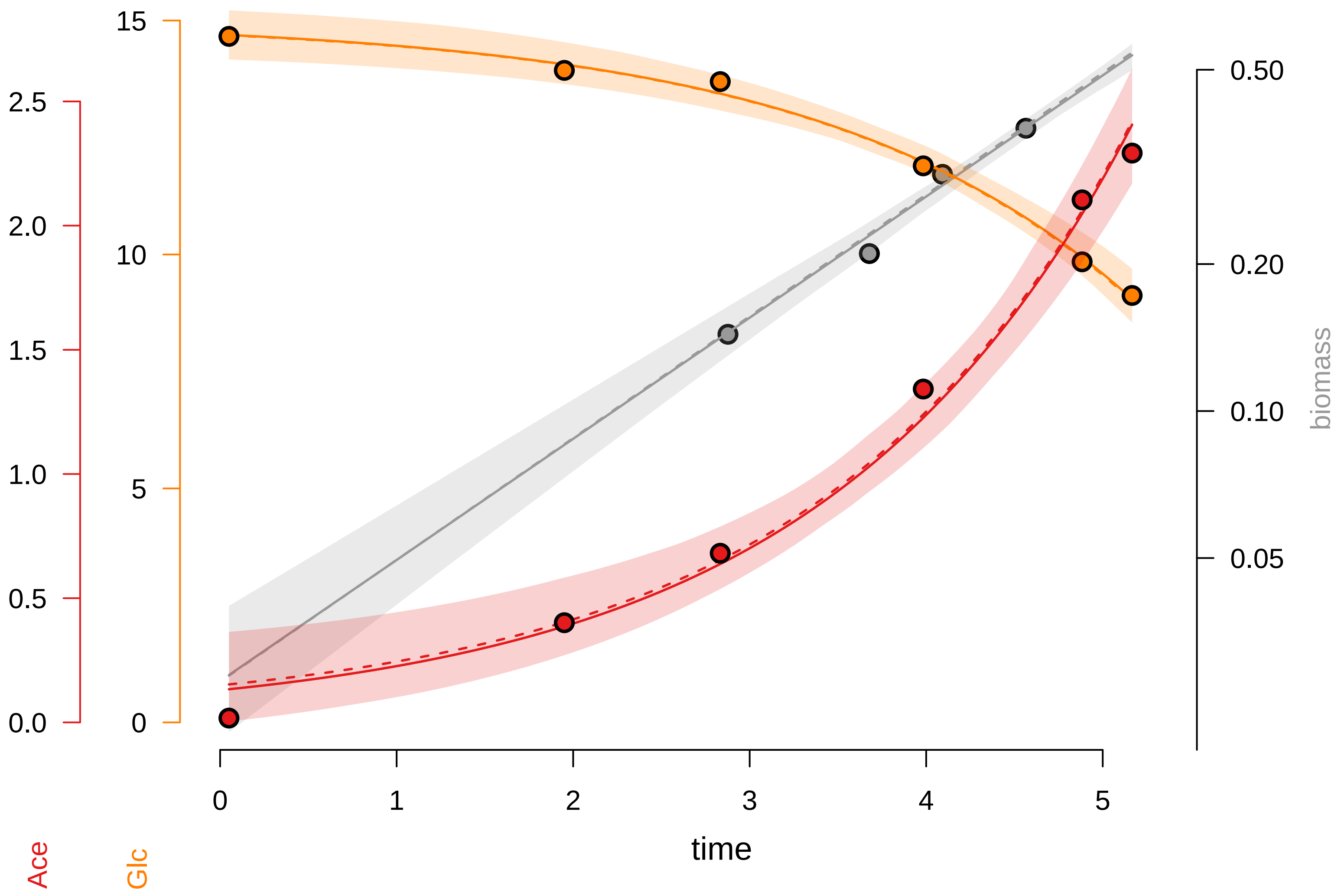

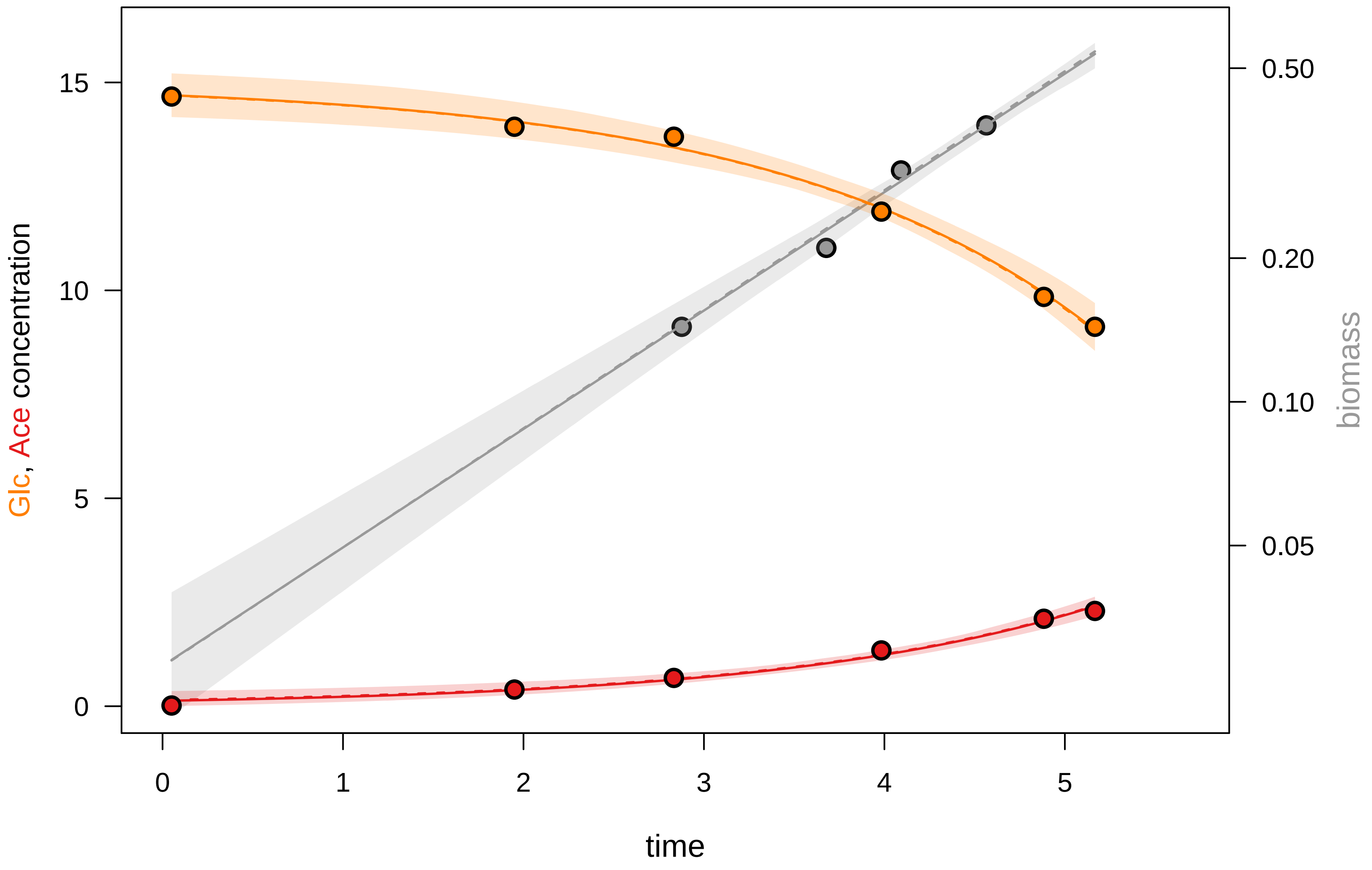

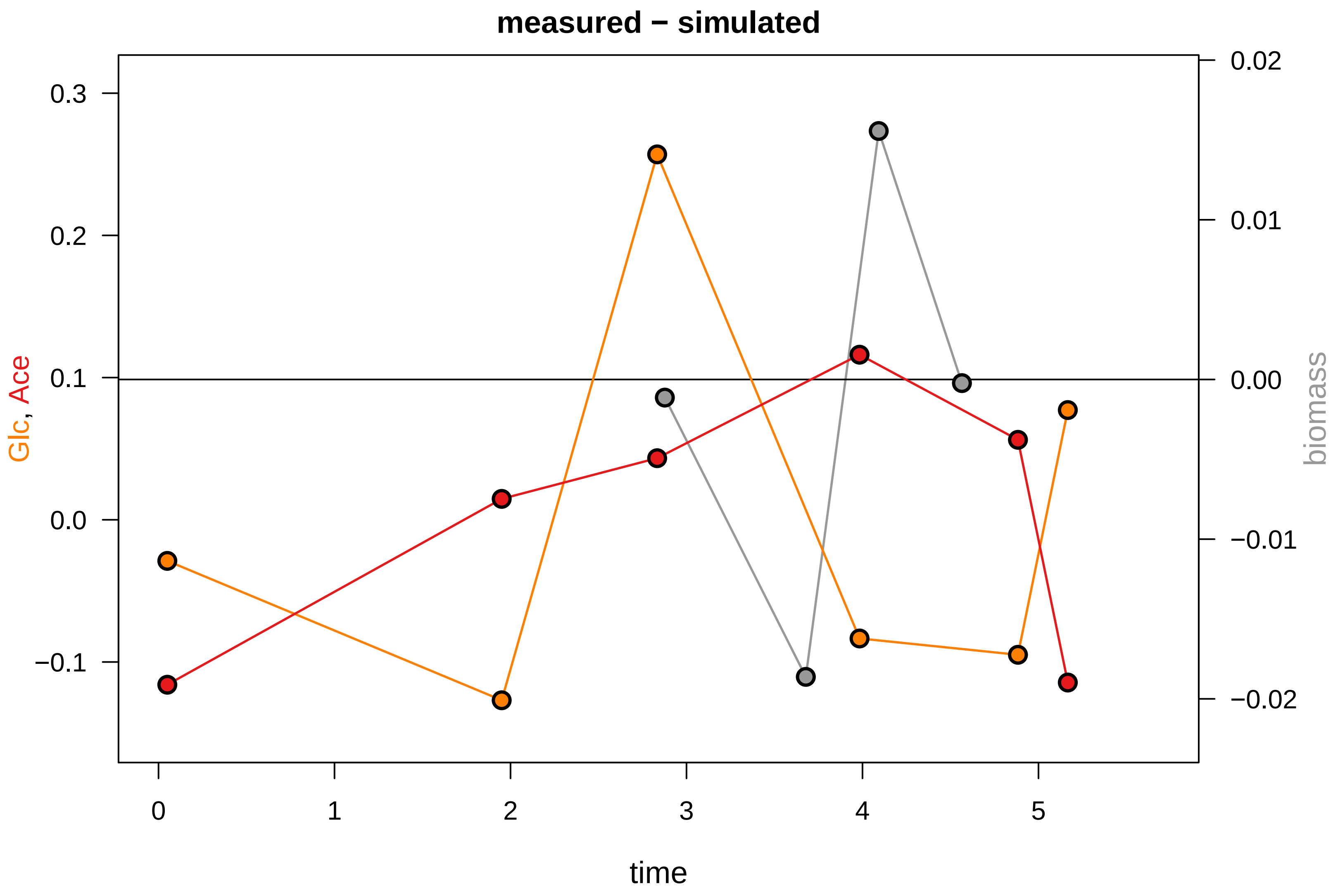

### KEIO_ROBOT3_17.pdf

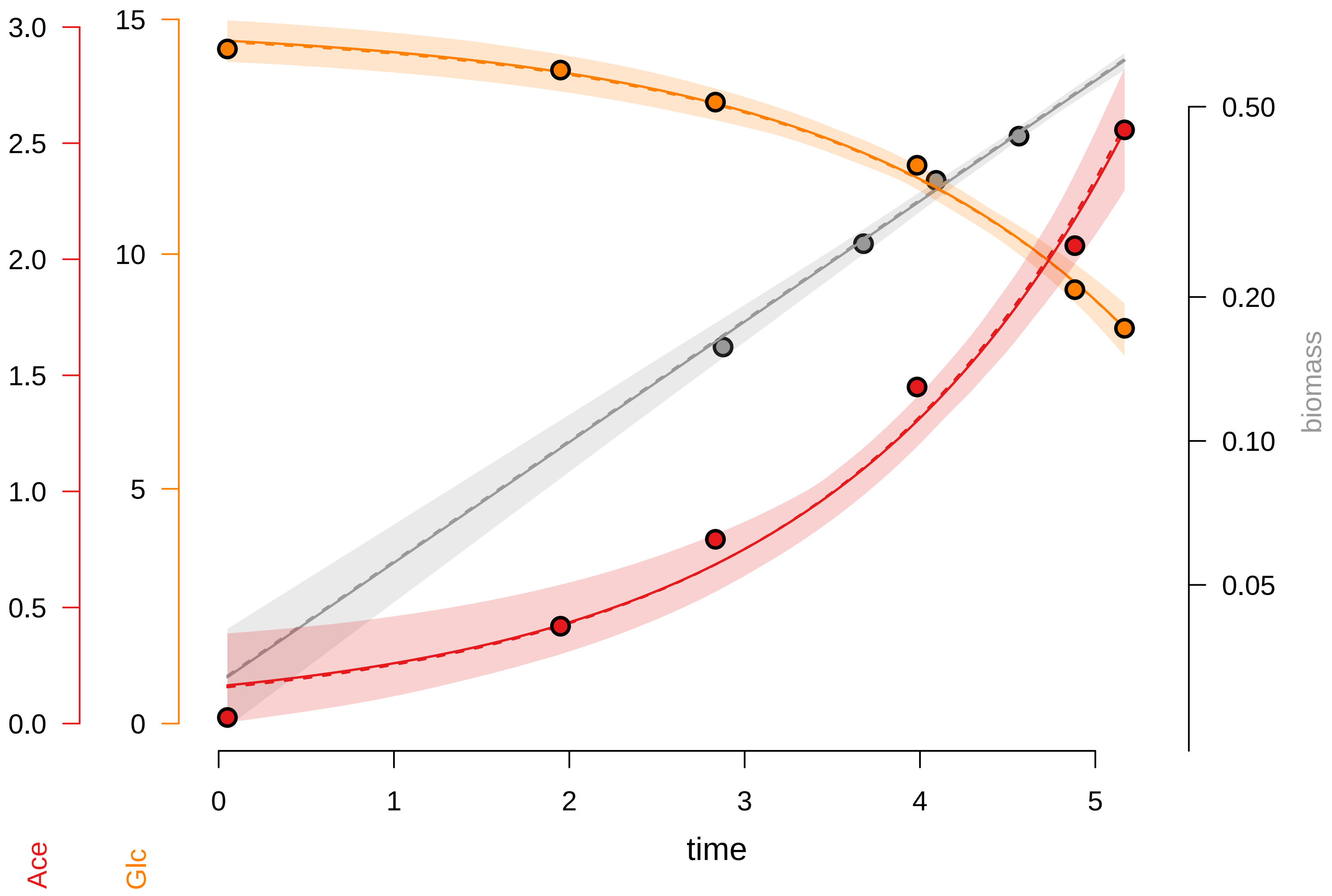

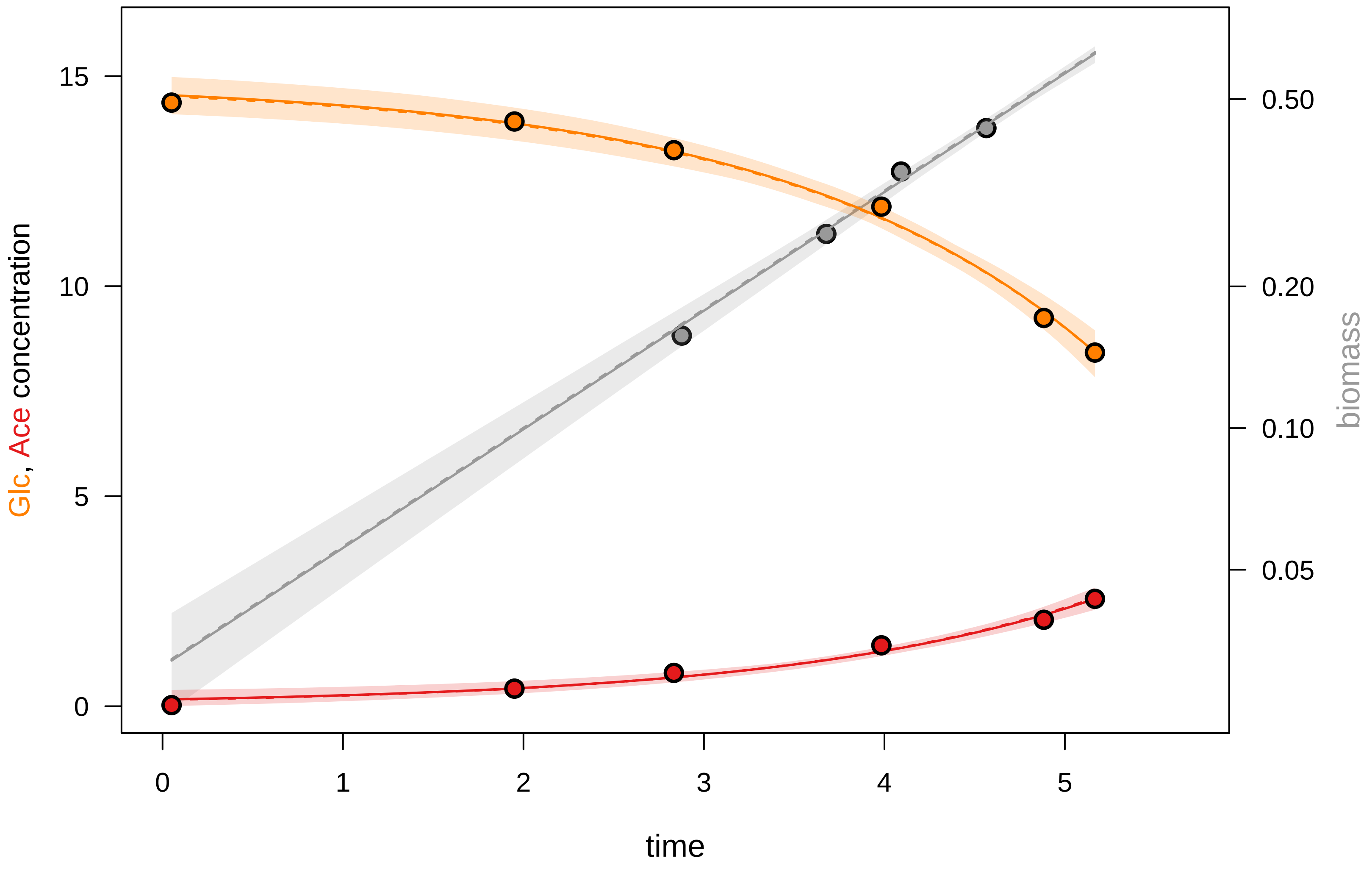

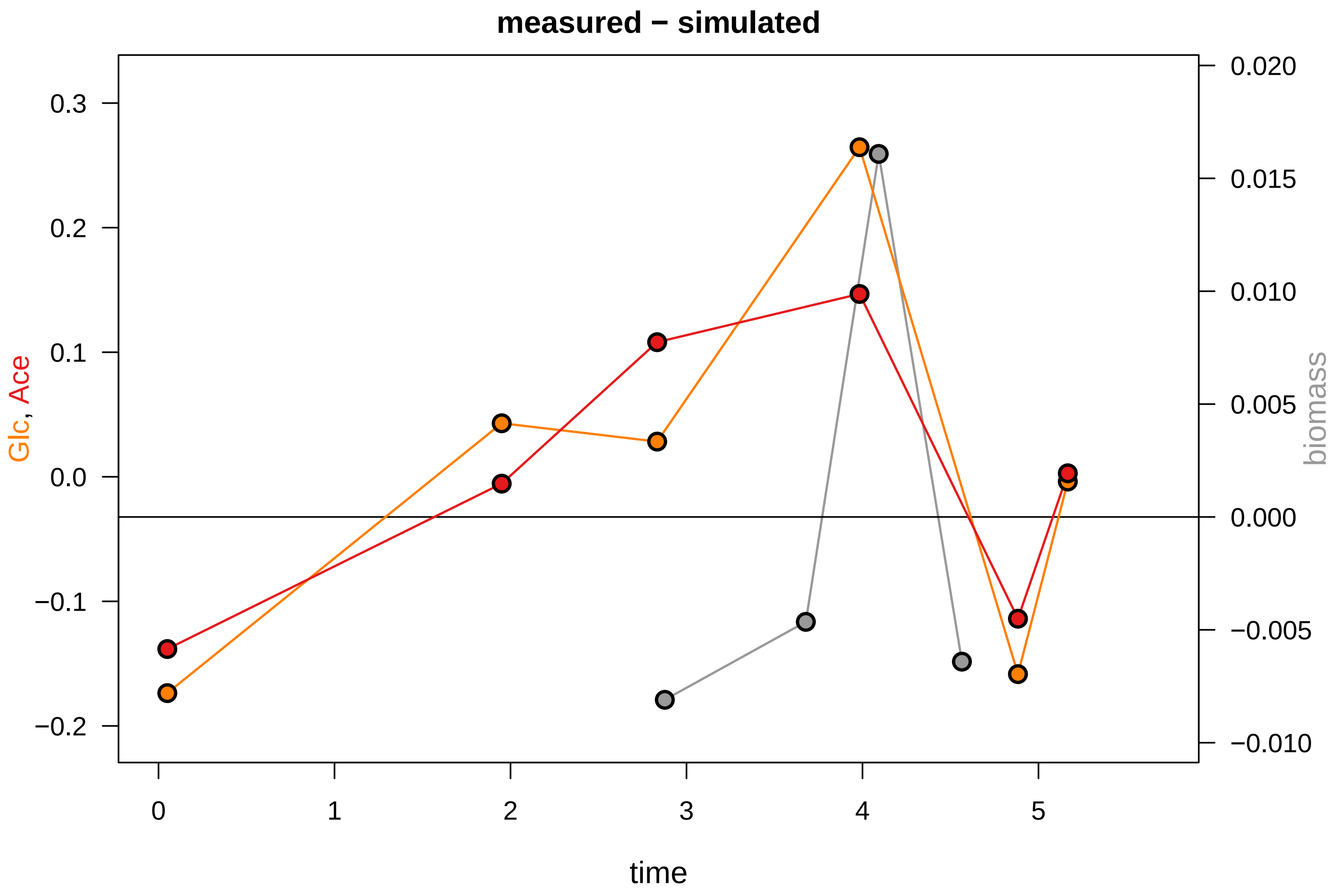

### KEIO_ROBOT3_18.pdf

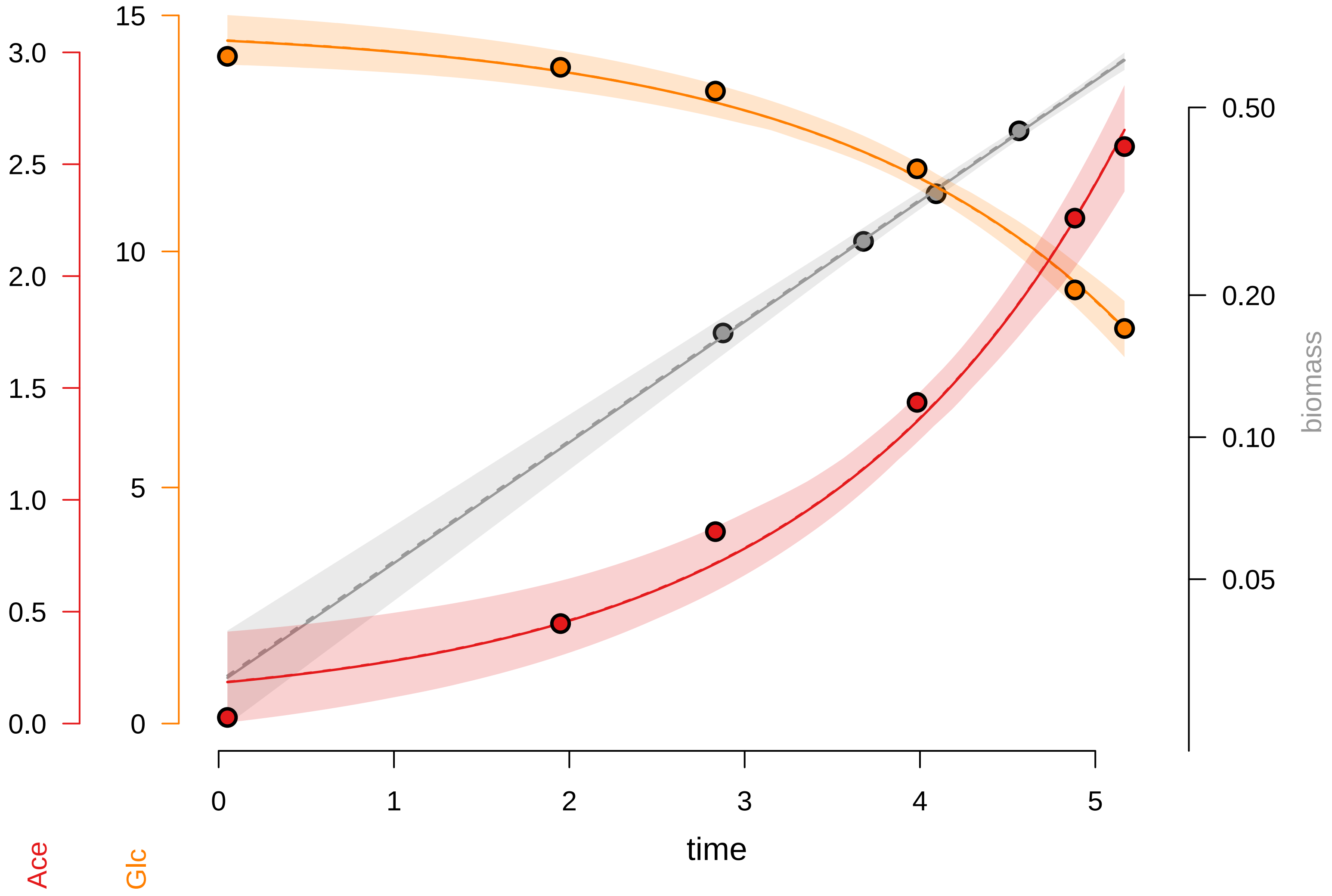

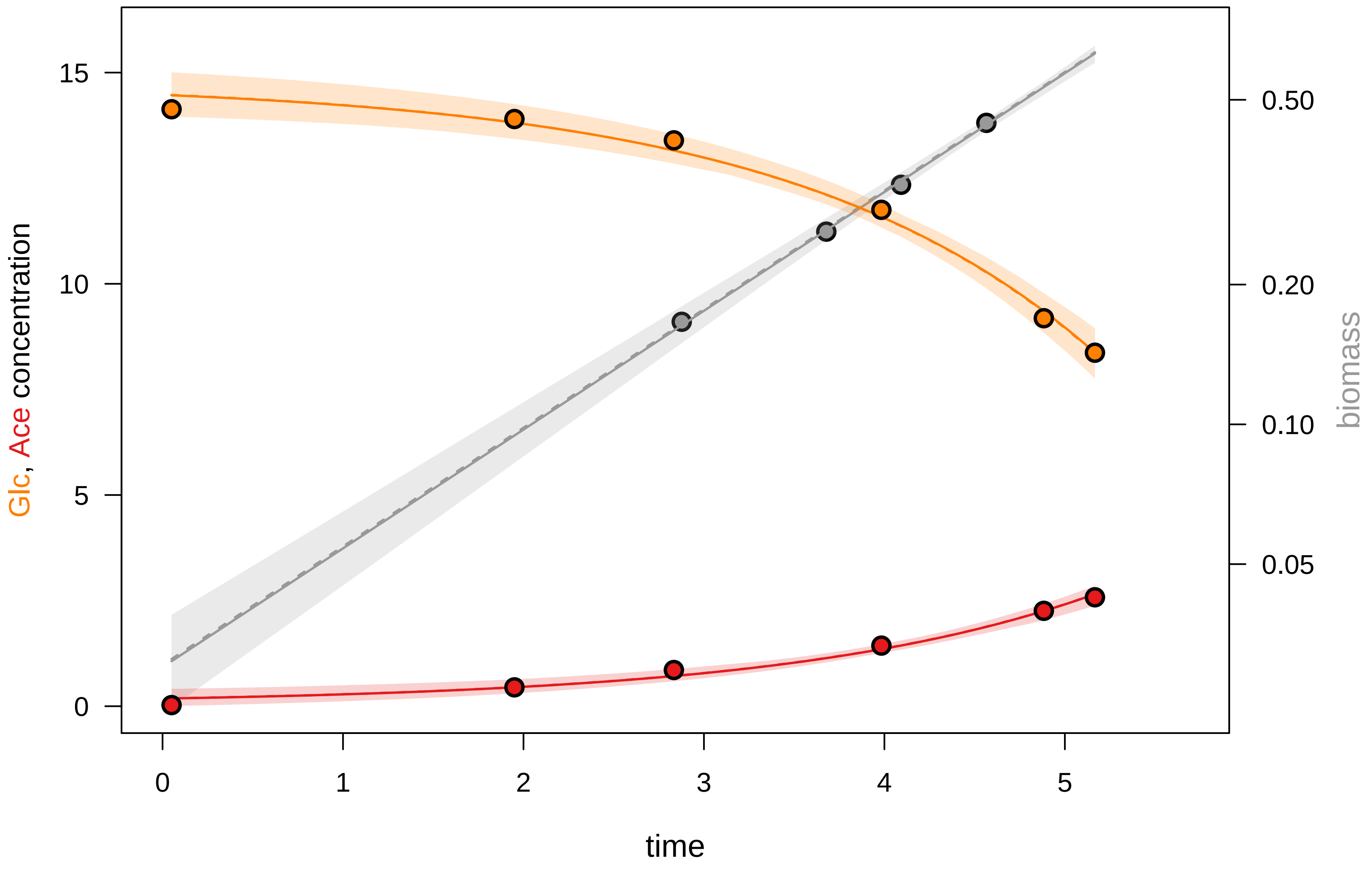

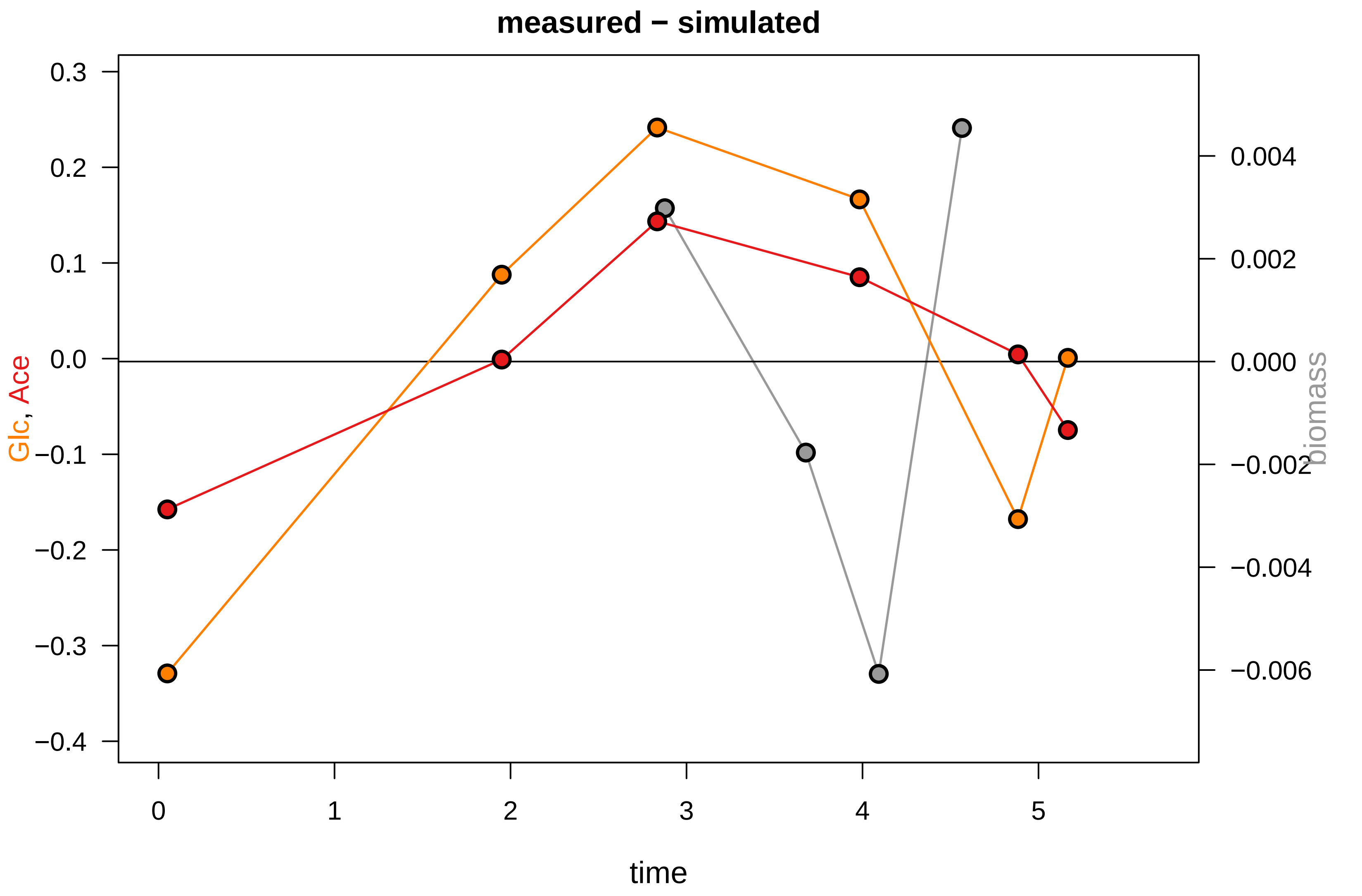

### KEIO_ROBOT3_19.pdf

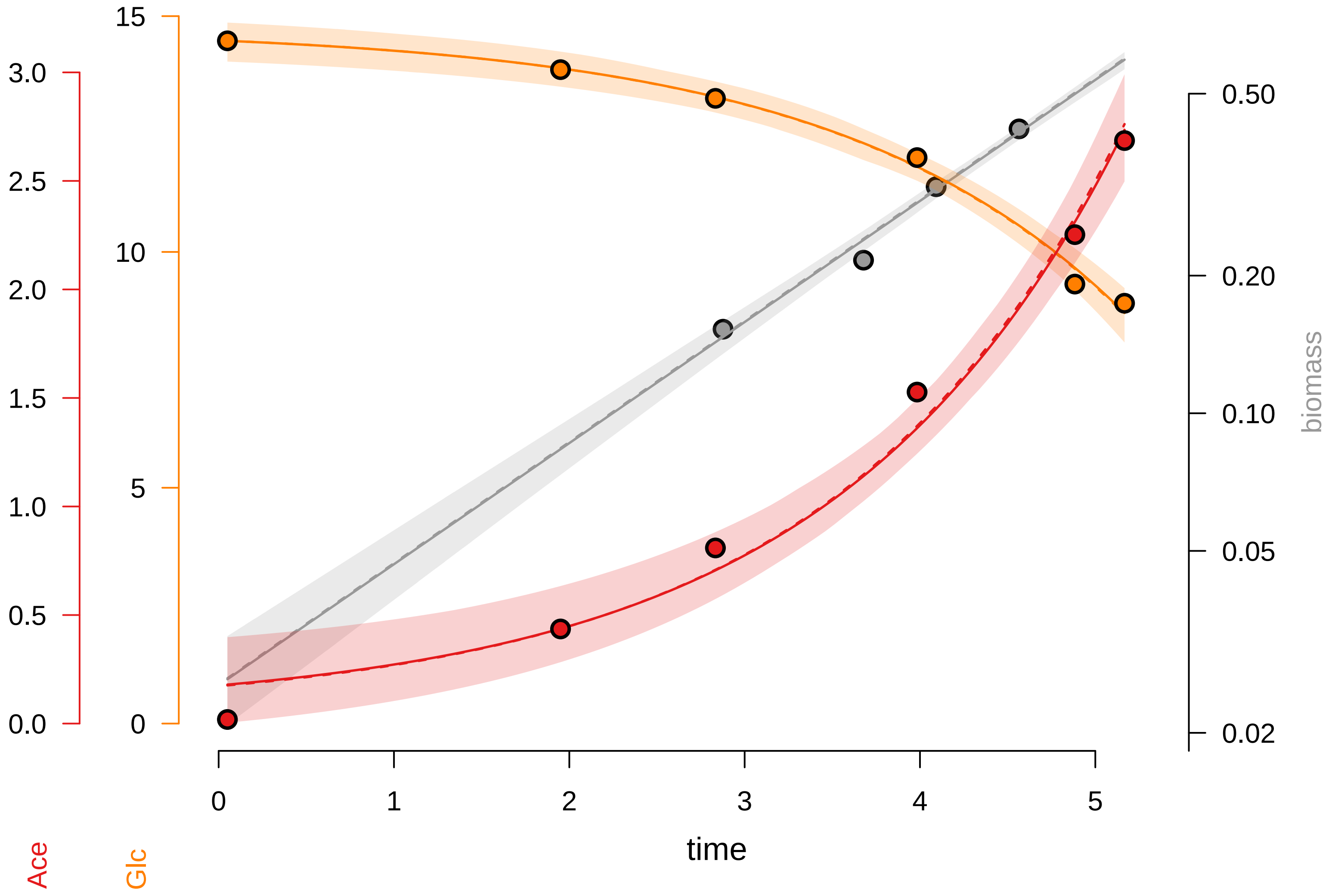

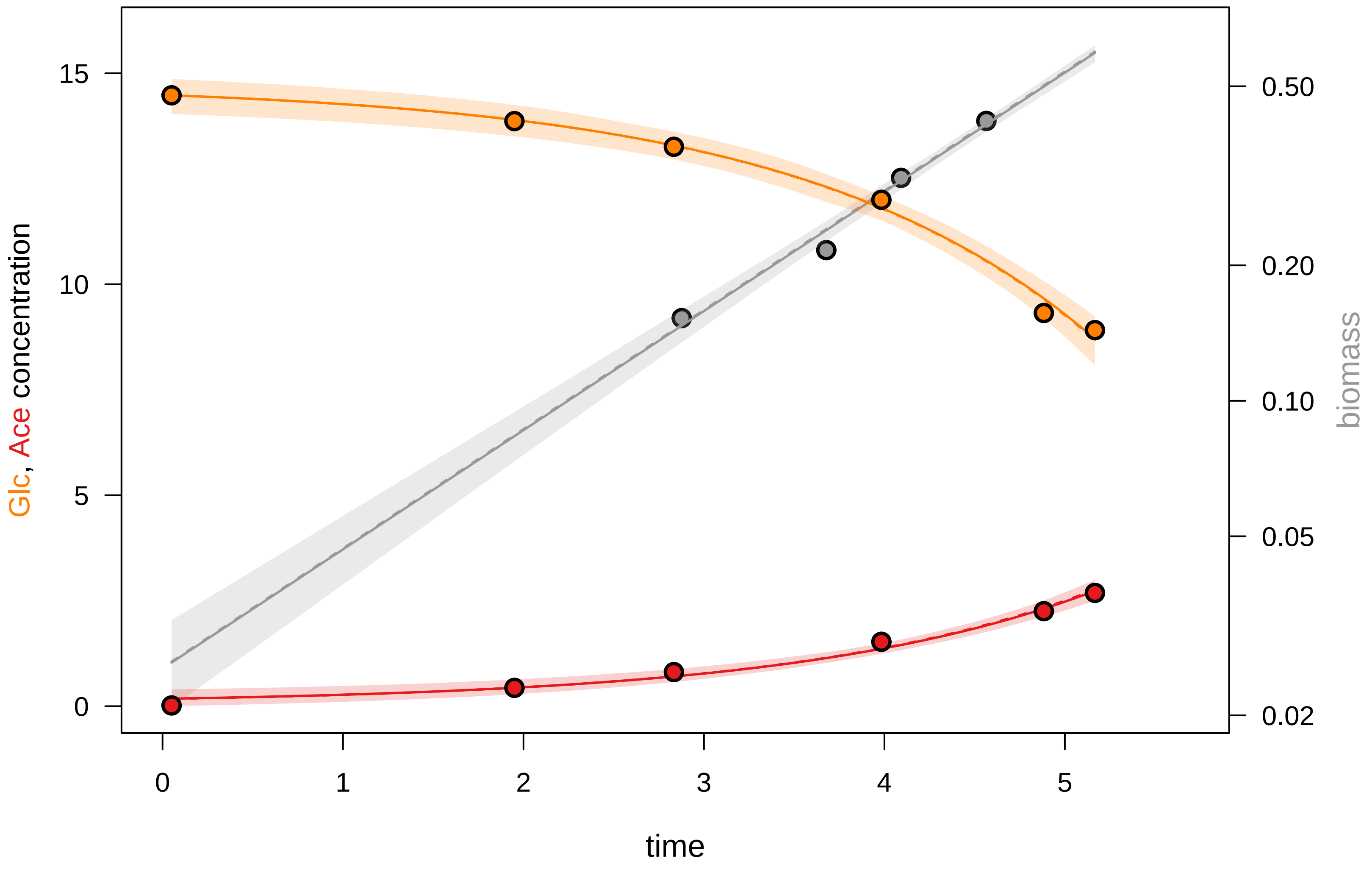

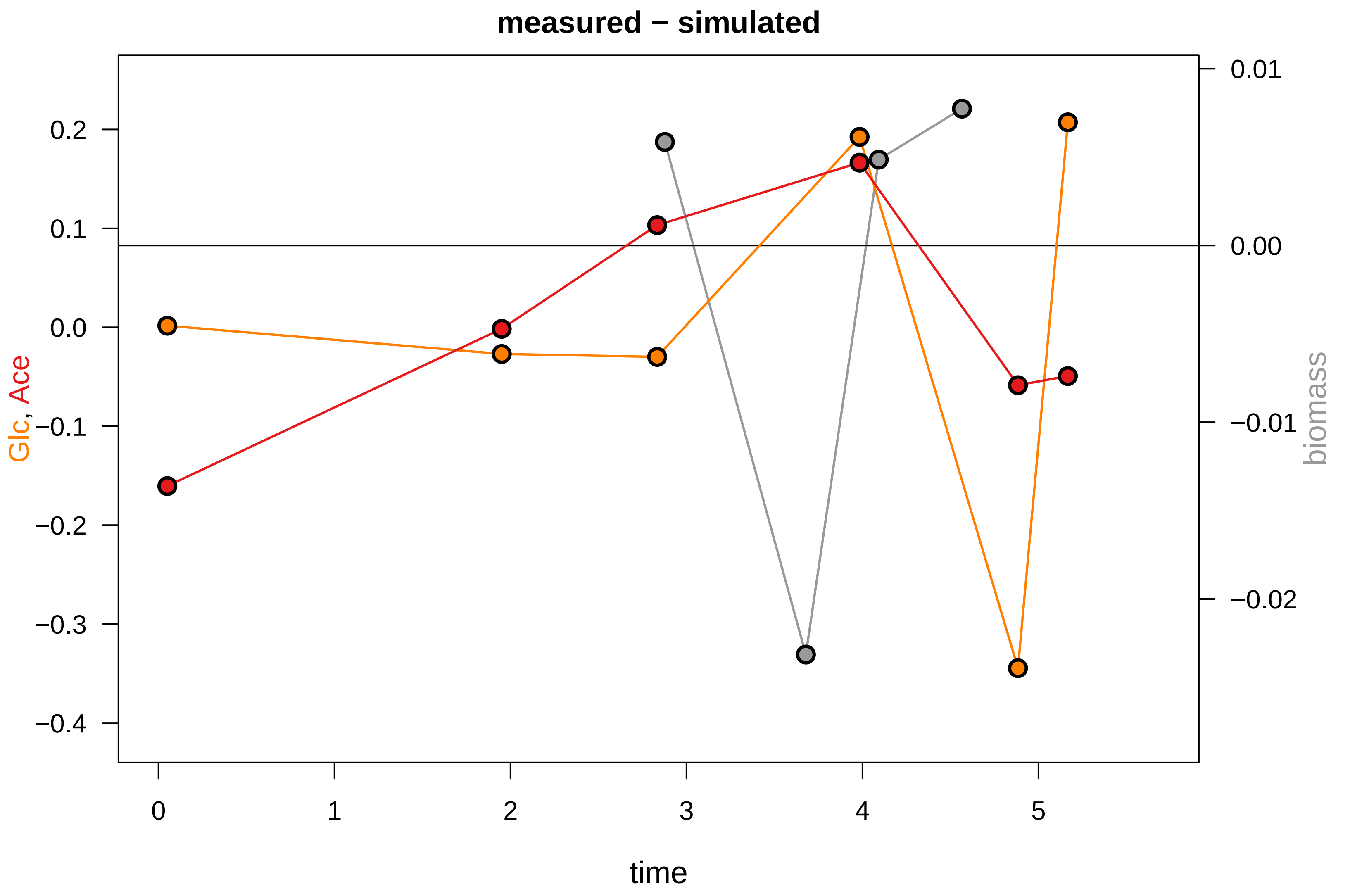

### KEIO_ROBOT3_24.pdf

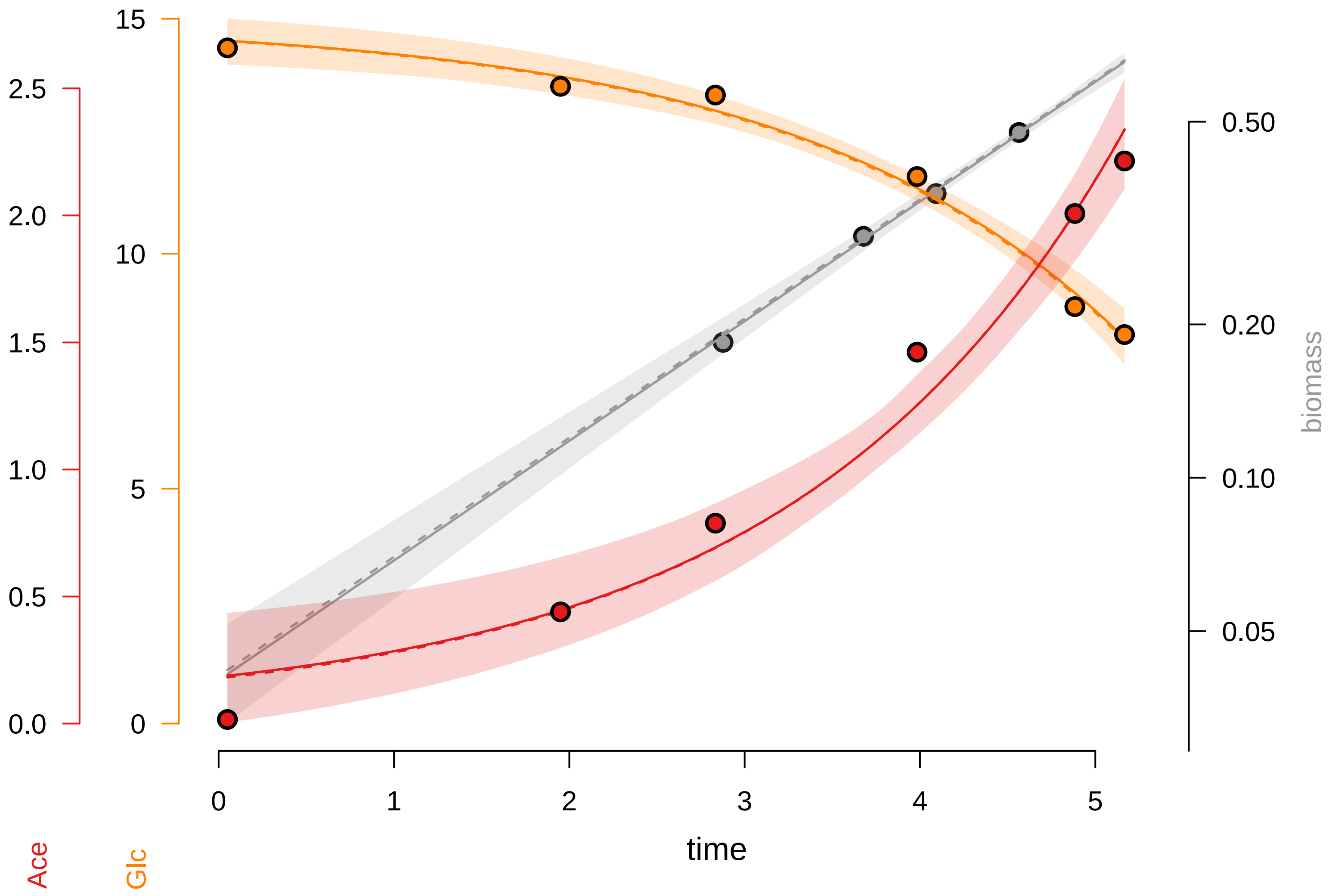

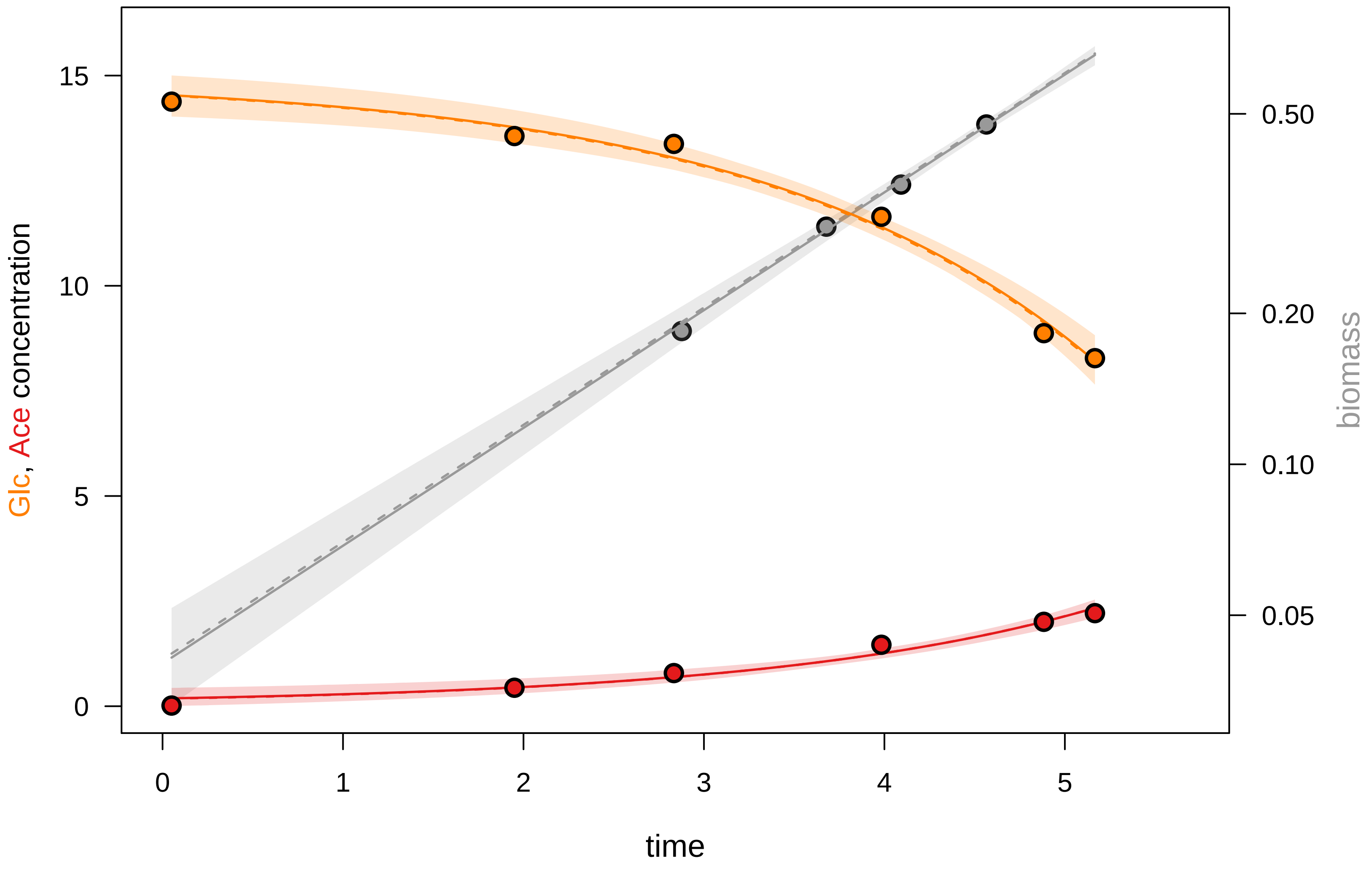

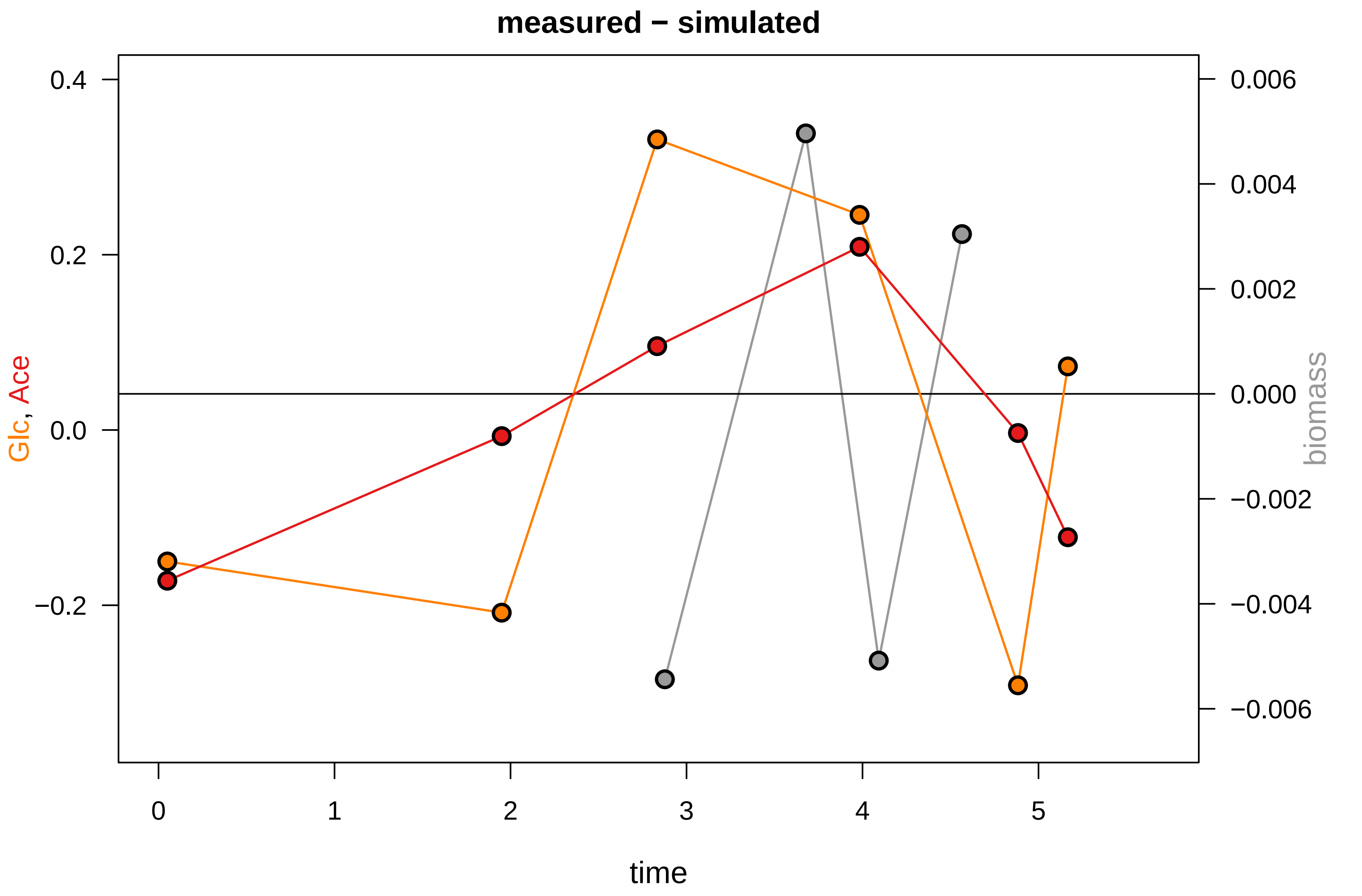

### KEIO_ROBOT3_25.pdf

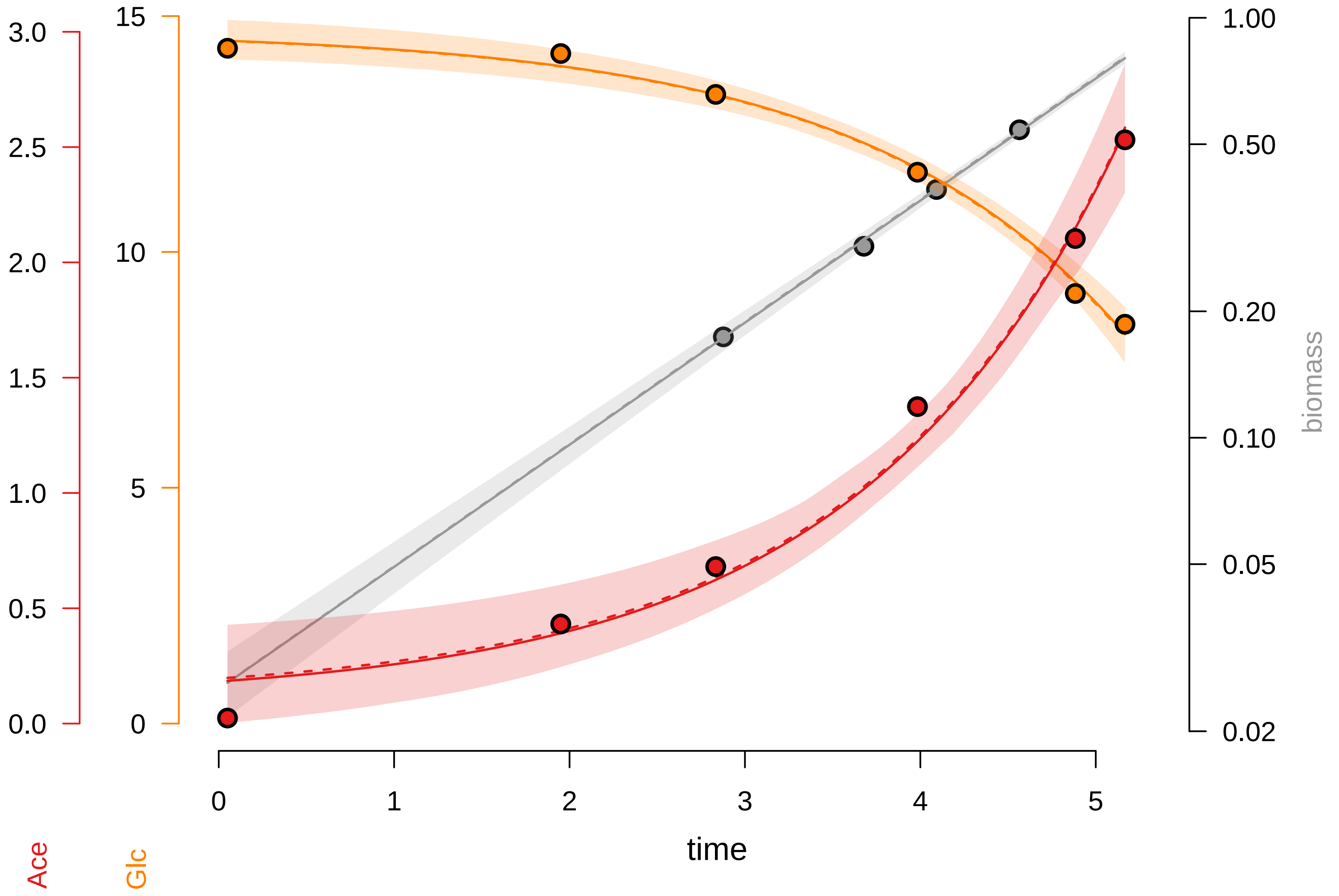

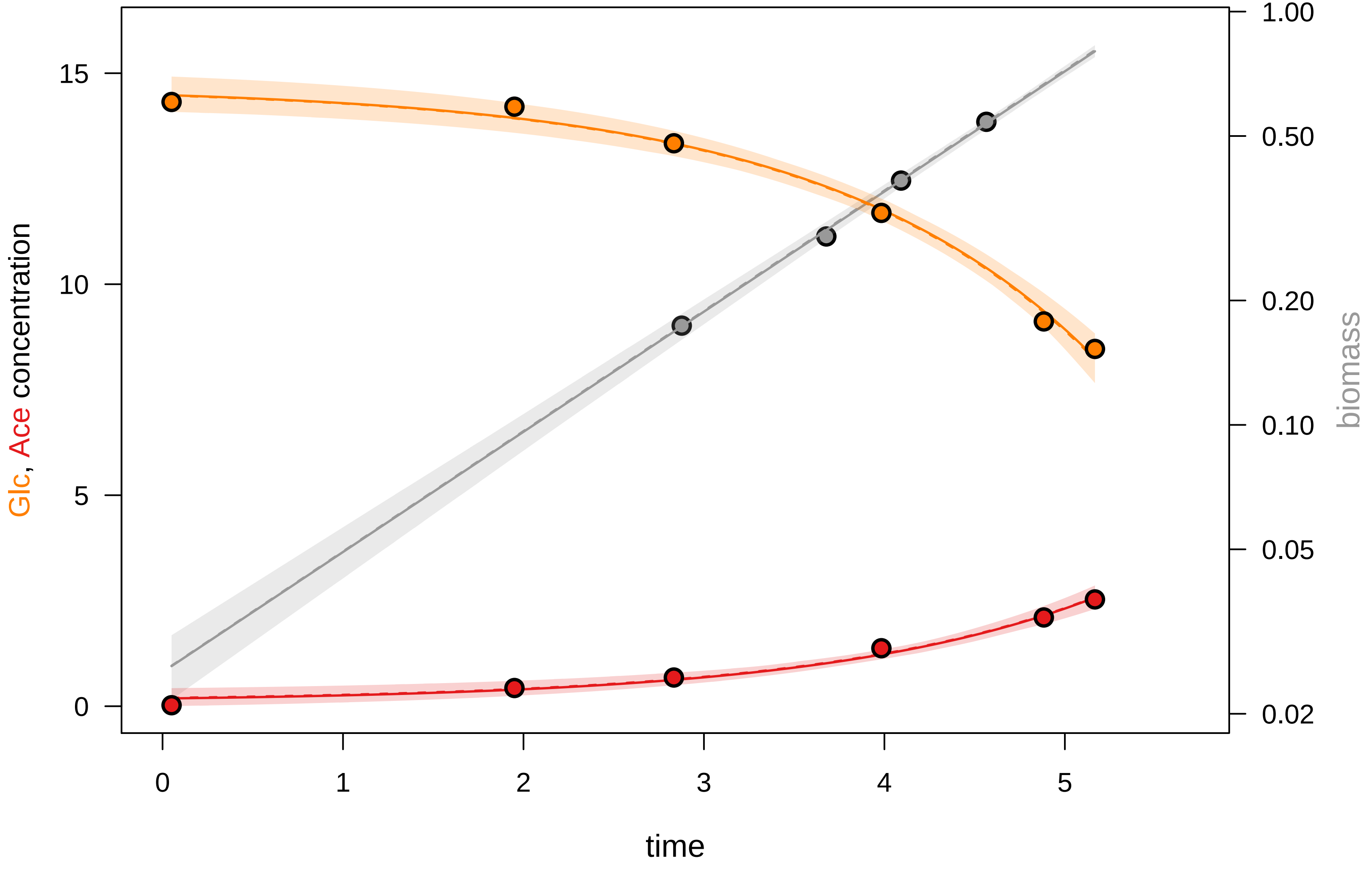

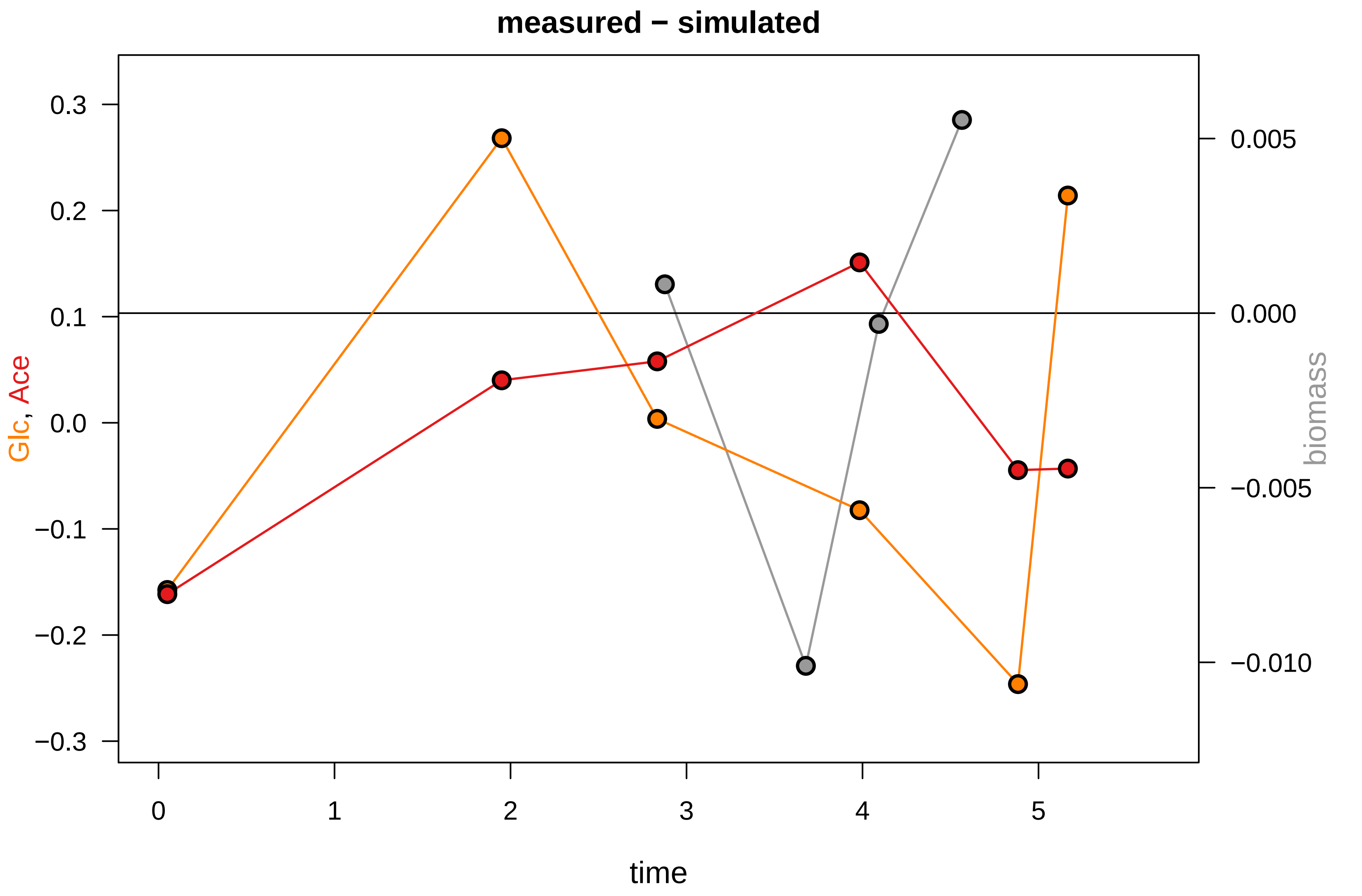

### KEIO_ROBOT3_27.pdf

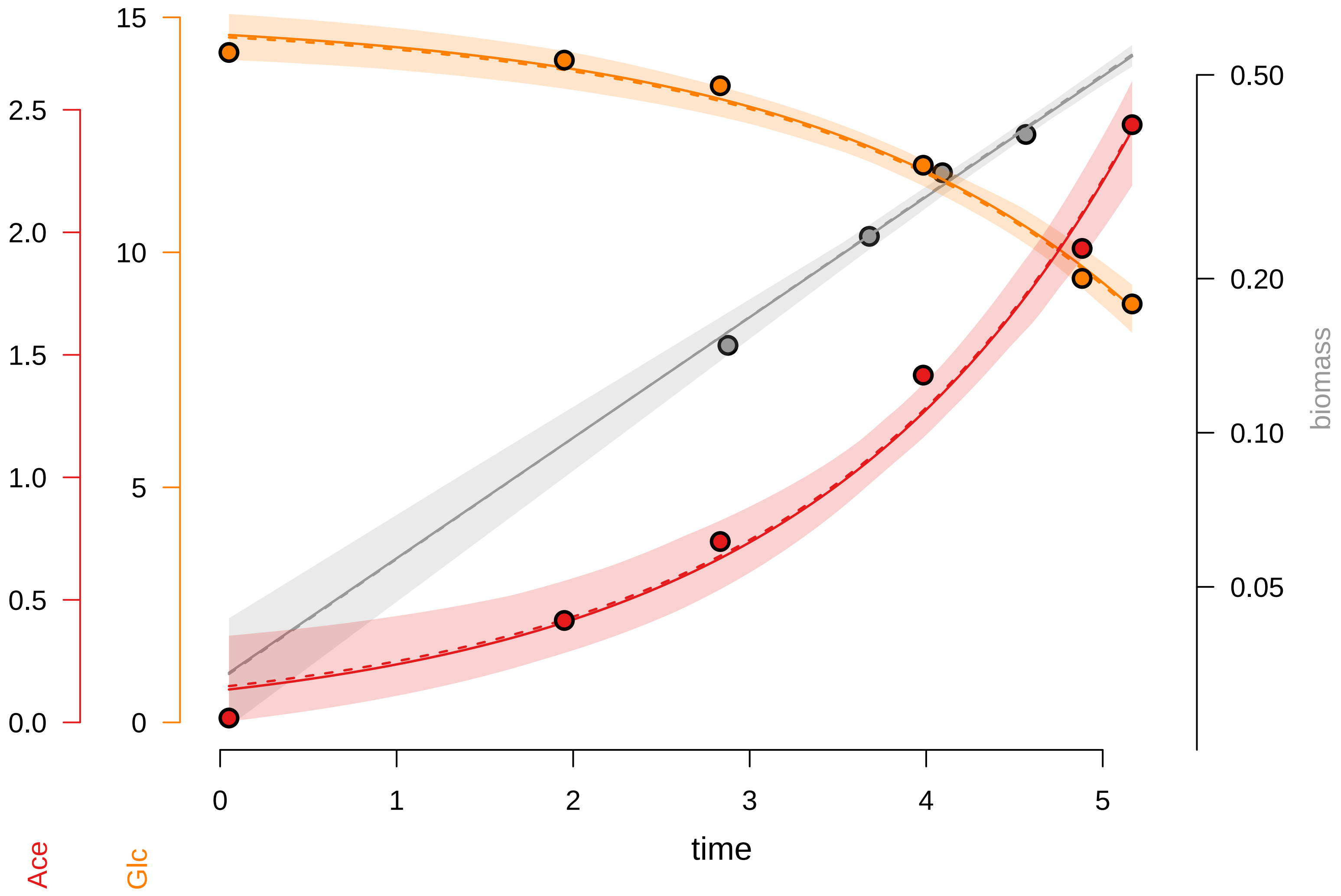

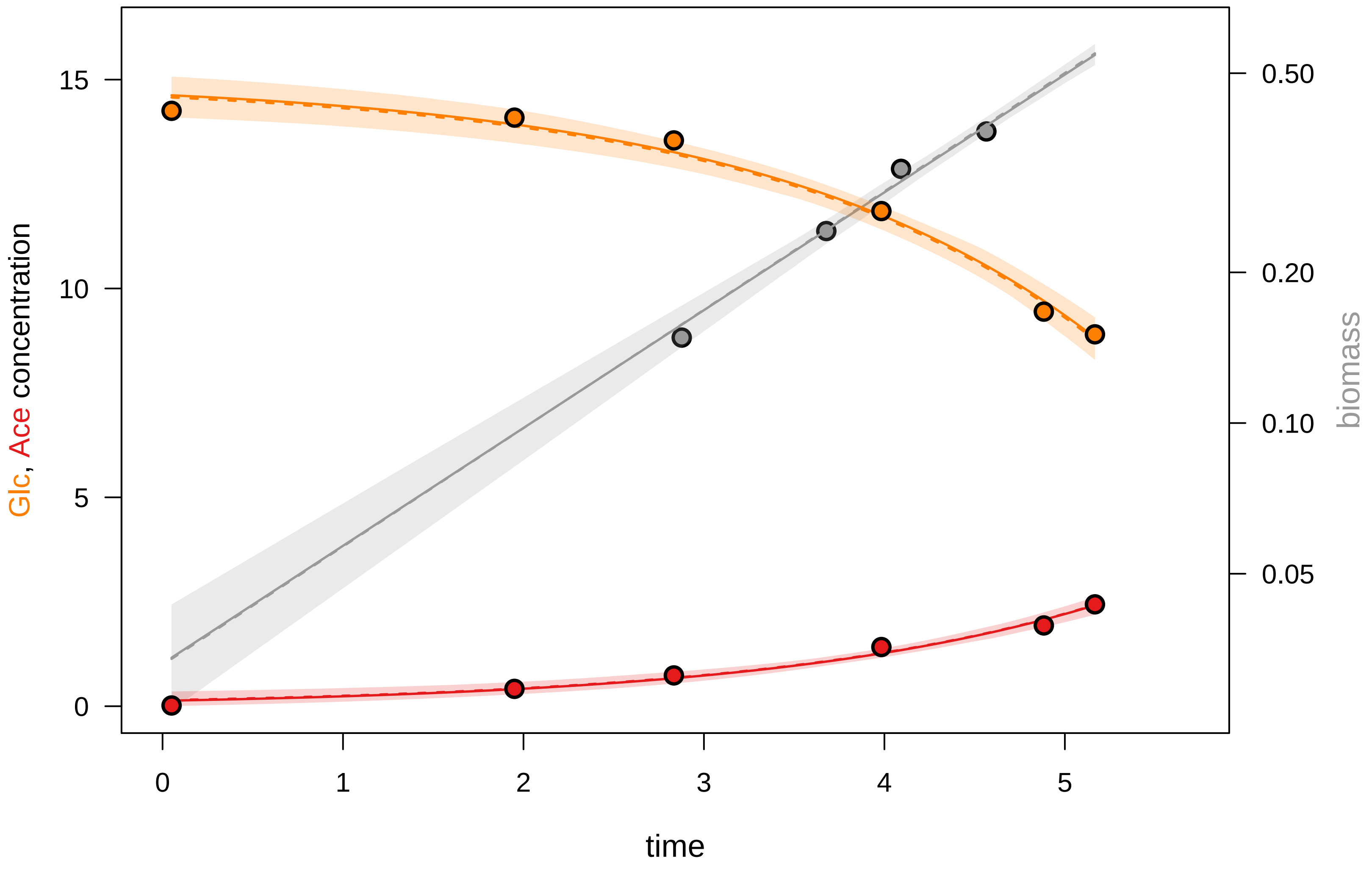

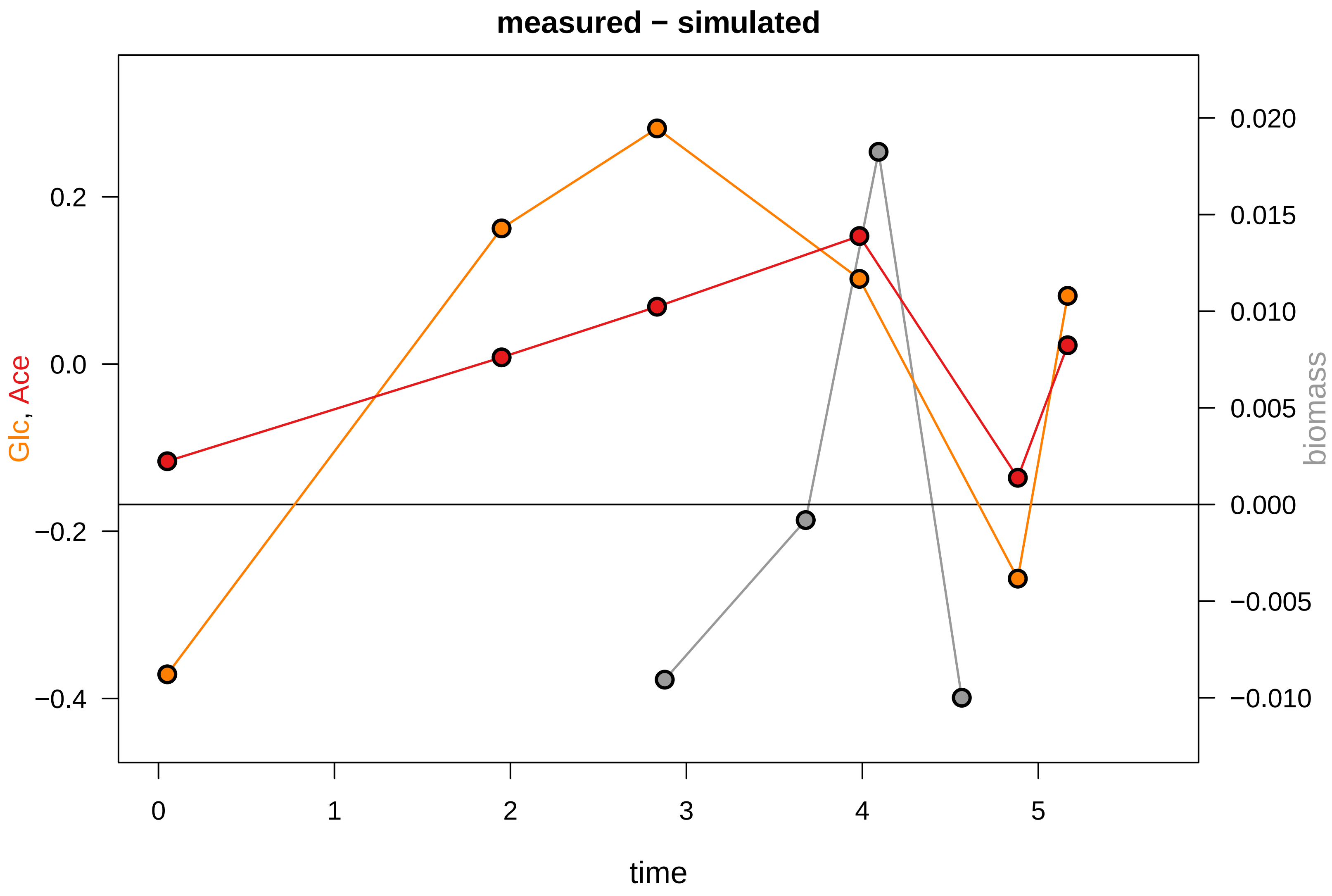

### KEIO_ROBOT3_28.pdf

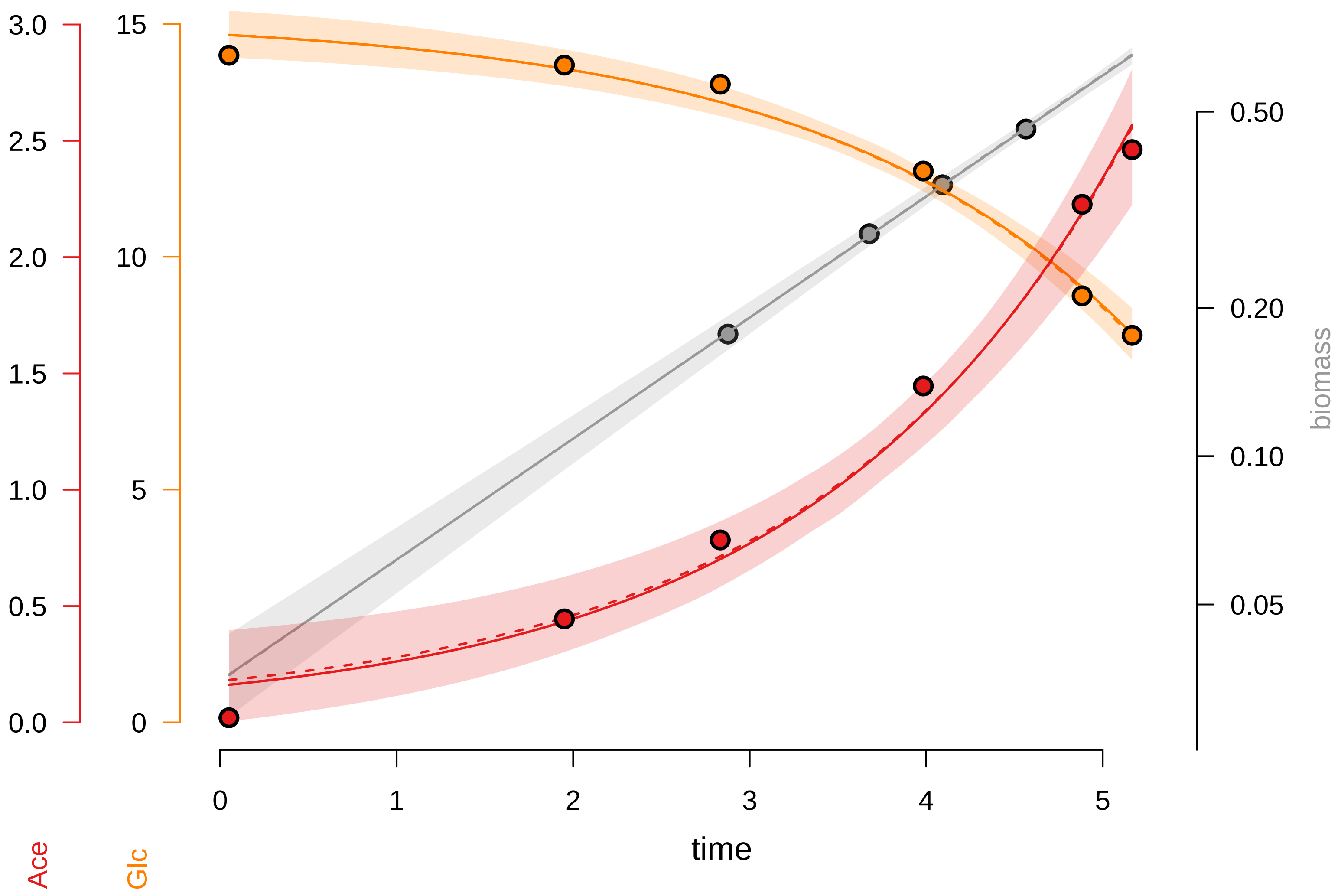

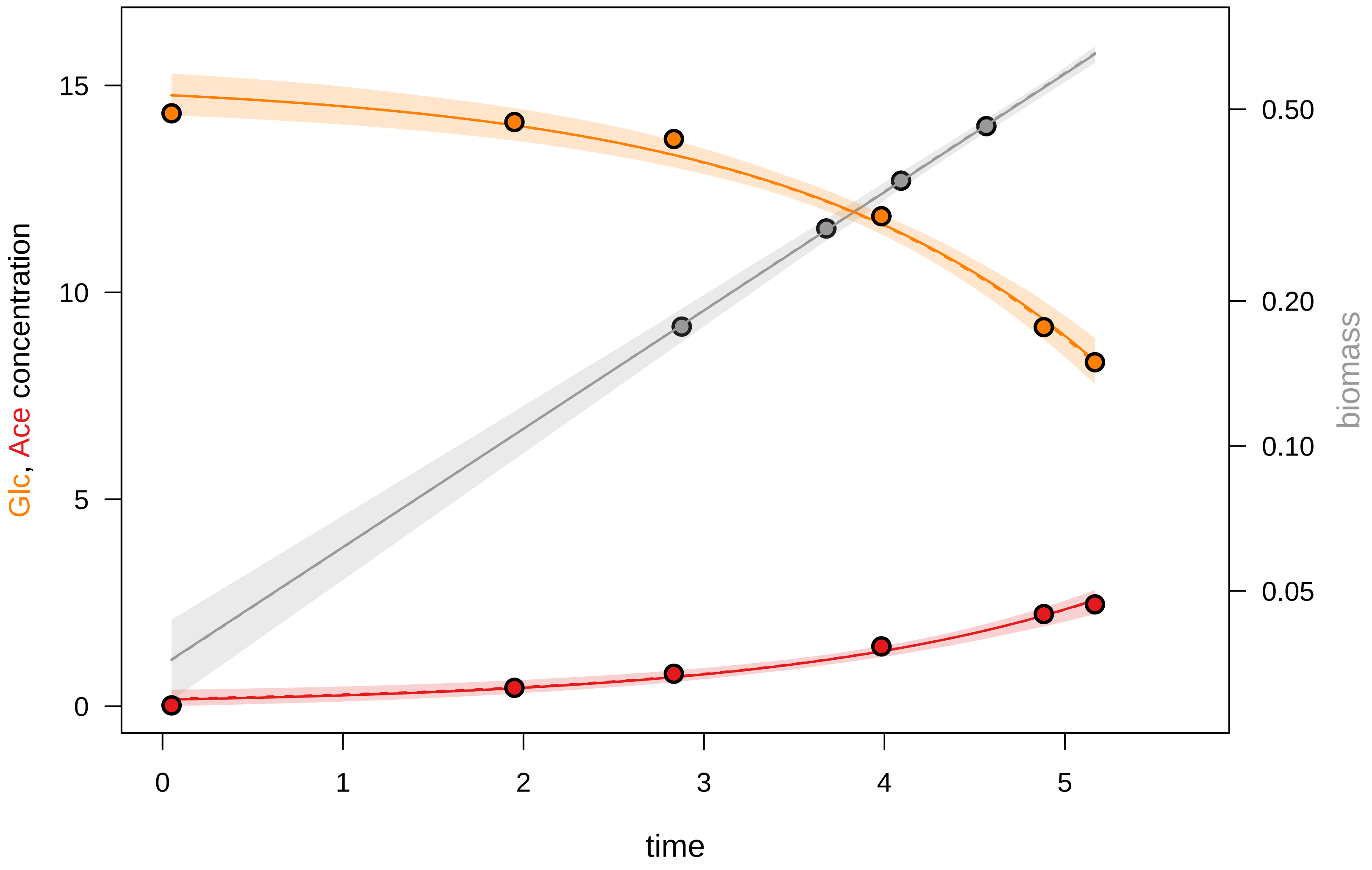

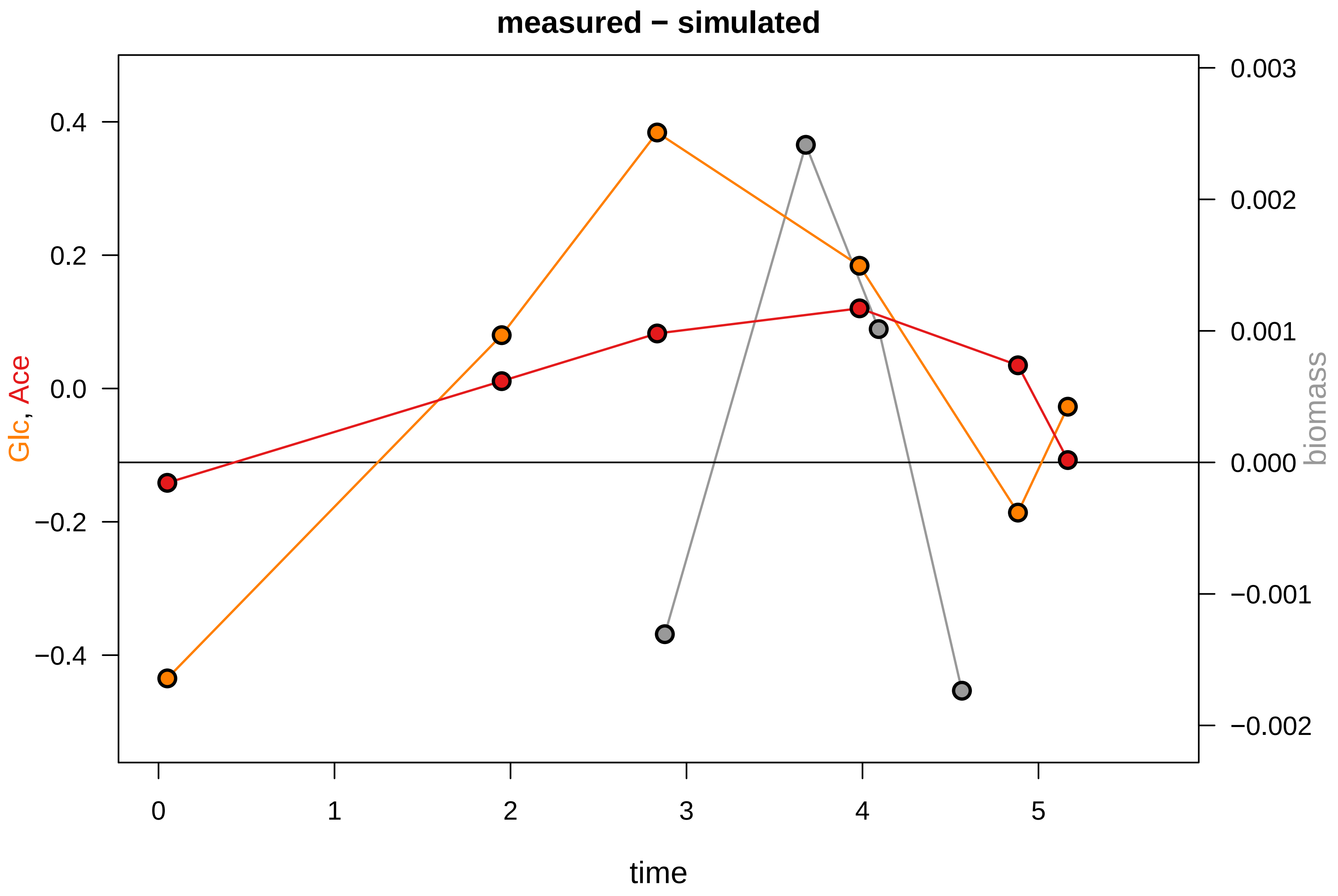

### KEIO_ROBOT3_31.pdf

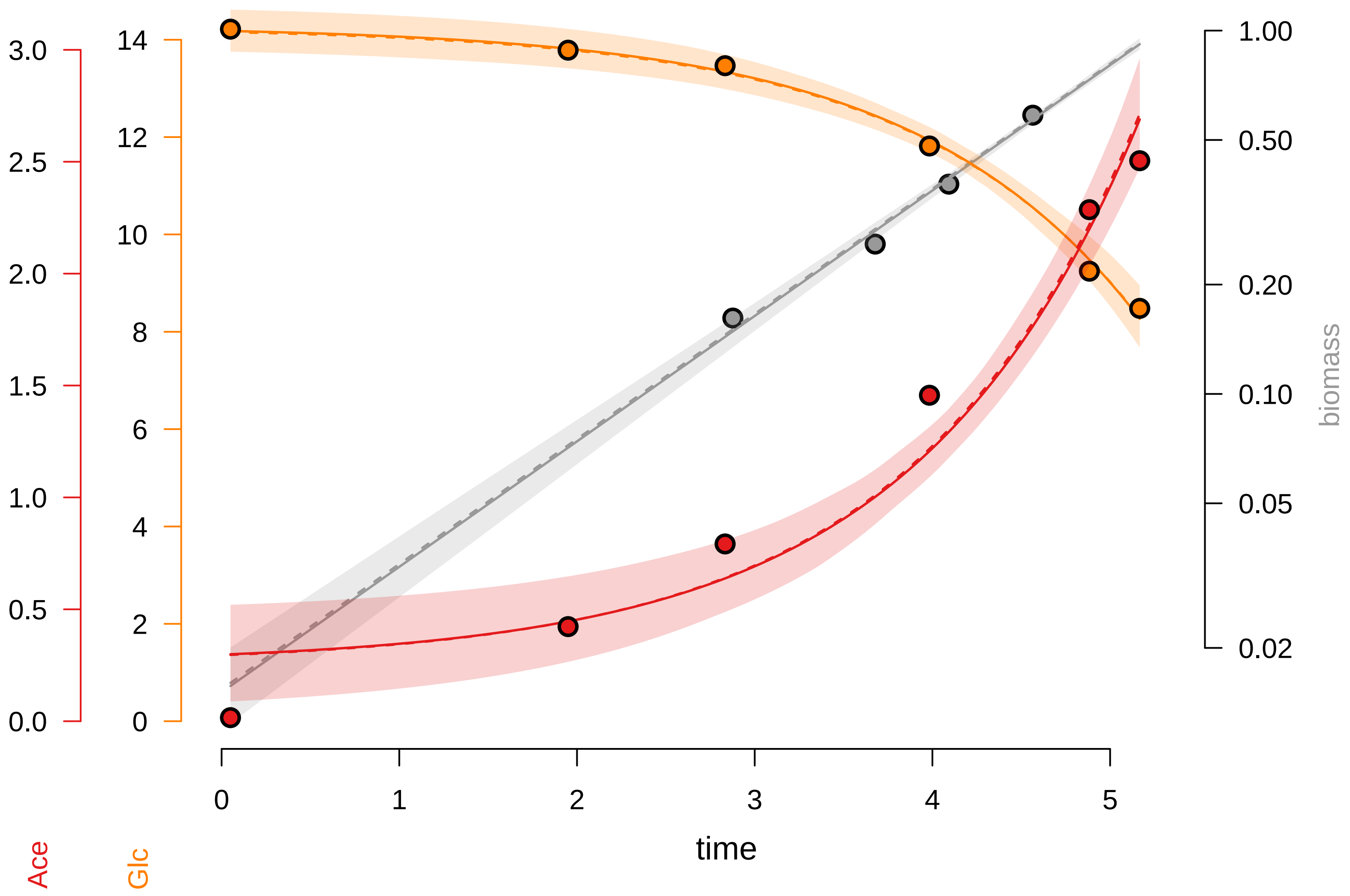

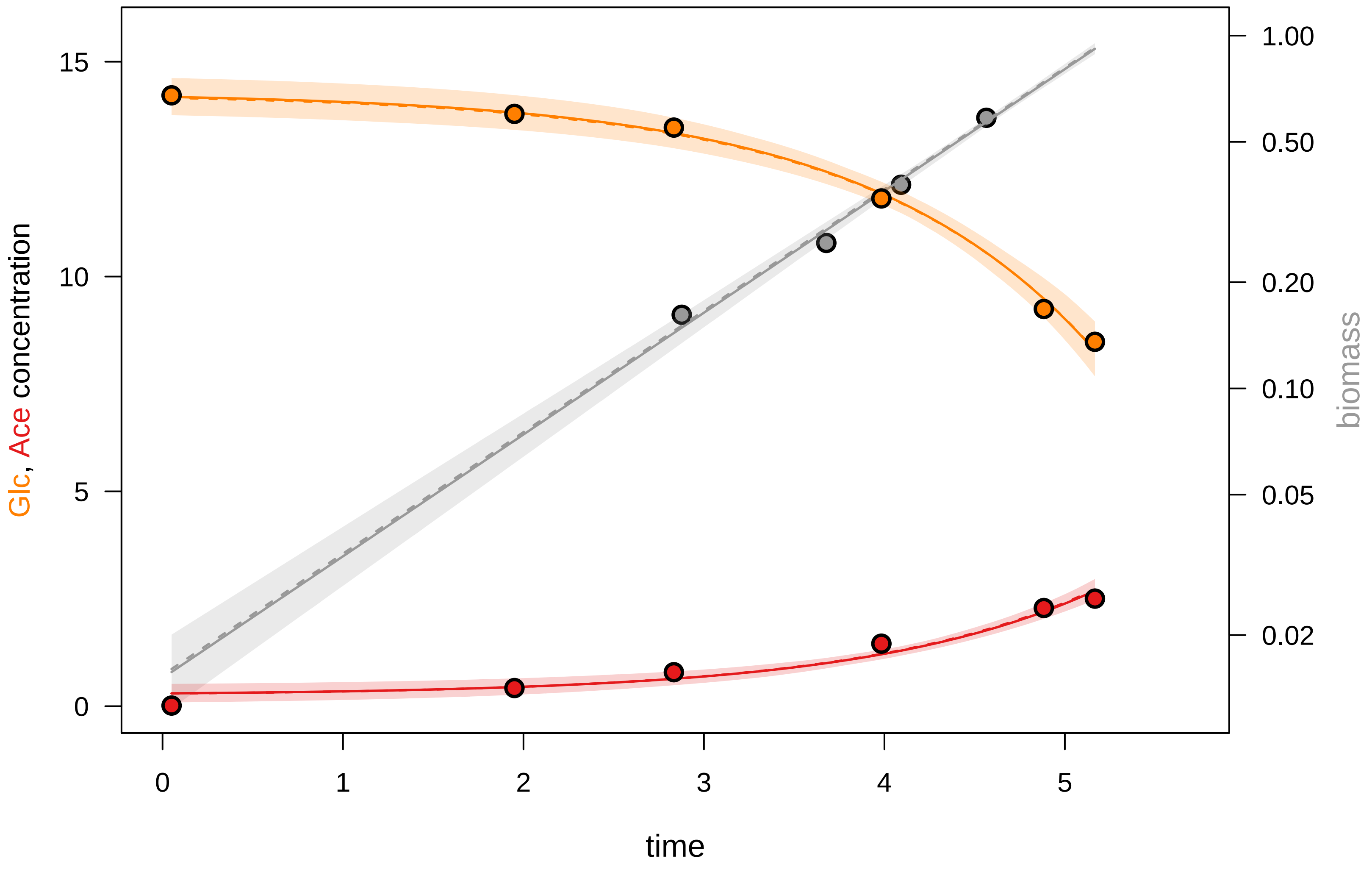

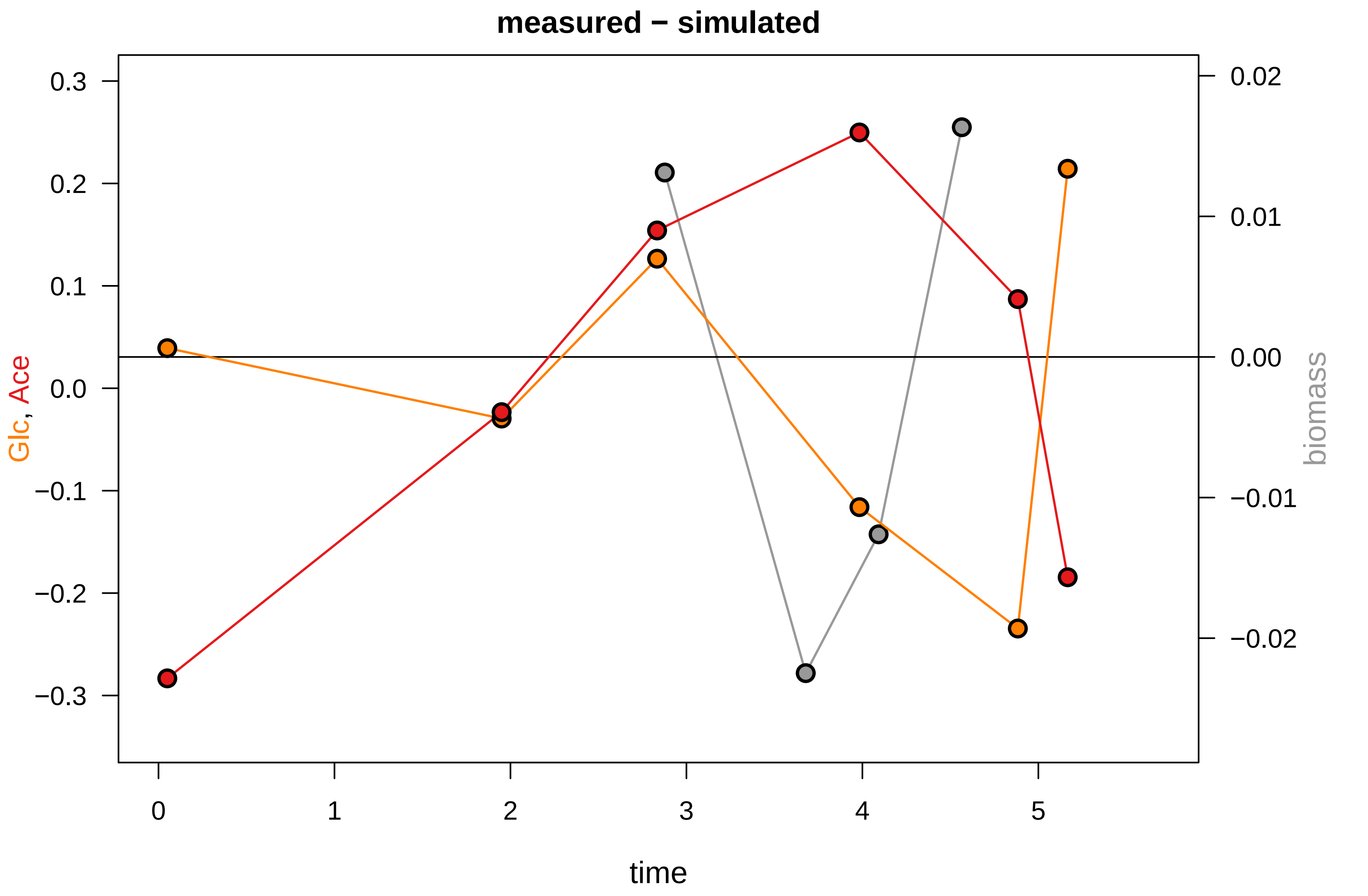

### KEIO_ROBOT3_46.pdf

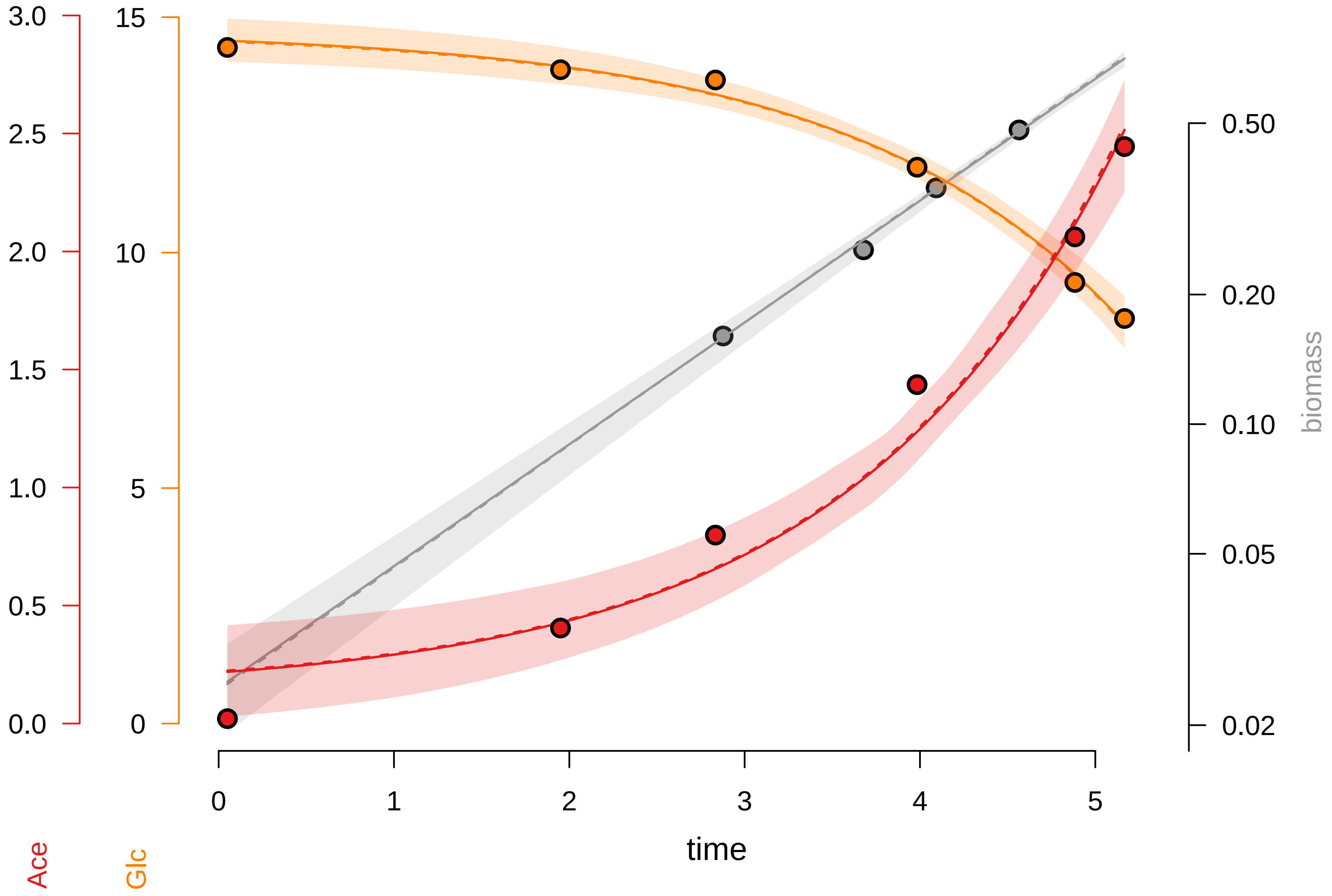

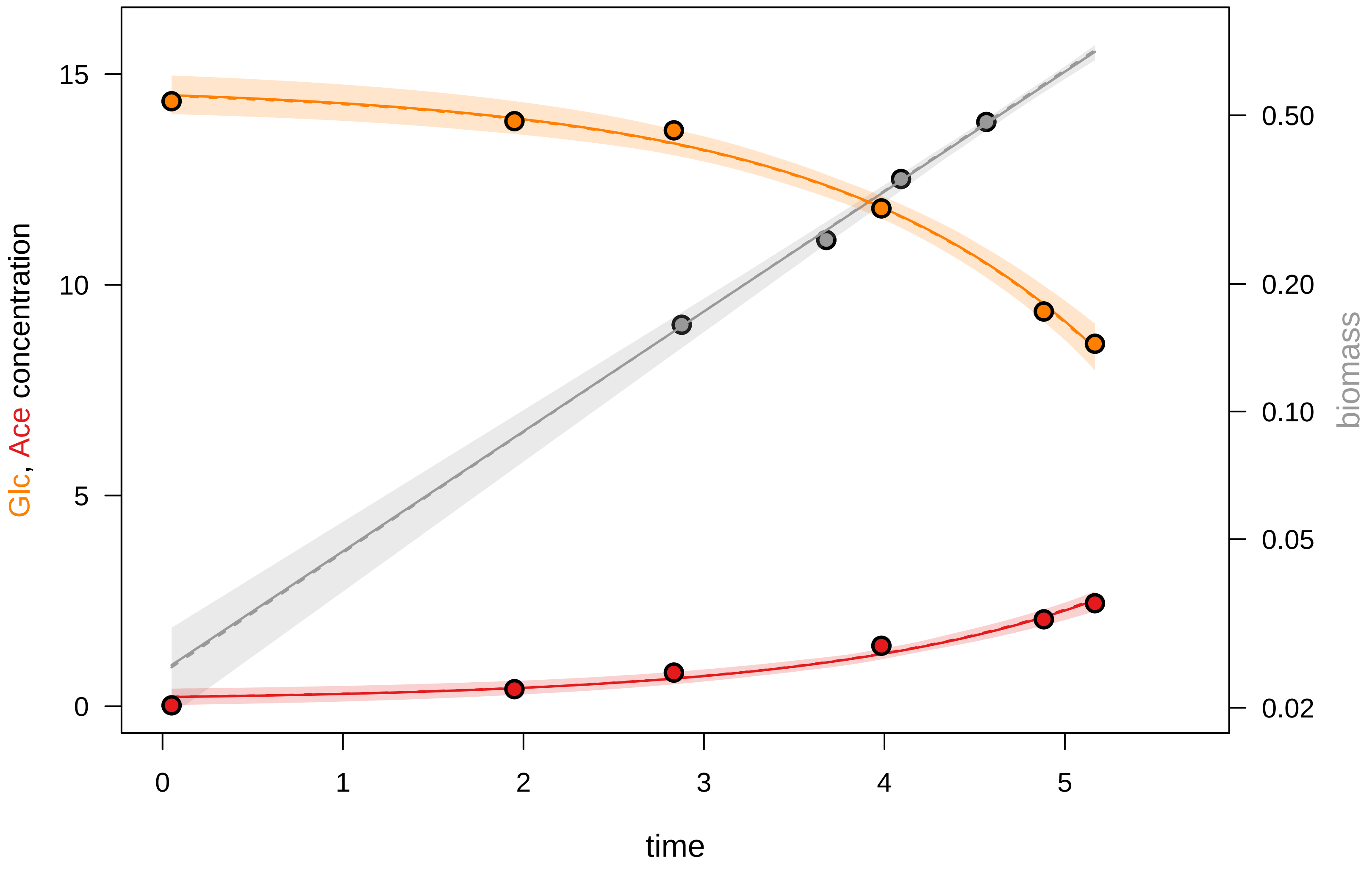

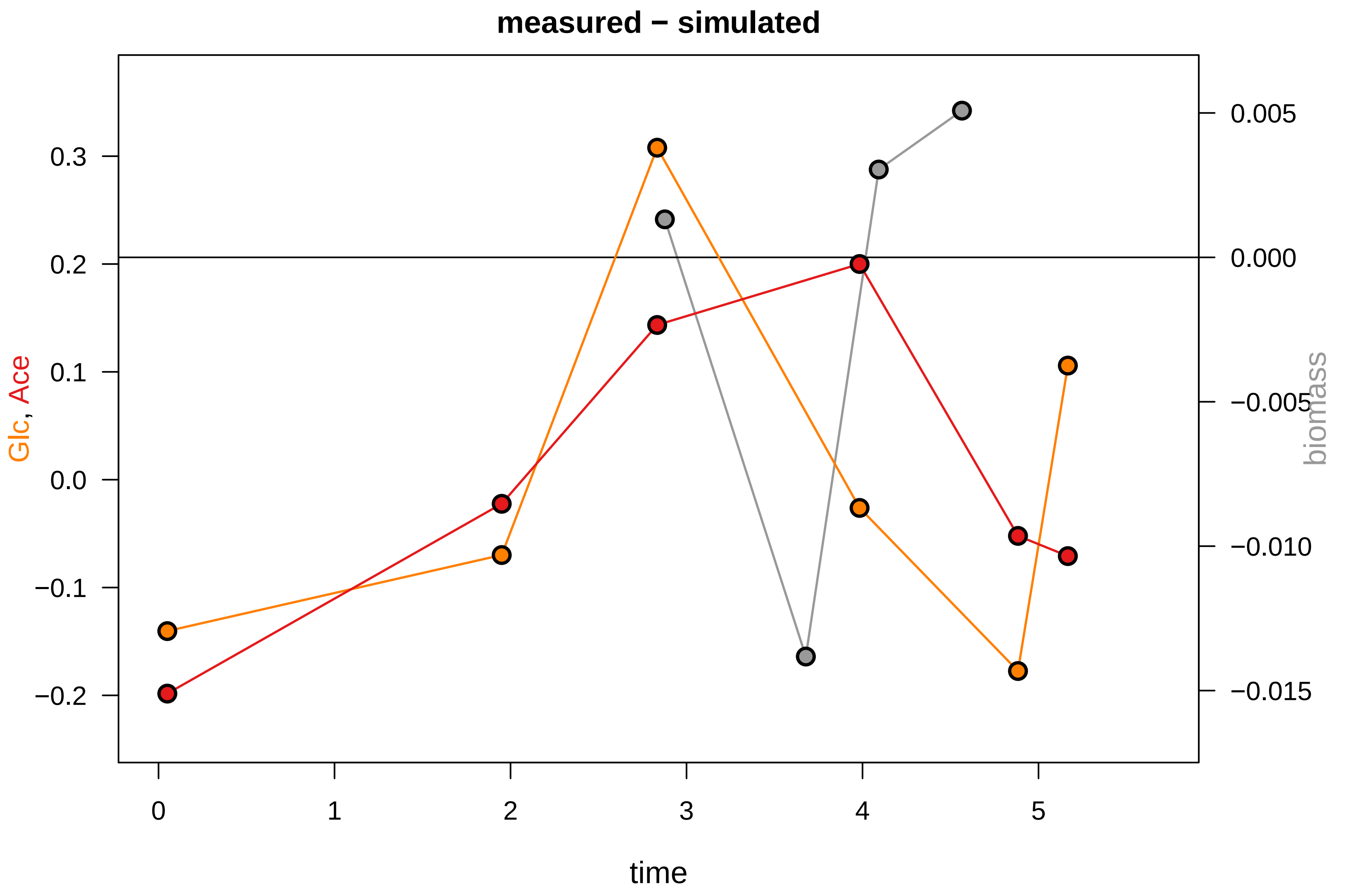

### KEIO_ROBOT4_13.pdf

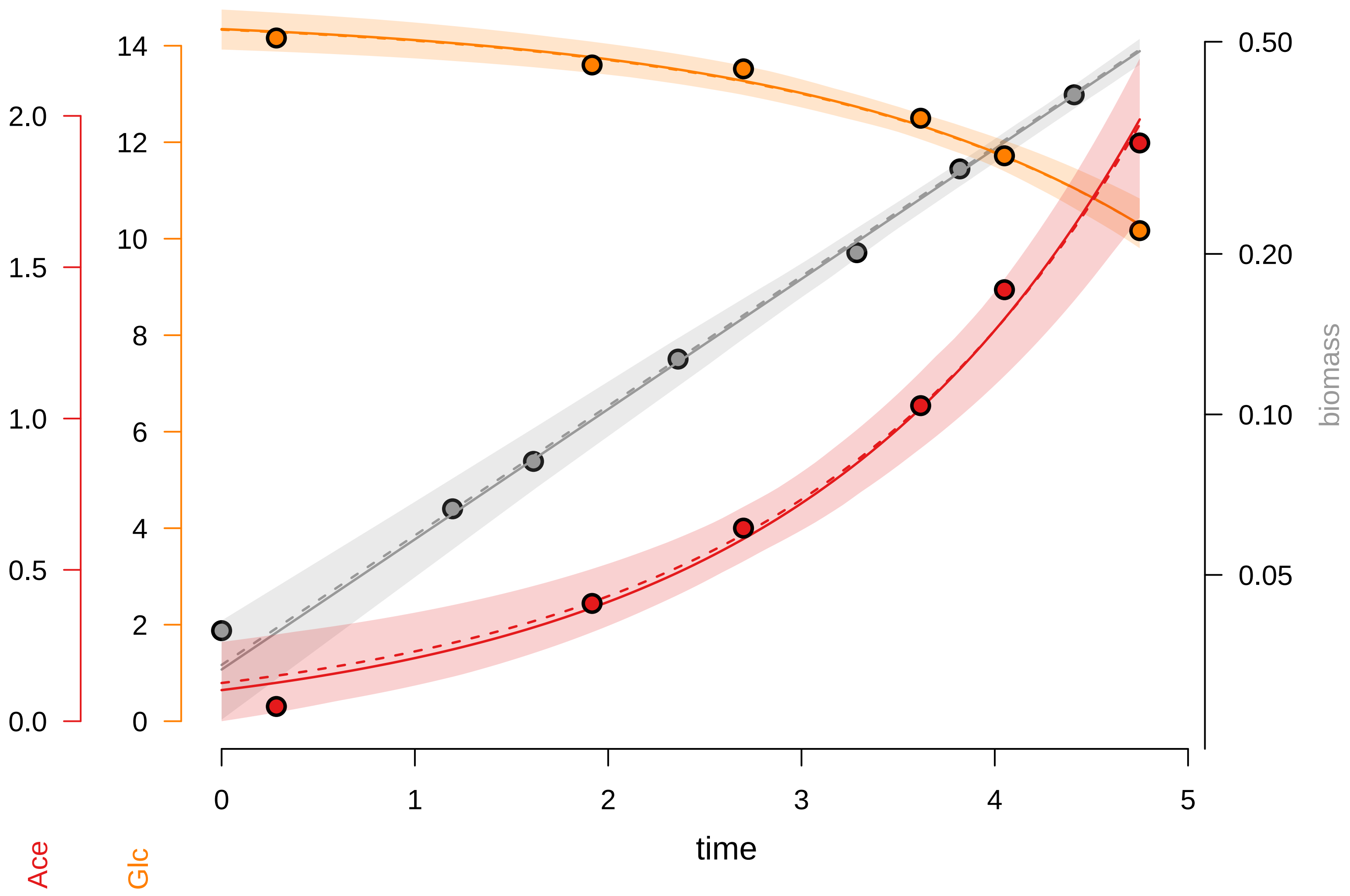

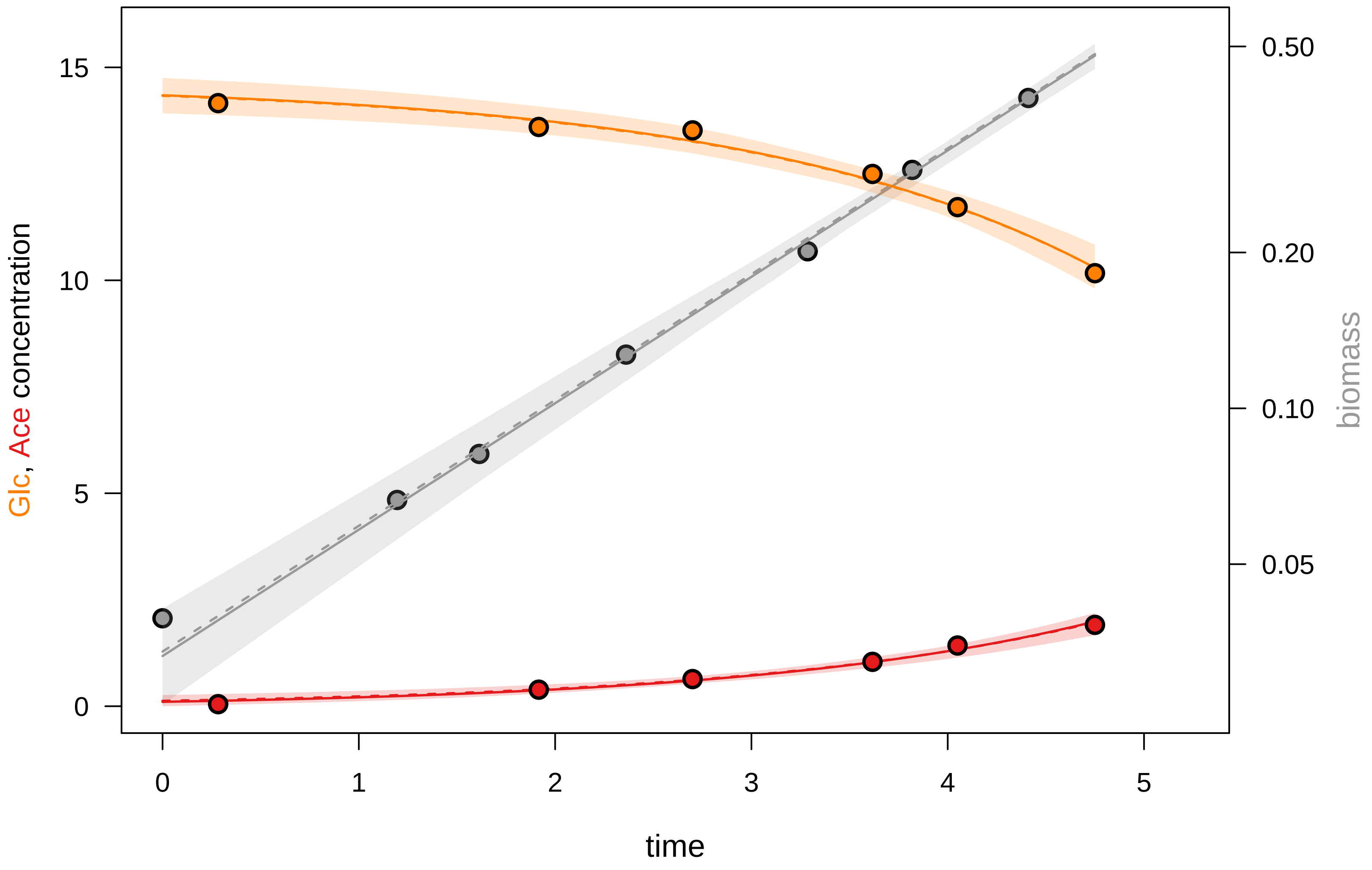

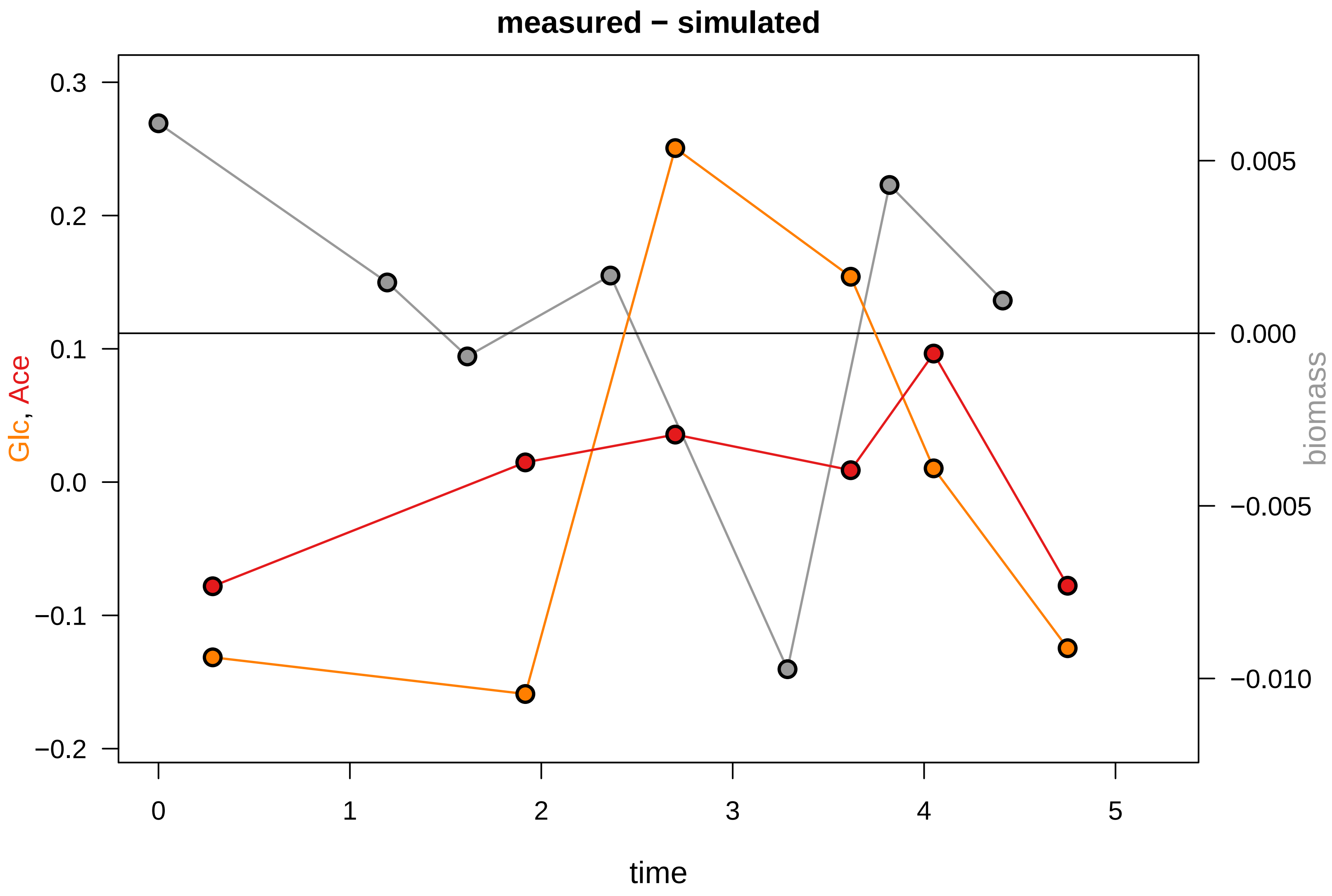

### KEIO_ROBOT4_22.pdf

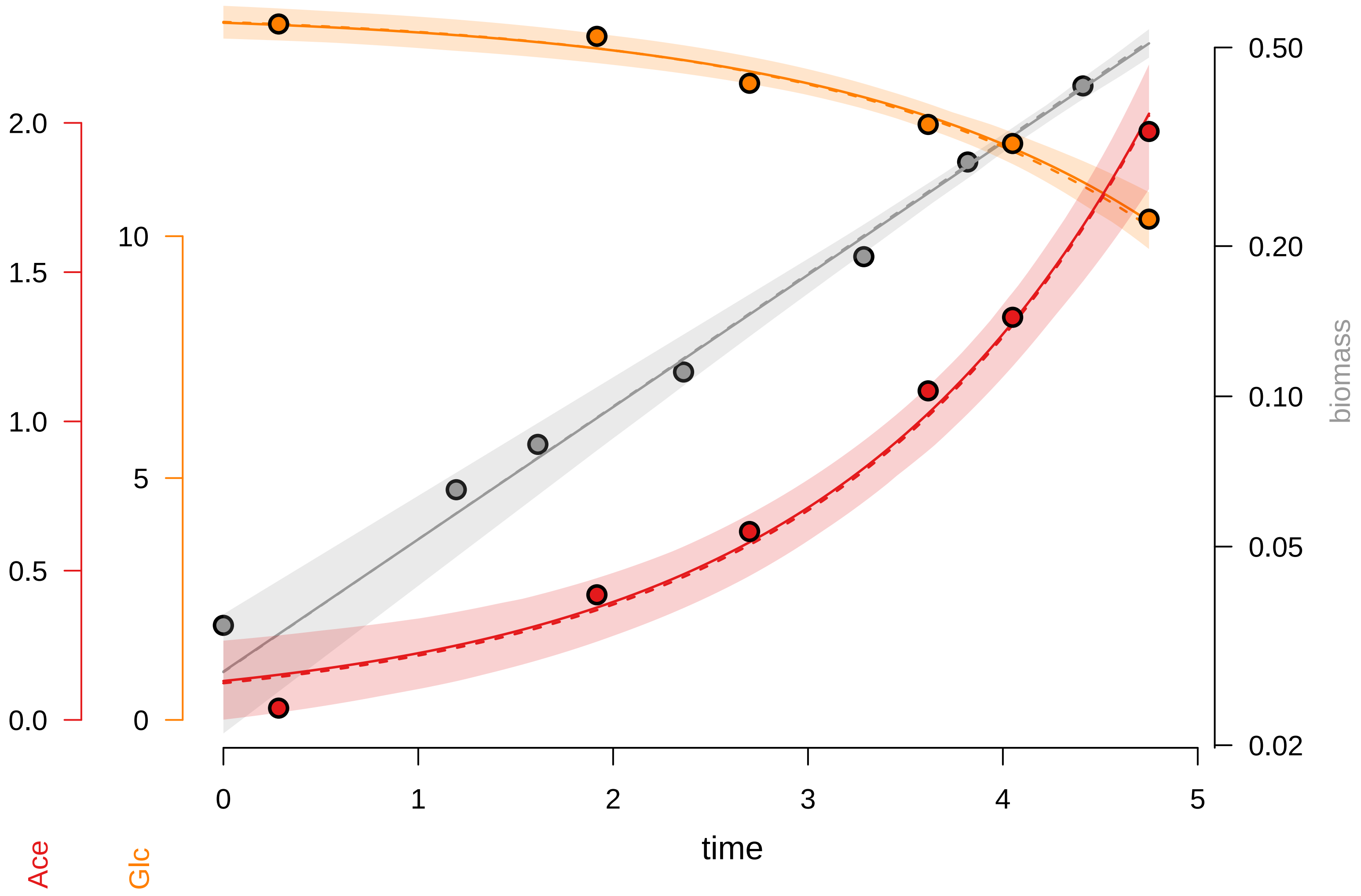

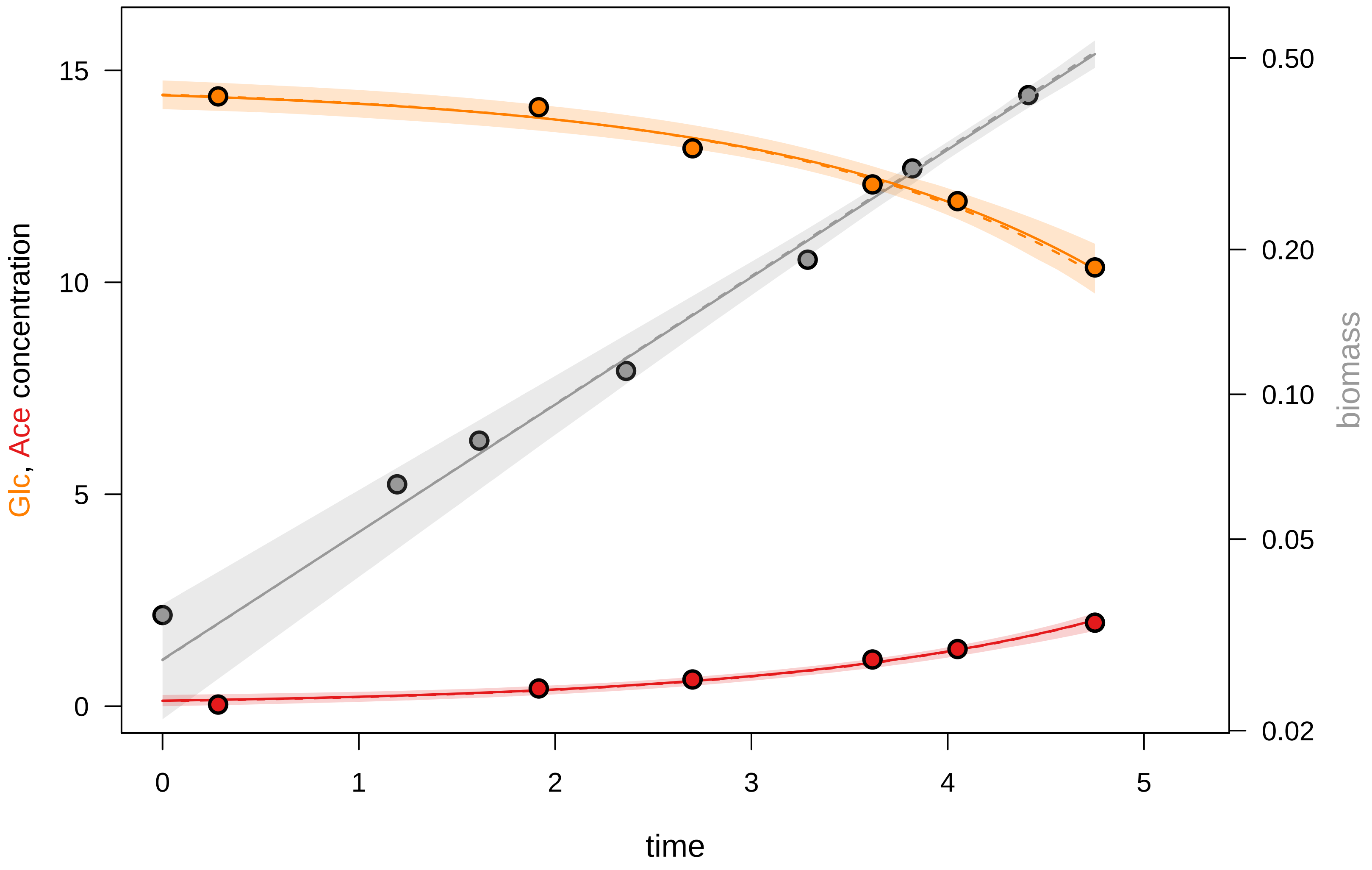

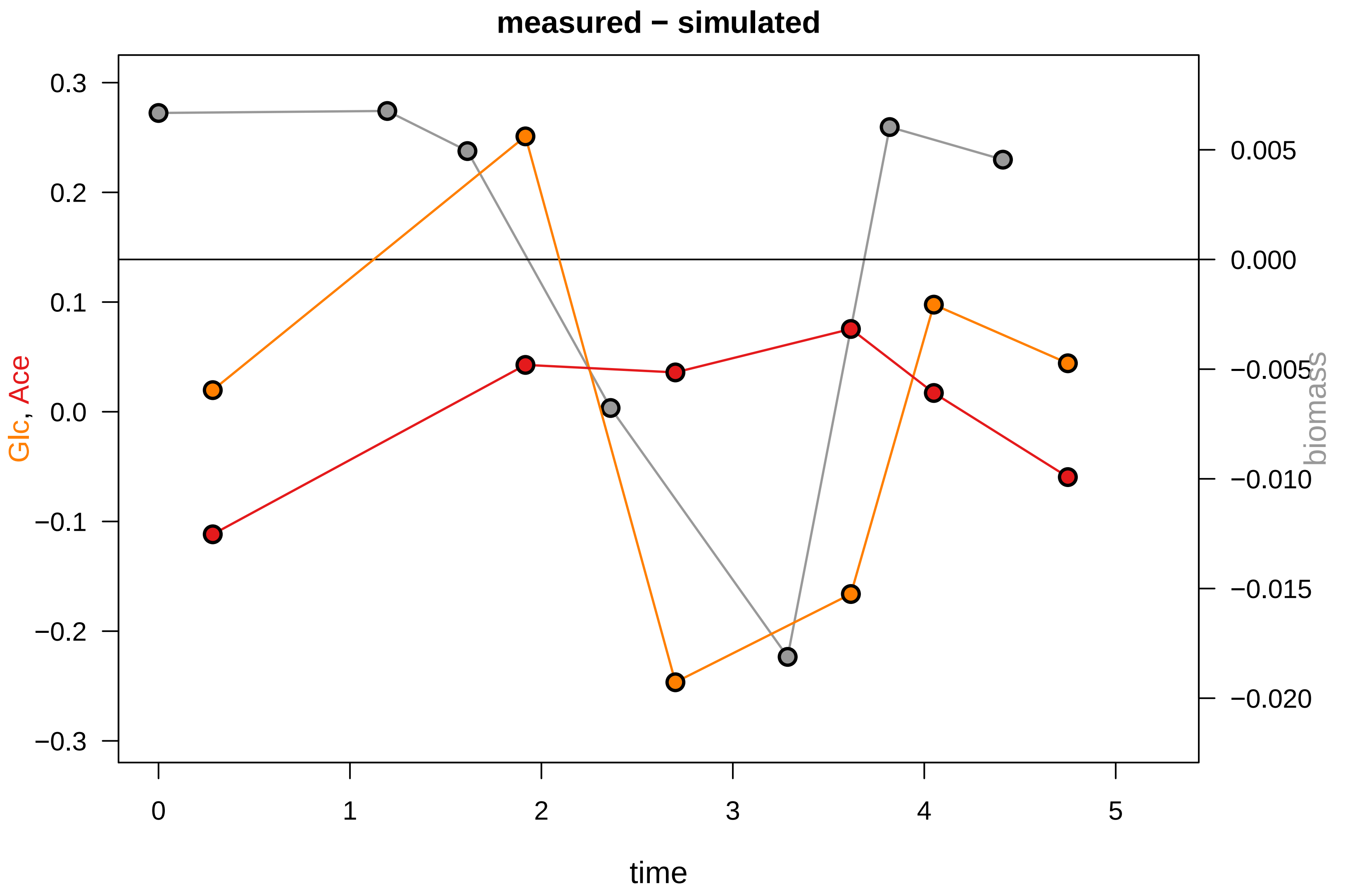

### KEIO_ROBOT4_26.pdf

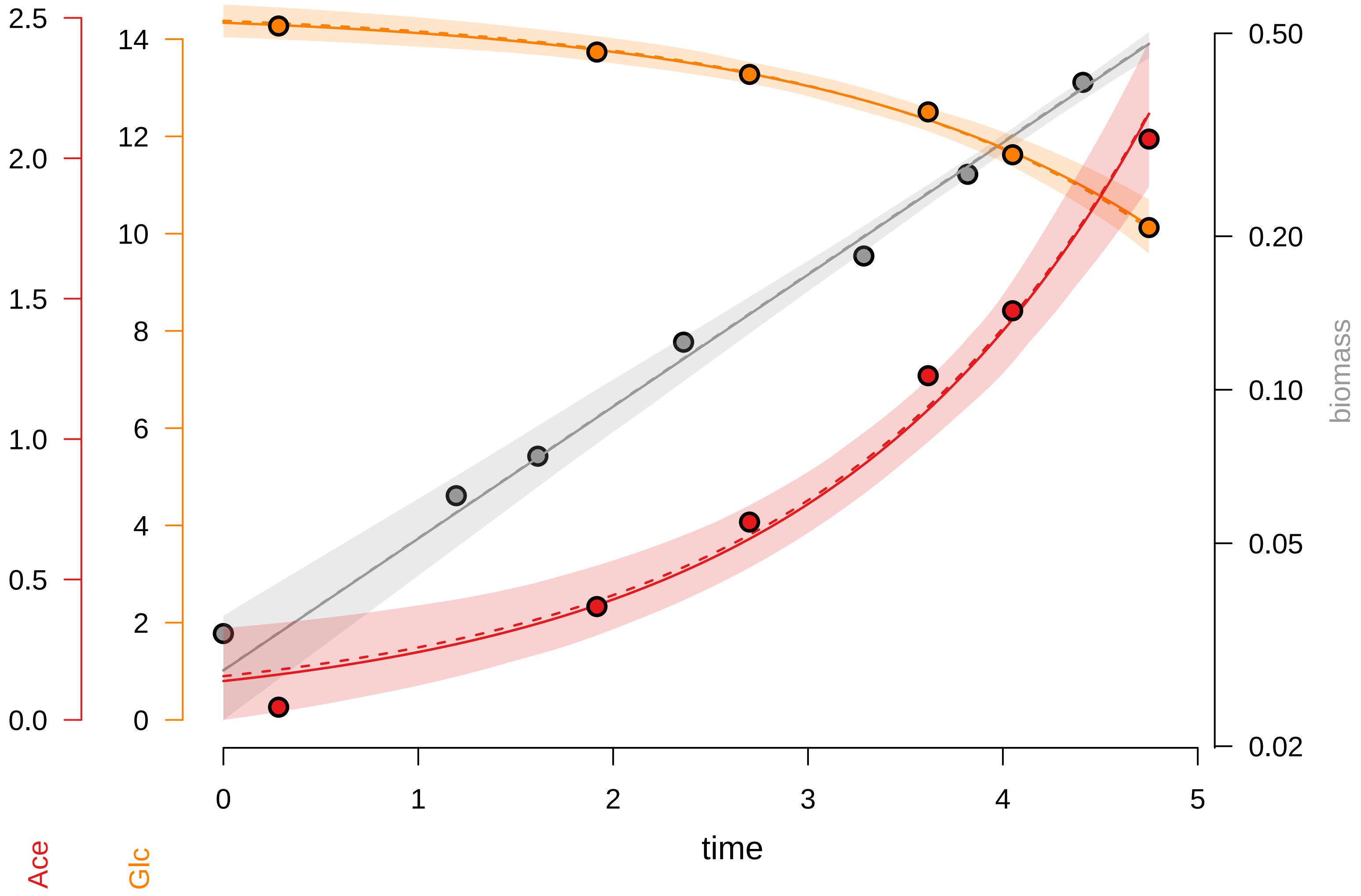

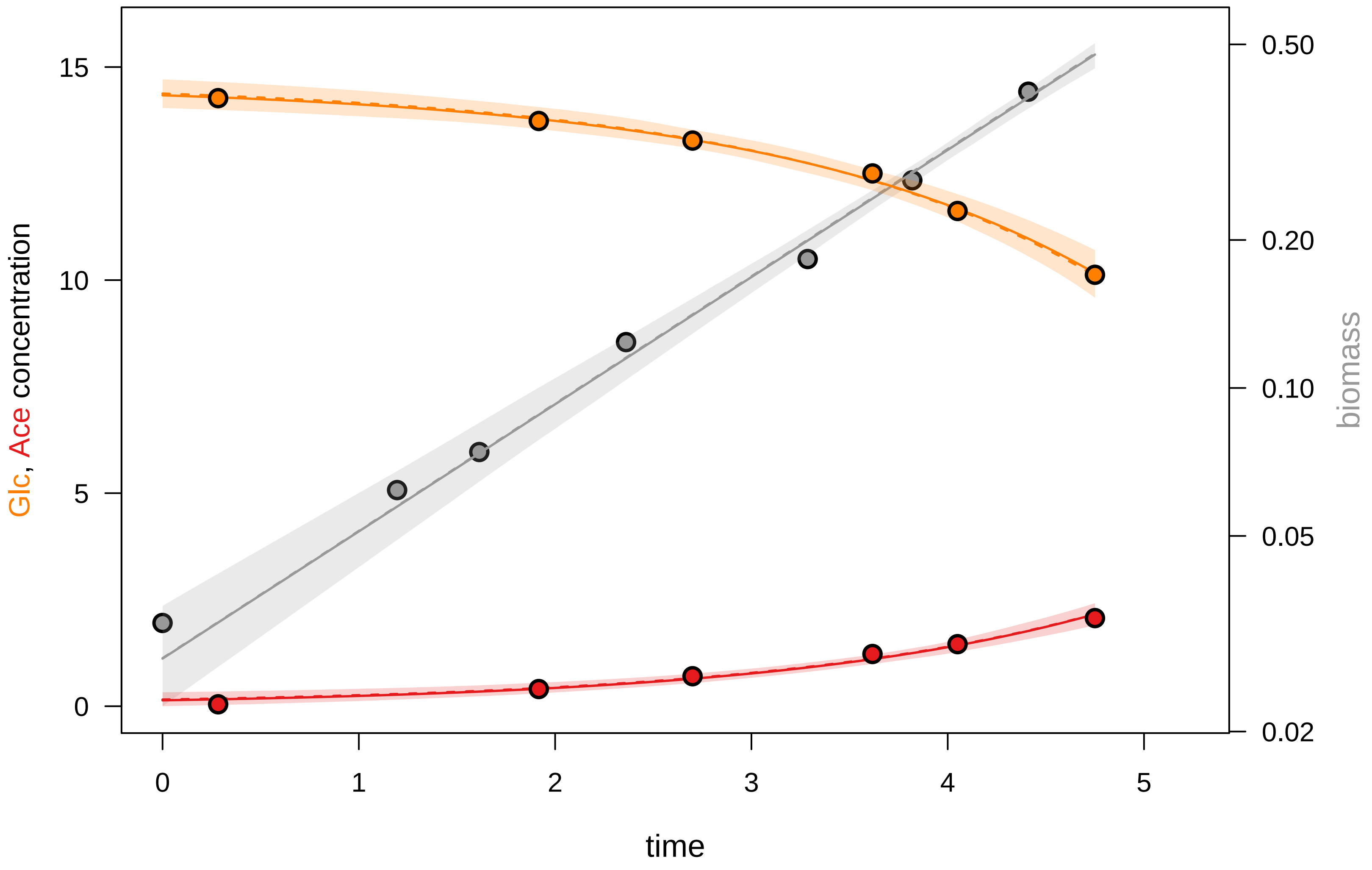

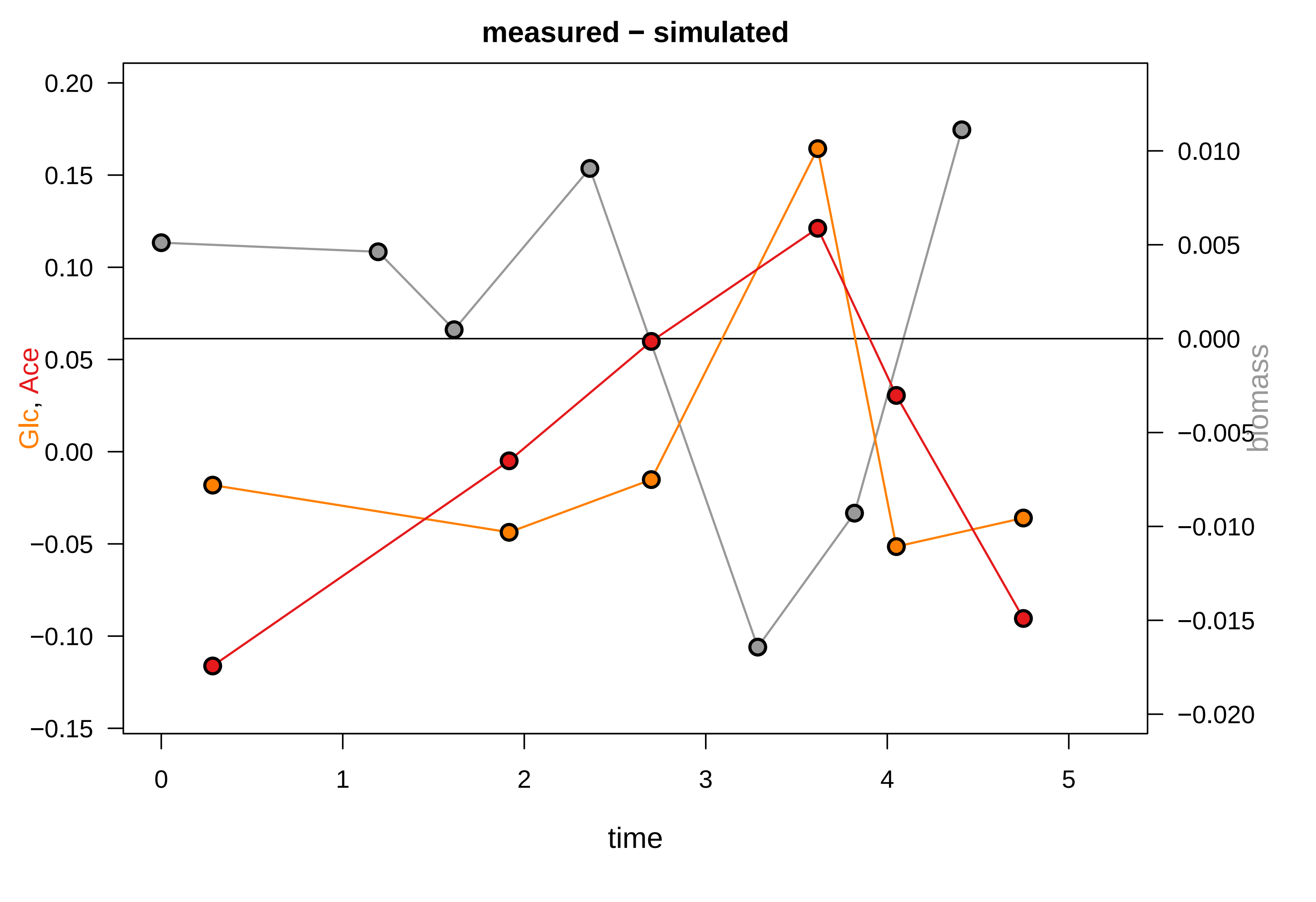

### KEIO_ROBOT4_27.pdf

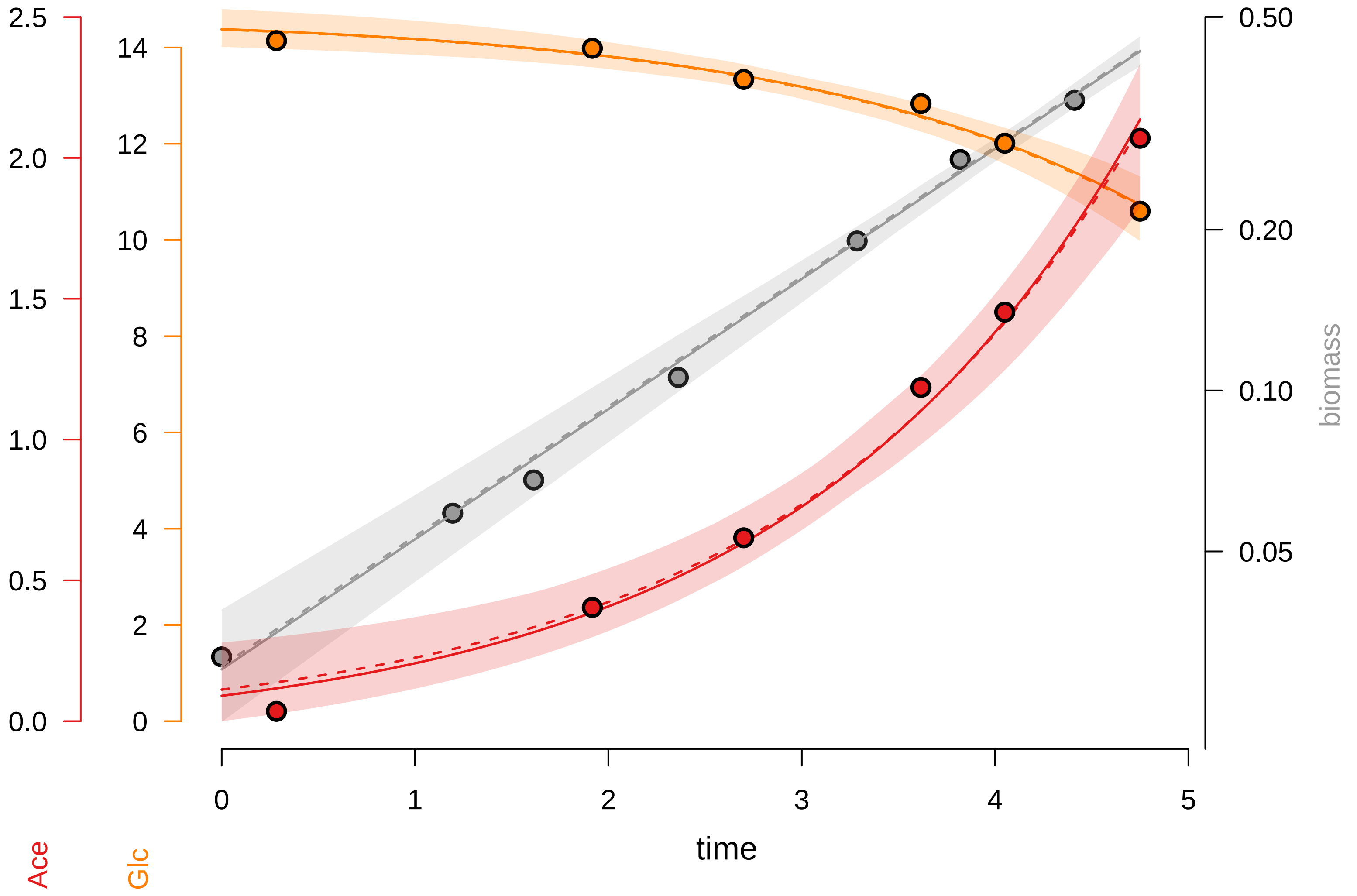

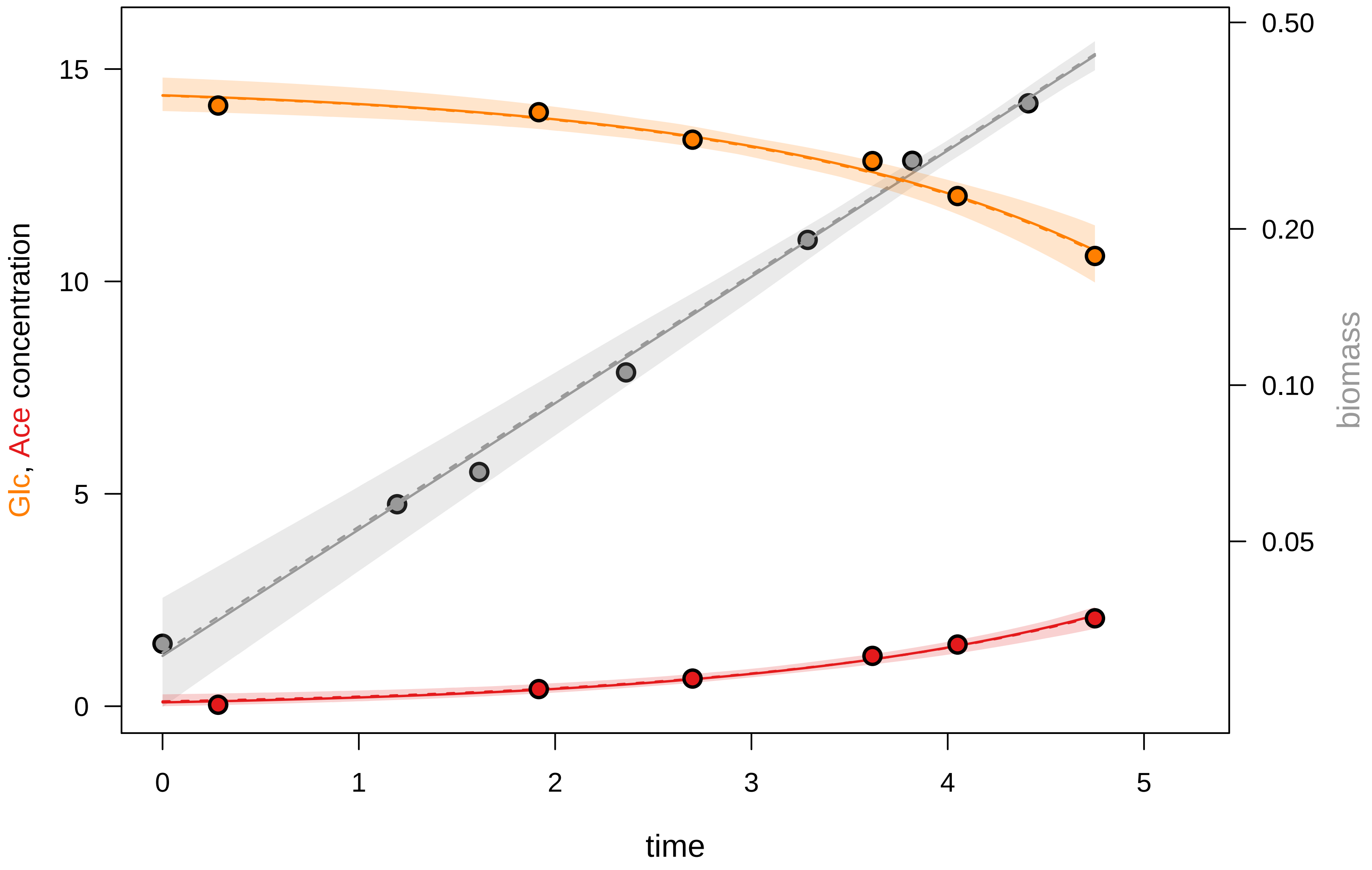

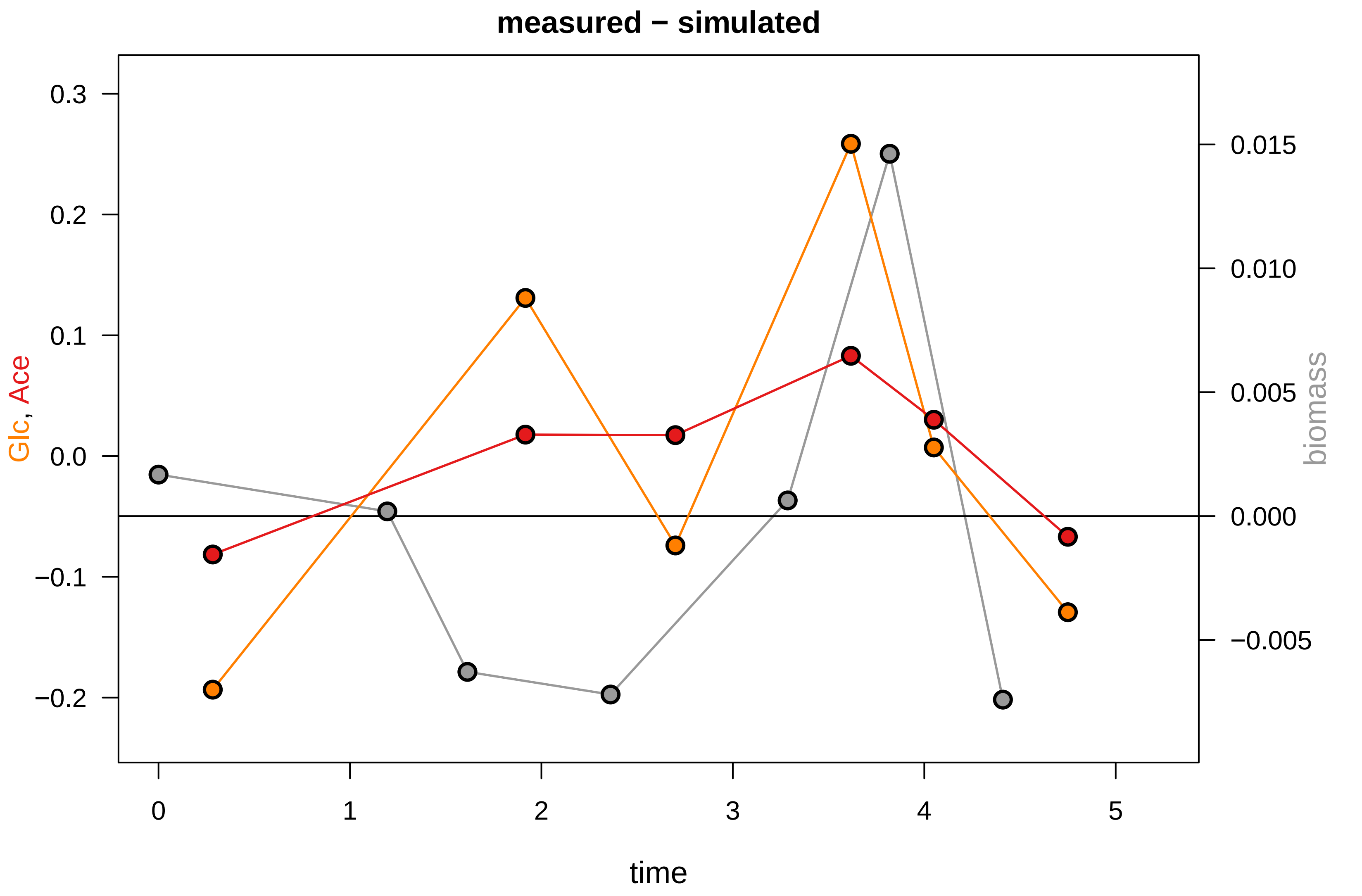

### KEIO_ROBOT4_30.pdf

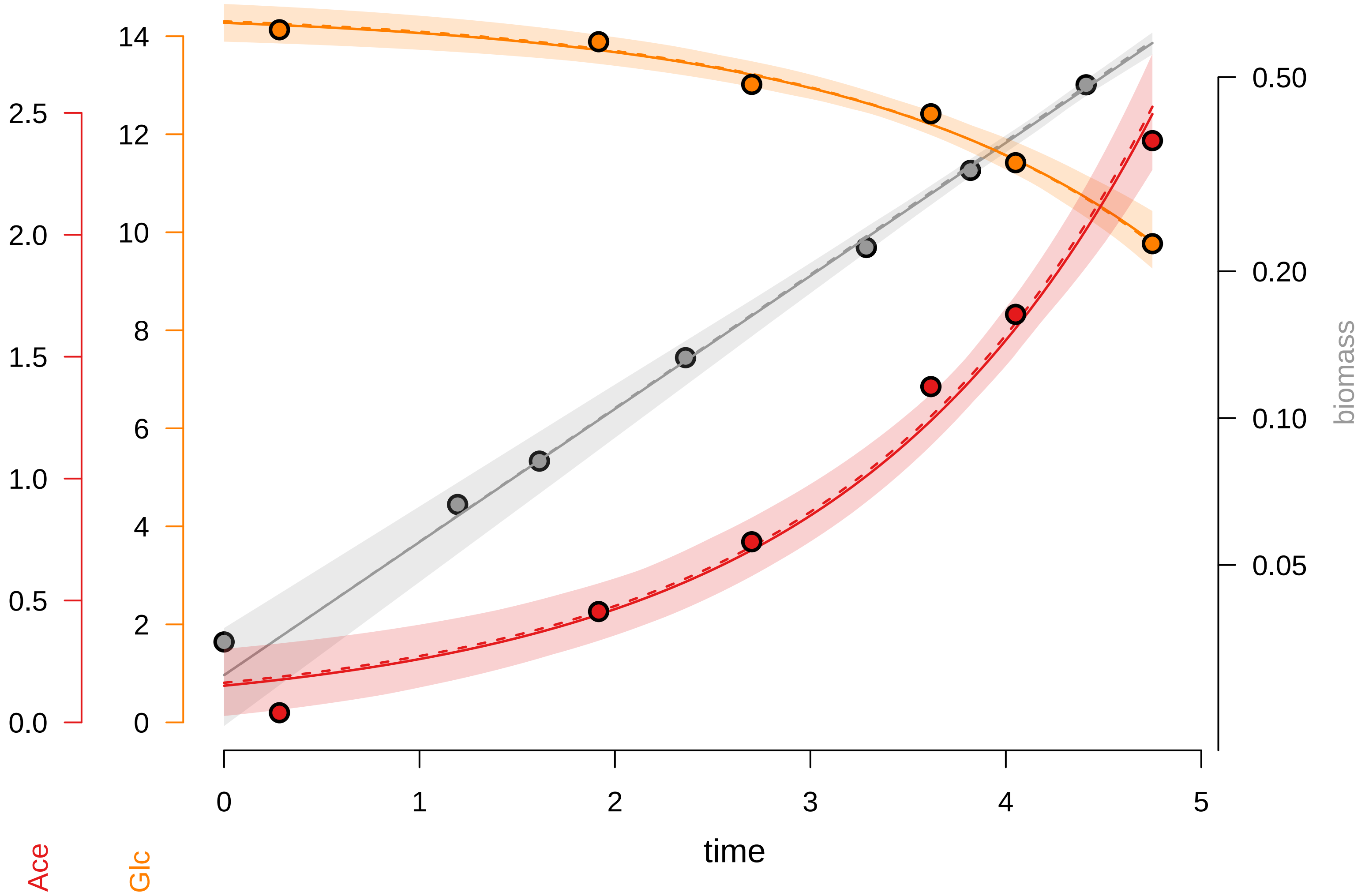

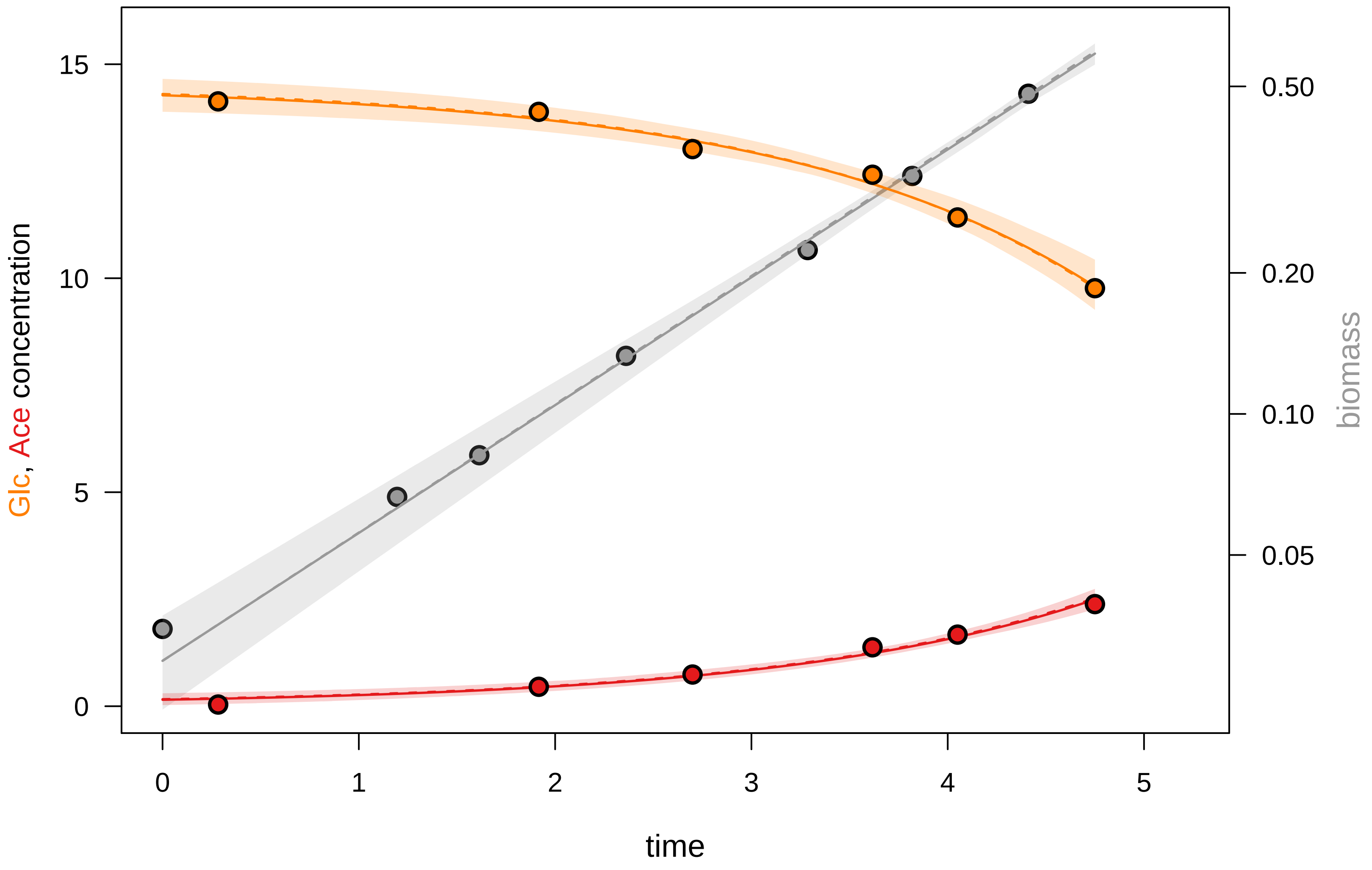

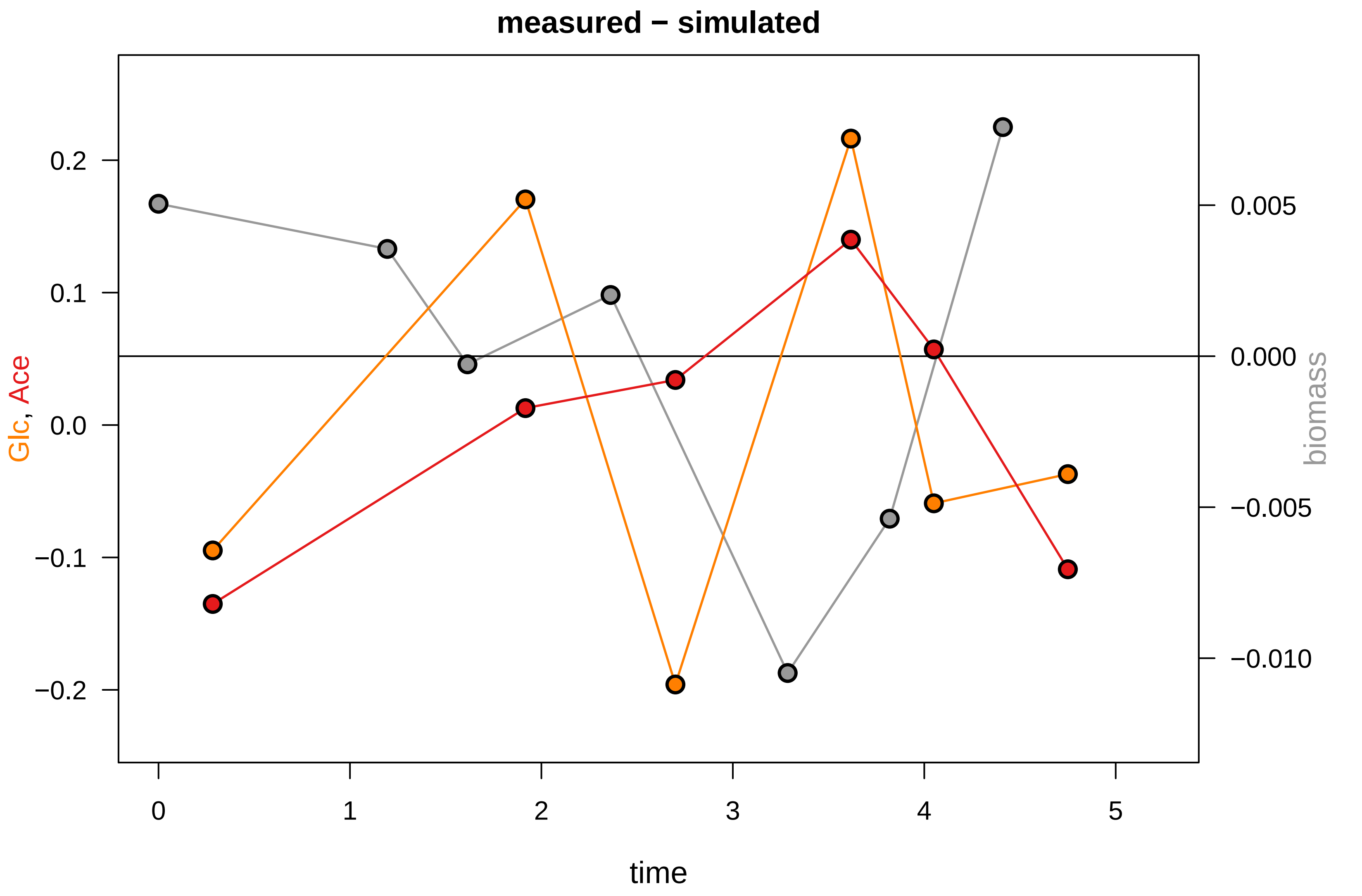

### KEIO_ROBOT4_40.pdf

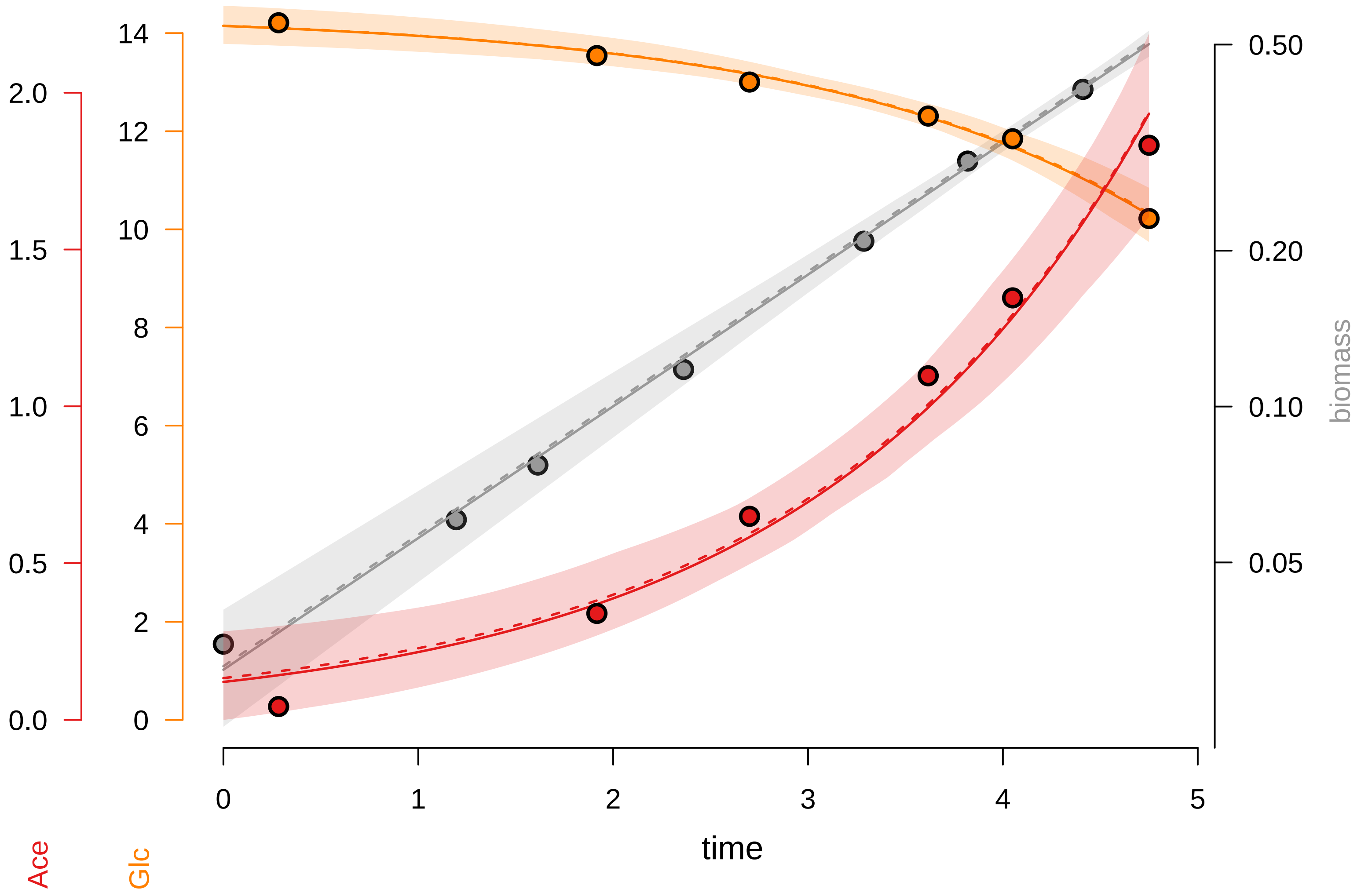

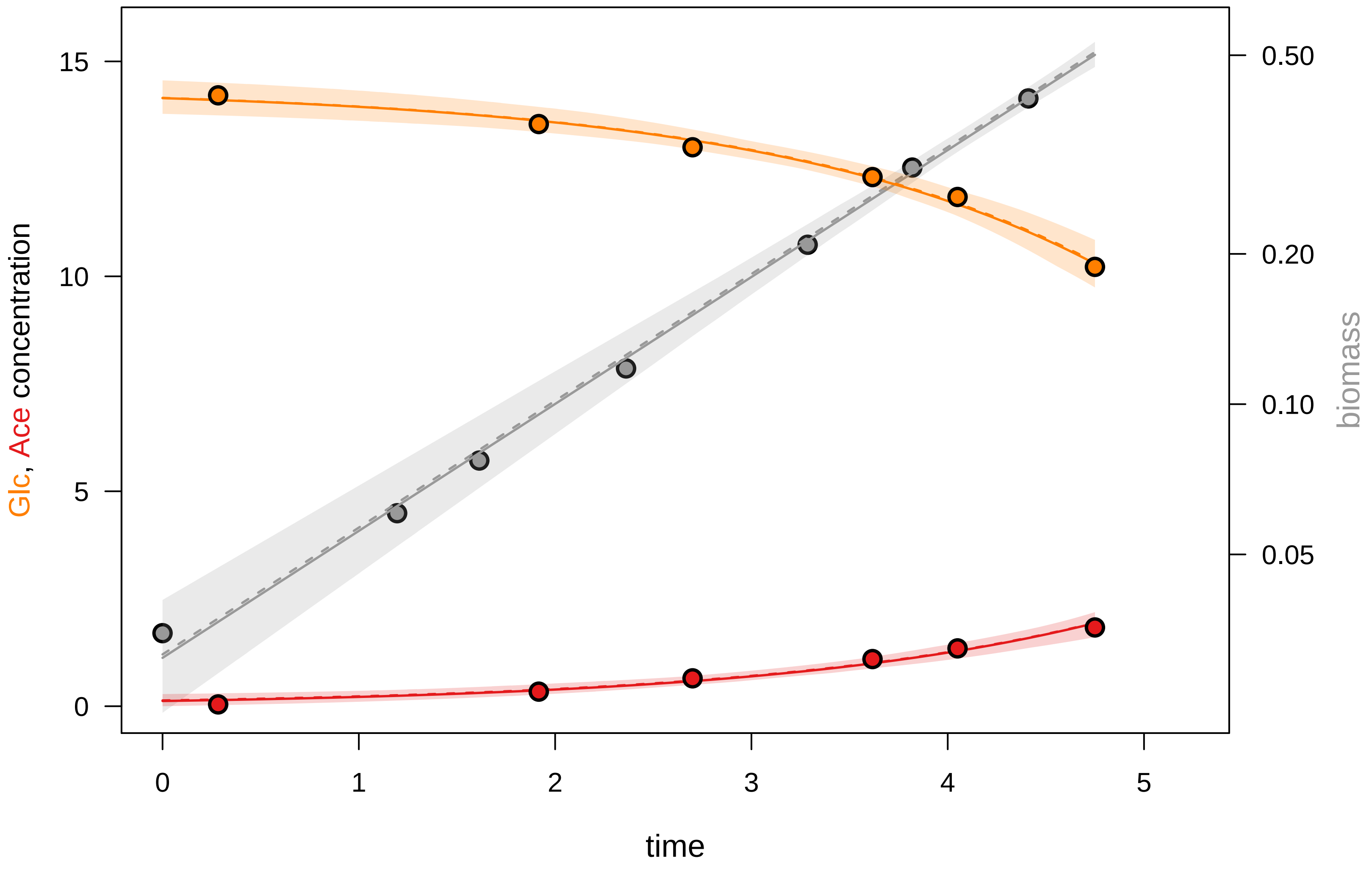

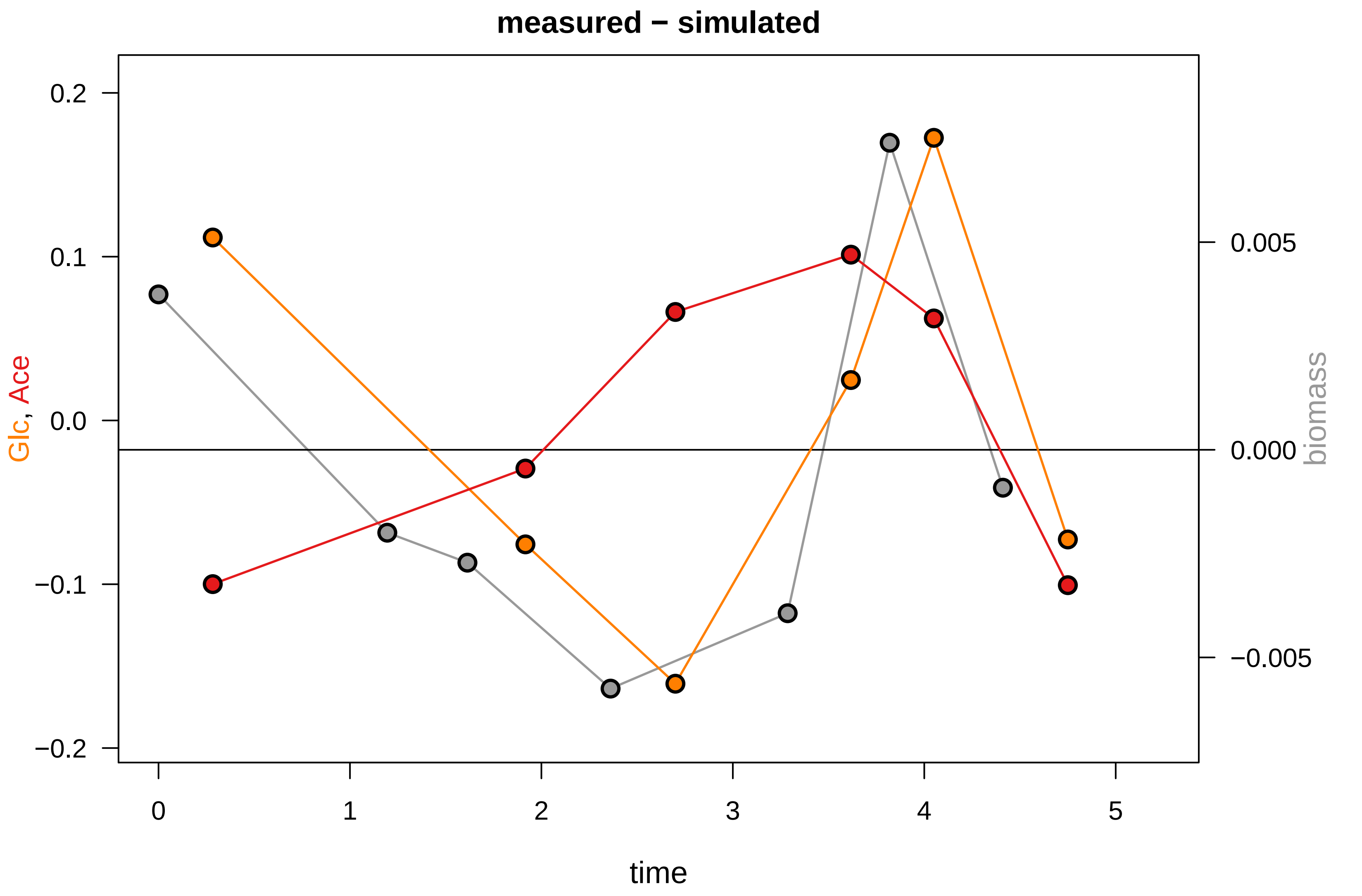

### KEIO_ROBOT5_1.pdf

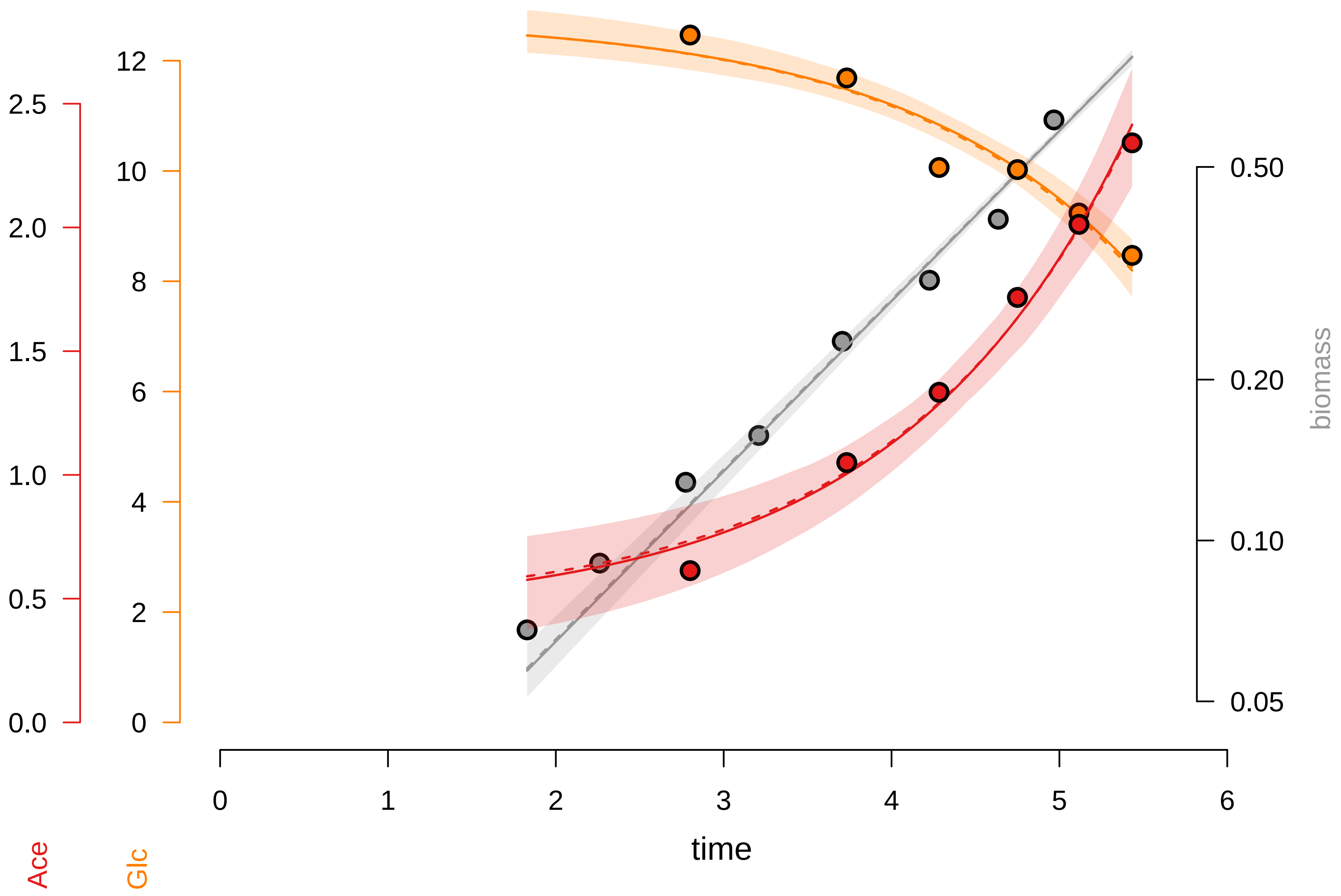

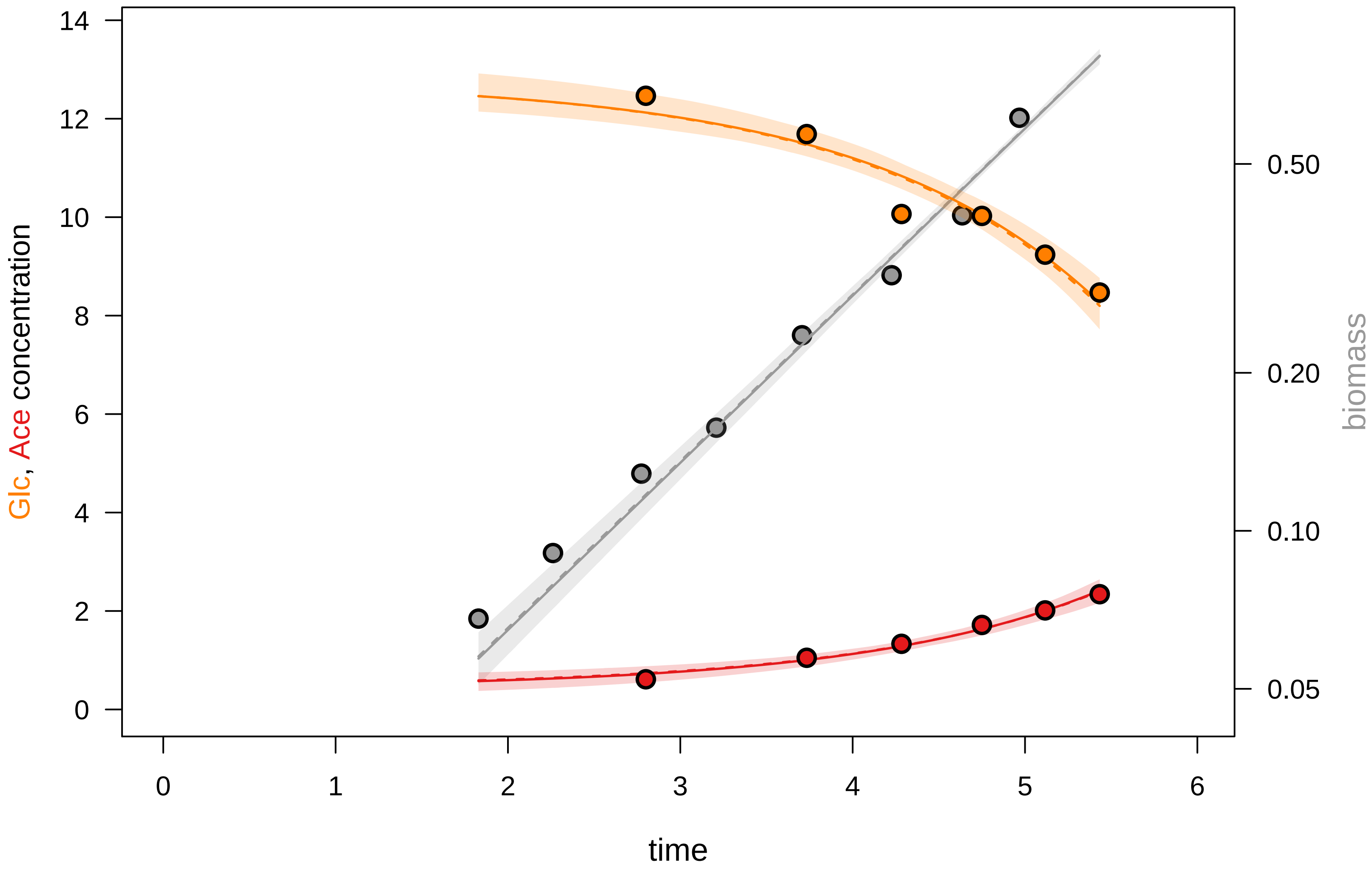

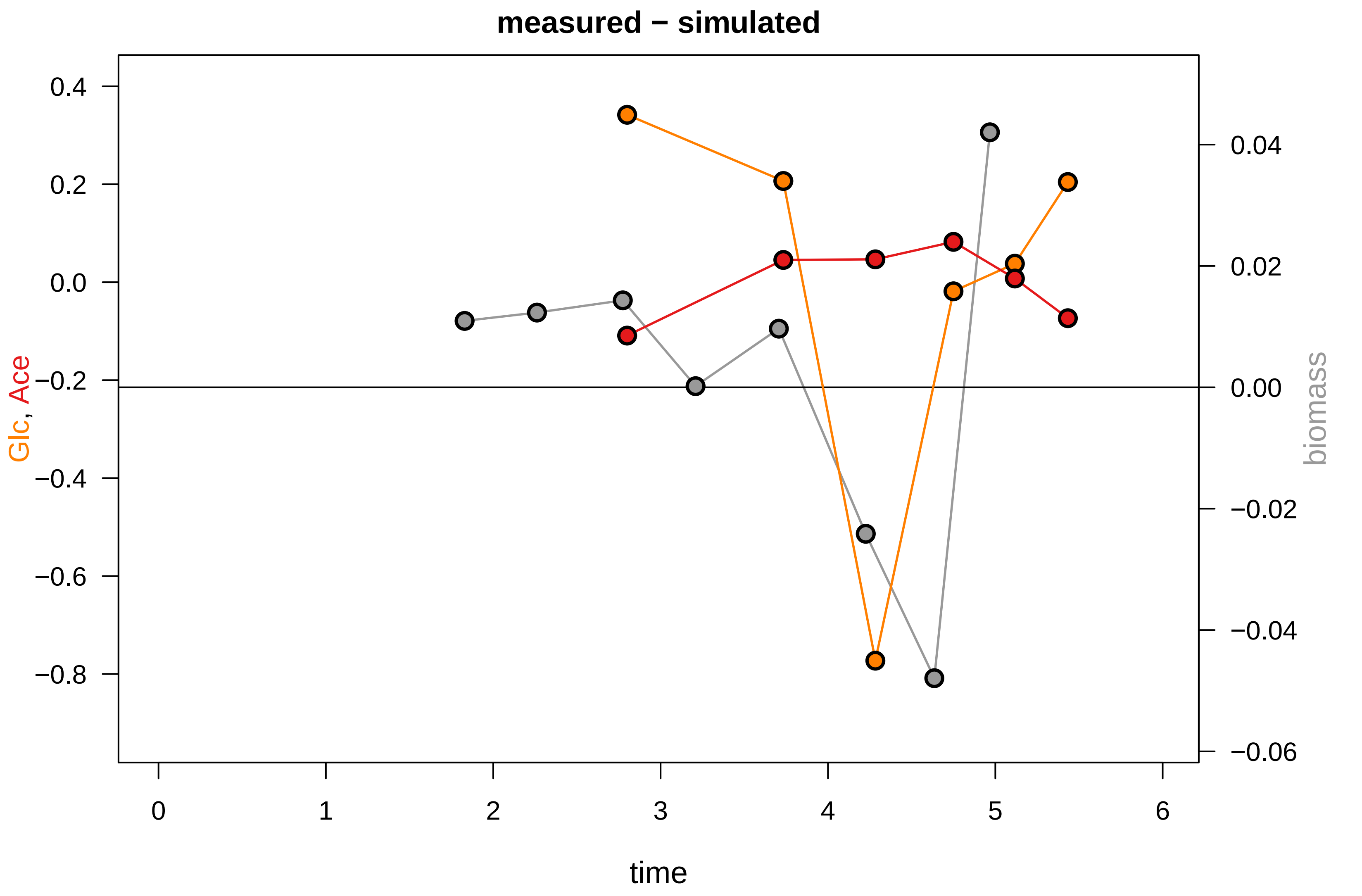

### KEIO_ROBOT5_14.pdf

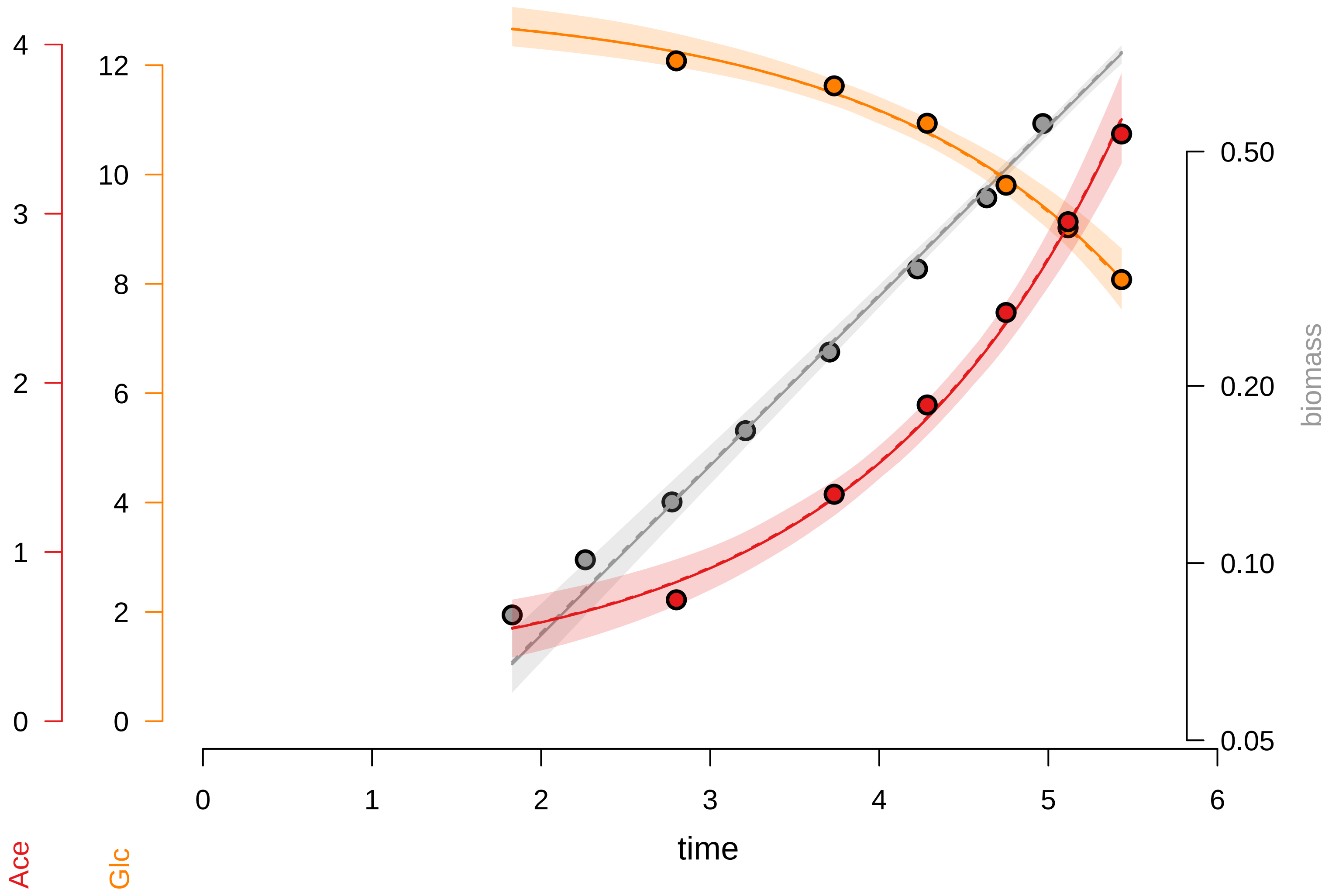

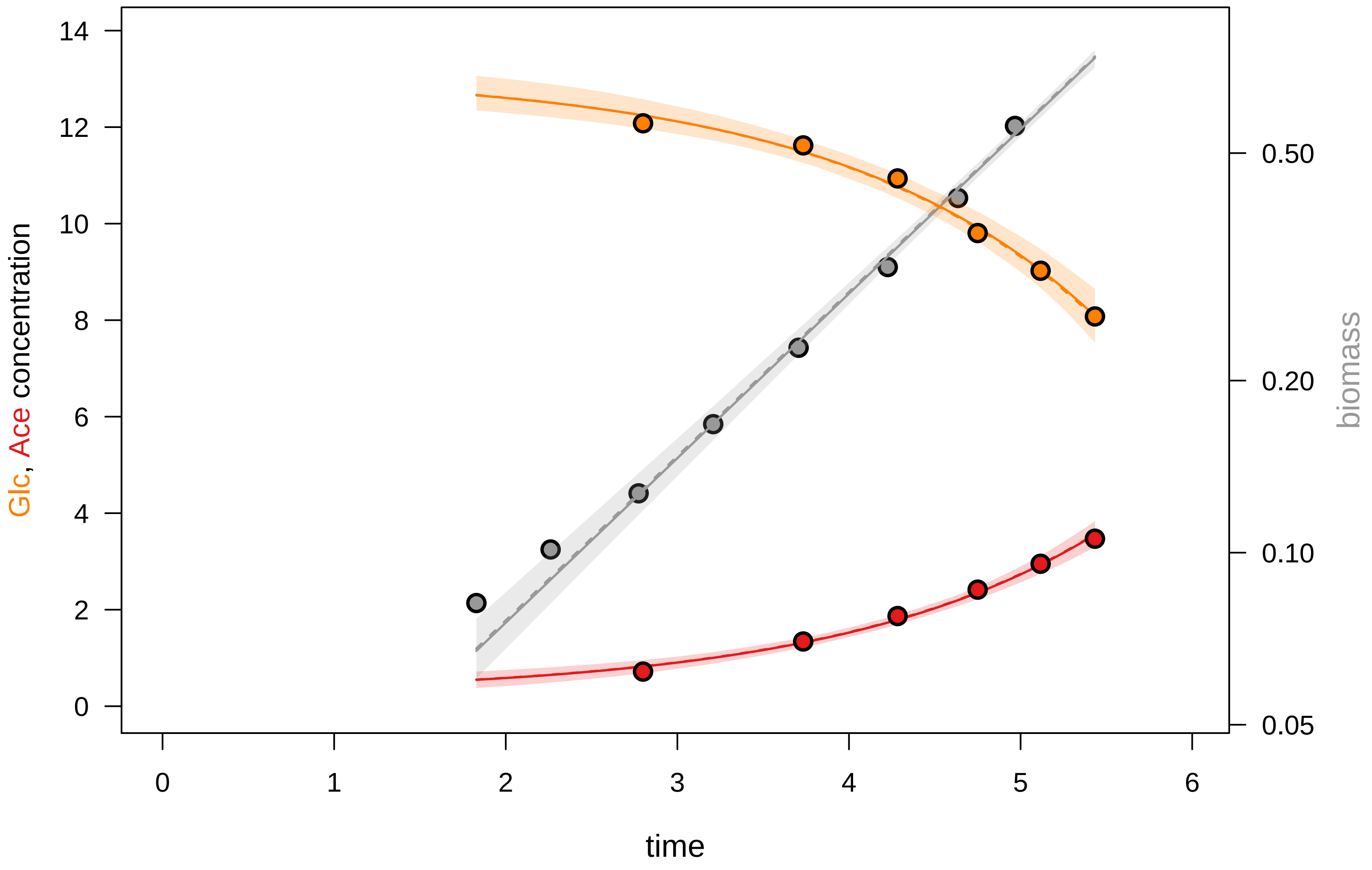

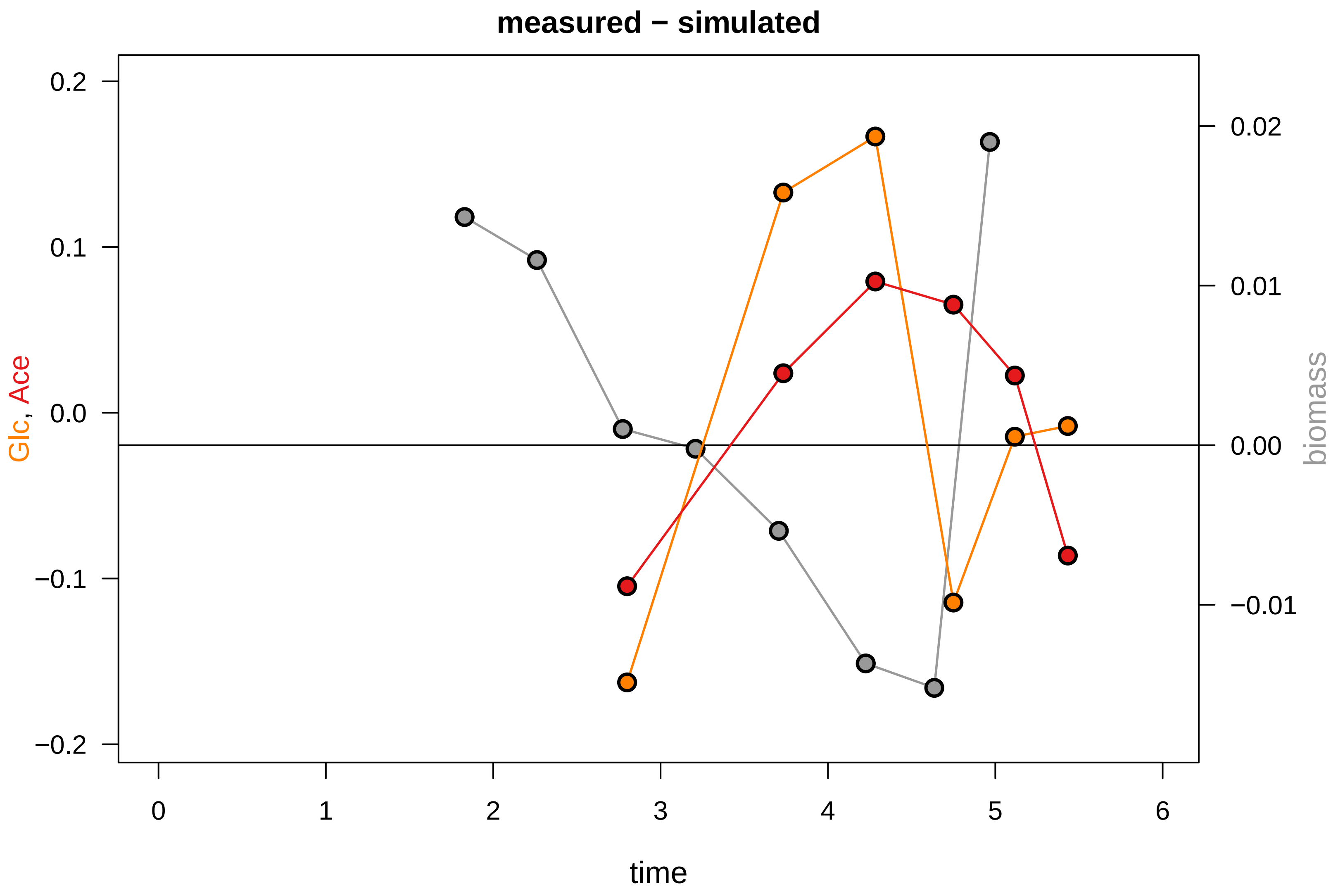

### KEIO_ROBOT5_15.pdf

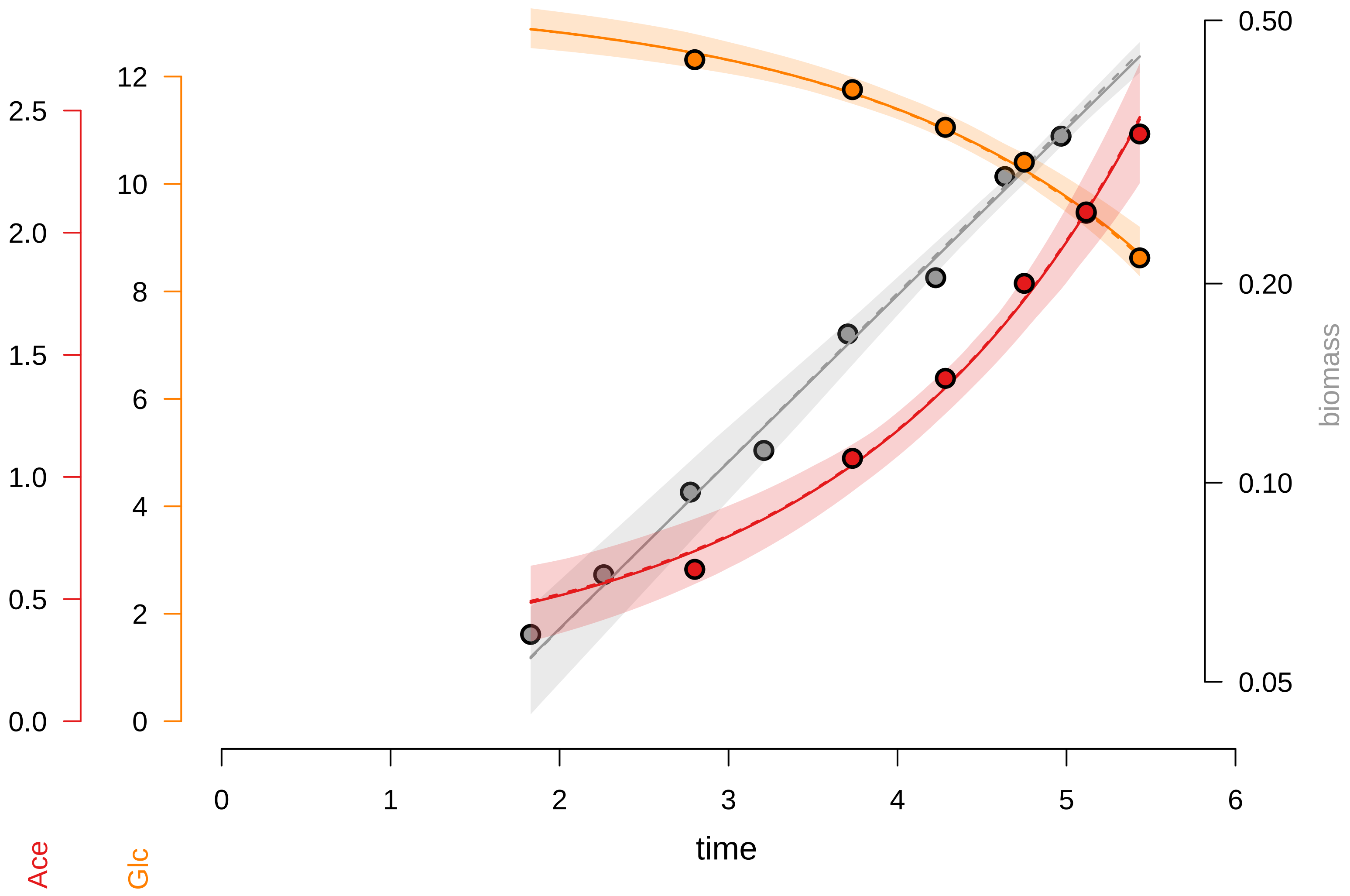

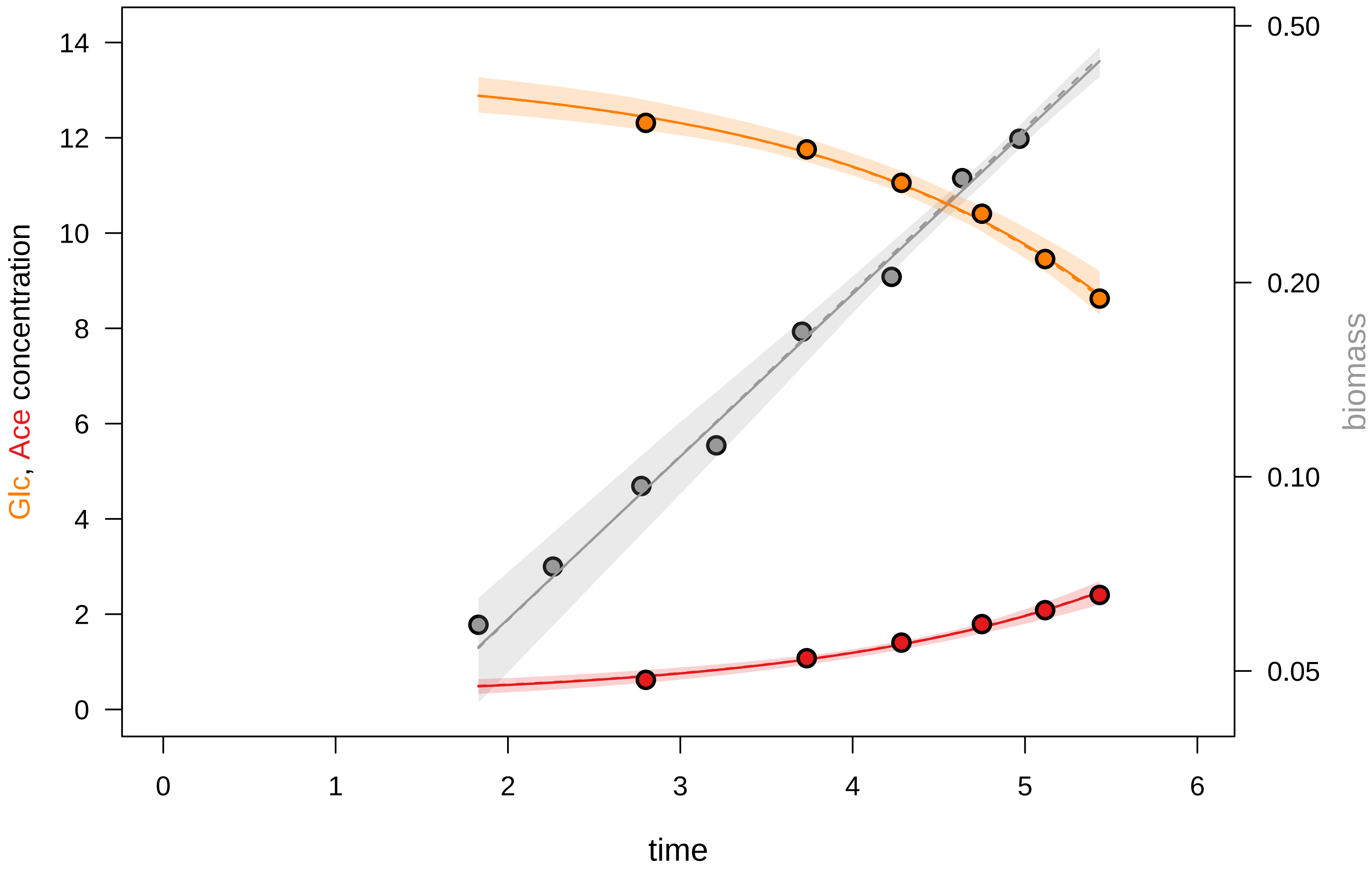

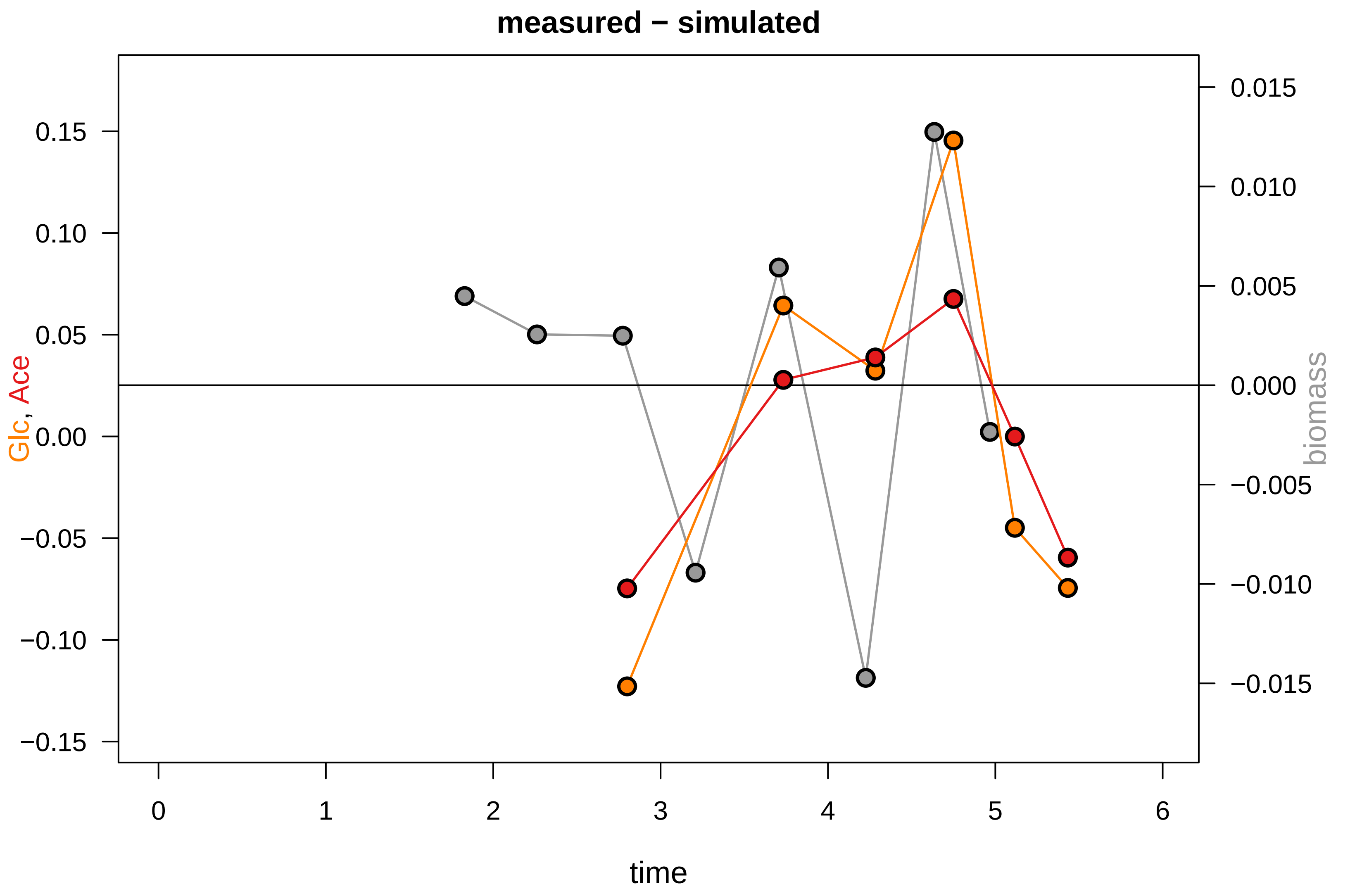

### KEIO_ROBOT5_17.pdf

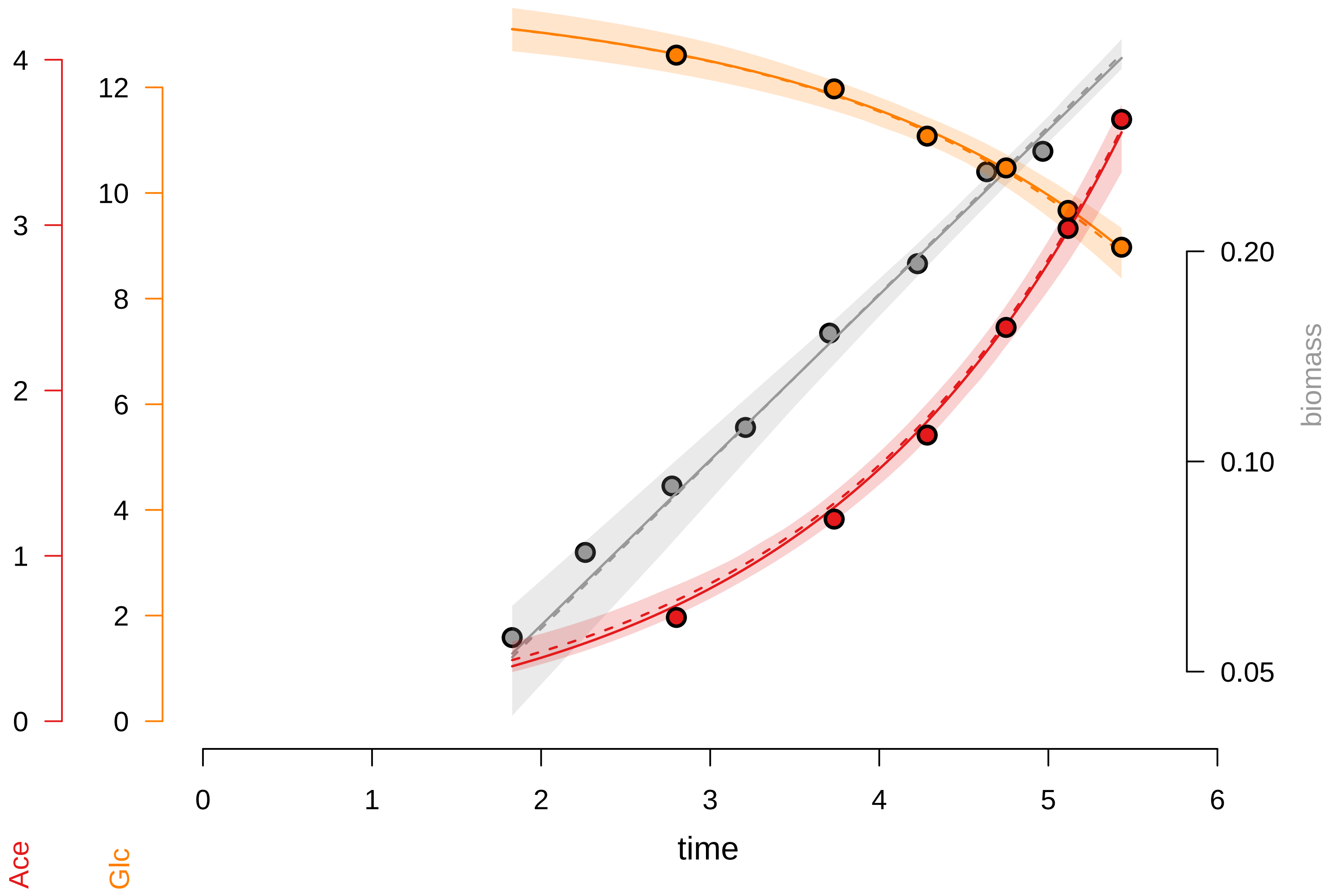

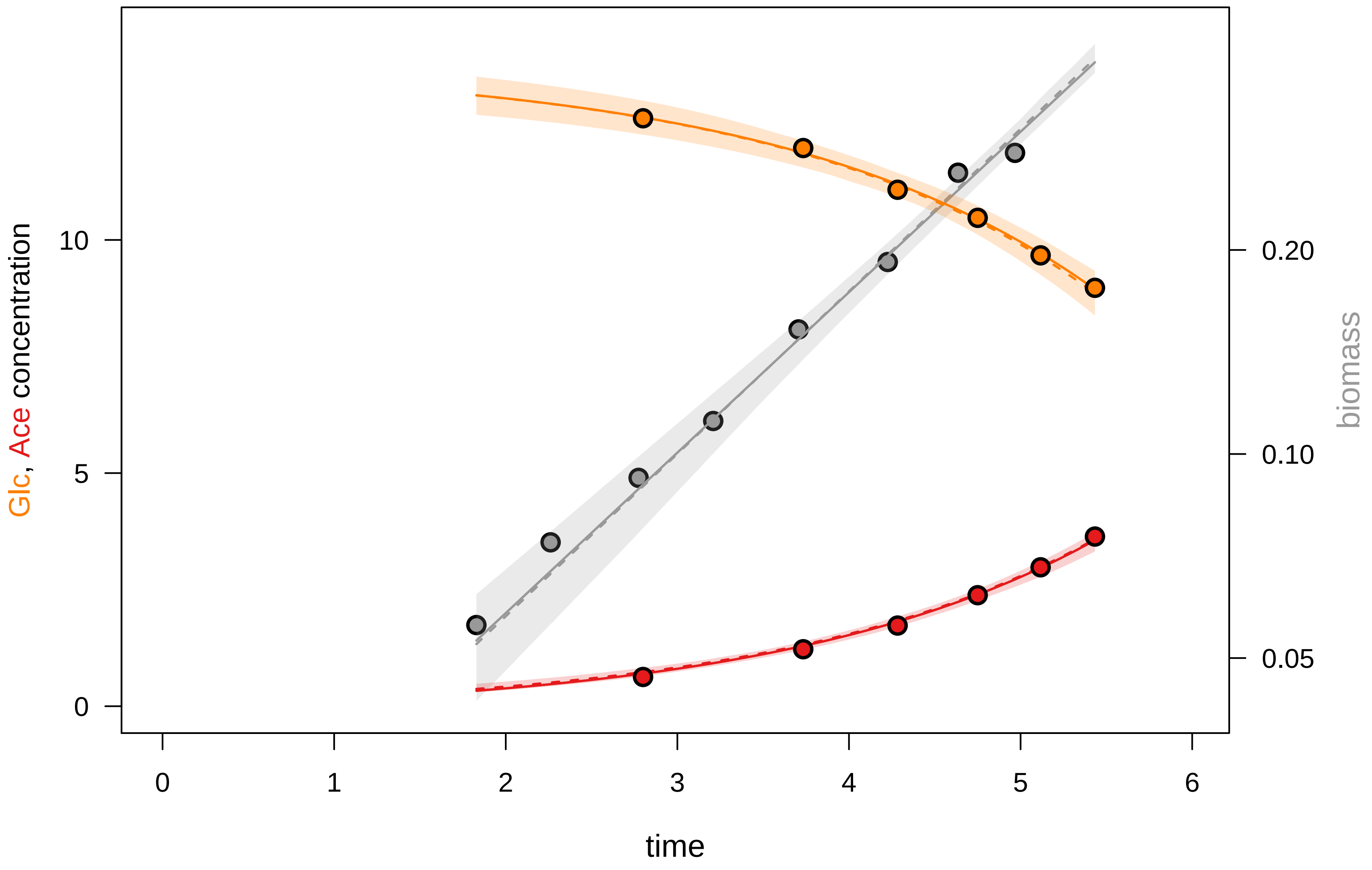

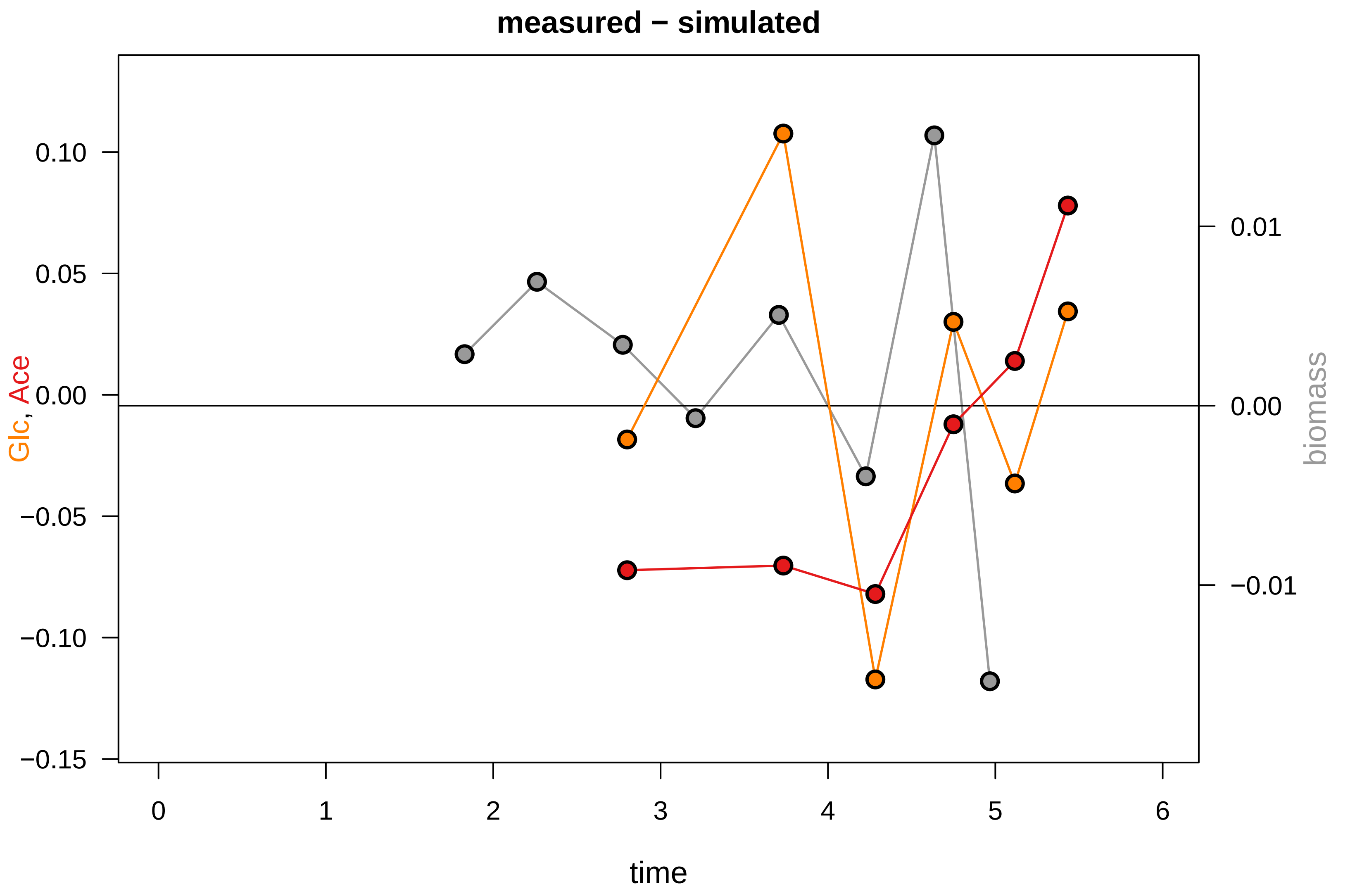

### KEIO_ROBOT5_48.pdf

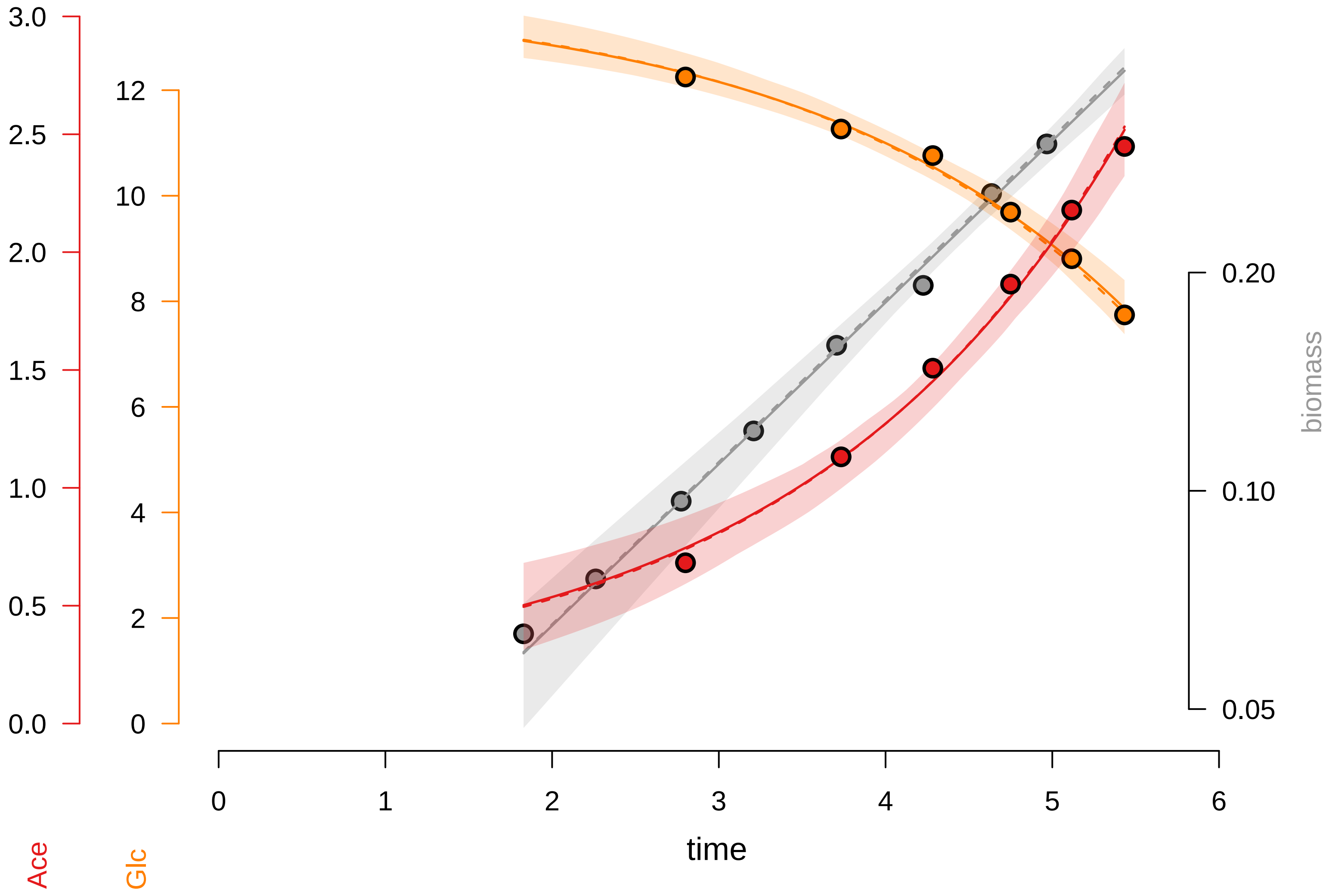

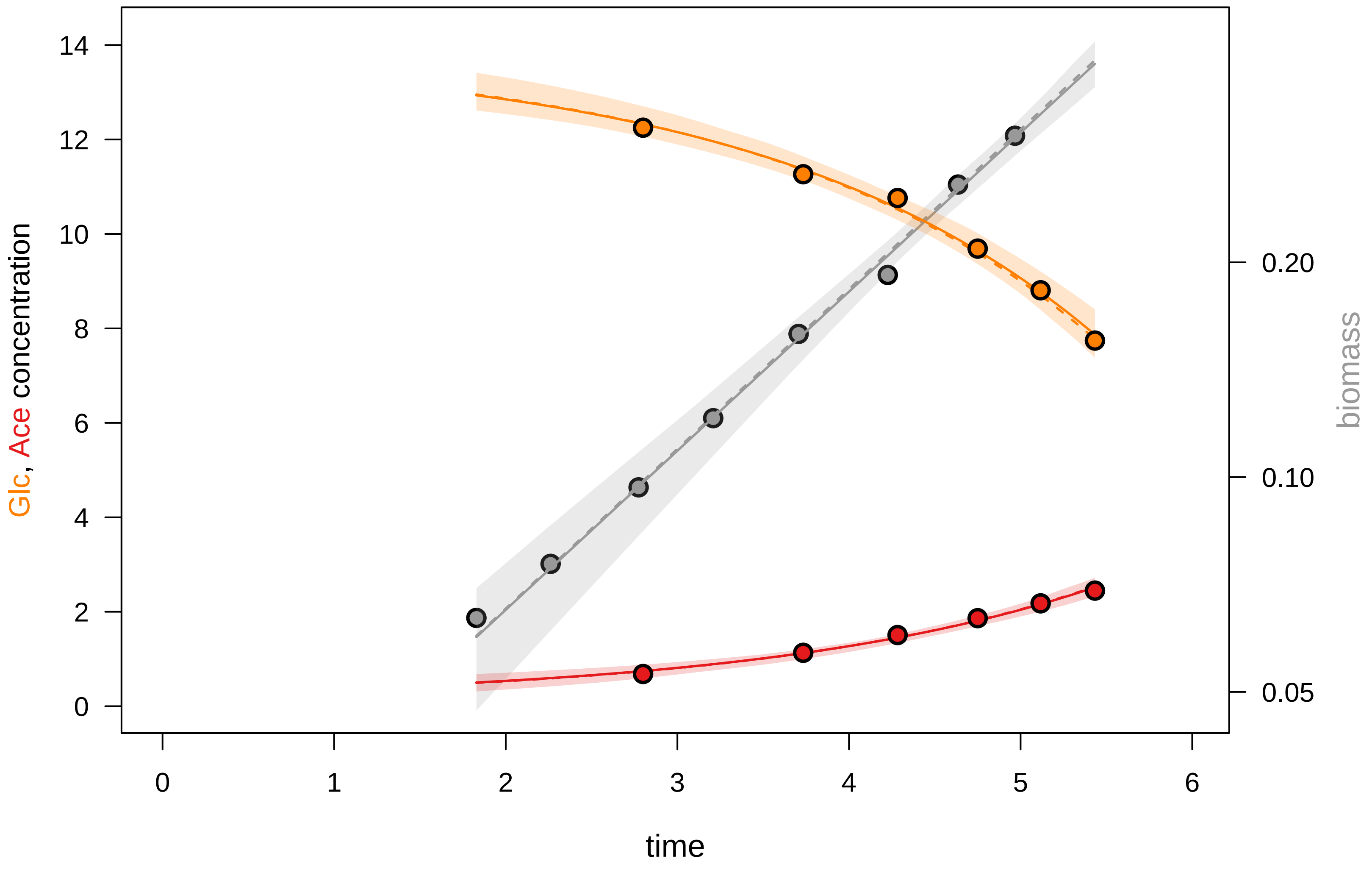

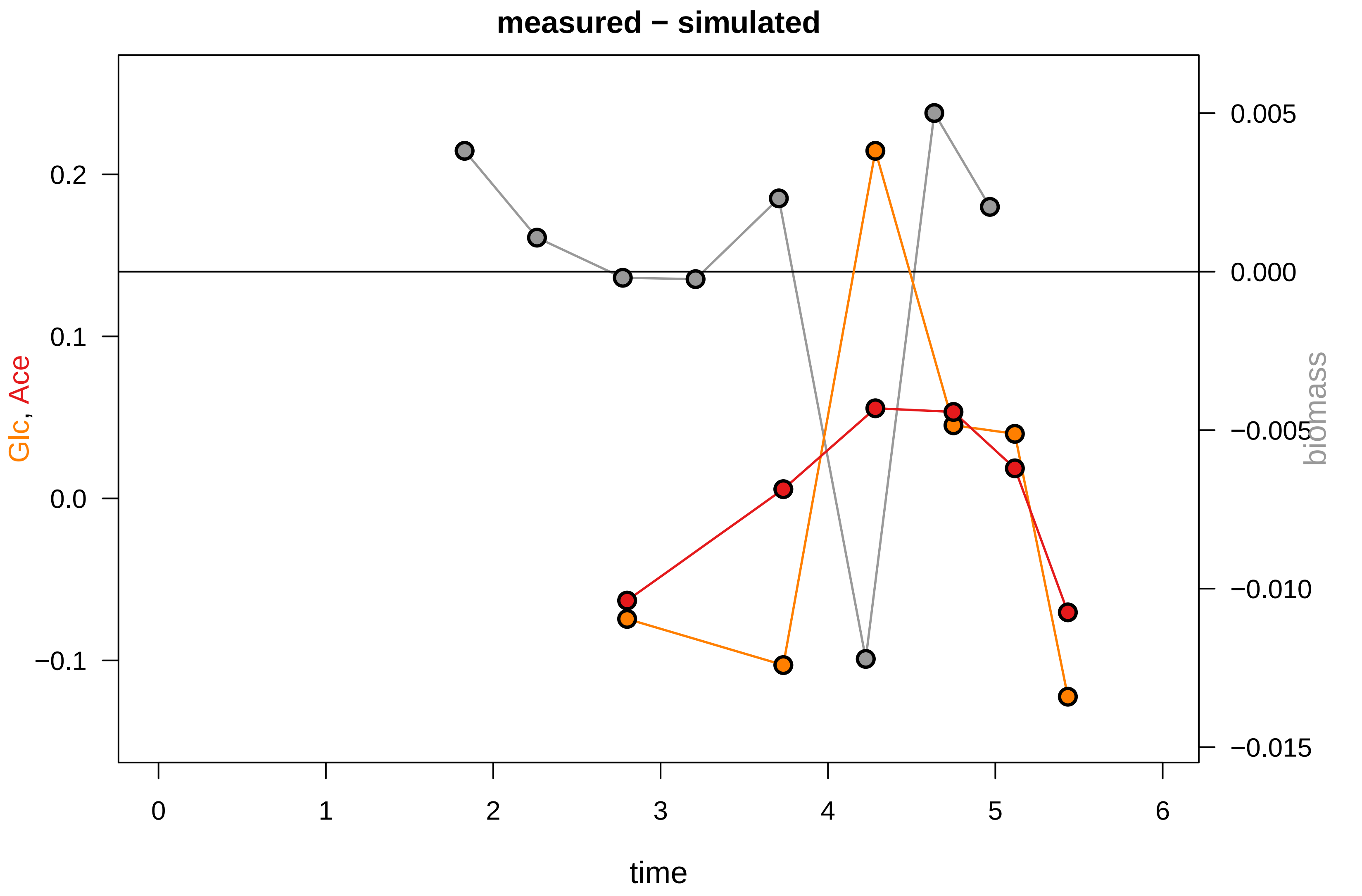

### KEIO_ROBOT6_1.pdf

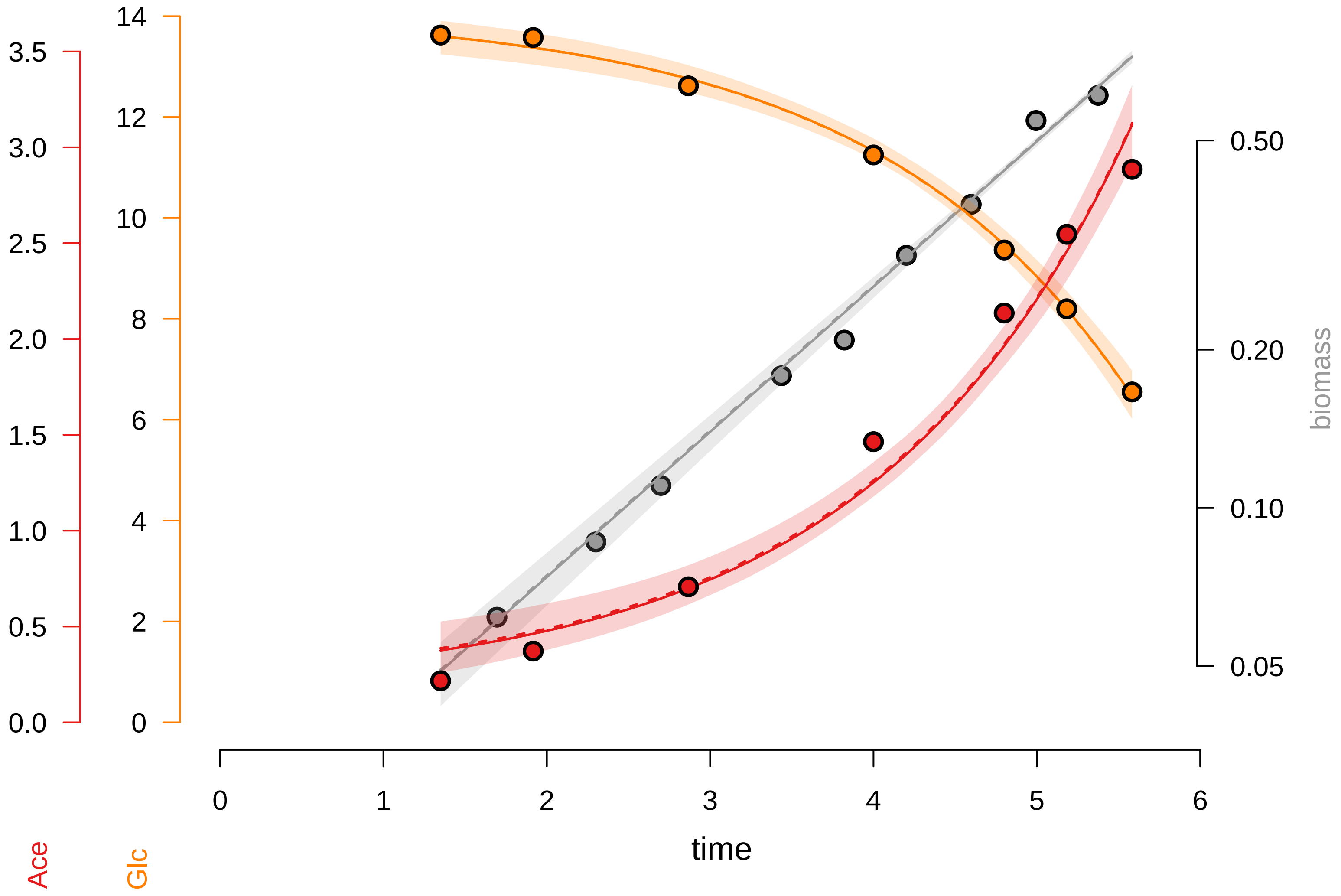

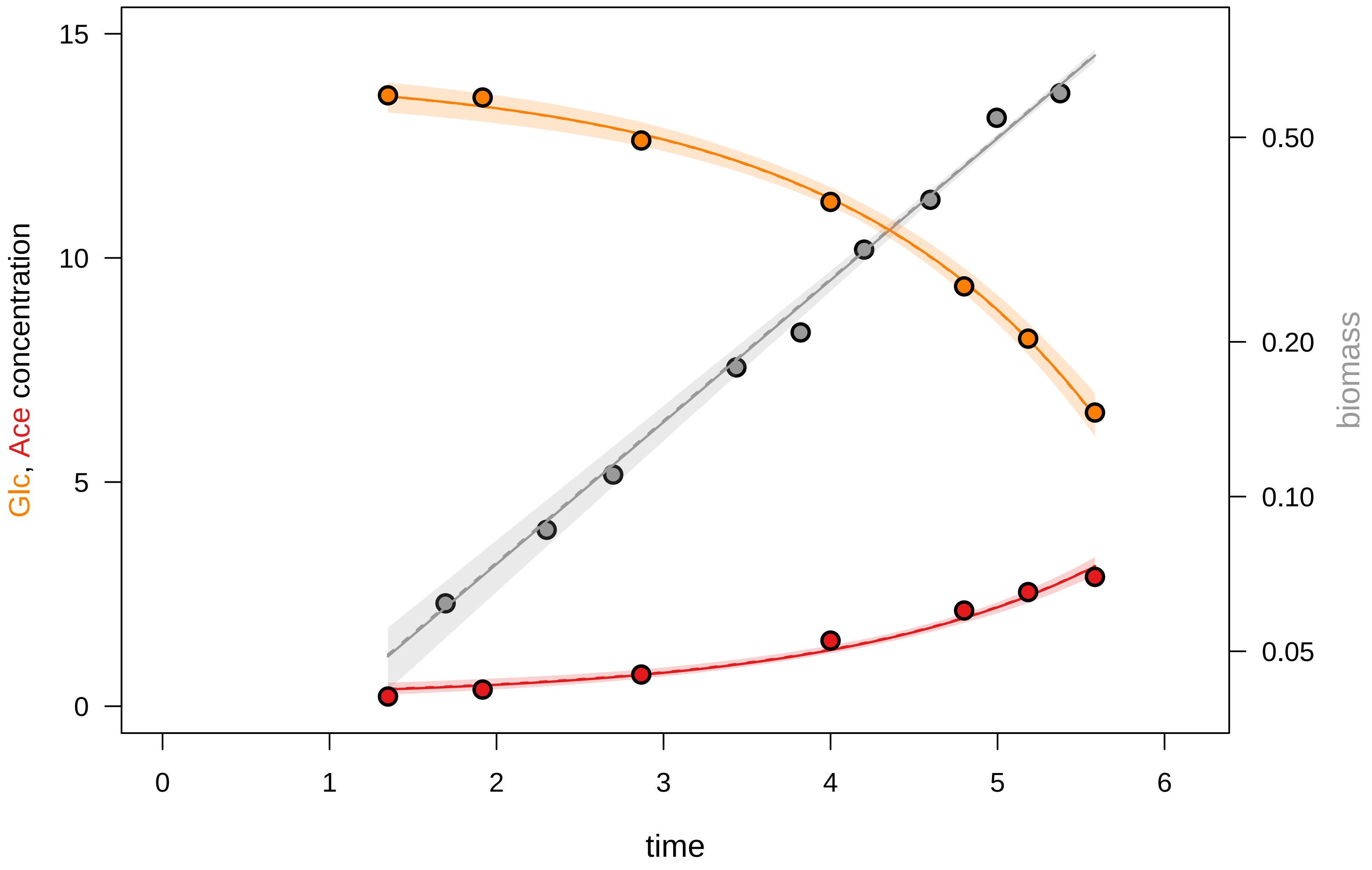

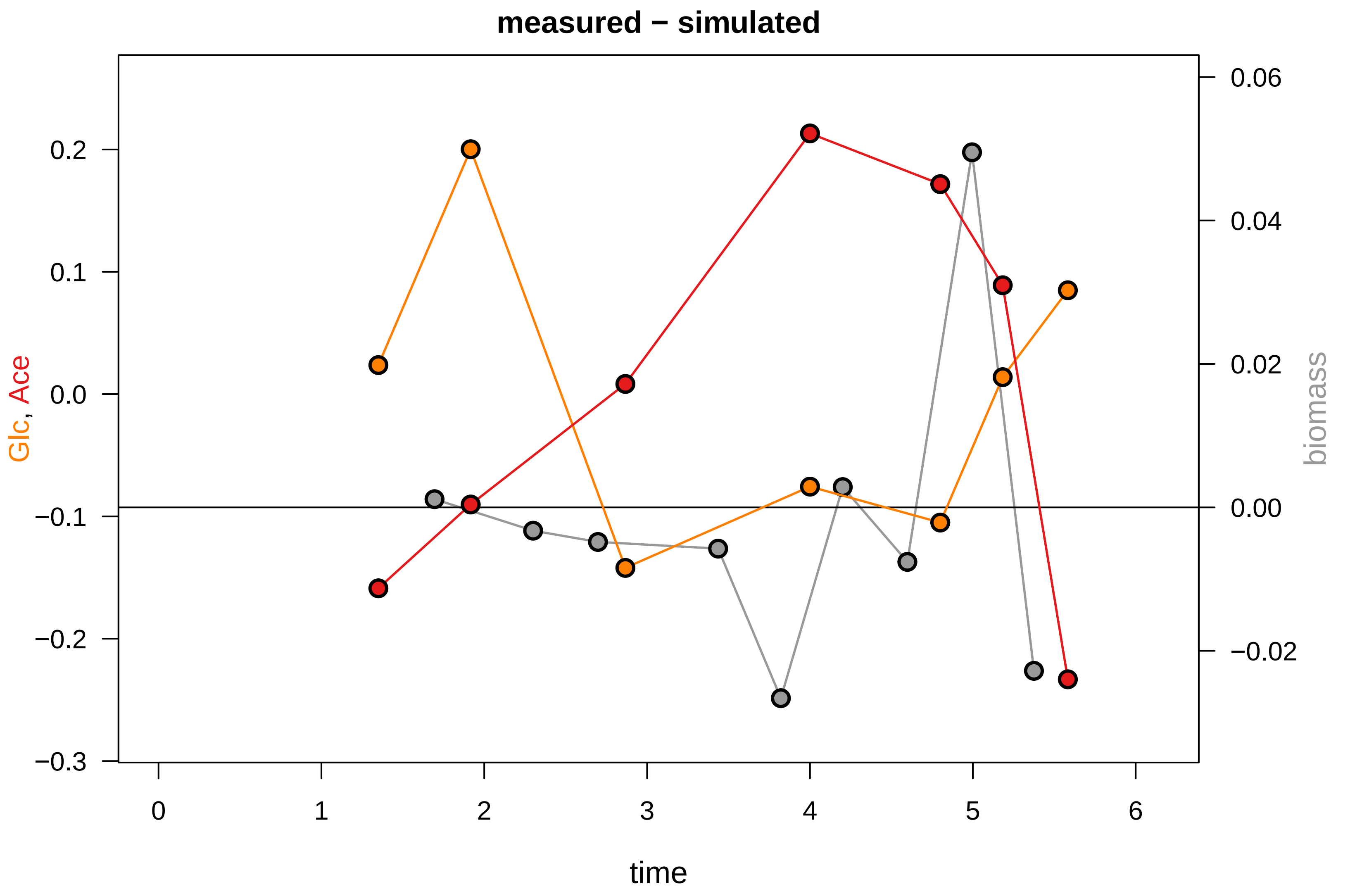

### KEIO_ROBOT6_2.pdf

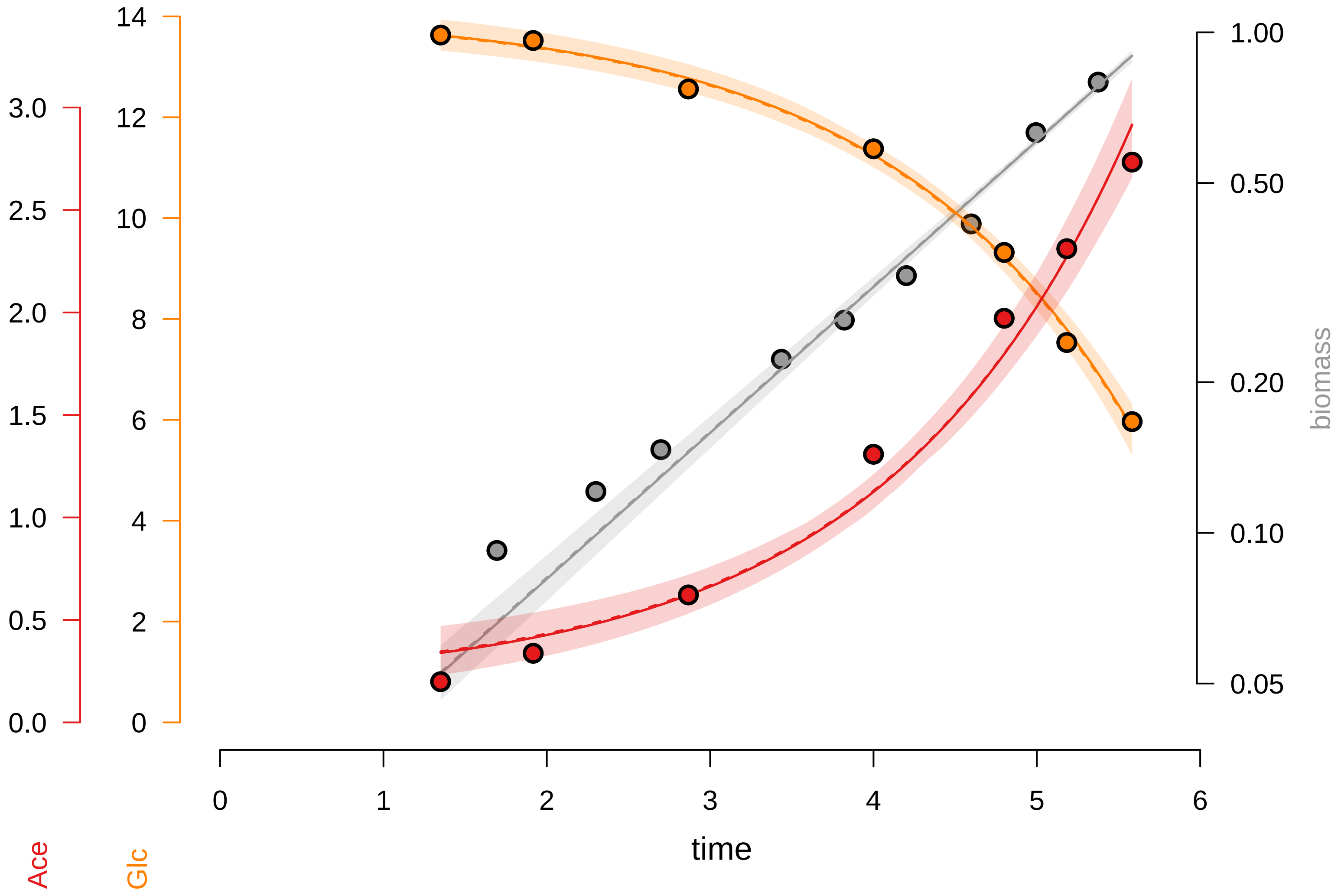

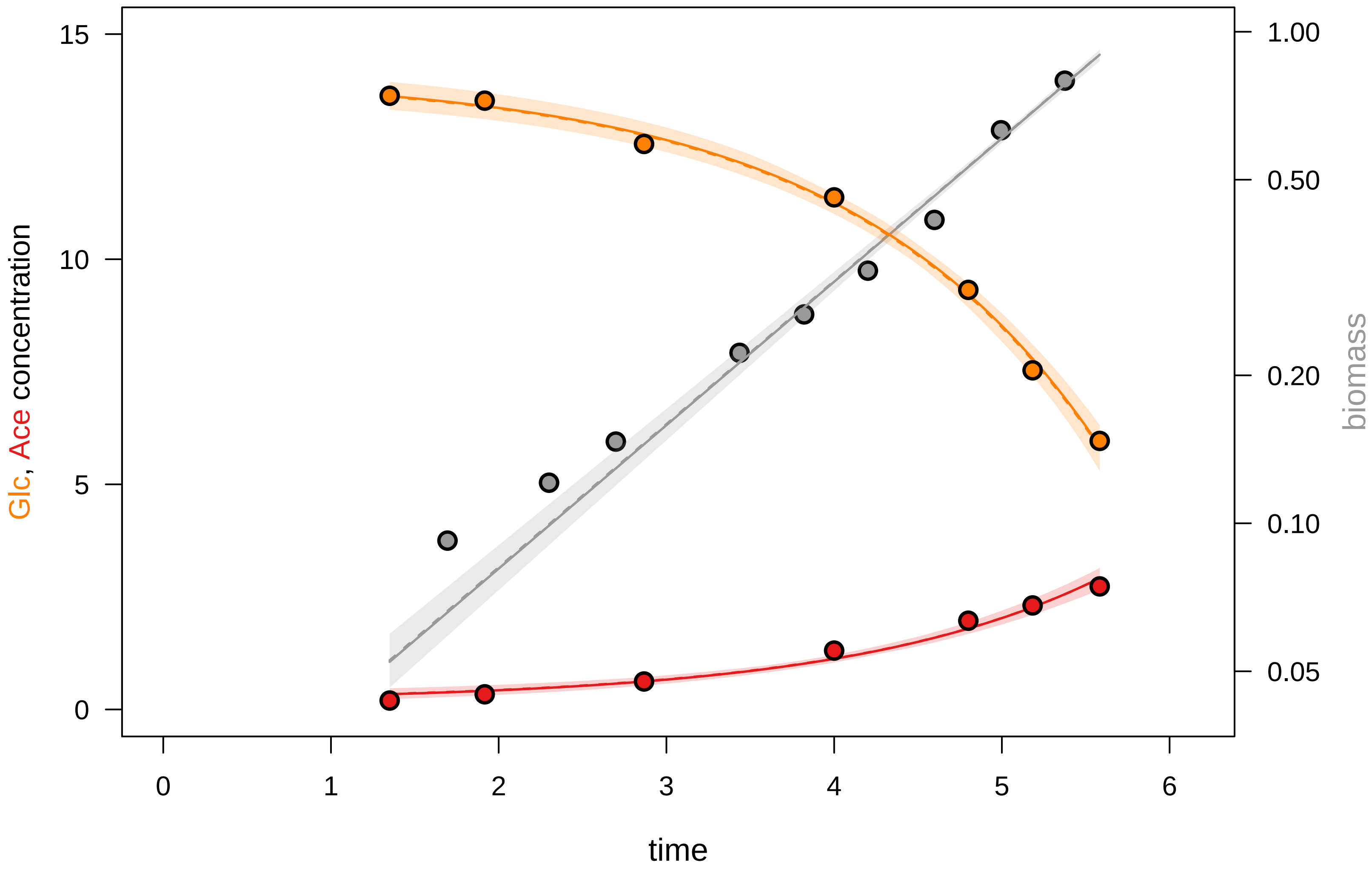

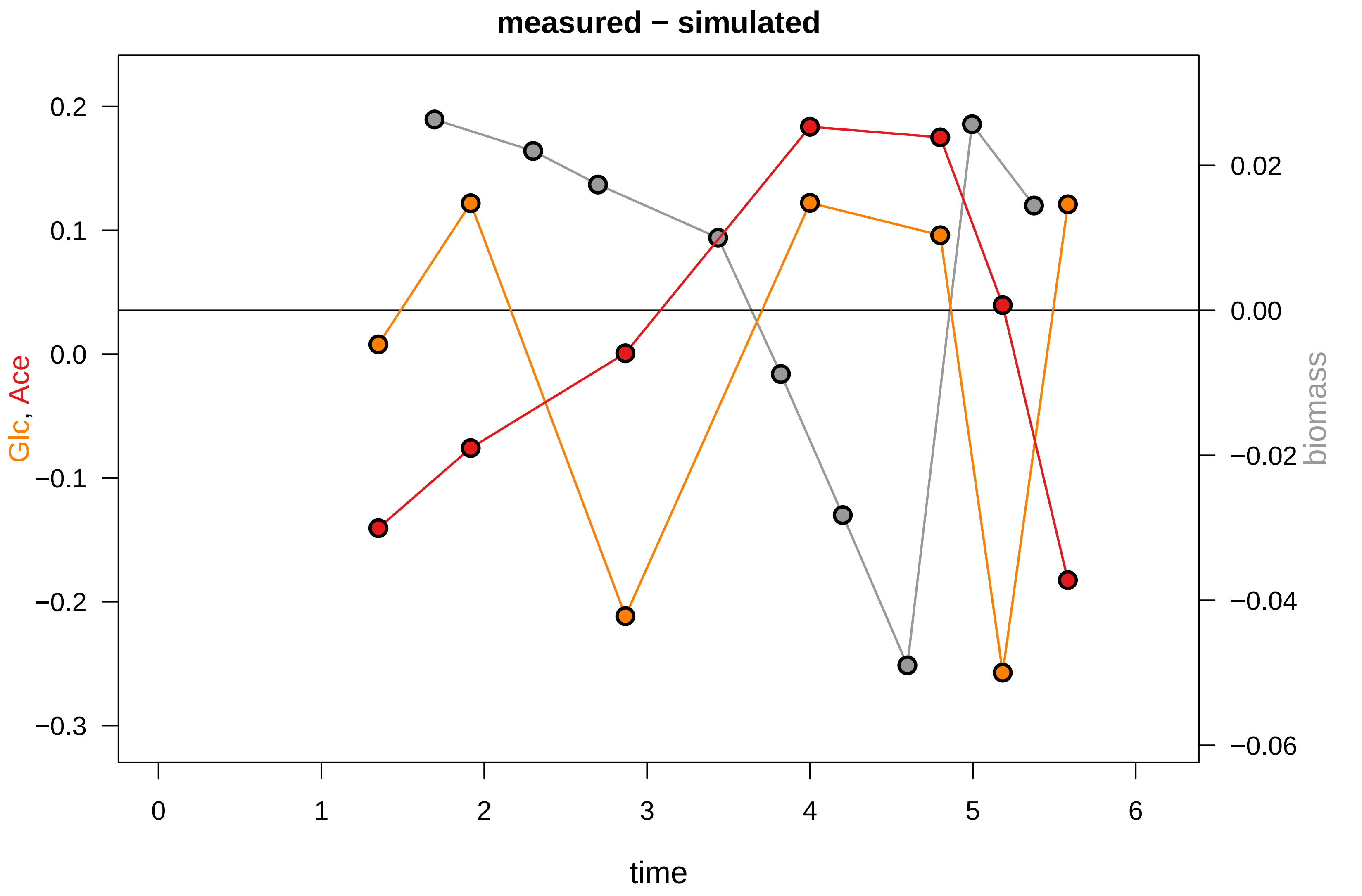

### KEIO_ROBOT6_3.pdf

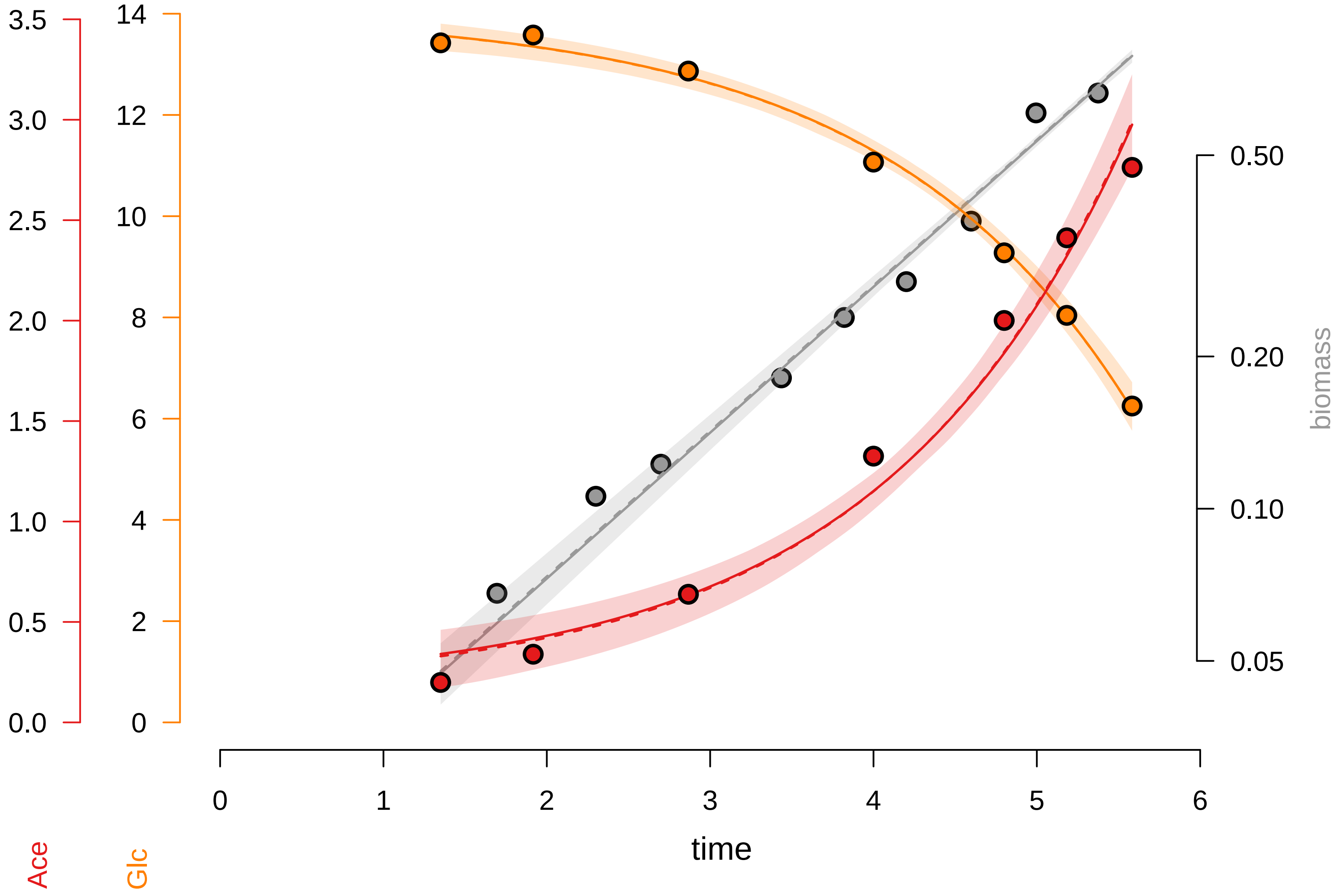

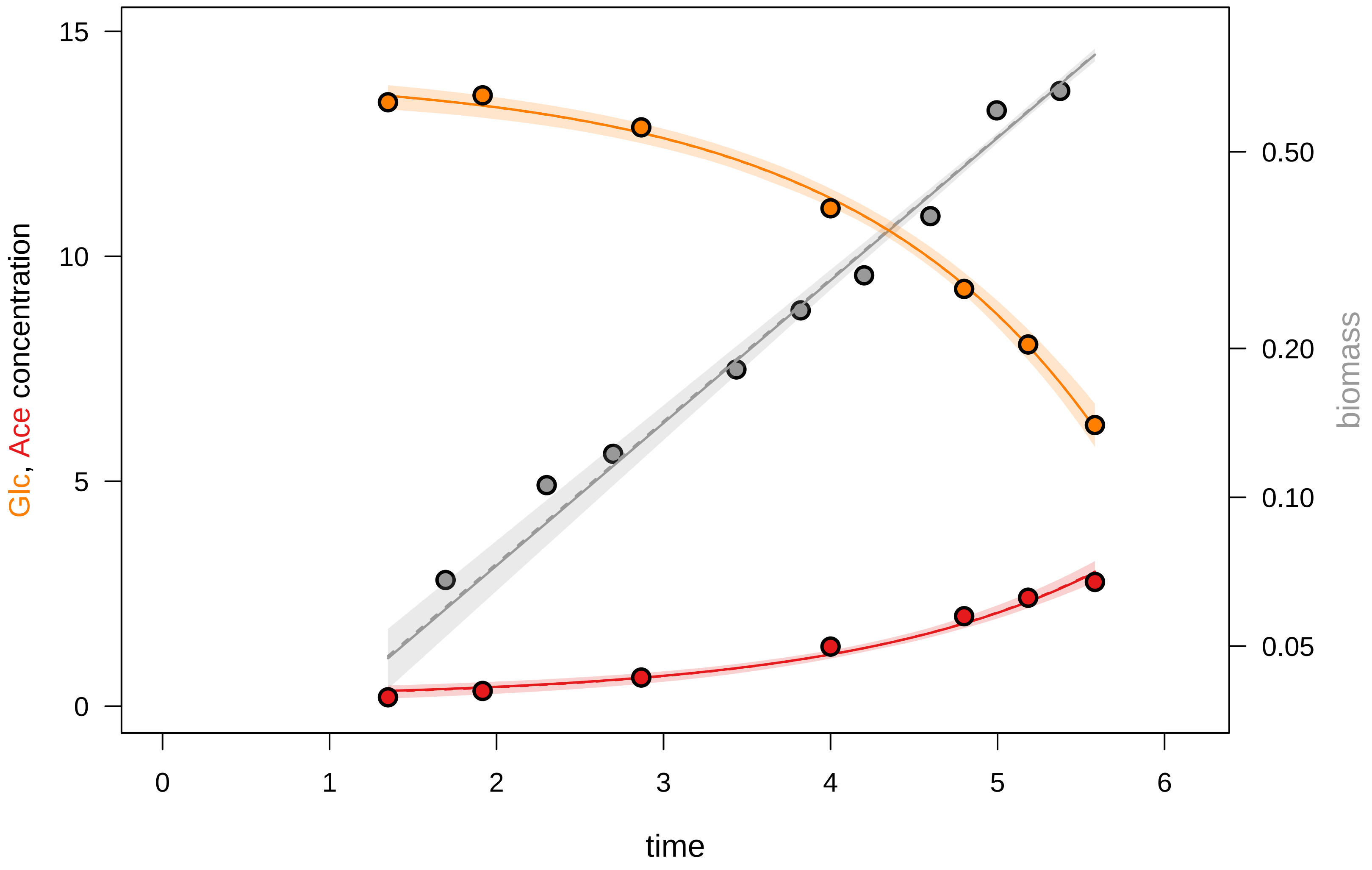

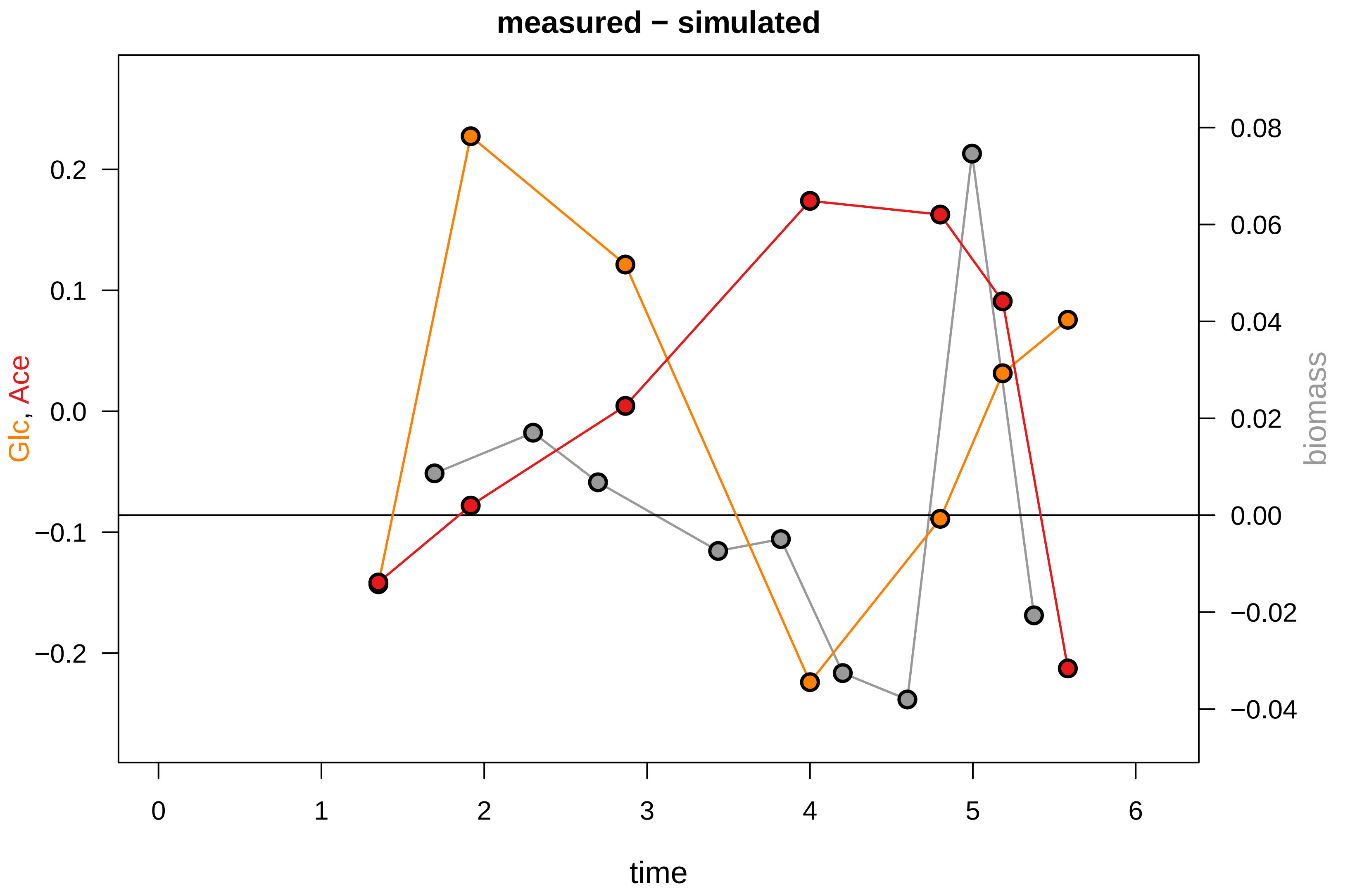

### KEIO_ROBOT6_4.pdf

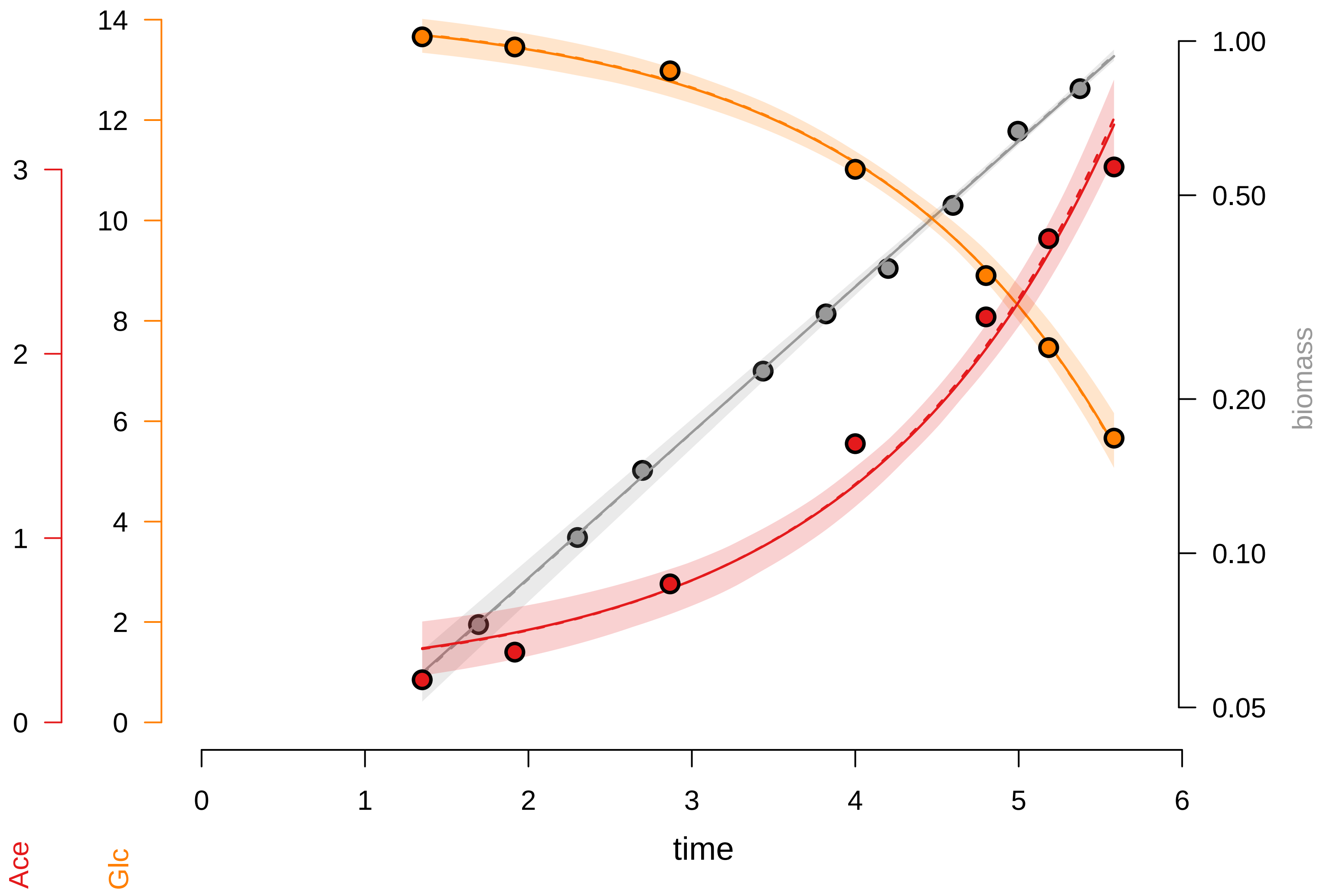

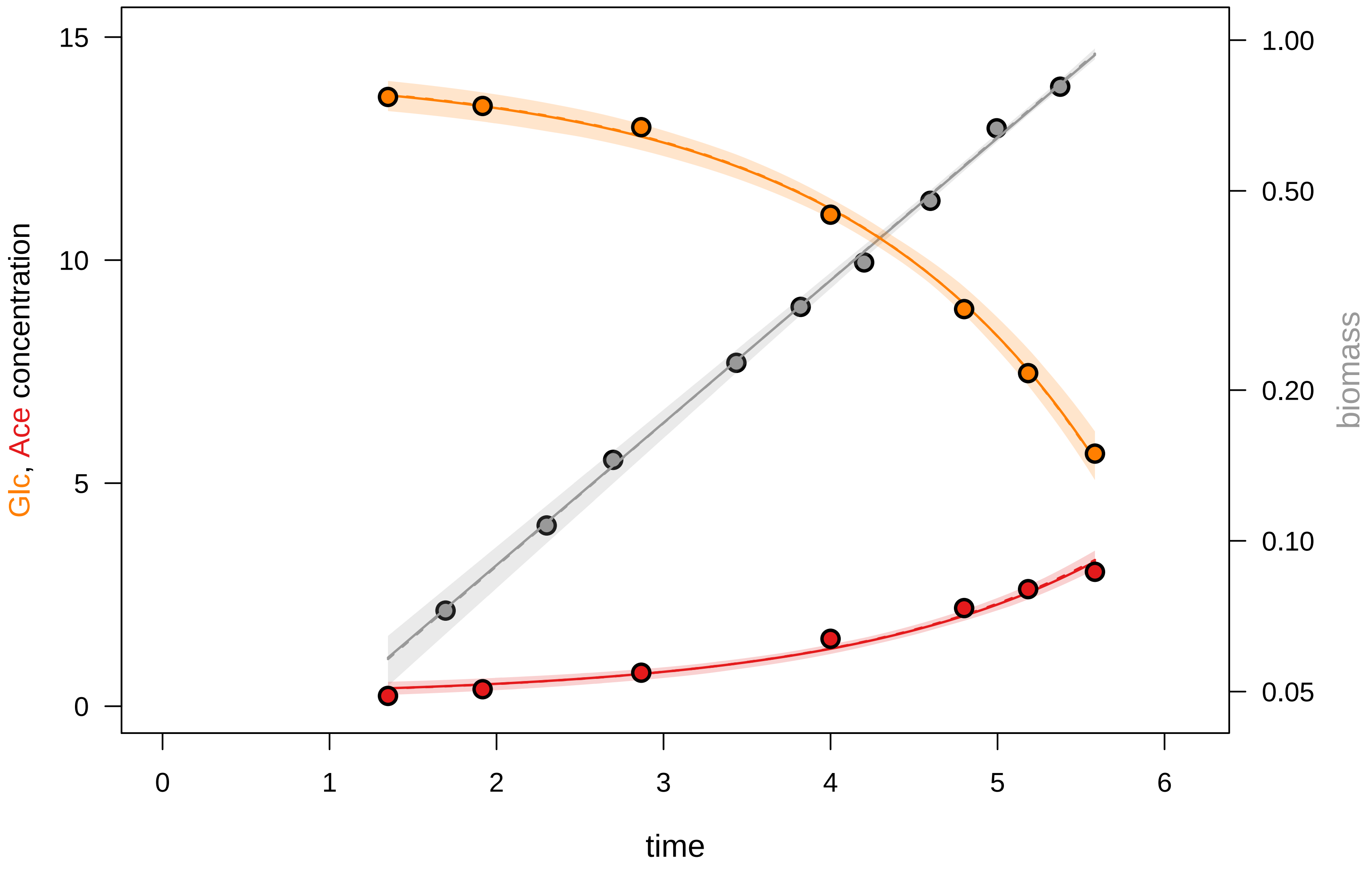

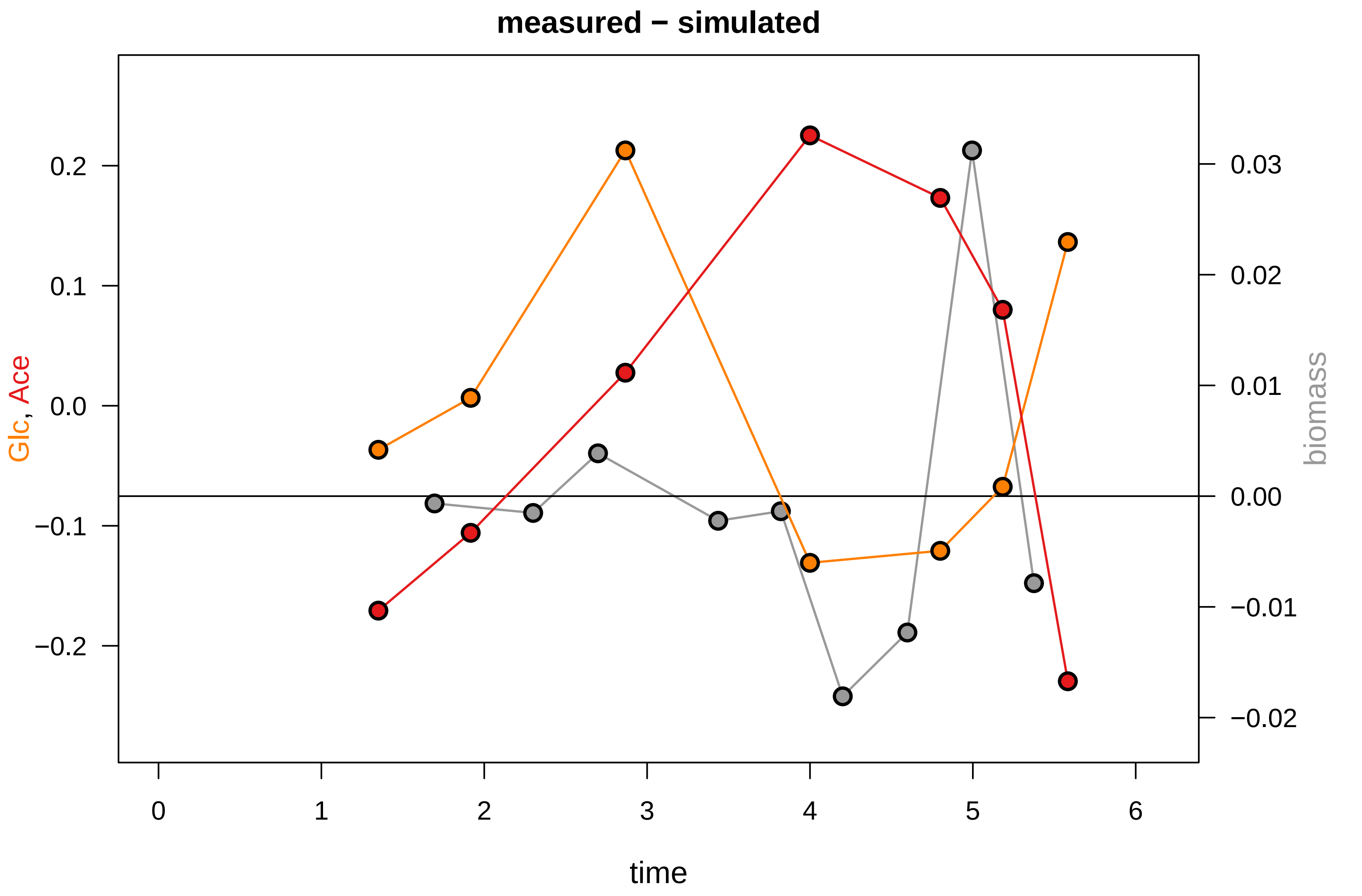

### KEIO_ROBOT6_5.pdf

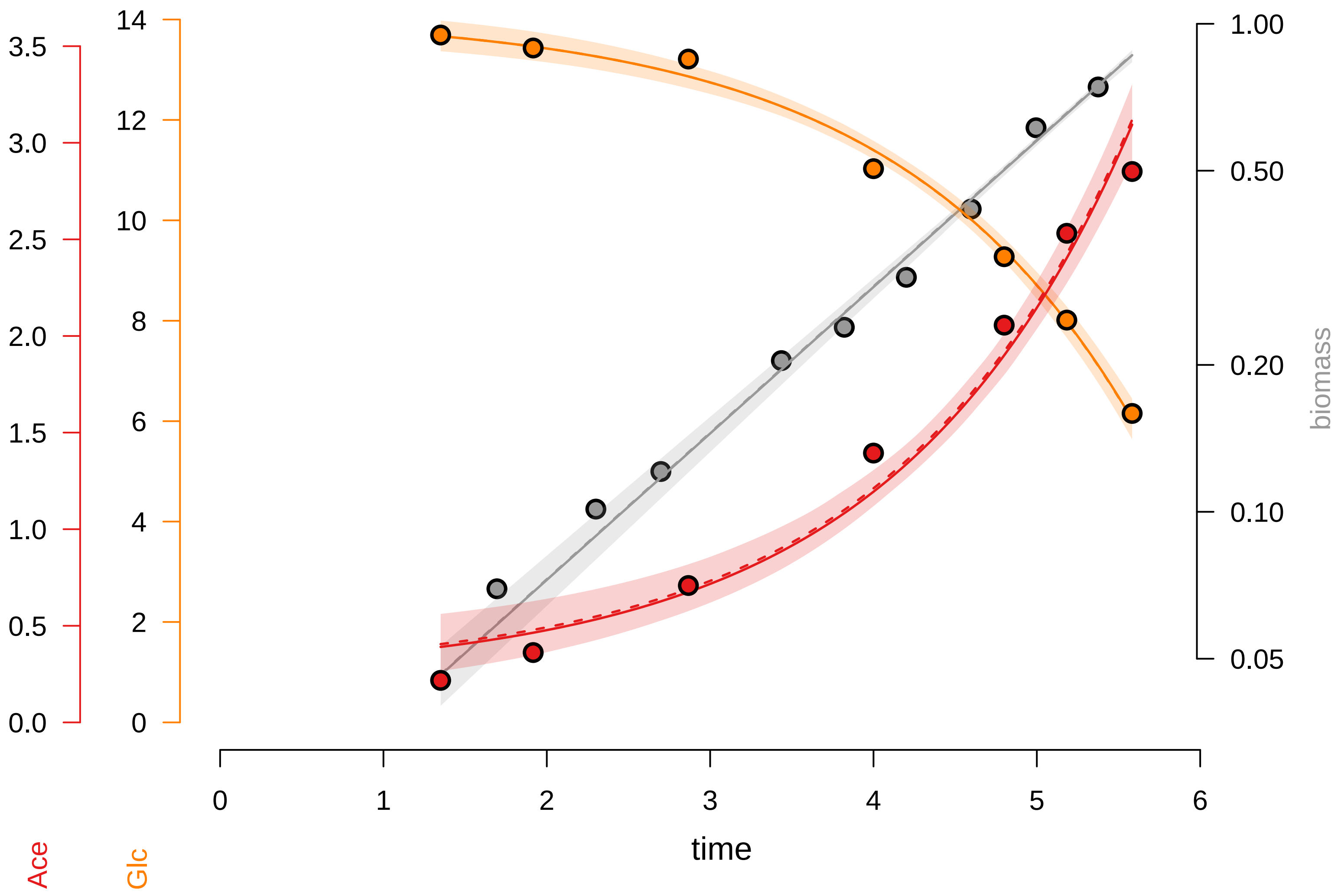

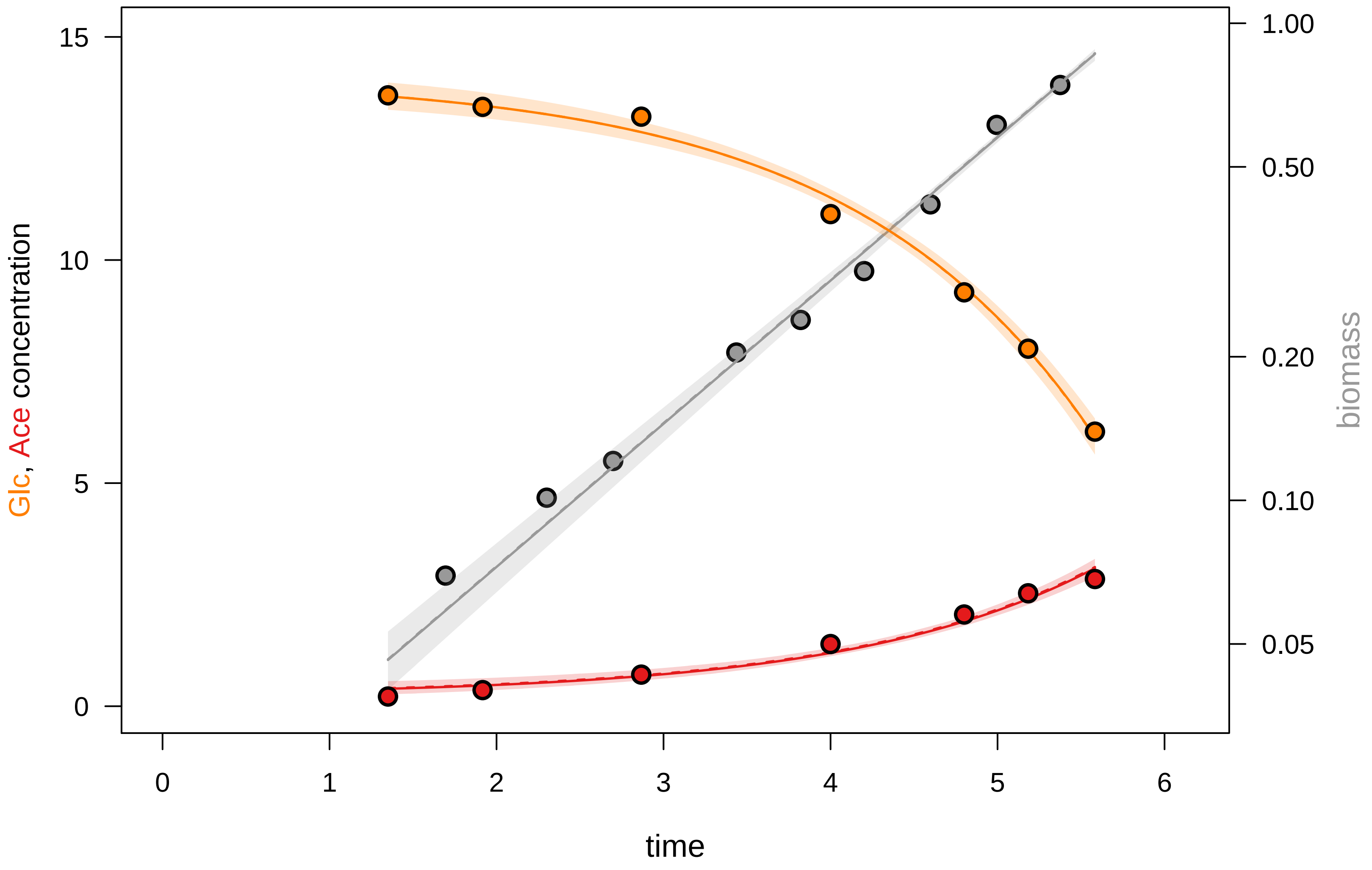

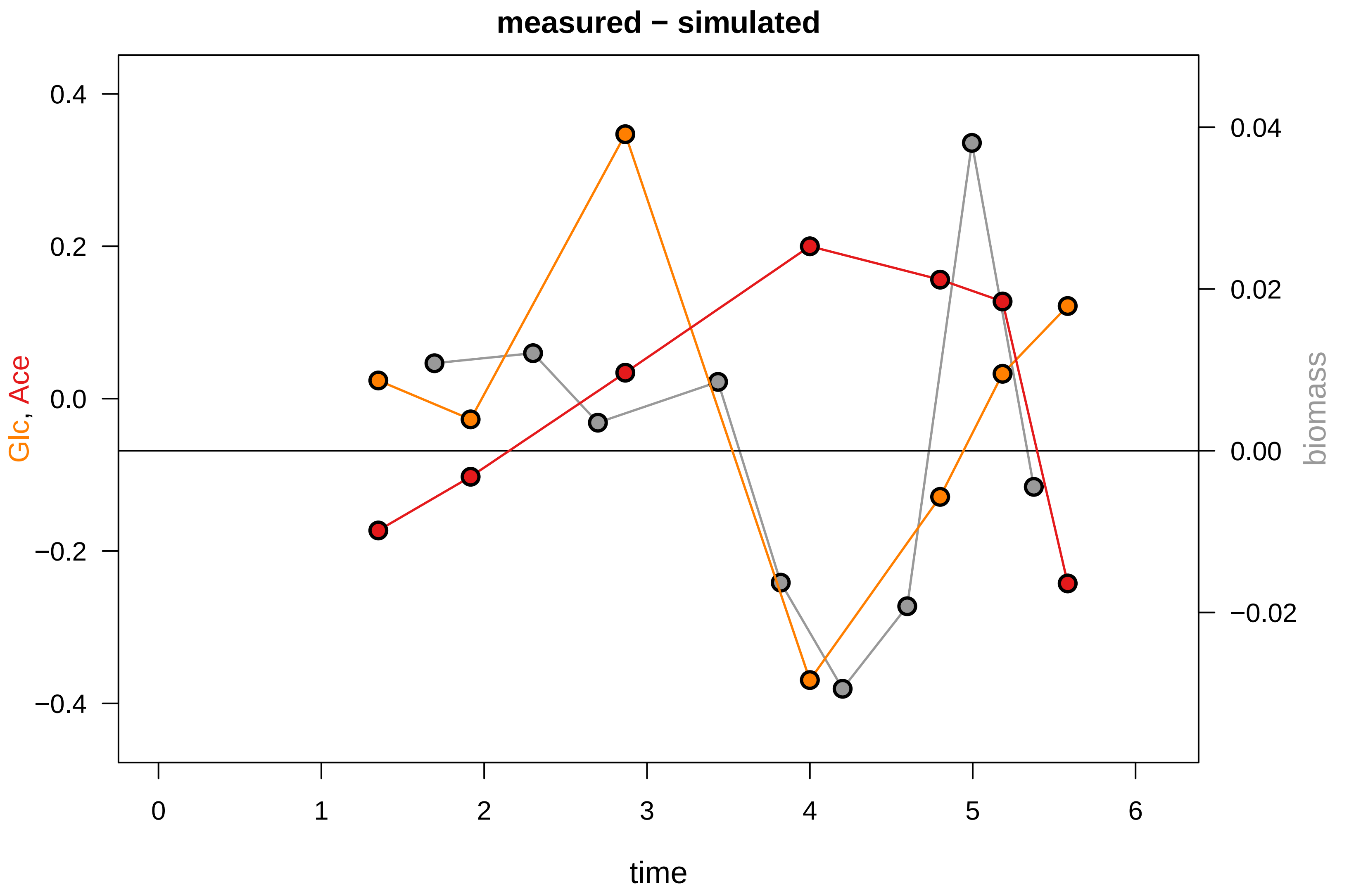

### KEIO_ROBOT6_6.pdf

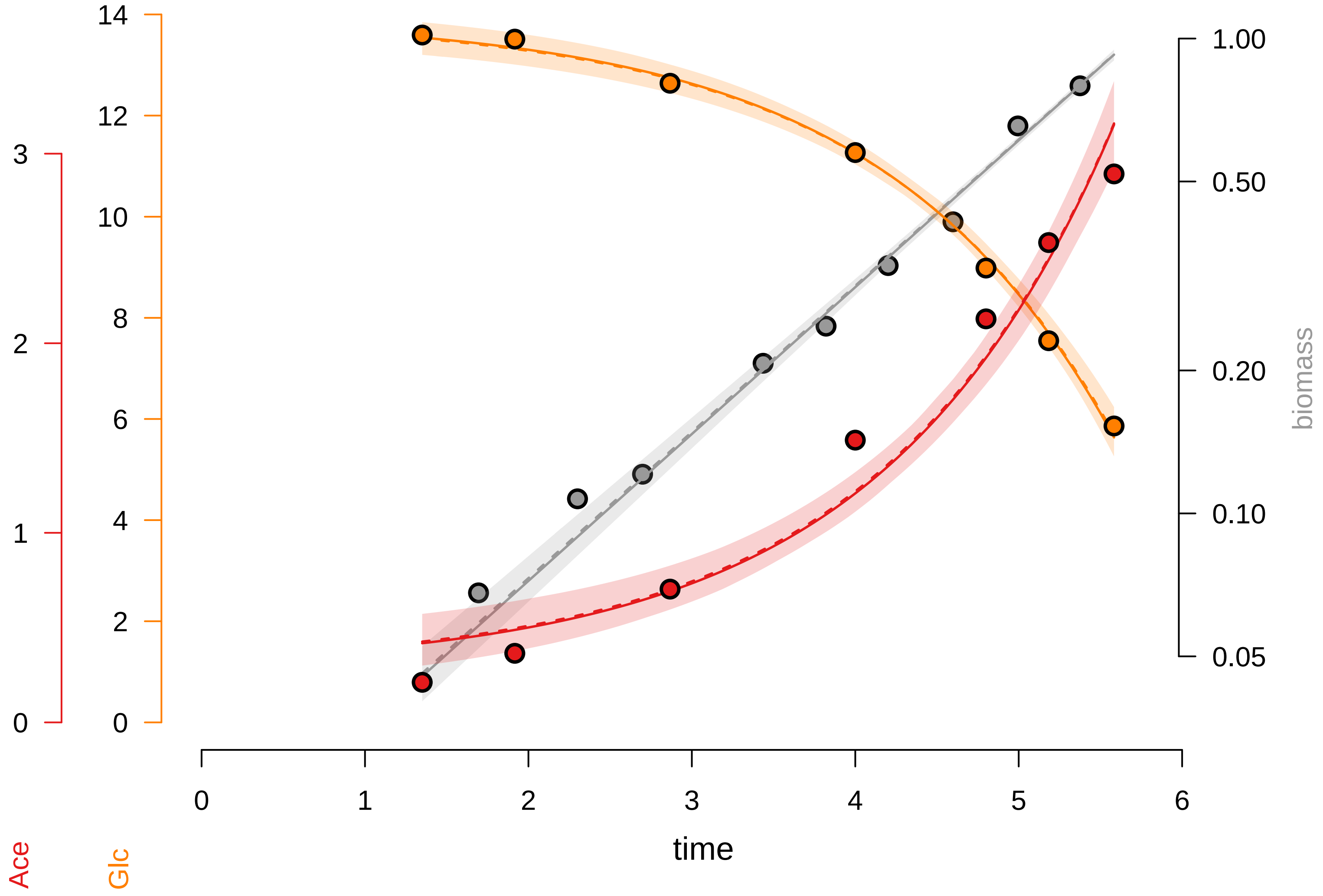

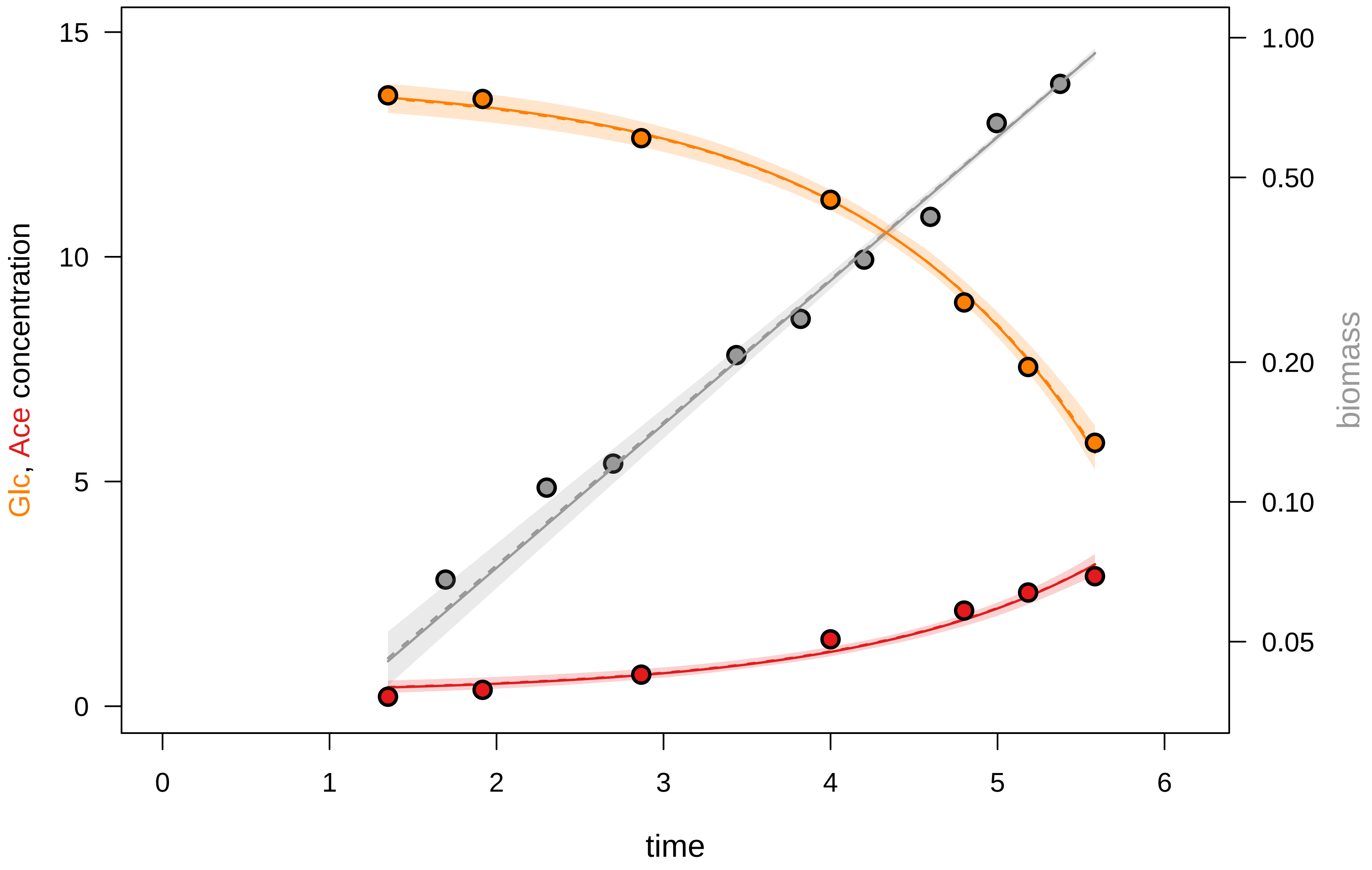

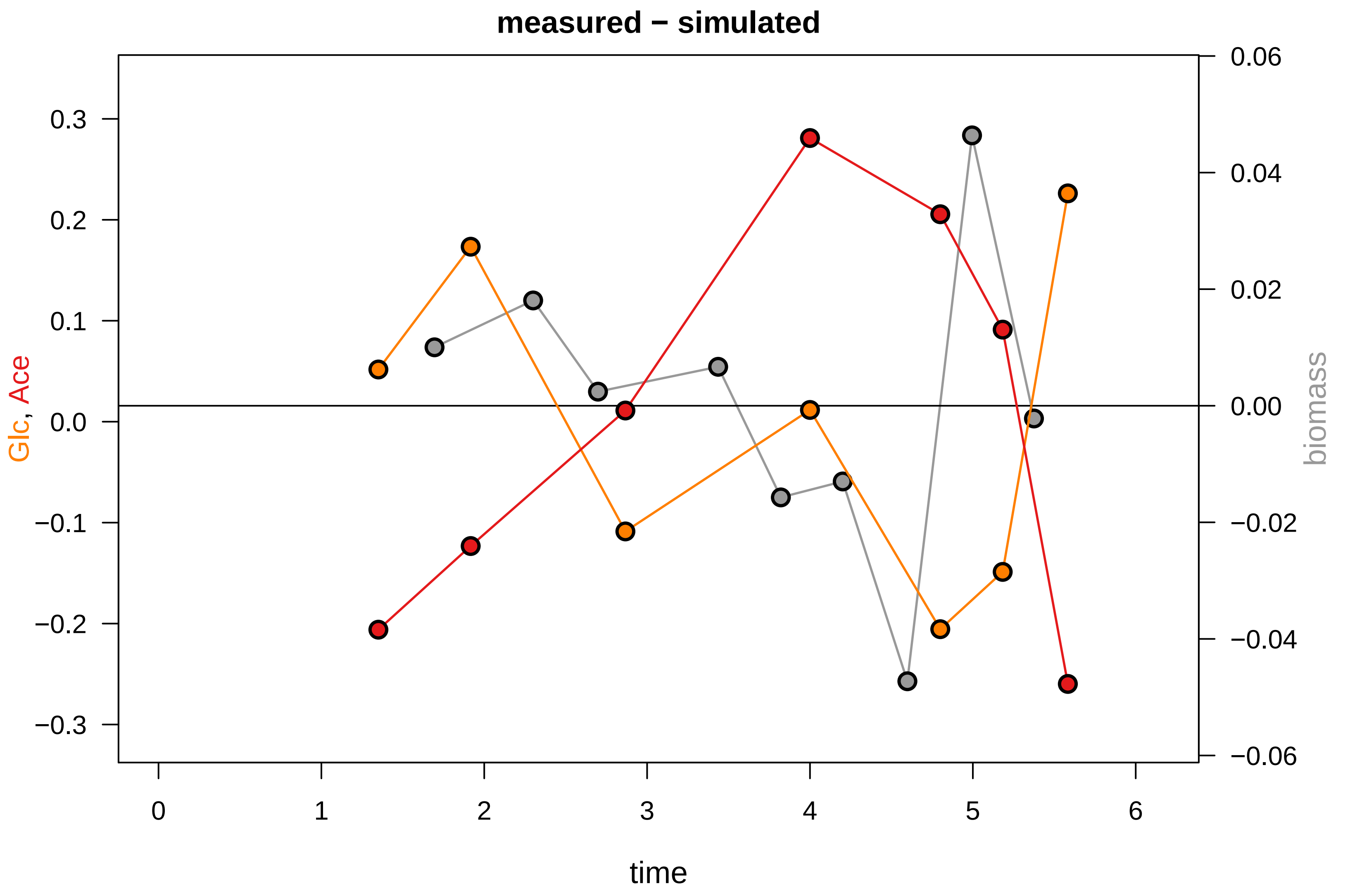
